# A Stepwise Thiol Dioxygenation Mechanism in Mercaptosuccinate Dioxygenase Revealed by A Combined Experimental and Computational Study

**DOI:** 10.64898/2026.05.19.726356

**Authors:** Stephanie Jordan, Hannah Ralls, Henrik P. H. Wong, Jordan A. Ernst, Todd C. Harrop, Sam P. de Visser, Yifan Wang

## Abstract

Thiol dioxygenases (TDOs) catalyze the incorporation of molecular oxygen into thiol metabolites and *N*-terminal cysteine residues of regulatory proteins, thereby playing critical roles in sulfur metabolism and oxygen sensing. Despite extensive study over the past two decades, the molecular basis for substrate recognition and the catalytic mechanism of TDOs remains controversial, owing to the scarcity of substrate-bound structures and direct evidence for catalytic intermediates. Herein, we present a comprehensive study of mercaptosuccinate dioxygenase (MSDO), a TDO originally identified in *Variovorax paradoxus* B4, using a combination of structural, biochemical, spectroscopic, and computational approaches. MSDO oxidizes both (*S*)- and (*R*)-mercaptosuccinate (MS) with similar *K*_m_ values but exhibits approximately 2.5-fold higher turnover for the (*S*)-enantiomer. Crystal structures of MSDO reveal that both (*S*)- and (*R*)-MS coordinate the iron in a bidentate mode via their thiolate and proximal carboxylate groups, with the distal carboxylate adopting distinct orientations. Two active-site Arg residues recognize the substrate carboxylate groups and thereby stabilize a flexible *C*-terminal loop, underpinning a catalytic site gating mechanism in MSDO. EPR spectroscopy corroborates bidentate coordination, showing conversion of a high-spin {FeNO}^7^ complex to a low-spin species upon substrate binding. Time-resolved *in crystallo* reactions capture two key iron-bound intermediates, namely an unprecedented monooxygenated sulfenate and a dioxygenated sulfinate product. These structural snapshots are supported by DFT calculations that point to a stepwise oxygen atom transfer pathway. Computational analysis further accounts for the kinetic differences between the substrate enantiomers, as rationalized by structural comparisons, active-site geometry, and second coordination sphere interactions. Together, these results elucidate fundamental principles of TDO catalysis and advance our understanding of nonheme iron-dependent oxygen activation.

## Introduction

Thiol dioxygenases (TDOs) are mononuclear nonheme iron-dependent enzymes found across diverse organisms.^1-3^ These enzymes catalyze the incorporation of molecular oxygen into thiol-containing molecules to form sulfinic acids. They feature a conserved 3-His iron-binding motif embedded within a cupin fold, in contrast to the 2-His-1-carboxylate facial triad commonly found in many other nonheme iron-dependent oxygenases, including α-ketoglutarate-dependent dioxygenases, pterin-dependent hydroxylases, Rieske dioxygenase, and extradiol dioxygenases.

Members of the TDO family are characterized by their distinct thiol substrates (**Figure 1**), and play important roles in sulfur metabolism and oxygen sensing in both eukaryotes and prokaryotes.^1, 2^ The founding member cysteine dioxygenase (CDO) regulates cellular cysteine levels, and its dysregulation is linked to neurodegenerative and autoimmune disorders as well as certain cancers.^4, 5^ Besides CDO, cysteamine dioxygenase (ADO) is the only other TDO found in mammals. ADO regulates sulfur metabolism by directly contributing to hypotaurine production,^6^ a process associated with glioblastoma development and mitochondrial respiratory capacity.^7, 8^ Additionally, ADO oxidizes *N*-terminal cysteine residues on regulatory proteins, promoting their O_2_-dependent proteolysis via the *N*-degron pathway.^9, 10^ A similar function is also identified in plant cysteine oxidase (PCO), which controls the degradation of transcriptional regulators to play central roles in plant development.^9^ TDOs are also found in bacteria. 3-Mercaptopropionate dioxygenase (MDO) oxidizes 3-mercaptopropionate, one of the most abundant thiols in freshwater ecosystems and a central metabolite in bacterial sulfur metabolism.^11, 12^ Mercaptosuccinate dioxygenase (MSDO) converts mercaptosuccinate (MS) to sulfinosuccinate (SFS), which then decomposes to succinate and sulfite (**Figure 1a**) that are subsequently funneled into central metabolism.^13, 14^ Collectively, TDOs constitute a distinct enzyme family that regulates sulfur metabolism, transcriptional signaling, and environmental adaptation through metal-dependent dioxygenation chemistry. Notably, ergothioneine dioxygenase (ETDO) and certain sulfoxide synthases, such as OvoA, also feature a 3-His-ligated iron center and display sulfur dioxygenase activity.^15-17^ However, ETDO catalyzes thione dioxygenation in a dimeric form, whereas sulfoxide synthases are better known for promoting oxidative Cys−His coupling with a distinct structural fold.^24, 25^ Therefore, these enzymes are not generally classified within the TDO superfamily.

**Figure 1.**
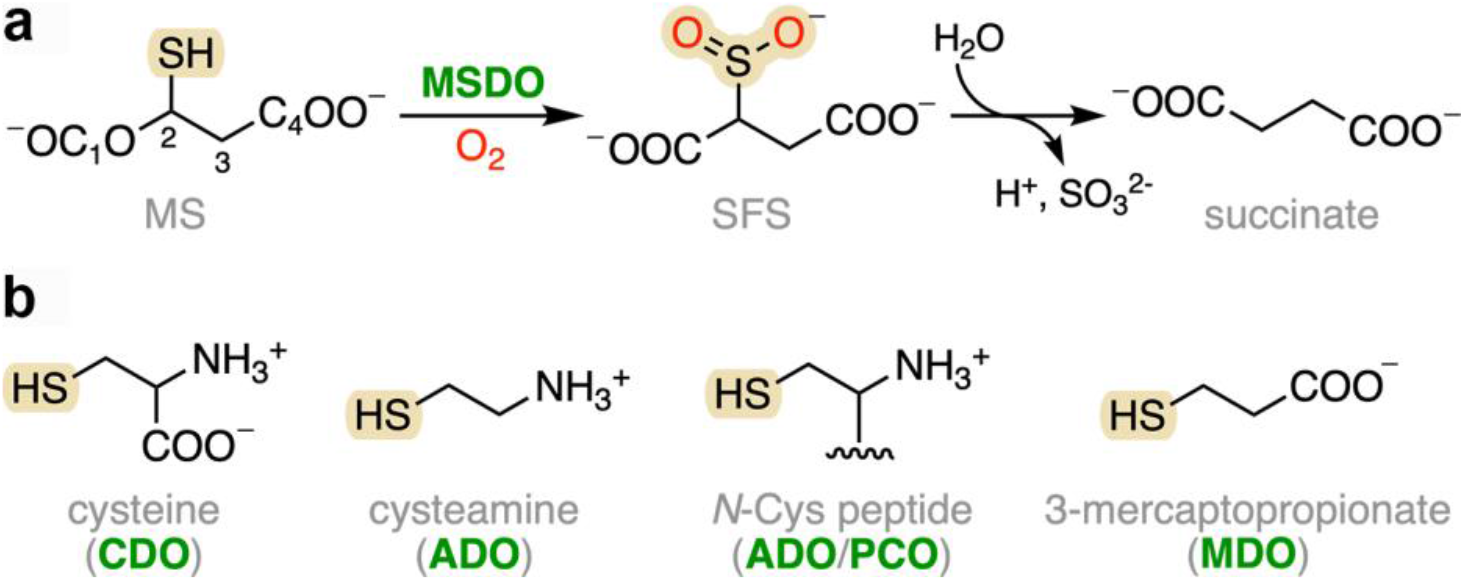
TDOs catalyze dioxygenation reactions on diverse thiol-containing substrates. (**a**) Reaction catalyzed by MSDO and the subsequent non-enzymatic desulfination step. (**b**) Other TDOs (green) and their corresponding thiol substrates.

Nevertheless, mechanistic insights into these closely related systems are expected to advance our overall understanding of their sulfur oxidation chemistry.

MSDO was first identified in the Gram-negative bacterium *Variovorax paradoxus* B4 and found to be conserved across many *Variovorax* strains,^13, 14^ which are species that are known for their metabolic versatility and ability to degrade environmental pollutants,^18, 19^ making them attractive candidates for bioremediation. These bacteria are also increasingly recognized as plant symbionts, capable of promoting plant growth and enhancing tolerance to abiotic stress such as drought and high salinity.^20, 21^ Since MS can be utilized by these bacteria as a sole carbon, sulfur, and energy source via the MSDO-dependent catabolic pathway,^13, 14^ a deeper understanding of this enzymatic process will inform future efforts to employ *Variovorax* as a platform for biotechnology and environmental remediation. Additionally, the substrate MS has broad applications in biomedical research, nanotechnology, cosmetics, and pharmaceuticals.^22^ Its two carboxylate groups make it sterically and electrostatically distinct from other thiol substrates, offering a unique opportunity to probe how TDOs recognize chemically diverse small molecules. However, MSDO has not yet been structurally or spectroscopically characterized. Among TDOs, CDO remains the only member with a well-defined substrate binding mode. Its substrate *L*-Cys coordinates the iron center in a bidentate manner through the thiolate and amino groups, supported by both X-ray crystallographic and spectroscopic evidence.^23-25^ Although extensive efforts have been devoted to studying substrate binding in ADO and MDO,^11, 12, 26-31^ the absence of native substrate-bound structures and inconsistencies arising from different approaches leave the principles of substrate recognition unresolved, highlighting the need to investigate another TDO system.

Unlike many 2-His-1-carboxylate iron-dependent dioxygenases, whose catalytic mechanisms for C-H bond activation are well established,^32-35^ the molecular basis by which the 3-His-ligated Fe center catalyzes dioxygenation on an *S*-atom is less understood. Specifically, whether the two O-atoms of molecular oxygen are incorporated into the Fe-bound thiolate in a concerted or stepwise manner remains a subject of active debate.^36-39^ To date, experimental evidence directly supporting or excluding proposed pathways is limited, and most mechanistic models are based on studies of CDO, in which the amino group of *L*-Cys participates in metal coordination. However, some thiol substrates lack an amino group (**Figure 1**) or do not coordinate via the amino moiety,^27, 28^ which is expected to alter the reactivity of the iron center. Mechanistic investigation of another TDO system, particularly one with a structurally distinct substrate, is therefore critical for a broader understanding of nonheme iron-promoted *S*-dioxygenation.

Here, we selected MSDO as a new platform for a structural and mechanistic investigation due to its distinct substrate, its importance in prokaryotic sulfur metabolism, and the limited understanding of its catalytic process. The enzyme activity was examined using both (*R*)- and (*S*)-enantiomers of the substrate, as some TDOs show enantiomer specificity. By combining X-ray crystallography and electron paramagnetic resonance (EPR) spectroscopy, we resolved the binding conformations of both enantiomers and compared them with other TDOs, filling a critical gap in the understanding of substrate recognition in TDOs. In addition, time-resolved *in crystallo* experiments captured key catalytic intermediates for the first time. Our studies, therefore, provide direct insight into key steps of the *S*-dioxygenation in TDOs. Complementary computational studies validated these intermediates, allowing us to propose a stepwise reaction mechanism and defined the roles of active site residues. Together, the insights obtained in this work not only define the molecular basis of MSDO catalysis but also offer broader implications for understanding structure-function relationships across the TDO family.

## Results and Discussion

### MSDO oxidizes both substrate enantiomers and forms a transient product

MS is a chiral molecule, yet MSDO’s ability to recognize and oxidize its individual enantiomers is unclear. Since commercially available MS is supplied as a racemic mixture (*rac*-MS), the two enantiomers were first resolved by chiral HPLC (**Figure 2a**). The fractions eluting at 5.6 and 9.0 min were assigned as (*R*)-MS or (*S*)-MS, respectively, based on their distinct ellipticity signals centered at 230 nm in circular dichroism spectra (**Figure 2b**).^40^ The corresponding HPLC fractions were pooled, dried under vacuum, and used for all subsequent experiments.

**Figure 2.**
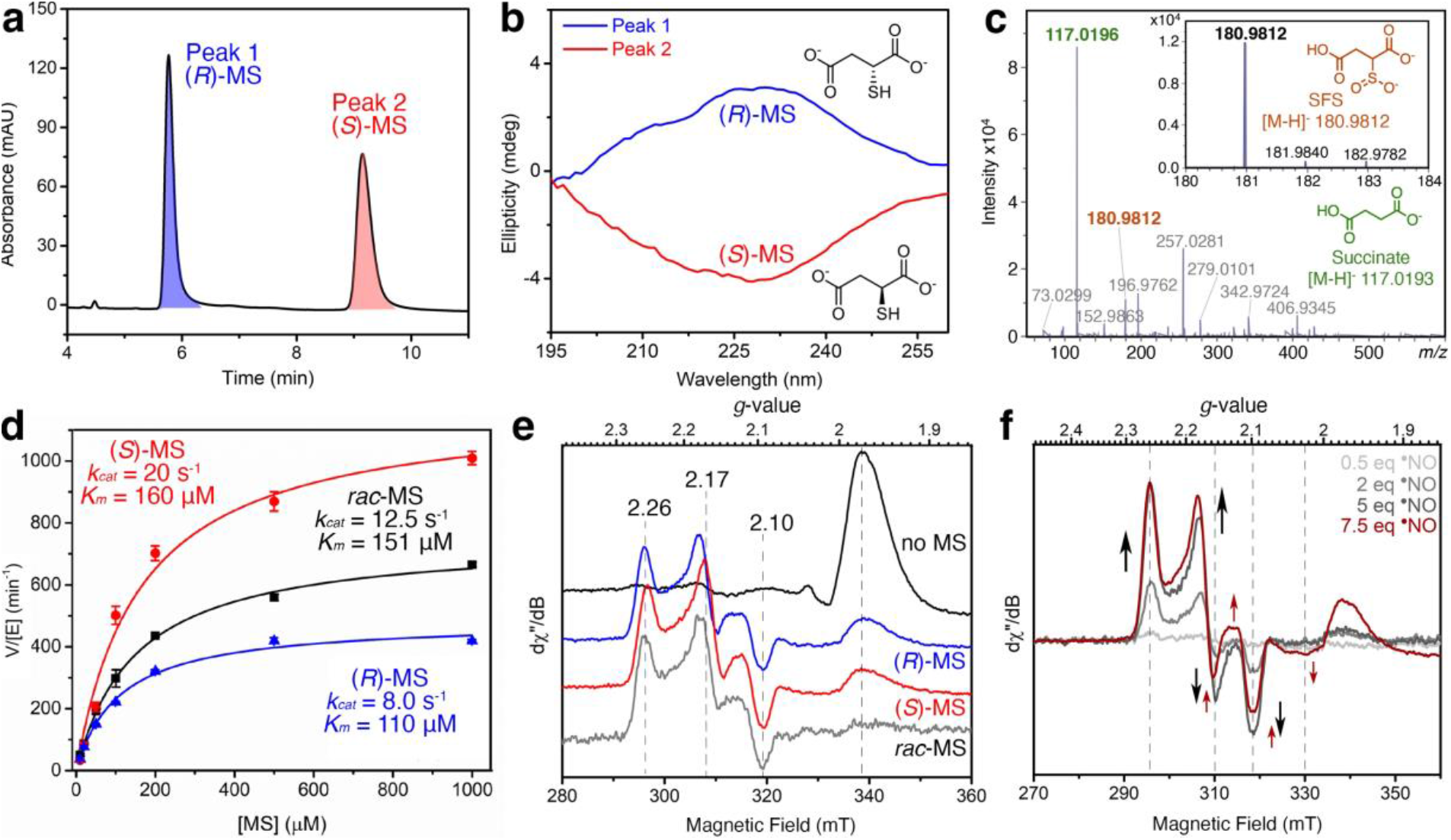
Biochemical and spectroscopic characterization of MSDO activity and substrate binding. (**a**) Chiral HPLC chromatogram showing separation of MS enantiomers. Peak 1 (blue) and Peak 2 correspond to (*R*)-MS and (*S*)-MS, respectively, as assigned by (**b**) circular dichroism spectroscopic analysis. (**c**) High-resolution mass spectrometry of a rapid freeze-quenched MSDO reaction. In addition to the major desulfinated product succinate (green, *m/z* 117.0196), a dioxygenated product SFS was detected at *m/z* 180.9812 (brown). The inset shows the experimental isotope pattern of SFS that matches well with the theoretical distribution. (**d**) Michaelis-Menten plots for MSDO reactions with (*S*)-MS (red), *rac*-MS (black), and (*R*)-MS (blue). Initial rates were determined using an O_2_ electrode with 1 µM freshly Fe-reconstituted MSDO. (**e**) EPR spectra of MSDO incubated with excess °NO. From top to bottom: in the absence of MS (black), in the presence of (*R*)-MS (blue), (*S*)-MS (red), and *rac*-MS (gray). The (*R*)-MS-bound nitrosyl complex exhibits a low-spin rhombic signal with *g* = 2.26, 2.17, and 2.10. (*S*)- and *rac*-MS-bound spectra are highly similar but show slight shifts in *g* values. (**f**) EPR spectra of MSDO pre-complexed with *rac*-MS upon incubation with 0.5, 2, 5, and 7.5 equiv of °NO. The substrate-bound nitrosyl species (g = 2.26, 2.17, and 2.10) formed first. Upon increasing °NO to 7.5 equiv, spectral changes associated with DNIC formation became apparent, as indicated by the red arrows. All EPR spectra were collected at 10 K with 1 mW microwave power.

To determine whether MSDO can oxidize both MS enantiomers, we analyzed enzymatic reactions using liquid chromatography high-resolution mass spectrometry (LC-HRMS). After 5 min of incubation, reactions containing either (*R*)-MS or (*S*)-MS yielded the same product with *m/z* values of 117.0191 and 117.0192, respectively, suggesting the formation of succinate (**Figure S1**). However, the dioxygenated product SFS was not detected under these conditions, consistent with prior reports that SFS could neither be observed in *rac*-MS reactions nor chemically synthesized owing to its instability.^13, 14^ To capture the fleeting product, we rapidly quenched the reaction by flash-freezing immediately after mixing enzyme and substrate. Interestingly, a species with *m/z* 180.9809 was observed in addition to succinate, which corresponds to SFS (**Figure 2c**). This result provides the first direct experimental observation of MSDO’s primary dioxygenation product. The inherent instability of SFS may arise from its sulfinic acid moiety positioned adjacent to a β-carboxylate, which promotes rapid. reductive C−S bond cleavage to yield succinate and sulfite. Spontaneous product desulfination has not been observed in other TDOs, as their products lack a β-carboxylate, although a similar but much slower process has been reported for the sulfinate product formed by ETDO.^17^

Next, the dioxygenase activity of MSDO was evaluated by monitoring O_2_ consumption using an oxygen electrode. Since MSDO is only active in the ferrous form, the enzyme was freshly iron-reconstituted following an established protocol^41^ for kinetic assays using (*R*)-, (*S*)-, and *rac*-MS. The catalysis exhibited a proportional decrease in O_2_ level with increasing substrate concentration, whereby O_2_ was consumed at an approximately 1:1 ratio to MS at lower substrate concentrations (< 100 μM) (**Figure S2**). Reactions of (*S*)-MS showed faster O_2_ consumption than those with (*R*)-MS at the same concentrations. Initial rates were obtained to generate Michaelis-Menton curves for steady-state kinetic analysis (**Figure 2d** and **Table S1**). For the *rac*-MS, a *K*_*m*_ of 0.15 mM and a *k*_*cat*_ of 12.5 s^-1^ were determined. Compared with previously reported values,^13, 42^ the lower *K*_*m*_ and higher *k*_*cat*_ values observed here indicate improved apparent substrate affinity and enhanced catalytic activity after Fe reconstitution. The Michaelis-Menten curve of *rac*-MS lies between those of the individual enantiomers, reflecting an average of their activities. MSDO exhibited similar *K*_*m*_ values for the enantiomers, 0.11 and 0.16 mM for (*R*)- and (*S*)-MS, respectively. However, (*R*)-MS shows catalytic turnover approximately 2.5-fold slower than that of (*S*)-MS. Therefore, the catalytic efficiency of (*R*)-MS (*k*_*cat*_/*K*_*m*_ = 73 mM^-1^ s^-1^) is 1.8-times slower than that of (*S*)-MS (*k*_*cat*_/*K*_*m*_ = 130 mM^-1^ s^-1^). These results indicate that MSDO processes both enantiomers but preferentially oxidizes (*S*)-MS, which is in contrast to CDO and PCO that display strict enantioselectivity for *L*-Cys or *N*-terminal *L*-Cys residues over the *D*-forms.^43-45^

### Substrate enantiomers exhibit similar bidentate iron coordination

The binding of the substrate enantiomers to MSDO was probed by X-band continuous-wave EPR, a technique commonly used in TDO studies to examine how iron centers interact with thiol substrates and small molecules. DEA NONOate was used as a ^•^NO donor to generate a spin probe for the ferrous iron and to act as an oxygen surrogate. When MSDO was introduced to excess ^•^NO, a minor high-spin (*S* = *3/2*) signal at *g* = 3.98 and other marginal low-spin species around *g* = 2 region were observed, along with a prominent positive absorptive feature at *g* = 1.97 characteristic of excess free ^•^NO in solution (**Figure S3a**). The high-spin species is best assigned as {FeNO}^7^ based on Feltham-Enemark formalism and closely resembles the nitrosyl complexes previously characterized for CDO and OvoA.^15, 24^ Upon addition of (*R*)-MS, the resonances corresponding to the high-spin {FeNO}^7^ and excess ^•^NO were greatly diminished, concomitant with the appearance of a new rhombic low-spin species showing *g* values of 2.26, 2.17, and 2.10 (**Figure 2e**). These spectral changes indicate an obligate binding mode in which substrate should first coordinate to the iron center before small molecules such as ^•^NO and O_2_ can effectively bind. Such obligate binding behavior is characteristic of most TDOs,^11, 15, 24^ with ADO being an exception.^27^

The low-spin nitrosyl species was also observed in the complexes of (*S*)-MS and *rac*-MS, with only slight shifts in *g* values, indicating that the enantiomers coordinate the iron center in a similar manner. However, this species differs from previously observed low-spin nitrosyl complexes in TDOs, including the mono-nitrosyl complex in CDO with thiolate and amine bidentate coordination,^24, 46^ the dinitrosyl iron complex in ADO with monodentate thiolate coordination,^27^ and the nitrosyl complex in OvoA with dual substrate coordination by His and Cys,^15^ suggesting a unique iron coordination environment influenced by the carboxylate. In contrast, MDO remains as a high-spin {FeNO}^7^ (*S* = 3/2) species upon substrate binding, suggesting a distinct substrate binding conformation from MSDO, despite the proposed bidentate coordination of substrate carboxylate and thiol groups.^11, 29^ To further investigate this unique low-spin species, power saturation experiments were conducted by monitoring the intensity of the *g* = 2.26 feature at 20 K (**Figure S3b**). The power at half-saturation *P*_*1/2*_ was determined as 0.17 mW, which is much lower than the 1.3 mW determined in CDO at the same temperature.^24^ This is indicative of a slower relaxation of the iron center caused by replacing the amine binding with carboxylate coordination. The difference reflects changes in ligand field strength and metal−ligand covalency, supporting that the electronic structure of the iron center is modulated by distinct substrate coordination modes. The lineshape factor *b* is 1.2, indicating a mostly homogeneous environment with slight inhomogeneous broadening. Taken together, the (*R*) and (*S*)-enantiomers exhibit similar binding modes to the iron center through a bidentate coordination via their thiolate and carboxylate groups.

In addition to the major low-spin species, an unresolved low-spin component was also required to simulate the spectrum (**Figure S4**). This second species is similar to some previously reported dinitrosyl iron complexes (DNICs).^47, 48^ To confirm that the major species with more rhombicity is a mono-nitrosyl species derived from the substrate-bound complex before the formation of the DNIC, varying concentrations of ^•^NO were added to MSDO pre-incubated with excess MS (**Figure 2f**). 0.5 eq. of ^•^NO was insufficient to form the nitrosyl complex. At 2 eq., only the species with *g* = 2.26, 2.17, and 2.10 was observed and the population increased at 5 eq. ^•^NO. Upon increasing ^•^NO to 7.5 eq., the DNIC signal became apparent, as indicated by the emergence of a feature at *g* = 2.03. The apparent decrease in signals at *g* = 2.10 and 2.15 arises from spectral overlap of opposite phased components of the initial low-spin species and DNIC, leading to partial signal cancellation. These two species account for approximately 65% and 35% of the total signal intensity through the spectral fitting (**Figure S4**). These results indicate that DNIC formation occurred only under large excess ^•^NO, likely due to displacement of the substrate by a second ^•^NO molecule.

### X-ray crystal structures provide the first structural views of MSDO

The first MSDO X-ray crystal structures are reported here and provide structural insight into this least characterized member of the TDO family. The substrate-free structure was solved at 1.57 Å resolution (PDB code: 8VNI), with one monomer present in an asymmetric unit. Consistent with other TDOs, MSDO adopts a cupin fold in which its catalytic center is embedded within a β-barrel (**Figure 3a**). The structure is comprised of two α-helices, three β-sheets (β1-β2-β8-β3-β6, β7-β4-β5, and β9-β10), and two 3_10_-helices. Most residues are resolved with clear electron density, except for a short *C*-terminal region (residues 188-192) following β10. Although partially disordered, this *C*-terminal loop lies across one end of the β-barrel and functions as a “lid” that occludes the active-site entrance. This lid is expected to protect the active site from binding of adventitious molecules and to contribute to substrate specificity and product release in MSDO. Similar *C*-terminal loop lids are observed in CDO and MDO,^12, 46^ which process small thiol substrates, whereas in ADO and PCO this region is repositioned towards the β-barrel periphery to accommodate the intrusion of large peptide substrates (**Figure S5**).^49, 50^ Consistent with these functional distinctions, MSDO aligns closely with MDO and CDO, while the extended loops and peripheral features characteristic of ADO and PCO are absent in MSDO.

**Figure 3.**
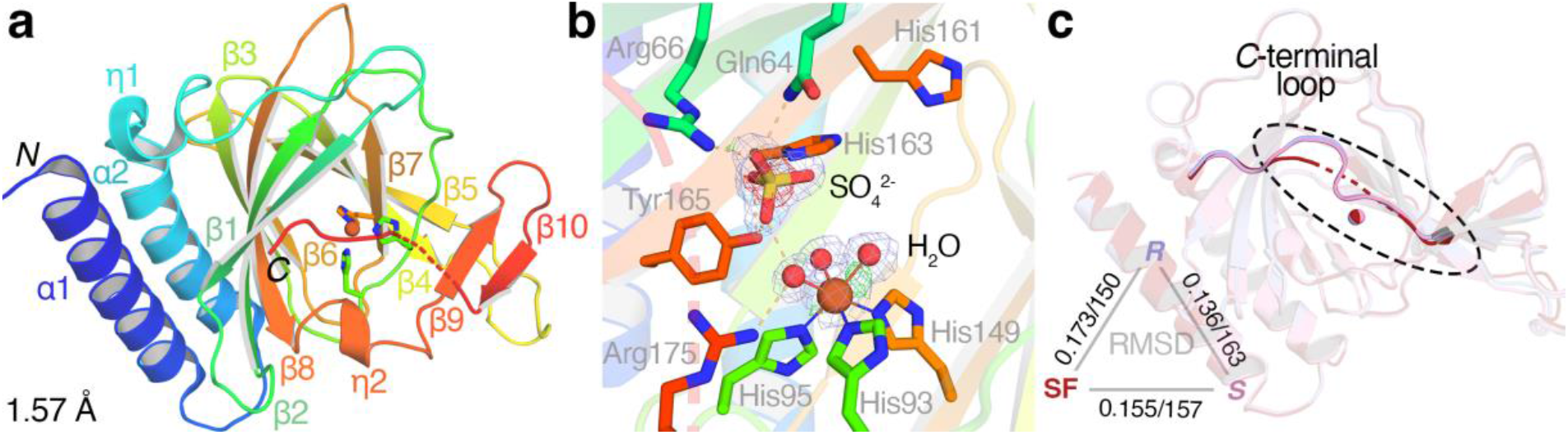
Crystal structures of MSDO. (**a**) Overall structure of substrate-free MSDO shown in a rainbow representation, with the *N*-terminus colored blue and the *C*-terminus colored red. The disordered *C*-terminal loop is indicated by red dashes. The structure was determined to a resolution of 1.57 Å (PDB code: 8VNI). (**b**) Close-up view of the active site in the substrate-free MSDO structure. The iron center adopts a hexacoordinate geometry, ligated by three water molecules and three His residues. A sulfate ion is observed in the active site. The 2*F*_o_-*F*_c_ electron density map is contoured at 1σ (light blue), while the *F*_o_-*F*_c_ difference maps are contoured at 3σ and -3σ (green and red, respectively). Interactions between active site residues and the sulfate are shown by wheat dashed lines. (**c**) Structural superposition of substrate-free MSDO (red) with the (*R*)-MS-bound (purple) and (*S*)-MS-bound (pink) complexes. The *C*-terminal loop, which is disordered in the substrate-free (SF) structure (red dashed line), becomes ordered upon binding of either substrate enantiomer. RMSD values (Å) over the numbers of C_α_ atoms, calculated from structural superposition, are shown in the triangle.

In the active site, MSDO adopts a 3-His coordination, comprising His149 from β7 and His93 and His95 from the loop adjacent to β7 (**Figure 3b**). Three water molecules complete the iron coordination sphere, yielding an octahedral geometry that is characteristic of TDOs in the resting state. Additional electron density is observed in the active site and is best modeled as a sulfate ion (SO_4_^2-^) derived from the crystallization buffer. The sulfate ion is stabilized by interactions with surrounding polar residues, including Gln64, Arg66, His163, and Tyr165. The sulfate-binding pocket is likely capable of accommodating negatively charged functional groups, such as the substrate carboxylate groups. Notably, His163 and Tyr165 are conserved in both MDO and CDO (**Figure S6**), where they participate in an outer-sphere Ser-His-Tyr (SHY) motif that facilitates O_2_ and substrate binding and enhances catalysis.^26, 29, 43^ In CDO, the Tyr residue further forms a catalytically important Tyr-Cys crosslinked cofactor.^23, 51^ In MSDO, however, the Ser of the Ser-His-Tyr motif is replaced by His161, and the crosslinked Cys is substituted by Ala99. The distinct side chain p*K*_a_ of His161, together with its spatial separation from His163 (**Figure 3b**), suggests that although His163 and Tyr165 are conserved, their roles in MSDO catalysis may differ from those established in CDO and MDO.

### Substrate enantiomers adopt different binding conformations

The active-site sulfate from the initial crystallization conditions precludes substrate binding, and both co-crystallization and substrate-soaking attempts failed to yield MS-bound complexes. Consequently, we adopted an alternative crystallization condition free of oxyanions to obtain substrate-bound MSDO structures. Under anaerobic conditions, electron density corresponding to bound MS was observed in the active site when 2 mM of the respective enantiomer was included in the crystallization buffer. The structures of (*R*)- and (*S*)-MS-bound complexes were determined at resolutions of 2.18 and 2.05 Å, respectively (PDB codes: 9NYW and 9NYV). Detailed X-ray crystallography data collection and refinement statistics are shown in **Table S2**. The (*R*)-MS-bound structure exhibits continuous, well-defined electron density, allowing unambiguous modeling of the substrate conformation (**Figure 4a**). The carbon chain of (*S*)-MS is partially disordered; however, the electron-rich thiolate and carboxylate groups are clearly resolved for reliable assignment of its conformation (**Figure 4c**). Attempts to fit the electron density with water molecules or the opposite enantiomer resulted in poor fits (**Figure S7**).

**Figure 4.**
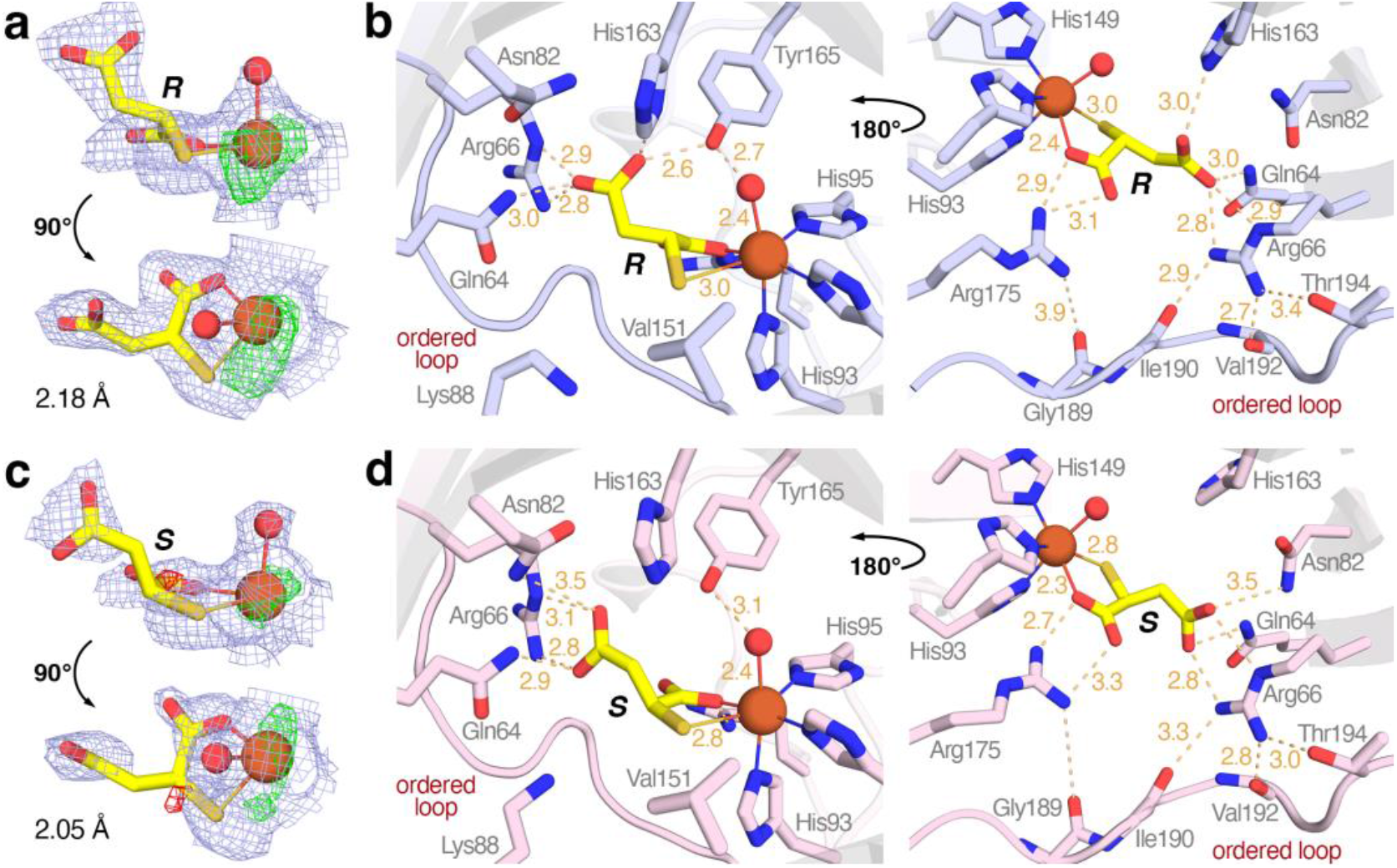
Crystal structures of MSDO in complex with MS enantiomers. (**a, c**) Electron density maps of Fe-bound (*R*)-MS (**a**) and (*S*)-MS (**c**). The 2*F*_o_-*F*_c_ electron density maps are contoured at 1σ (light blue), while the *F*_o_-*F*_c_ difference maps are contoured at 3σ and -3σ (green and red, respectively). (**b, d**) Front (left) and back (right) views of the active site interactions with (*R*)-MS (**b**) and (*S*)-MS (**d**). Interactions between active-site residues and MS are indicated by wheat dashed lines, with distances given in Å. Back views show that Arg66 and Arg175 form interactions with residues on the ordered *C*-terminal loop (shown in opaque). Residue side chains and interactions are selectively hidden in either the front or back views for clarity. The (*R*)-MS- and (*S*)-MS-bound structures were determined at resolutions of 2.18 and 2.05 Å, respectively (PDB codes: 9NYW and 9NYV).

In the finalized structures (**Figure 4b** and **4d**), both (*R*)- and (*S*)-MS bind in a bidentate fashion via their thiolate and the adjacent carboxylate (labeled as -C_1_OO^−^ in **Figure 1**), displacing two water ligands. The thiolate does not engage in direct interactions with active site residues and instead faces toward Val151. A hydrophobic residue is maintained at this position across other TDOs, indicative of a conserved thiolate-binding site and a hydrophobic environment that likely promotes *O*-atom transfer. The C_1_OO^−^ group forms a salt bridge with Arg175, a residue conserved only in MDO, suggesting a similar substrate carboxylate coordination may occur in MDO.^29^ This binding mode is consistent with the substrate-bound low-spin nitrosyl complexes of MSDO probed by EPR spectroscopy. The carboxylate ligates to iron at the same distance (2.4 Å) in both enantiomers, whereas the *S*-atom in (*R*)-MS is further from the iron than in (*S*)-MS (3.0 vs 2.8 Å), which may account for the slight differences in the *g*-values observed in the EPR spectra. The difference in sulfur coordination likely also contributes to the different kinetics between the two enantiomers, with (*S*)-configuration turning over more rapidly. Additionally, a water molecule *trans* to His93 is retained in both enantiomer-bound structures, forming an interaction with Tyr165. A vacant sixth coordination site for O_2_ binding is typical in nonheme iron oxygenases upon substrate binding,^32, 33^ however, retention of this water ligand in the MSDO active site is not entirely unexpected, as Fe-coordinated solvent ligands have also been observed in some TDO structures complexed with substrates or substrate analogs.^23, 29, 52^

The distal carboxylate group of MS (labeled as -C_4_OO^−^ in **Figure 1**) adopts distinct orientations in the (*R*)- and (*S*)-MS-bound structures, resulting in different interaction networks within the active site. In the (*R*)-MS complex, the carboxylate is stabilized by Gln64, Arg66, His163, and Tyr165, whereas in the (*S*)-MS complex it rotates away from His163 and Tyr165 and only interacts with Gln64, Arg66, and Asn82. This conformational difference is accompanied by rotation of Asn82 and His163 during structural refinement, with their side chains adopting alternative conformations to form favorable van der Waals contacts and hydrogen-bonding interactions. Arg66 is essential for binding of both enantiomers and is conserved only in CDO, where it also forms a salt bridge with the carboxylate group of the substrate (**Figure S6**). On the other hand, no such equivalent Arg is found in MDO, as Arg175 stabilizes the substrate carboxylate that coordinates to the Fe center.^29^ Given that MS enantiomers exhibit comparable *K*_m_ values, the different binding conformations observed between (*R*)- and (*S*)-MS are not expected to significantly affect substrate binding affinity but rather influence catalytic turnover.

Arg66 and Arg175 are not only critical for MS recognition but are postulated to also play important roles in shielding the active site. This is consistent with the previous functional analysis of MSDO that shows the enzyme activity was greatly diminished upon deleting Arg66.^42^ The overall MSDO structure remains largely unchanged when MS binds, with root-mean-square deviations (RMSDs) of 0.173 Å over 150 C_α_ and 0.155 Å over 157 C_α_ relative to the substrate-free structure for (*R*)- and (*S*)-MS-bound complexes, respectively. However, the *C*-terminal loop, which is flexible in the substrate-free form, becomes ordered in the MS-bound structures (**Figure 3c**), and the resolved loop residues, Gly189, Ile190, Val192, and Thr194, form interactions with Arg66 and Arg175 (**Figure 4b** and **4d**, right panels). In the absence of MS, these Arg residues are likely more flexible, allowing the *C*-terminal loop to adopt an open conformation that permits substrate entry. Substrate interactions constrain the Arg residues, thereby stabilizing the loop in a closed conformation that facilitates catalysis. Notably, the conserved Arg60 in CDO and Arg168 in MDO similarly interact with their *C*-terminal loop lids upon substrate/analog binding (**Figure S6**). Therefore, the Arg residues mediate substrate recognition while regulating active-site accessibility, constituting a conserved active-site gating mechanism among small thiol-oxidizing TDOs.

### Time-resolved in crystallo reactions capture monooxygenated and dioxygenated intermediates

Spectroscopic studies suggest that O_2_ binding in TDOs generates an Fe(III)-superoxo species,^37, 53,54^ a commonly invoked intermediate in nonheme iron enzymes. However, how this superoxo species subsequently incorporates two O-atoms into a thiol substrate remains an open mechanistic question due to the lack of characterized intermediates. Time-resolved *in crystallo* reactions have emerged as a powerful approach for trapping catalytic intermediates during enzymatic turnover, particularly in iron-dependent oxygenases.^55-57^ To gain molecular insights into MSDO-catalyzed thiol dioxygenation, substrate-bound crystals prepared under anaerobic conditions were transferred into aerobic solutions for defined incubation times prior to flash-freezing in liquid nitrogen. Crystals of the (*S*)-MS-bound complex did not yield interpretable results, resulting in heterogeneous and incomplete electron density likely due to the faster catalytic turnover. In contrast, the (*R*)-MS-bound crystals produced compelling results.

Following a 20 s oxygen exposure of the (*R*)-MS-bound crystals, an intermediate structure was determined at 1.98 Å resolution (PDB code: 9PYP). Modeling the active site electron density with (*R*)-MS alone left unresolved electron density adjacent to the *S*-atom (**Figure S8a** and **S8b**), indicating a ligand identity that differs from the starting model. Modeling a monooxygenated intermediate in the (*R*)-configuration, in which a single *O*-atom is inserted between the Fe−S bond to form an Fe-bound sulfenate species, resulted in an excellent fit (**Figure 5a**). Attempts to model a further dioxygenated intermediate introduced negative difference density, suggesting that the electron density is insufficient to support incorporation of a second O-atom at this stage of the reaction (**Figure S8c**). Following O−O bond cleavage of the superoxo intermediate, formation of a ferryl species, i.e. Fe(IV)=O, is predicted by computational analysis.^39^ However, X-ray-induced photoreduction during data collection resulted in an iron center that was best assigned as a water-bound Fe(II) species, as supported by the observed Fe−O distance of 2.4 Å in the crystal structure. Tyr165 and nearby water molecules may serve as proton donors during reduction of the Fe(IV)=O. Future spectroscopic assessment and reduction-free techniques will be required to better understand the iron oxidation state of this intermediate.

**Figure 5.**
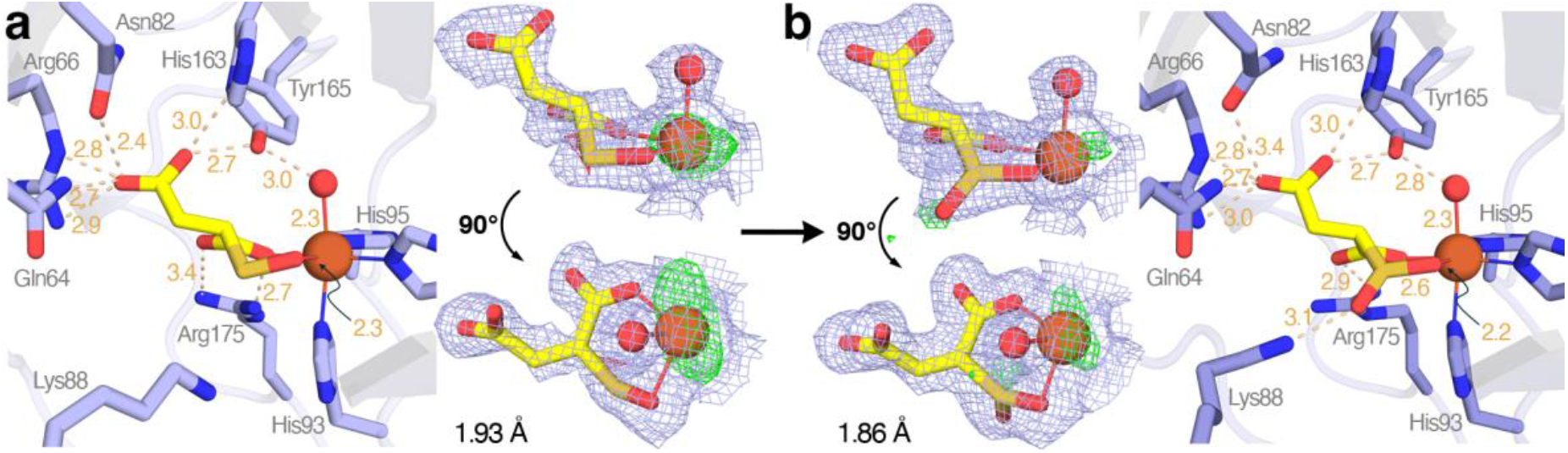
Crystal structures of catalytic intermediates derived from (*R*)-MS captured by *in crystallo* reaction. Active-site views and corresponding electron density maps of (**a**) an Fe-bound monooxygenated, sulfenate intermediate and (**b**) an Fe-bound dioxygenated, sulfinate intermediate. The 2*F*_o_-*F*_c_ electron density maps are contoured at 1σ (light blue), while the *F*_o_-*F*_c_ difference maps are contoured at 3σ and -3σ (green and red, respectively). Interactions between active-site residues and the bound intermediates are indicated by wheat dashed lines, with distances given in Å. The sulfenate- and sulfinate-bound structures were determined at resolutions of 1.93 and 1.86 Å, respectively (PDB codes: 9PYP and 10AB).

Upon extending the oxygen exposure time to 30 s, the electron density surrounding the *S*-atom became more pronounced and modeling with the sulfenate intermediate could not account for the additional electron density (**Figure S8d-f**). Instead, modeling the active site with the fully dioxygenated sulfinate product, SFS, in the (*R*)-configuration resulted in a satisfactory fit to the electron density (**Figure 5b**, PDB code: 10AB). The dioxygenated product adopts a bidentate binding conformation that closely resembles that of the monooxygenated intermediate, differing only by one additional O-atom. Structural superposition of the two states shows that the overall structure and active-site architecture are largely preserved (RMSD = 0.096 Å over 163 C_α_). However, notable structural rearrangements in the active site are observed for Lys88 and Arg175, whose side chains reposition to interact with the newly formed sulfinate moiety and the proximal carboxylate group, respectively (**Figure S9**). The active site dynamics likely accommodate the increased polarity that arises upon product formation. A water molecule *trans* to His93 is coordinated to the iron center and may be supplied through a continuous hydrogen-bonded water network that connects the iron coordination sphere to the bulk solvent (**Figure S10**). This water chain provides a pathway for rapid proton or water transfer, thereby allowing the coordination site to be readily reoccupied following catalytic turnover and potentially restoring the 3H_2_O-ligated iron center upon product release.

Notably, although the sulfinate species is highly labile in solution and readily undergoes desulfination, the SFS product remains unexpectedly stable within the MSDO active site, persisting on the hour timescale before significant degradation of electron density is observed. In the crystal structure, SFS coordinates to the iron center through both the sulfinate moiety and the adjacent carboxylate, a binding mode that likely constrains the product geometry and provides electrostatic stabilization. In contrast, the sulfinate product in solution lacks such coordination and is therefore more susceptible to spontaneous C−S bond cleavage. This observation validates that desulfination is a non-enzymatic stepthat occurs after product is released from the active site. The structure also reveals a closed conformation of the *C*-terminal loop, which may be constrained by crystal lattice, potentially accounting for the retention of the dioxygenated product. Together, these findings suggest a stepwise mechanism of oxygen addition mediated by the iron center and highlight its critical role in stabilizing an otherwise transient product.

Although the stepwise thiol dioxygenation mechanism has long been proposed within the TDO family, the structural snapshots of catalytic intermediates in MSDO provide the first direct experimental evidence supporting this pathway. Previous crystallographic studies of CDO identified an Fe(II)-bound persulfenate intermediate consistent with a concerted oxygen addition pathway,^38, 58^ but subsequent experimental and computational investigations questioned whether this species represent an on-pathway intermediate.^39, 59^ More recently, a mechanistic study showed that cobalt-substituted ADO is catalytically competent, supporting the concerted mechanism that does not invoke a high-valent metal-oxo intermediate.^36^ However, neither CDO nor ADO features substrate carboxylate coordination to the iron center, as observed in MSDO. This distinction is expected to modulate the electronic structure and reactivity of the metal center, potentially favoring different oxygen activation pathways based on the charge maintenance concept.^60, 61^ Further differences are evident in product-bound complexes. A biomimetic CDO complex shows that cysteine sulfinic acid coordinates to Fe(II) in a tridentate fashion through both sulfinate O-atoms and the amine nitrogen,^62^ in contrast to the bidentate coordination observed in MSDO. This difference likely arises from metal metal center charges and geometric constraints imposed by the bound products. In CDO, the product contains a β-amino relative to the sulfinate, whereas in MSDO the substrate features an α-carboxylate adjacent to the sulfinate. Collectively, mechanistic investigations across TDOs with diverse substrate binding modes are essential to define how substrate coordination governs oxygen addition pathways. In this context, our study fills a critical gap by providing a carboxylate-bound example that supports a stepwise dioxygenation mechanism.

### Computational studies on the reaction mechanism and selectivity patterns

Based on the above reported crystal structure coordinates of the (*R*)- and (*S*)-MS-bound complexes (9NYW and 9NYV, respectively) we created enzymatic models and ran a 200 ns molecular dynamics (MD) simulation (see Supporting Information for system preparation and setup). Both simulations stabilized very quickly and showed a rigid substrate binding structure and stable first- and second-coordination sphere around the metal and substrate. In agreement with the crystal structure coordinates, we identify dominant hydrogen bonding interactions to (*S*)-MS from Gln64, Arg66, Asn82, and Arg175 that provide stability to the complex and force the substrate into a specific orientation. Additional interactions involving His163 and Tyr165 are observed in the (*R*)-MS-bound structure.

Using the average structure of the two MD simulations, large QM cluster models were created that include the substrate, Fe(III)-superoxo and its first- and second-coordination sphere. These QM cluster models should capture the intra- and intermolecular interactions of the protein, substrate, and cofactor well and describe the features and properties of the actual structure and its reactivity (**Figure S11**).^63, 64^ Thus, the QM cluster model of the Fe(III)-superoxo reactant with (*S*)-MS bound is designated **Re**_S_ and has 287 atoms, while the one with (*R*)-MS bound is **Re**_R_ and contains 298 atoms. Both structures were calculated in the lowest energy singlet, triplet, quintet and septet spin states, whereby we show the ^5,7^**Re**_S_ and ^5,7^**Re**_R_ optimized geometries (**Figure 6a**). The quintet and septet spin states are close in free energy with a small preference for the septet, while the other two spin states are much higher in free energy (**Tables S3** and **S4**). These energy differences imply that the quintet and septet spin reactant states will likely be stable at room temperature and have a finite probability. The optimized geometries of both enantiomers have the substrate bound through the thiolate and C_1_OO^−^ groups and the Fe(III)-superoxo as end-on. The (*S*)-MS-bound structure has the superoxo group aligned with the Fe−S bond (dihedral angle S−Fe−O−O of 37°), whereby the proximal oxygen atom of the superoxo accepts a hydrogen bond from Tyr165. By contrast, in the (*R*)-MS-bound structures the superoxo group points in the other direction towards Ile101 (dihedral angle S−Fe−O−O of 170°). In addition, in ^5,7^**Re**_R_ the Tyr165 phenol group forms a hydrogen bond with the distal C_4_OO^−^ group of the substrate. These substrate and protein binding orientations match those reported in **Figure 4** that showed a flip of the His163 residue upon binding of the other enantiomer. The C_4_OO^−^ group in ^5,7^**Re**_S_ forms a double salt bridge with Arg66, while several bridging water molecules also connect the group to the Gln64 and Asn82 side chains that provide a tight binding orientation. In ^5^**Re**_R_ the interaction with Arg66 is still in place together with the phenol group of Tyr165 to the C_4_OO^−^ group. To validate our optimized geometries, we created overlays of ^5^**Re**_S_ and ^5^**Re**_R_ with the PDB files of 9NYV and 9NYW, respectively (**Figure S12**). The overlay gives a good match between the two structures and shows a very tight substrate and oxidant binding pocket that is held together through intramolecular and intermolecular interactions. Although in ^5,7^**Re**_S_ the superoxo group forms an interaction with iron of 2.077 (quintet) and 2.185 (septet) Å, this bond is much longer and weaker in the corresponding (*R*)-MS structure, where the distance reaches well over 2.7 Å. Attempts to reoptimize with shorter Fe−O distances for the (*R*)-MS structure failed, consequently, the interactions in the protein bound with (*R*)-MS disfavor dioxygen binding and as a result may slow down catalysis as indeed observed experimentally.

**Figure 6.**
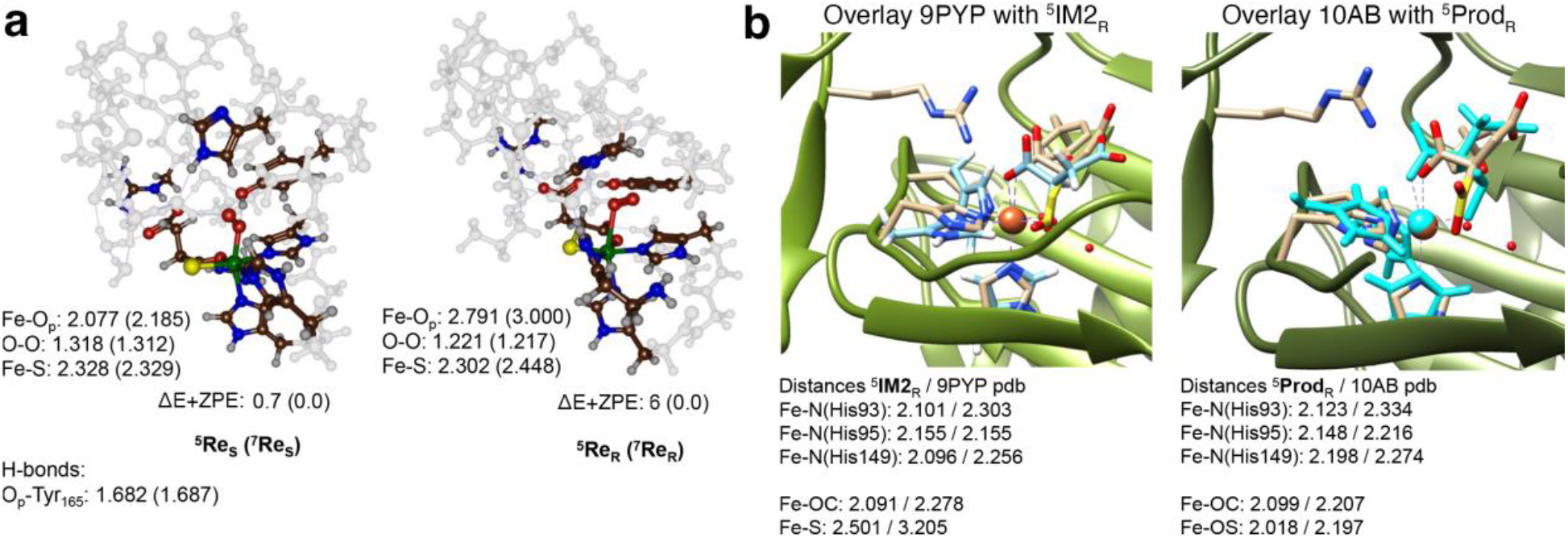
(**a**) DFT optimized geometries of Fe(III)-superoxo complexes with (*S*)- and (*R*)-MS bound. **(A)** Superoxo complexes. Bond lengths are in Å and the relative energy (ΔE+ZPE) in kcal mol^−1^. O_p_ represents the proximal oxygen atom of the superoxo group. (**b**) Comparison of the crystal structure coordinates of 9PYP and 10AB (wheat sticks) with the DFT optimized geometries (blue and cyan sticks). Specified bond lengths are given in Å.

Subsequently, we studied the dioxygenation of (*S*)-MS and (*R*)-MS in MSDO using DFT cluster models. On the septet spin state, the reactant structure is low in energy and in equilibrium with the quintet spin state structure, however, no viable oxygen atom transfer pathway was found. Geometry scans starting from ^7^**Re**_S_ and ^7^**Re**_R_ for S−O bond shortening were performed but did not lead to a transition state structure and a local minimum but instead showed a continuous rise in energy (**Figure S13**). Consequently, the electronic configuration of the septet spin structures prevents dioxygenation of the substrate. In the other spin states the full mechanism was explored and for the full reaction profile starting from either ^5^**Re**_S_ or ^5^**Re**_R_ the quintet spin state was lowest in free energy. **Figure 7** gives the quintet spin free energy profiles of dioxygenation of (*S*)-MS and (*R*)-MS as calculated with DFT. Similarly to previous calculations on CDO and MDO,^65-71^ the mechanism proceeds through a stepwise manner with consecutive oxygen atom transfer, in which attack by the proximal oxygen atom (O_p_) on sulfur is significantly higher in energy than attack by the distal oxygen atom (O_d_) (**Figure S14**). Thus, from the reactant complexes the reaction starts with attack of the terminal oxygen atom of the Fe(III)-superoxo on the sulfur atom of the substrate to form a ring structure with four-membered Fe−O−O−S ring (intermediate **IM1**) via a transition state **TS1**. Thereafter, the peroxo bond cleaves via transition state **TS2** to form an Fe(IV)=O sulfenate intermediate **IM2**. Next, the second oxygen atom transfer takes place via transition state **TS3** to form the sulfinate product complex **Prod**.

**Figure 7.**
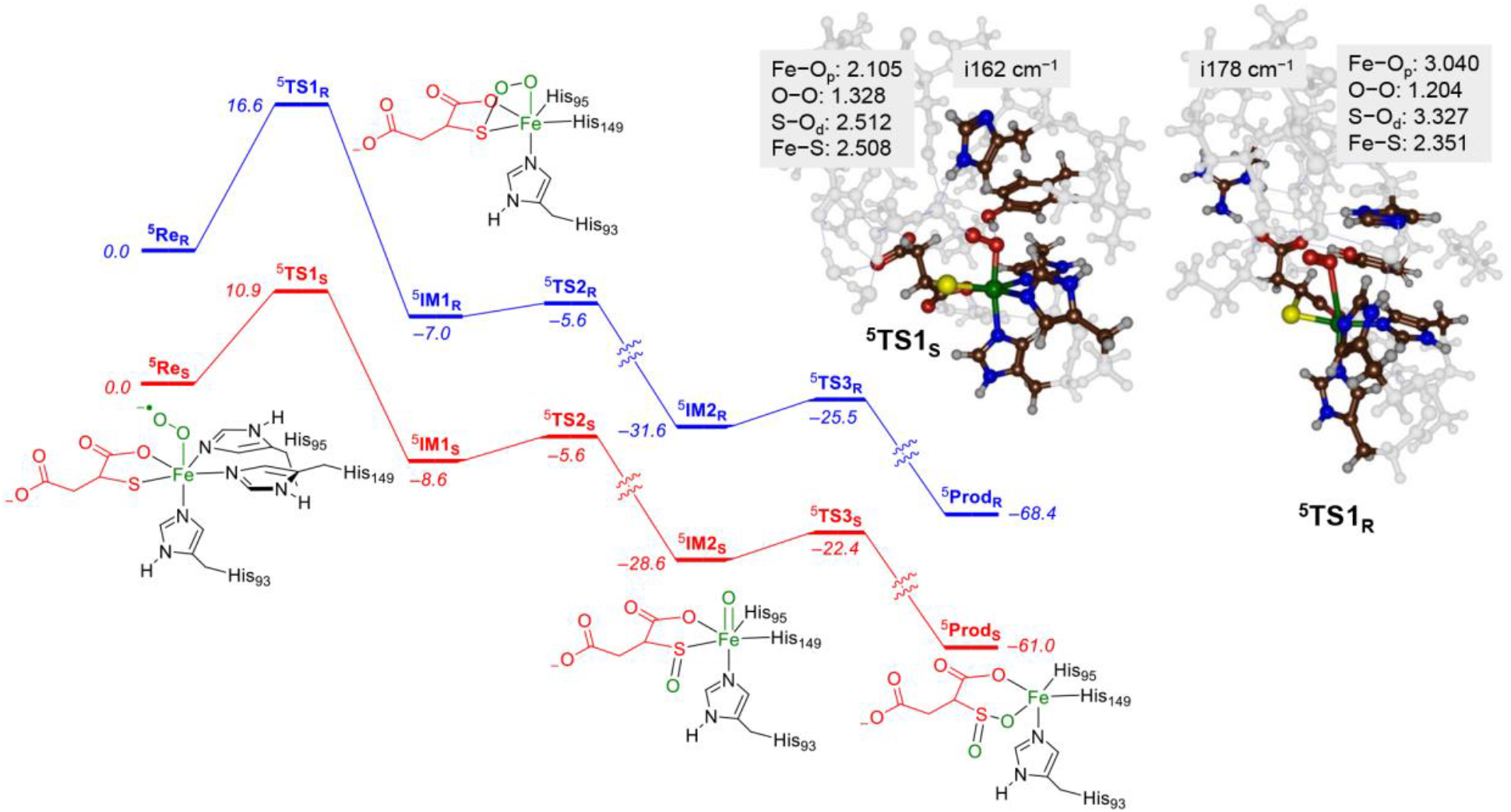
DFT calculated free energy profile at 298 K with values in kcal mol^−1^ for the dioxygenation of (*S*)-MS and (*R*)-MS by MSDO model complexes. Free energies contain zero-point, thermal and entropic corrections to the energy. Optimized transition state structures give bond lengths in Å and the imaginary frequency in cm^−1^.

Both enantiomers react in the active site with a rate-determining S−O bond formation via transition state **TS1** (**Figure 7**), however, ^5^**TS1**_S_ has a free energy of ΔG = 10.9 kcal mol^−1^, while for the (*R*)-MS an activation free energy of 16.6 kcal mol^−1^ is obtained. Despite the large difference in barrier height for the (*S*)-MS and (*R*)-MS substrates, both structures relax to an **IM1** intermediate with similar exergonicity (ΔG = −8.6 kcal mol^−1^ for ^5^**IM1**_S_ and ΔG = −7.0 kcal mol^−1^ for ^5^**IM1**_R_). The same is true for the O−O cleavage barrier and the formation of the Fe(IV)=O intermediates that have similar energetics for the two enantiomers. The only other structure, where the two enantiomers differ by more than 5 kcal mol^−1^ is the product complex that is more stable for the (*R*)-enantiomer than for the (*S*)-enantiomer. The calculated ^5^**Prod**_R_ structure has the product bound as bidentate ligand to Fe(II) through one of the oxygen atoms of the sulfinate group and one of the oxygen atoms of the C_1_OO^−^ group with distances of 2.018 and 2.099 Å, respectively. The overall structure matches the orientation of the bound product in the crystal structure reported above very well (**Figure 6B)**.

To understand the differences in catalysis between (*R*)- and (*S*)-MS activation by MSDO, we compared the ^5^**TS1** transition state structures (**Figure 7**). Both structures have a small imaginary frequency (i162 cm^−1^ for ^5^**TS1**_S_ and i178 cm^−1^ for ^5^**TS1**_R_) for the S−O stretch vibration. Previous studies located transition states with somewhat larger imaginary frequencies of well over i250 cm^−1^,^69, 72^ which means that the potential energy curve around the transition states for MSDO are broader than in CDO due to differences in active site structure, polarity, and hydrogen bonding interactions. The ^5^**TS1**_S_ structure is very similar to the ones reported on CDO with an elongated O−O bond to 1.328 Å and shortened S−O bond to 2.512 Å. The bridging peroxo group in ^5^**TS1**_S_ accepts a hydrogen bond from Tyr165 (to O_p_) and a water molecule (to O_d_) that connects it to His163. By contrast, ^5^**TS1**_R_ has the peroxo group only weakly bound with a long distance to iron (Fe−O_p_ is 3.040 Å) and to sulfur (S−O_d_ is 3.327 Å). At the same time the dioxygen bond is still short (O−O is 1.204 Å). Indeed, an analysis of the group spin densities in ^5^**TS1**_S_ and ^5^**TS1**_R_ shows major differences (**Tables S5** and **S6**), whereby in ^5^**TS1**_S_ the superoxo group has a spin of −0.7, while the sulfur bears 0.8 unpaired electron and the iron 3.7 unpaired electrons. Therefore, the unpaired electrons on sulfur and dioxygen are with opposite spin and will pair up to form the S−O orbital in the sulfoxide complex. By contrast, in ^5^**TS1**_R_ the peroxo group has virtually no unpaired spin, hence, it is in its O_2_^2−^ form, while iron has 3.6 unpaired electrons and sulfur only 0.2. Consequently, from the orbital occupation in ^5^**TS1**_R_, several electrons need to be transferred to obtain the Fe(IV)=O with sulfenate configuration, which is costly and requires a substantially higher barrier as seen for ^5^**TS1**_S_. Together, these computational results delineate distinct oxygen activation modes in different enantiomer complexes and reveal how the active site environment modulates oxygen transfer. The calculations further support a distal oxygen-initiated attack on the thiolate and the formation of an Fe(IV)=O intermediate, providing additional evidence for a stepwise thiol dioxygenation mechanism.

## Conclusion

MSDO occupies a unique position within the biologically important TDO superfamily due to its electrostatically distinct substrate bearing two carboxylate groups. Here, we define the molecular basis of MSDO catalysis through an integrated spectroscopic, kinetic, structural, and computational analysis. MSDO reacts with both (*R*)- and (*S*)-MS enantiomers, displaying comparable binding affinities but increased turnover for (*S*)-MS. The direct oxygenated product, SFS, underwent rapid decay in solution but was detected for the first time by rapid freeze-quench experiments. EPR and crystallographic data establish a bidentate substrate binding mode through the thiolate and the proximal carboxylate, representing a distinct substrate coordination mode within TDOs. While the overall active site geometry is largely conserved for both enantiomers, alterations in distal carboxylate interactions likely underlie the observed differences in reactivity. Comparison of substrate-free and substrate-bound structures reveals a *C*-terminal loop that functions as an active-site gate, regulated by two substrate-stabilizing arginine residues. Most importantly, time-resolved *in crystallo* reactions capture an unprecedented monooxygenated sulfenate intermediate and the unstable dioxygenated product SFS, suggesting a stepwise dioxygenation mechanism. This assignment is further corroborated by DFT calculations that agree with a consecutive oxygen atom transfer mechanism with a rate-determining initial S−O bond formation transition state. Subsequently, exergonic O-O bond cleavage leads to the formation of an Fe(IV)=O species followed by a second oxygen atom transfer to form SFS products. The latter is a stable local minimum with a calculated structure that matches the experimental findings well. Collectively, these findings establish a coherent framework for MSDO catalysis, expand the mechanistic landscape of thiol dioxygenases, and provide insights into substrate recognition and oxygen activation in nonheme iron enzymes.

## Supporting Information

Experimental details, X-ray crystallography data statistics, supplemental tables and figures, and Cartesian coordinates of all DFT-optimized structures.

## Acknowledgements

This work was supported by a National Institute of General Medical Sciences grant R35GM147510 (to YW) from the National Institutes of Health. We thank Dr. Jeffery Urbauer for assistance with circular dichroism measurements and the PAMS Facility for mass spectrometry analyses at the University of Georgia. We also thank the National Science Foundation (MRI grant 1827968 to TH) for support of EPR spectroscopic experiments. We acknowledge the staff and X-ray synchrotron resources at several beamlines, including 22-ID at the Advanced Photon Source, Argonne National Laboratory; 5.0.1 and 8.2.2 at the Advanced Light Source, Lawrence Berkeley National Laboratory; and 19-ID at the National Synchrotron Light Source II, Brookhaven National Laboratory. These facilities are supported by the U.S. Department of Energy (DOE) Office of Science User Facilities operated under Contract Nos. DE-AC02-06CH11357, DE-AC02-05CH11231, and DE-AC02-98CH10886, respectively. HPHW thanks the University of Manchester for a Postgraduate Research and Teaching Award. SdV thanks the Computational Shared Facilities for computer time and access.

## Notes

### Competing Interest Statement

The authors have declared no competing interest.

