## supplemental tables and figures for "A Stepwise Thiol Dioxygenation Mechanism in Mercaptosuccinate Dioxygenase Revealed by A Combined Experimental and Computational Study"

### Experimental methods

*Protein expression and purification*

The codon-optimized gene encoding MSDO from *Variovorax paradoxus* B4 was synthesized and cloned into a pET-28a(+) expression vector (Twist Bioscience) with an *N*-terminal $His_6$-tag cleavable by Tobacco Etch Virus (TEV) protease. The plasmid was transformed into *E. coli* BL21 (DE3) competent cells, which were grown in Luria-Bertani growth medium containing kanamycin (50 µg/mL) at 200 rpm, 30 °C until the culture reached an $OD_{600}$ of 0.6. The temperature was then reduced to 25 °C, and protein expression was induced with 0.5 mM isopropyl β-thiogalactopyranoside (IPTG) and supplemented with 125 µM ferrous ammonium sulfate. Cultures were grown for 24 hours before harvesting by centrifugation, and the resulting cell pellets were stored at -80°C.

Frozen cell pellets were resuspended in Buffer A (50 mM Tris-HCl, 200 mM NaCl, pH 7.6) supplemented with 0.1 mM phenylmethylsulphonyl fluoride and 0.05% (v/v) β-mercaptoethanol. Resuspended cells were lysed via sonication, and the lysate was clarified by centrifugation. The supernatant was loaded onto a HisTrapFF column (Cytiva) equilibrated with Buffer A, using an ÄKTA primer plus FPLC system (GE Healthcare). Protein elution was done with a gradient of Buffer B (50 mM Tris-HCl, 200 mM NaCl, and 350 mM imidazole at pH 7.6), with MSDO eluting at ~15-25% Buffer B. Fractions containing the target protein were pooled based on SDS-PAGE results (MW = 23.9 kDa), concentrated using centrifugal filters (Amicon, 10 kDa cutoff), and buffer exchanged on a HiTrap desalting column (Cytiva) into 50 mM Tris-HCl (pH 7.6), 100 mM NaCl, and 5% glycerol. The purified protein was then concentrated, aliquoted, flash frozen with liquid nitrogen, and stored at -80 °C for further use. Enzyme concentration was determined by UV-visible absorbance at 280 nm using the predicted extinction coefficient ($\varepsilon_{280}$ = 24,210 $cm^{-1}$ $M^{-1}$).[1]

*Purification and determination of* (R)- *and* (S)-*MS*

Commercially obtained MS (>98%, TCI) was supplied as a racemic mixture. The (*R*)- and (*S*)-enantiomers were separated using a Chiralcel OD-H normal phase column (4.6 × 250 mm, 5 µm, Daicel) with an isocratic mobile phase of 90:10 hexane:ethanol containing 0.1% TFA with a flow rate of 1 mL/min on an Ultimate-3000SD HPLC system (Thermo Fisher Scientific). Fractions were collected and analyzed by circular dichroism (CD) spectroscopy (Jasco J-710), and the absolute configurations were assigned based on reported spectra.[2] CD spectra were recorded from 260 to 190 nm at 20 °C using 1 mm-path length cell, with continuous scanning at 100 nm/min, 1-nm bandwidth, 4 s response time, and 1-nm data pitch. Final spectra were represented by the average of four scans. Purified (*R*)- and (*S*)-MS were dried *in vacuo* and re-dissolved in the appropriate solvents for subsequent experiments.

*Product determination by LC-MS*

MSDO was freshly desalted into 50 mM ammonium acetate buffer (pH 7.6). Separate reaction solutions contained either (*R*)- or (*S*)-MS at a final concentration of 2 mM in ammonium acetate buffer. MSDO at a final concentration of 10 µM was added to initiate the reaction. Reaction solutions were incubated at 25 °C for 30 min, then filtered using centrifugal filters (Millipore) with a 3 kDa cutoff to remove enzyme. Filtrate was collected, then stored at -20 °C until analysis.

A Bruker electrospray ionization quadrupole time-of-flight mass spectrometer, Impact II coupled with an Elute UHPLC (Billerica, MA) was used for sample analysis. 4 μL of a solution containing 0.1 μL of sample diluted in 100 μL of water and 0.1% formic acid was injected onto a Restek Ultra II Aqueous C18 column (2.2 um, 50 × 2.1mm) (Bellefonte, PA) with an isocratic solvent system of 95% water and 5% acetonitrile and 0.1% formic acid. Raw data were acquired using Bruker Compass HyStar 5.0 software at the resolving power of 20,000 at 400 Da, scanning m/z 45-650 in negative ion mode. MS2 (CID) was performed on compounds of interest. Bruker Compass DataAnalysis 5.1 was used for data analyses.

*SFS production from Rapid freeze-quenched MSDO reactions*

Desalted MSDO was buffer exchanged into anaerobic 50 mM ammonium acetate buffer (pH 7.6) in a glovebox. *rac*-MS was prepared with aerobic ammonium acetate buffer to a final concentration of 1 mM and 280 μL of the MS solution was transferred to centrifugal filters with a 3 kDa MW cutoff. 20 μL of anaerobic MSDO (final concentration 10 μM) was added to initiate the reaction and the solution was immediately centrifuged at 15,000 rpm, 4 °C for 1 min after addition of enzyme. Filtrate was rescued and flash frozen in liquid nitrogen and stored at -20 °C until analysis. Mass spectrometry analysis was performed as described above with a scanning range of 50-1000 *m/z*.

*Preparation and EPR analysis of MSDO-nitrosyl complexes*

Desalted MSDO in 50 mM Tris-HCl (pH 7.6), 100 mM NaCl, and 5% glycerol was used for EPR sample preparation. Anaerobic enzyme was generated using an argon/vacuum gas manifold and transferred to a glovebox, where all subsequent steps were performed to prevent oxidation and denaturation. EPR samples contained 100 μM enzyme, 0.5 mM ascorbic acid, 10 mM racemic, (*R*)-, or (*S*)-MS, and slight excess •NO released from DEA NONOate (≥98%, Cayman). Following •NO addition, samples were incubated for 10 min, transferred to 4 mm quartz EPR tubes, and flash-frozen in liquid nitrogen. For •NO concentration-dependent samples, a stock of 2 mM DEA NONOate was prepared in anaerobic buffer and allowed to incubate for 1 hour. To ensure accurate concentrations, a hemoglobin assay was used to determine free •NO in solution.[3] Samples were prepared by adding increasing amounts of •NO per the protocol described above, with an increase in the incubation time to 30 min.

X-band continuous-wave EPR measurements were carried out on a Bruker EMX-Plus spectrometer equipped with a ColdEdge Stinger 4 K closed-cycle helium system and an ER4119HS resonator. Spectra were recorded at 9.35 GHz with a microwave power of 1.0 mW, a field modulation frequency of 100 kHz, and a modulation amplitude of 0.4 mT. Four scans were averaged for each sample. All spectra were collected at 10 K. *g*-values were determined by inspection of the EPR line shape. Power saturation studies of the low-spin species were performed at 20 K over a microwave power range of 0.20 μW to 50.24 mW.

*Kinetic assays using $O_2$ electrode*

For all kinetic experiments, MSDO was freshly prepared by iron reconstitution. The as-purified protein was incubated with 10 mM EDTA (pH 8.0) overnight at 4 °C with gentle stirring. The following day, the protein was concentrated and desalted into reaction buffer (50 mM Tris-HCl and 100 mM NaCl at pH 7.6), re-concentrated, and rendered anaerobic using an argon/vacuum manifold. Under anaerobic conditions in a glovebox, 5 equivalents of ascorbic acid and 2 equivalents of ferrous ammonium sulfate were added slowly to the protein. After 20 min of incubation on ice, the enzyme was subsequently desalted into reaction buffer using a centrifugal filter inside the glovebox, then sealed in an anaerobic vial until use.

Kinetic assays were performed using an $O_2$ electrode (Oxygraph+, Hansatech). Racemic MS, (*R*)-MS, and (*S*)-MS were tested over concentrations ranging from 10 µM to 1 mM. Reactions were initiated by adding 1 µM MSDO from the sealed vial via an air-tight syringe to 0.5 mL of substrate solution in reaction buffer, and $O_2$ consumption was recorded to obtain the initial rates. All reactions were conducted in triplicate, and the results were fitted to Michaelis–Menten curves in Origin to obtain kinetic parameters.

#### *Crystallization and* in crystallo *reactions*

Iron-reconstituted MSDO, prepared as described above, was used for crystallization. The protein was purified on a HiLoad 16/600 Superdex-75 pg column (Cytiva) equilibrated with anaerobic 50 mM Tris-HCl, 50 mM NaCl, pH 7.5, and concentrated to 30 mg/mL. Crystals were grown using the hanging-drop vapor diffusion method by mixing 2.0 µL of protein with 1.5 µL of crystallization solution containing 0.1 M sodium acetate (pH 4.5), 0.2 M $Li_2SO_4$, and 30% (w/v) PEG 8000. Trays were stored at 12 °C, and thin, clear needle clusters appeared after 2 weeks.

For substrate-bound crystals, all steps were performed strictly anaerobically in a glovebox. MSDO was concentrated to 40 mg/mL after Superdex-75 purification. A different crystallization condition was used to avoid sodium acetate binding at the active site, which blocks MS binding. The optimized solution contained 0.1 M Bis-Tris (pH 5.5), 0.2 M $MgCl_2$, and 25% (w/v) PEG 3350. Anaerobically prepared (*R*)-MS and (*S*)-MS were added to the enzyme at a final concentration of 2 mM for co-crystallization. Crystals were grown using the same setup as above, then soaked in mother liquor supplemented with 20% glycerol as a cryoprotectant before flash-freezing in liquid nitrogen for data collection.

For *in crystallo* reaction experiments, co-crystals of MSDO with (*R*)-MS were picked in a glovebox and incubated in air-containing mother liquor for defined times prior to cryoprotection and flash-freezing in liquid nitrogen. A monooxygenated intermediate was obtained after 20 s of incubation, while a dioxygenated product formed after 30 s and remained stably bound in the active site until crystal loss of diffraction quality.

#### *X-ray crystallographic data collection and structure determination*

Crystallographic data were collected at ALS beamline 5.0.1 (resting-state structure), NSLS-II beamline 19-ID (substrate-bound structures), APS beamline 22-ID (monooxygenated intermediate structure), and ALS beamline 8.2.2 (dioxygenated intermediate structure). Data were integrated and scaled using HKL2000,[4] and molecular replacement was performed with an AlphaFold-predicted MSDO model[5] as the search model. Structure building and refinement were carried out using PHENIX and COOT.[6, 7] Details of data collection and refinement statistics are summarized in Table S2.

Ligand refinement was performed in the final stages of model refinement. Initial placement was guided by ligand-omit difference maps, with (*R*)-MS as the starting ligand model in all cases. Where necessary, alternative ligand models were fit into the $2F_{obs}–F_{calc}$ electron density maps, with corresponding $F_{obs}–F_{calc}$ maps examined for verification. Geometry restraints for monooxygenated and dioxygenated derivatives were generated using Phenix eLBOW. Ligand occupancies were refined to 1.0 for the (*R*)-MS, (*S*)-MS, monooxygenated intermediate, and dioxygenated intermediate structures. The $2F_{obs}–F_{calc}$ maps were contoured at 1.0 σ to evaluate ligand electron density. For omit-map validation, ligands were removed from the final models, and the resulting $F_{obs}–F_{calc}$ maps contoured at 3.0 σ confirmed the presence and placement of the active-site ligands.

*Computational approaches: Enzyme model set-up and MD simulations*

Primary model preparation: The crystal structure coordinates of the (*R*)- and (*S*)-bound mercaptosuccinate structures (9NYW and 9NYV) were used to build the primary model for the molecular dynamics (MD) simulations. The PDB structures only contained one chain, hence chain A was selected for our work. Hydrogen atoms were added to both structures under pH 7.5 conditions in PropKa v3.2 and PDB2PQR v3.7.1.[8] At this pH, all carboxylate groups in the protein are deprotonated and all Arg and Lys residues are protonated. Histidine residues were visually inspected and adjusted resulting in the following protonation states for both protein structures: His63, His93, His95, His109, His149, His161 and His163 were protonated on $N_\delta$.

Force field parameterization: Forcefield parameters for the non-standard iron center were derived using the bonded model approach via the Metal Center Parameter Builder (MCPB.py) as implemented in AmberTools25.[9] The two sets of truncated models comprising the metal center and its first-coordination sphere ligands were geometry-optimized in the gas phase at the UB3LYP/def2-SVP level of theory using Gaussian-09.[10-13] The small model was used to derive bonding parameters (force constants and equilibrium values) via the Seminario method, while the large model employed to assign partial atomic charges using the Restrained Electrostatic Potential (RESP) fitting scheme. Protein residues were modeled using the ff19SB force field, and non-protein components were described using the Generalized Amber Force Field (GAFF2).[14, 15] The system was solvated in a truncated octahedral box of OPC water molecules with a 10.0 Å buffer, neutralized with sodium ($Na^+$) and chloride ($Cl^–$) ions on the surface via the TLEaP module and mimic a physiological ionic strength of 0.15 M.

MD Simulations: Subsequently, the system was subjected to multi-stage energy minimization algorithm protocol consisted of seven stages of restrained minimization where positional restraints on heavy atoms, solute, solute + solvent and backbone + cofactor were gradually reduced (force constants: 50, 250, 25, 10, 1.0 and 0.1 kcal $mol^{–1}$ $Å^{–2}$) by a final unrestrained minimization with a total of 20,000 steps and the protein chains fixed, consisting of 5,000 steps of steepest descent followed by 15,000 steps of conjugate gradient algorithm Thereafter, the system was gradually heated from 0 K to 300 K over 200 ps 400,000 steps using a time step of 0.5 fs. During this stage, harmonic positional restraints were applied to all heavy solute atoms + backbone (force constant: 5.0 to 2.0 kcal $mol^{–1}$ $Å^{–2}$) with the protein backbone fixed. The Langevin thermostat ($\gamma$ = 5.0 $ps^{–1}$), was used to control the system temperature during the heating stage. The system was then applied to a 1 ns NPT ensemble equilibration with positional restraints on heavy solute atoms whereby the constraints on the backbone atoms were gradually reduced. Finally, an NPT production run of 200 ns was performed in Amber,[16] at a constant temperature of 300 K and pressure of 1 bar for 50,000,000 steps using a time step of 0.2 fs, the temperature was regulated by Langevin thermostat with a collision frequency of $\gamma$ of 2.0 $ps^{-1}$ and the constant pressure was controlled via Monte Carlo barostat. Detailed analyses resulted from MD simulation are shown in Figure S15–S26.

*Computational approaches: Quantum mechanics calculations*

QM Cluster Model Set-Up: Cluster models were set-up as described previously.[17] Based on the average structure of the MD simulation on the (*S*)-isomer and (*R*)-isomer bound enzyme structures, we created cluster models $^5$**Re**$_S$ and $^5$**Re**$_R$, which are Fe(III)-superoxo complexes with substrate-bound. A large environment surrounding the metal active site and the substrate was taken into consideration as described in Figure S23 including six water molecules. We selected the same protein environment for both isomers. Overall, the (*S*)-bound structure has 287 atoms and total charge of Q = 1, while the (*R*)-bound structure has 298 atoms and Q = +2. The two enzyme models contain the three His ligands bound to the iron with the connecting protein included and $Asn_{94}$ truncated to Gly. Several additional protein chains were included namely Gln64-Ala65-Arg66, Tyr80-Leu81-Asn82-Val83-Met84, and His163-Met164-Tyr165 with Ala65, Leu81, Val83 and Met164 abbreviated to a Gly residue. In addition, the side chains of Ile101, Leu169 and Arg175 were included in the models.

DFT Approaches: Calculations were performed using density functional theory approaches as implemented in the Gaussian-09 software package.[11] All calculations utilize the unrestricted B3LYP-GD3BJ method that includes Becke-Johnson corrected dispersion corrections.[10, 12, 18] Geometries were optimized using the LANL2DZ with core potential on iron and 6-31G* on the rest of the atoms.[19, 20] Single point calculations were performed with the LACV3P+ plus core potential on iron and 6-311+G* on the rest of the atoms. Absolute energies (E), zero-point corrections (ZPE) and free energies (G) of DFT optimized geometries (*S*)- and (*R*)-MS are shown in Tables S7 and S8, respectively. Figures S27–S30 show the B3LYP-D3BJ optimized geometries of the proposed intermediates and transition states for the (*S*)-configuration in various spin states; the corresponding (*R*)-configuration structures are shown in Figures S31–S34.

**Table S1. Steady-state kinetic parameters determined for MSDO using $O_2$ electrode**

| | $V_{max}$ (μM min$^{-1}$) | $K_m$ (μM) | $k_{cat}$ (s$^{-1}$) | $k_{cat}/K_m$ (mM$^{-1}$ s$^{-1}$) |
|---|---|---|---|---|
| ***rac*-MS** | 750 ± 10 | 151 ± 8 | 12.5 ± 0.2 | 83 ± 5 |
| **(*R*)-MS** | 480 ± 20 | 110 ± 10 | 8.0 ± 0.3 | 73 ± 7 |
| **(*S*)-MS** | 1180 ± 70 | 160 ± 30 | 20 ± 1 | 130 ± 20 |

**Table S2. X-ray crystallography data collection and refinement statistics**

| | Resting state | (*S*)-MS bound | (*R*)-MS bound | Monooxygenated intermediate | Dioxygenated product |
|---|---|---|---|---|---|
| **PDB code** | **8VNI** | **9NYV** | **9NYW** | **9PYP** | **10AB** |
| **Data Collection** | | | | | |
| **Wavelength (Å)** | 0.97741 | 0.97860 | 0.97860 | 1.00000 | 0.97949 |
| **Space group** | $P2_1$ | $P2_12_12_1$ | $P2_12_12_1$ | $P2_12_12_1$ | $P2_12_12_1$ |
| **Cell dimensions** | | | | | |
| ***a, b, c*** **(Å)** | 41.2, 35.3, 72.3 | 35.4, 41.3, 131.8 | 35.2, 41.2, 133.0 | 35.4, 41.2, 133.1 | 35.2 41.8 133.9 |
| **α, β, ɣ (˚)** | 90.0, 94.3, 90.0 | 90.0, 90.0, 90.0 | 90.0, 90.0, 90.0 | 90.0, 90.0, 90.0 | 90.0, 90.0, 90.0 |
| **Resolution (Å)** | 50.00−1.57 (1.60−1.57)[a] | 50.00−2.06 (2.10−2.06)[a] | 50.00−2.20 (2.24−2.20)[a] | 50.00−2.01 (2.04−2.01)[a] | 50.00−1.87 (1.90−1.87)[a] |
| **Redundancy** | 6.4 (4.4) | 8.6 (5.6) | 9.5 (6.9) | 16.4 (12.6) | 9.9 (9.4) |
| **$R_{merge}$[b] (%)** | 11.2 (65.8) | 17.0 (59.8) | 17.9 (65.1) | 27.6 (96.3) | 11.2 (79.4) |
| ***I*/*σ*** | 15.5 (1.4) | 11.4 (2.32) | 27.5 (1.1) | 7.9 (4.4) | 9.6 (3.0) |
| **Completeness (%)** | 99.9 (98.2) | 99.3 (95.7) | 99.3 (96.1) | 97.6 (94.3) | 94.5 (97.2) |
| **$CC_{1/2}$, highest res. shell** | 0.68 | 0.82 | 0.83 | 0.87 | 0.87 |
| **Refinement** | | | | | |
| **Resolution (Å)** | 41.07−1.57 | 38.89−2.05 | 39.09−2.18 | 39.33−1.98 | 39.27−1.86 |
| **No. of reflections** | 29,415 | 12,258 | 10,067 | 13,535 | 15,681 |
| **$R_{work}$[c]/$R_{free}$ (%)** | 15.86/17.96 | 17.71/23.08 | 17.97/25.03 | 16.78/22.30 | 16.08/21.54 |
| **No. atoms/*B*-factors (Å$^2$)** | | | | | |
| **Protein** | 1548/17.51 | 1521/24.62 | 1527/32.10 | 1536/19.14 | 1543/20.87 |
| **Iron** | 1/11.84 | 1/21.23 | 1/27.78 | 1/13.39 | 1/18.08 |
| **Ligand[d]** | n. a. | 12/44.85 | 12/40.17 | 10/29.14 | 11/22.67 |
| **Solvent** | 292/31.40 | 127/26.19 | 116/31.28 | 191/27.21 | 199/29.47 |
| **Bond lengths (Å)** | 0.007 | 0.007 | 0.006 | 0.009 | 0.008 |
| **Bond angles (˚)** | 0.981 | 0.887 | 0.834 | 0.998 | 0.966 |

| | | | | | |
|---|---|---|---|---|---|
| **Ramachandran analysis** | | | | | |
| **Favored (%)** | 97.35 | 95.36 | 96.39 | 96.91 | 98.45 |
| **Allowed (%)** | 2.65 | 4.64 | 3.09 | 3.09 | 1.55 |
| **Outlier (%)** | 0.00 | 0.00 | 0.52 | 0.00 | 0.00 |

[a] Numbers in parentheses refer to data in the highest-resolution shell.

[b] $R_{merge} = \Sigma|I_h - \langle I_h \rangle|/\Sigma I_h$ , where $I_h$ is the observed intensity and $\langle I_h \rangle$ is the average intensity.

[c] $R_{work} = \Sigma||F_o| - k|F_c||/\Sigma|F_o|$.

[d] Ligand refers to the substrate, monooxygenated intermediate, or the dioxygenated product.

**Table S3. Relative energies and free energies (in kcal·mol$^{-1}$) of DFT optimized geometries along the MSDO dioxygenation mechanism of (*S*)-MS**

| | ΔE[BS1] | ΔE+ZPE | ΔG [BS1] | ΔE [BS2] | ΔE+ZPE | ΔG [BS2] |
|---|---|---|---|---|---|---|
| $^{5}$**Re**$_{S}$ | 0.00 | 0.00 | 0.00 | 0.00 | 0.00 | 0.00 |
| $^{3}$**Re**$_{S}$ | 3.56 | 3.16 | 6.29 | 8.52 | 8.11 | 11.24 |
| $^{1}$**Re**$_{S}$ | 5.21 | 7.28 | 9.44 | 11.83 | 13.89 | 16.05 |
| $^{7}$**Re**$_{S}$ | -0.58 | -1.16 | -2.11 | -0.09 | -0.67 | -1.62 |
| $^{5}$**TS1**$_{S}$ | 2.49 | 2.08 | 3.66 | 9.70 | 9.29 | 10.88 |
| $^{5}$**IM1**$_{S}$ | -27.44 | -25.48 | -21.62 | -14.45 | -12.48 | -8.63 |
| $^{5}$**TS2**$_{S}$ | -20.25 | -19.07 | -14.78 | -11.14 | -9.96 | -5.68 |
| $^{5}$**IM2**$_{S}$ | -42.53 | -41.04 | -36.68 | -34.47 | -32.98 | -28.62 |
| $^{5}$**TS3**$_{S}$ | -37.27 | -36.12 | -33.16 | -26.52 | -25.37 | -22.41 |
| $^{5}$**Prod**$_{S}$ | -85.53 | -82.38 | -78.58 | -67.91 | -64.76 | -60.97 |
| $^{3}$**IM1**$_{S}$ | -23.96 | -22.30 | -19.18 | -6.31 | -4.65 | -1.53 |
| $^{3}$**TS2**$_{S}$ | -21.44 | -20.69 | -17.33 | -4.00 | -3.25 | 0.11 |
| $^{3}$**IM2**$_{S}$ | -45.43 | -41.56 | -38.06 | -32.43 | -28.55 | -25.06 |
| $^{3}$**Prod**$_{S}$ | -72.59 | -68.63 | -65.79 | -52.91 | -48.95 | -46.12 |

**Table S4. Relative energies and free energies (in kcal·mol$^{-1}$) of DFT optimized geometries along the MSDO dioxygenation mechanism of (*R*)-MS**

| | DE [BS1] | DE+ZPE | DG [BS1] | DE [BS2] | DE+ZPE | DG [BS2] |
|---|---|---|---|---|---|---|
| $^{5}$**Re**$_{R}$ | 0.00 | 0.00 | 0.00 | 0.00 | 0.00 | 0.00 |
| $^{3}$**Re**$_{R}$ | 19.43 | 19.56 | 19.11 | 15.16 | 15.29 | 14.84 |
| $^{1}$**Re**$_{R}$ | -0.09 | 1.11 | 3.05 | -1.86 | -0.67 | 1.28 |
| $^{7}$**Re**$_{R}$ | -2.85 | -3.13 | -6.05 | -4.33 | -4.60 | -7.53 |
| $^{5}$**TS1**$_{R}$ | 17.37 | 17.29 | 18.20 | 15.76 | 15.69 | 16.59 |
| $^{5}$**IM1**$_{R}$ | -12.25 | -10.72 | -10.62 | -8.60 | -7.07 | -6.97 |
| $^{5}$**TS2**$_{R}$ | -5.98 | -5.28 | -4.89 | -6.72 | -6.02 | -5.63 |
| $^{5}$**IM2**$_{R}$ | -25.61 | -24.61 | -22.58 | -34.61 | -33.60 | -31.57 |
| $^{5}$**TS3**$_{R}$ | -17.23 | -17.49 | -17.10 | -25.58 | -25.83 | -25.45 |
| $^{5}$**Prod**$_{R}$ | -73.14 | -70.66 | -71.30 | -70.25 | -67.77 | -68.41 |

**Table S5. Group spin densities and charges of DFT optimized geometries along the MSDO dioxygenation mechanism of (*S*)-MS**

Sub = substrate and Prot = protein residues.

| | Spin densities | | | | | Charges | | | | |
|---|---|---|---|---|---|---|---|---|---|---|
| | Fe | OO | Sub | Prot | Total | Fe | OO | Sub | Prot | Total |
| $^{5}$**Re**$_{S}$ | 3.88 | -0.84 | 0.76 | 0.19 | 4.00 | 0.48 | -0.53 | -1.07 | 2.12 | 1.00 |
| $^{5}$**TS1**$_{S}$ | 3.72 | -0.68 | 0.84 | 0.12 | 4.00 | 0.50 | -0.54 | -1.03 | 2.07 | 1.00 |
| $^{5}$**IM1**$_{S}$ | 3.72 | 0.08 | 0.09 | 0.11 | 4.00 | 0.55 | -0.86 | -0.72 | 2.03 | 1.00 |
| $^{5}$**TS2**$_{S}$ | 3.75 | 0.18 | -0.06 | 0.13 | 4.00 | 0.60 | -1.03 | -0.63 | 2.06 | 1.00 |
| $^{5}$**IM2**$_{S}$ | 3.72 | 0.38 | -0.27 | 0.17 | 4.00 | 0.49 | -1.23 | -0.37 | 2.11 | 1.00 |
| $^{5}$**TS3**$_{S}$ | 3.75 | 0.33 | -0.24 | 0.16 | 4.00 | 0.54 | -1.30 | -0.33 | 2.08 | 1.00 |
| $^{5}$**Pr**$_{S}$ | 3.73 | 0.06 | 0.08 | 0.13 | 4.00 | 0.63 | -1.45 | -0.22 | 2.04 | 1.00 |
| $^{3}$**Re**$_{S}$ | 3.37 | -1.21 | -0.24 | 0.08 | 2.00 | 0.42 | -0.41 | -1.14 | 2.12 | 1.00 |
| $^{3}$**IM1**$_{S}$ | 1.96 | 0.01 | 0.02 | 0.02 | 2.00 | 0.31 | -0.80 | -0.68 | 2.16 | 1.00 |
| $^{3}$**TS2**$_{S}$ | 2.00 | -0.04 | 0.02 | 0.02 | 2.00 | 0.33 | -0.85 | -0.67 | 2.19 | 1.00 |
| $^{3}$**IM2**$_{S}$ | 2.15 | 0.20 | -0.37 | 0.02 | 2.00 | 0.32 | -1.20 | -0.33 | 2.21 | 1.00 |
| $^{3}$**Pr**$_{S}$ | 1.96 | -0.02 | 0.03 | 0.03 | 2.00 | 0.49 | -1.43 | -0.18 | 2.12 | 1.00 |
| $^{7}$**Re**$_{S}$ | 3.85 | 1.18 | 0.76 | 0.21 | 6.00 | 0.48 | -0.49 | -1.08 | 2.10 | 1.00 |
| $^{1}$**Re**$_{S}$ | 1.34 | -1.09 | -0.19 | -0.06 | 0.00 | 0.31 | -0.47 | -1.09 | 2.25 | 1.00 |
| $^{1}$**IM1**$_{S}$ | 0.00 | 0.01 | 0.00 | 0.00 | 0.00 | 0.37 | -0.84 | -0.67 | 2.14 | 1.00 |
| $^{1}$**IM2**$_{S}$ | 0.62 | -0.40 | -0.22 | 0.00 | 0.00 | 0.28 | -1.21 | -0.32 | 2.25 | 1.00 |

**Table S6. Group spin densities and charges of DFT optimized geometries along the MSDO dioxygenation mechanism of (*R*)-MS**

Sub = substrate and Prot = protein residues.

| | Spin densities | | | | | Charges | | | | |
|---|---|---|---|---|---|---|---|---|---|---|
| | Fe | OO | Sub | Prot | Total | Fe | OO | Sub | Prot | Total |
| $^{5}$**Re**$_{R}$ | 2.09 | 1.92 | -0.04 | 0.03 | 4.00 | 0.30 | -0.01 | -1.35 | 3.06 | 2.00 |
| $^{3}$**Re**$_{R}$ | 1.60 | 0.46 | -0.04 | -0.01 | 2.00 | 0.33 | -0.23 | -1.32 | 3.21 | 2.00 |
| $^{1}$**Re**$_{R}$ | 1.72 | -1.63 | -0.06 | -0.03 | 0.00 | 0.33 | -0.16 | -1.31 | 3.15 | 2.00 |
| $^{7}$**Re**$_{R}$ | 3.70 | 1.96 | 0.18 | 0.16 | 6.00 | 0.48 | 0.01 | -1.50 | 3.01 | 2.00 |
| $^{5}$**TS1**$_{R}$ | 3.64 | 0.02 | 0.18 | 0.16 | 4.00 | 0.48 | -0.02 | -1.48 | 3.01 | 2.00 |
| $^{5}$**IM1**$_{R}$ | 3.72 | 0.12 | 0.03 | 0.13 | 4.00 | 0.62 | -0.84 | -0.79 | 3.02 | 2.00 |
| $^{5}$**TS2**$_{R}$ | 3.73 | 0.10 | 0.01 | 0.15 | 4.00 | 0.68 | -0.99 | -0.75 | 3.06 | 2.00 |
| $^{5}$**IM2**$_{R}$ | 3.06 | 0.66 | 0.14 | 0.13 | 4.00 | 0.46 | -1.16 | -0.51 | 3.21 | 2.00 |
| $^{5}$**TS3**$_{R}$ | 3.74 | 0.23 | -0.15 | 0.19 | 4.00 | 0.59 | -1.31 | -0.39 | 3.11 | 2.00 |
| $^{5}$**Prod**$_{R}$ | 3.74 | 0.06 | 0.06 | 0.15 | 4.00 | 0.67 | -1.43 | -0.28 | 3.03 | 2.00 |

**Table S7. Absolute energies (E), zero-point corrections (ZPE) and free energies (G) of DFT optimized geometries along the MSDO dioxygenation mechanism of (*S*)-MS**

All data are given in au at B3LYP/BS1 level of theory and single point B3LYP/BS2 results.

| | E [BS1, au] | ZPE [BS1, au] | G [BS1, au] | E [BS2, au] |
|---|---|---|---|---|
| $^{5}$**Re**$_{S}$ | -7640.040415 | 2.440590 | -7637.816249 | -7642.050282 |
| $^{3}$**Re**$_{S}$ | -7640.034735 | 2.439944 | -7637.806232 | -7642.036707 |
| $^{1}$**Re**$_{S}$ | -7640.032108 | 2.443884 | -7637.801203 | -7642.031437 |
| $^{7}$**Re**$_{S}$ | -7640.041341 | 2.439672 | -7637.819608 | -7642.050425 |
| $^{5}$**TS1**$_{S}$ | -7640.036446 | 2.439933 | -7637.810411 | -7642.034818 |
| $^{5}$**IM1**$_{S}$ | -7640.084145 | 2.443718 | -7637.850706 | -7642.073305 |
| $^{5}$**TS2**$_{S}$ | -7640.072685 | 2.442477 | -7637.839808 | -7642.068041 |
| $^{5}$**IM2**$_{S}$ | -7640.108194 | 2.442960 | -7637.874708 | -7642.105207 |
| $^{5}$**TS3**$_{S}$ | -7640.099811 | 2.442420 | -7637.869090 | -7642.092544 |
| $^{5}$**Prod**$_{S}$ | -7640.176713 | 2.445607 | -7637.941477 | -7642.158507 |
| $^{3}$**IM1**$_{S}$ | -7640.078601 | 2.443232 | -7637.846814 | -7642.060339 |
| $^{3}$**TS2**$_{S}$ | -7640.074575 | 2.441780 | -7637.843865 | -7642.056654 |
| $^{3}$**IM2**$_{S}$ | -7640.112819 | 2.446762 | -7637.876906 | -7642.101958 |
| $^{3}$**Prod**$_{S}$ | -7640.156087 | 2.446900 | -7637.921095 | -7642.134602 |

**Table S8. Absolute energies (E), zero-point corrections (ZPE) and free energies (G) of DFT optimized geometries along the MSDO dioxygenation mechanism of (*R*)-MS**

All data are given in au at B3LYP/BS1 level of theory and single point B3LYP/BS2 results.

| | E [BS1, au] | ZPE [BS1, au] | G [BS1, au] | E [BS2, au] |
|---|---|---|---|---|
| $^{5}$**Re**$_{R}$ | -7775.681730 | 2.551281 | -7773.347959 | -7777.697462 |
| $^{3}$**Re**$_{R}$ | -7775.650763 | 2.551480 | -7773.317498 | -7777.673303 |
| $^{1}$**Re**$_{R}$ | -7775.681866 | 2.553180 | -7773.343096 | -7777.700422 |
| $^{7}$**Re**$_{R}$ | -7775.686277 | 2.550839 | -7773.357606 | -7777.704358 |
| $^{5}$**TS1**$_{R}$ | -7775.654050 | 2.551161 | -7773.318960 | -7777.672345 |
| $^{5}$**IM1**$_{R}$ | -7775.701254 | 2.553718 | -7773.364877 | -7777.711173 |
| $^{5}$**TS2**$_{R}$ | -7775.691259 | 2.552400 | -7773.355755 | -7777.708169 |
| $^{5}$**IM2**$_{R}$ | -7775.722548 | 2.552883 | -7773.383940 | -7777.752609 |
| $^{5}$**TS3**$_{R}$ | -7775.709195 | 2.550882 | -7773.375210 | -7777.738232 |
| $^{5}$**Prod**$_{R}$ | -7775.798287 | 2.555228 | -7773.461578 | -7777.809412 |

**Figure S1. LC-HRMS analysis of succinate formation in MSDO reactions with MS enantiomers**

MSDO reactions with (**a**) (*R*)-MS and (**b**) (*S*)-MS. Peaks marked with asterisks at *m/z* 117.0191(2) correspond to succinate (theoretical *m/z* = 117.0193) formed after spontaneous desulfination of the product SFS (not observed, theoretical *m/z* = 180.9812). Insets show zoomed-in views of these peaks. The predominant signals at *m/z* 112.9855(6) arise from TFA (theoretical *m/z* = 112.9850) carried over from the HPLC solvent used during enantiomer separation.

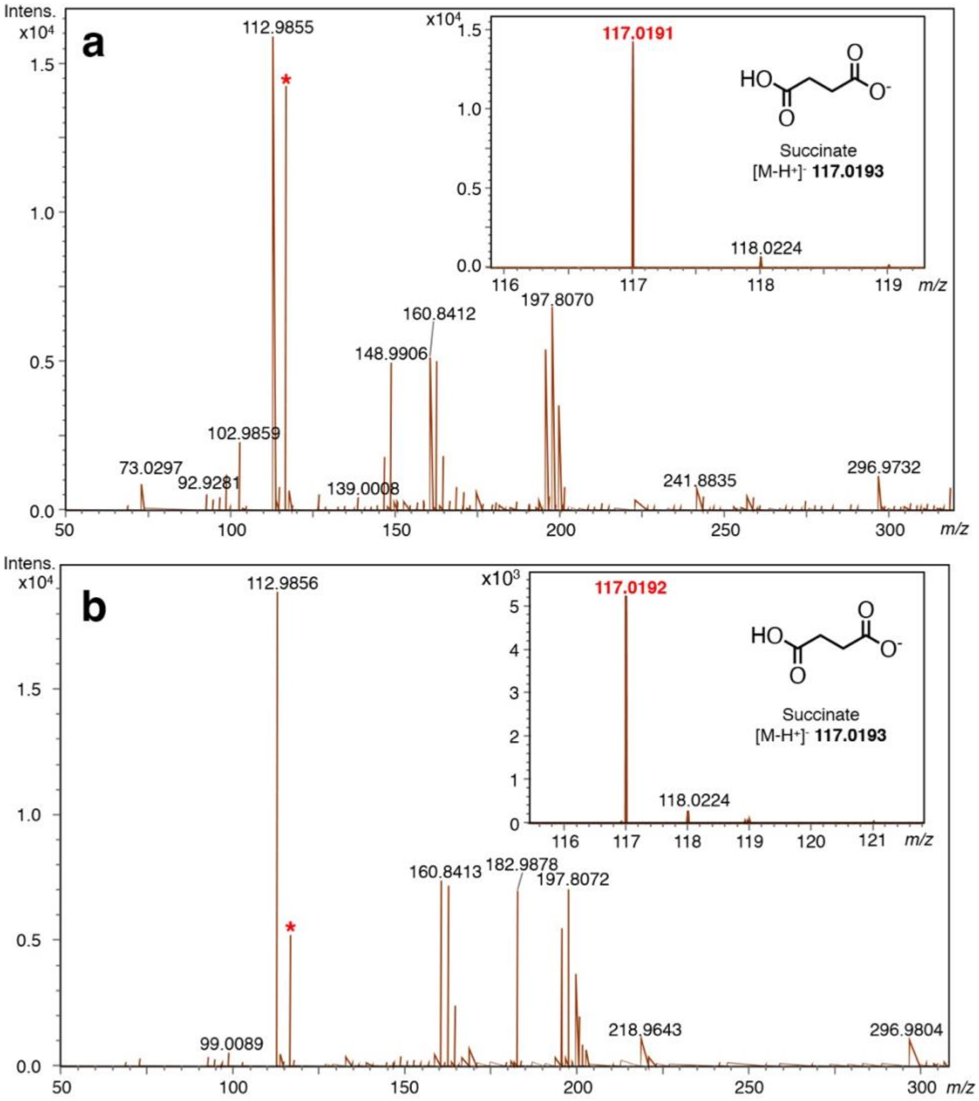

**Figure S2. $O_2$ consumption in MSDO reactions at varying substrate concentrations**

An $O_2$ electrode was used to monitor $O_2$ consumption in MSDO reactions containing 10, 20, 50, 100, 200, 500, 1000 μM of (**a**) (*R*)-MS and (**b**) (*S*)-MS. The reactions containing (*S*)-MS showed faster $O_2$ consumption than those with (*R*)-MS at the same concentrations.

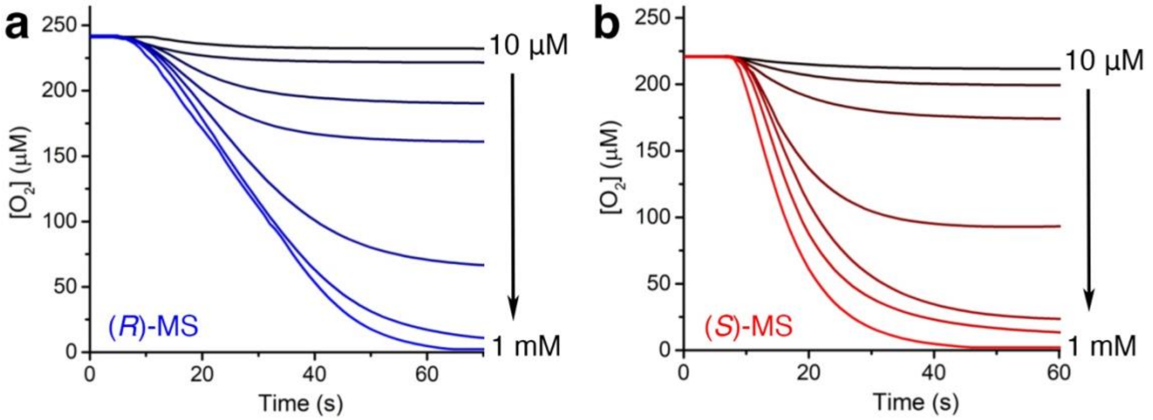

**Figure S3. EPR spectra of the nitrosyl complexes in MSDO**

(**a**) Full-scan EPR spectra of MSDO nitrosyl complexes recorded in the absence of the substrate (black) and in the presence of (*R*)-MS (blue), (*S*)-MS (red), or *rac*-MS (grey) at 10 K and 1 mW microwave power. A minor high-spin signal was observed in the substrate-free sample and was mostly diminished upon substrate addition. (**b**) Power saturation behavior of the (*R*)-MS-bound low-spin species monitored at $g = 2.26$ at 20 K. The power saturation curve (red) was fitted using the equation $\boldsymbol{I} = \boldsymbol{I_0}\frac{\boldsymbol{P}}{(\boldsymbol{P}+\boldsymbol{P_{1/2}})^{\boldsymbol{b}}}$ .

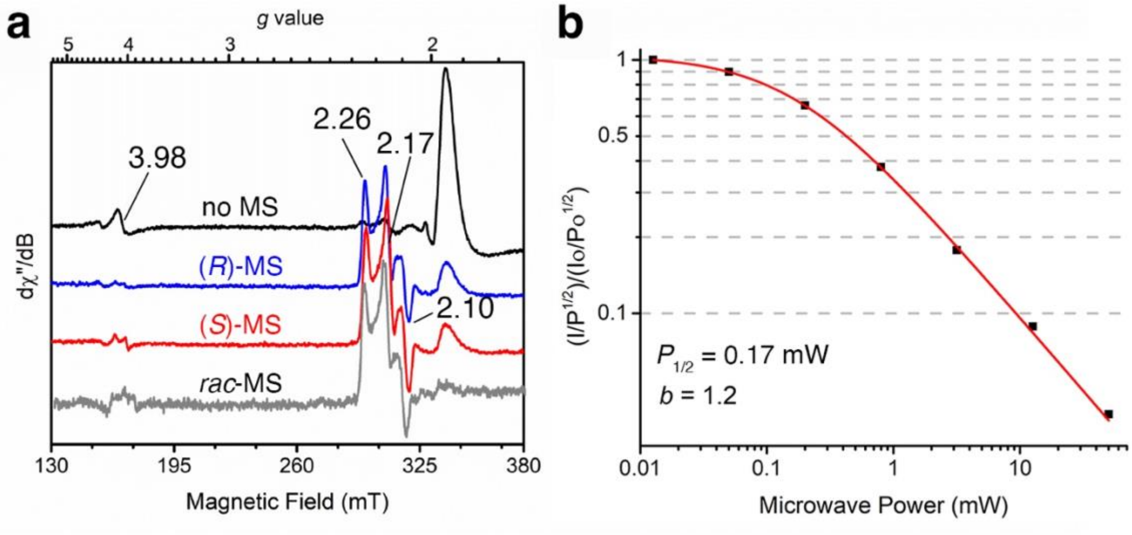

**Figure S4. EPR spectral simulation of low-spin nitrosyl species**

The EPR spectrum of low-spin nitrosyl complexes of (*R*)-MS-bound MSDO was simulated with two components using the Pepper function in EasySpin (Matlab) and least-squares fitting. From top to bottom: experimental data, simulated sum, Species 1, and Species 2. Species 1 and 2 contribute 64.7% and 35.3% of the total signal intensity, respectively.

Species 1: $S = 1/2$; $g_x$, $g_y$, and $g_z$ = 2.259, 2.169, and 2.098; $\sigma_{gx}$, $\sigma_{gy}$, and $\sigma_{gz}$ = 0.020, 0.022, and 0.025; lw = 0.55 mT; weight = 64.7%.

Species 2: $S = 1/2$; $g_x$, $g_y$, and $g_z$ = 2.185, 2.098, and 2.025; $\sigma_{gx}$, $\sigma_{gy}$, and $\sigma_{gz}$ = 0.086, 0.062, and 0.045; lw = 0.51 mT; weight = 35.3%.

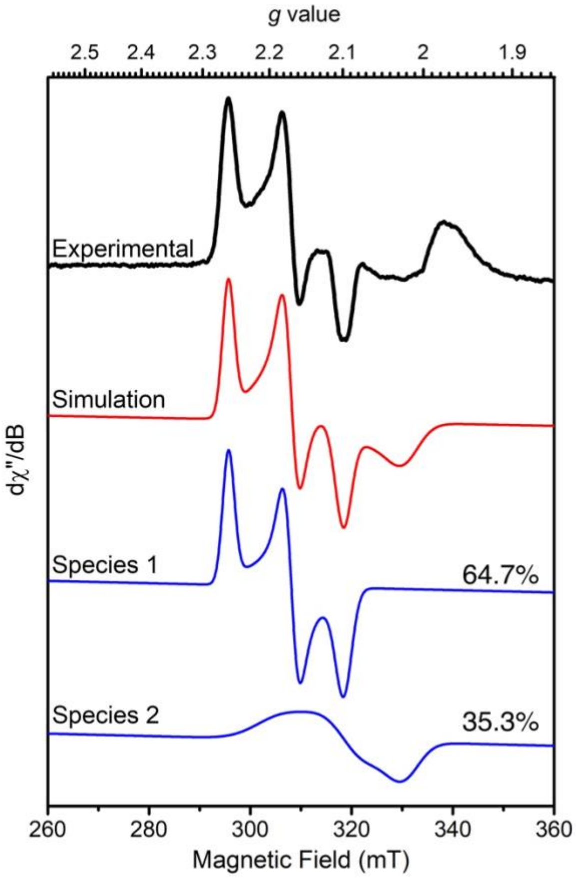

**Figure S5. Structural superposition of MSDO with other TDOs**

MSDO (grey, PDB: 8VNI) was superposed with (**a**) MDO (magenta, PDB:4TLF, chain A), (**b**) CDO (green, PDB: 6E87), (**c**) ADO (blue, PDB: 8U9I, chain A), and (**d**) PCO (yellow, PDB:6S7E). The resulting RMSD values are 1.863 over 140 Cα atoms, 2.779 over 157 Cα atoms, 1.742 over 91 Cα atoms, and 3.303 over 95 Cα atoms, respectively. All structural alignments and corresponding RMSD values were obtained using the Super command in PyMOL. In MSDO, CDO, and MDO, the *C*-terminal regions, shown as opaque loops, act as lids that close over the active-site entrance. In contrast, in ADO and PCO, this region forms part of the border of the β-barrel

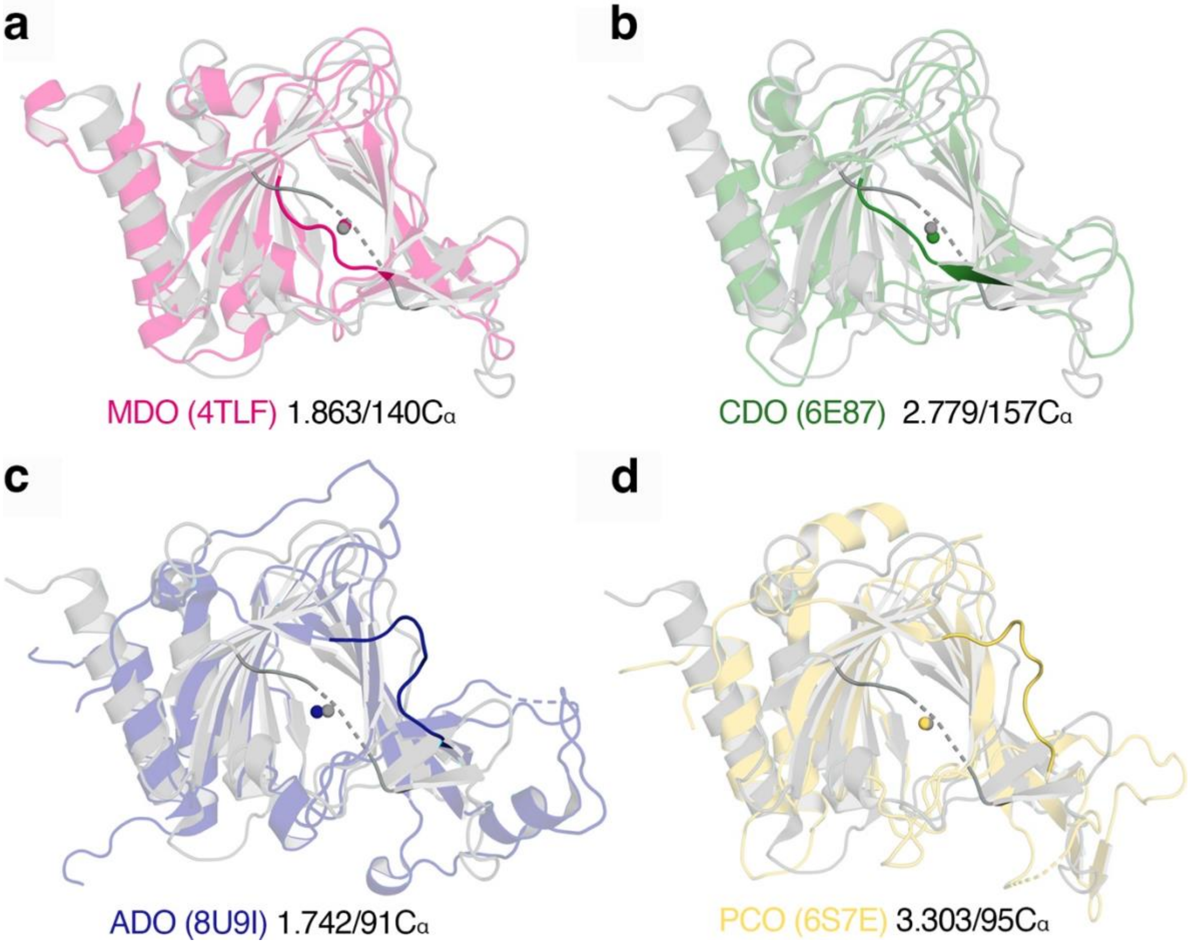

**Figure S6. Active site comparison between small thiol-oxidizing TDOs**

Structural comparison of (**a**) (*R*)-MS-bound MSDO, (**b**) *L*-Cys-bound CDO, and (**c**) 3-hydroxypropionic acid (substrate analog)-bound MDO. Substrates or analogs are shown as yellow sticks with red labels. Arg residues that interact with the substrate/analog are labeled in bold blue. Residues involved in the Tyr-Cys crosslink and the SHY motif are labeled in purple. *C*-terminal loop lids are shown as opaque loops. PDB codes are 9NYW (MSDO), 6N42 (CDO), and 6XB9 (MDO), respectively.

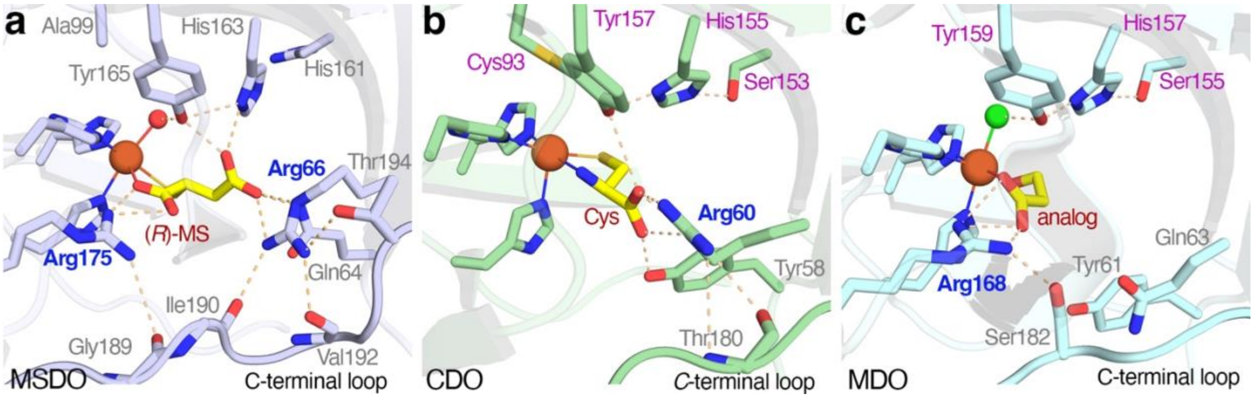

**Figure S7. Modeling of substrate-bound structures using alternative models**

Unsatisfactory fits were obtained when the active site of the (*S*)-MS-bound structure (PDB code of 9NYV) was modeled by (**a**) omitting the bound ligand, (**b**) placing four water molecules, or (**c**) placing (*R*)-MS and a water molecule. Similarly, unsatisfactory fits were obtained when the active site of the (*R*)-MS-bound structure (PDB code of 9NYW) was modeled by (**d**) omitting the bound ligand, (**e**) placing four water molecules, or (**f**) placing (*S*)-MS and a water molecule. The $2F_o$-$F_c$ electron density maps are contoured at 1σ (light blue), while the $F_o$-$F_c$ difference maps are contoured at +3σ and −3σ (green and red, respectively).

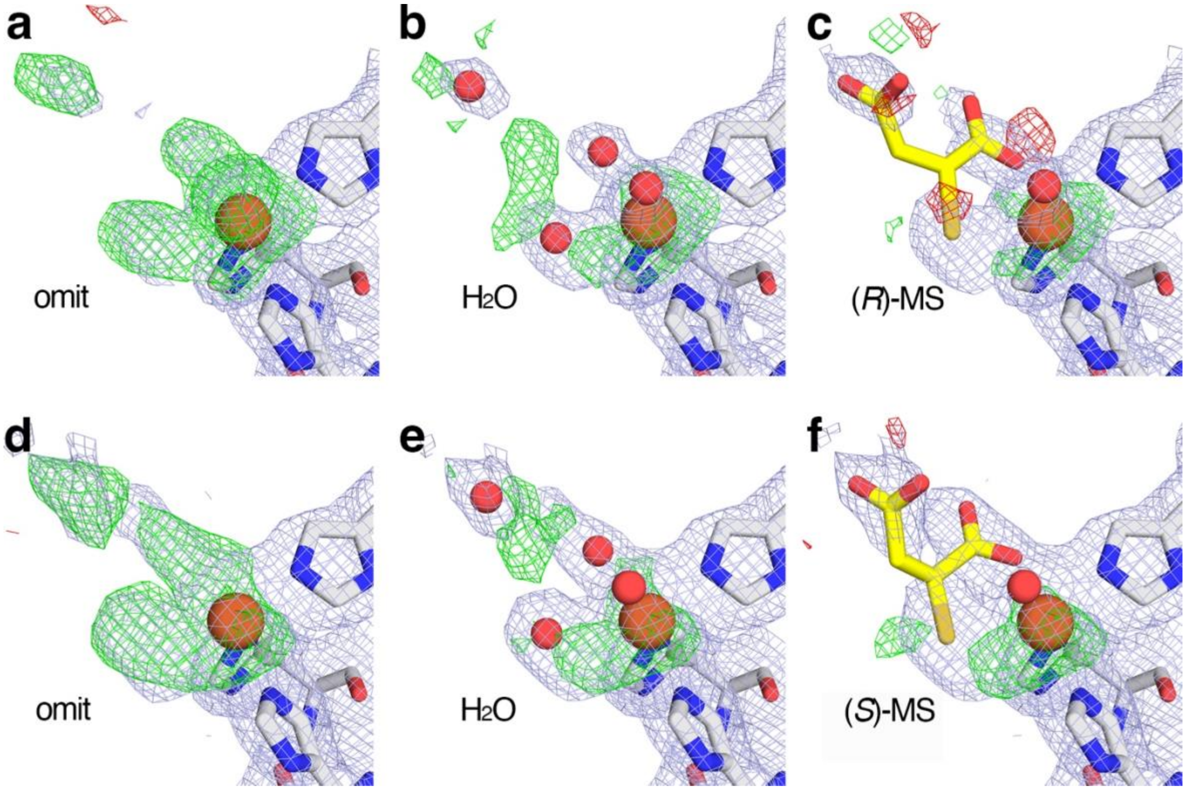

**Figure S8. Modeling of intermediate-bound structures using alternative models**

Unsatisfactory fits were obtained when the active site of the monooxygenated sulfenate intermediate-bound structure (PDB code of 9PYP) was modeled by (**a**) omitting the bound ligand, (**b**) placing (*R*)-MS and a water molecule, or (**c**) placing sulfinate and a water molecule. Similarly, unsatisfactory fits were obtained when the active site of the dioxygenated intermediate, sulfinate structure (PDB code of 10AB) was modeled by (**d**) omitting the bound ligand, (**e**) placing (*R*)-MS and a water molecule, or (**f**) placing sulfenate and an oxygen atom. The $2F_o$-$F_c$ electron density maps are contoured at 1σ (light blue), while the $F_o$-$F_c$ difference maps are contoured at +3σ and −3σ (green and red, respectively).

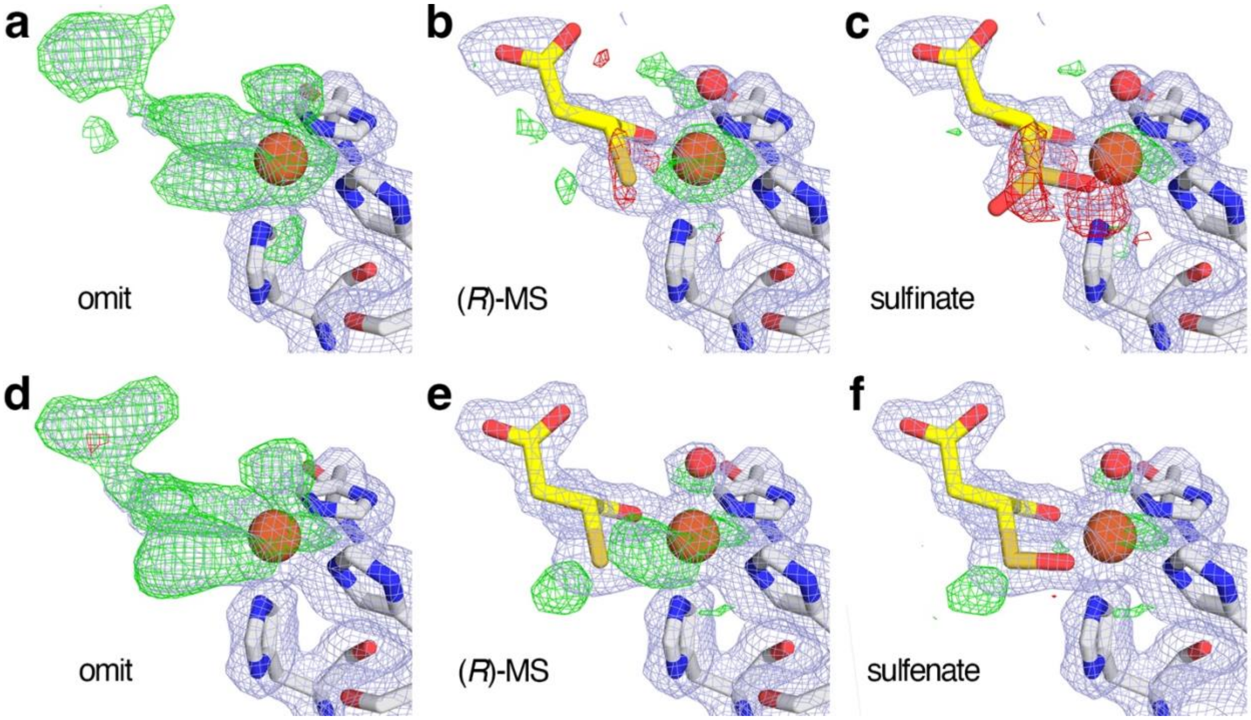

**Figure S9. Active-site superposition of the monooxygenated and dioxygenated intermediate structures**

The overall architecture of the active site is largely conserved between the monooxygenated and dioxygenated structures, with notable conformational differences observed only for the side chains of Lys88 and Arg175, which directly interact with the sulfinate moiety and the proximal carboxylate group, respectively. The monooxygenated and dioxygenated complexes are shown in white and blue (substrate in yellow), respectively, with key interactions depicted as gray and wheat dashed lines.

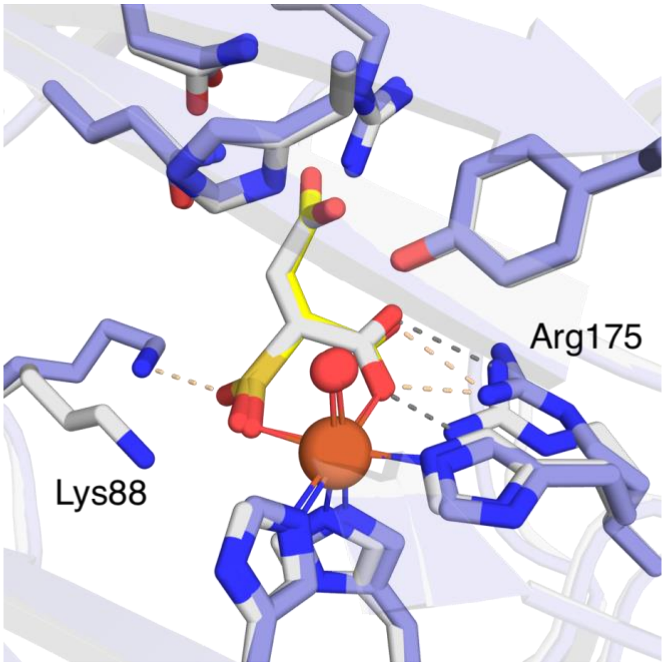

**Figure S10. Continuous hydrogen-bonded water chain in the active site**

In the dioxygenated product-bound structure, a continuous hydrogen-bonded water network connects the iron-coordinated water molecule to the bulk solvent. Water molecules are shown as red spheres, H-bonding interactions are indicated by dark dashed lines, and the cavity accommodating the water chain and the active site is depicted as a green pocket.

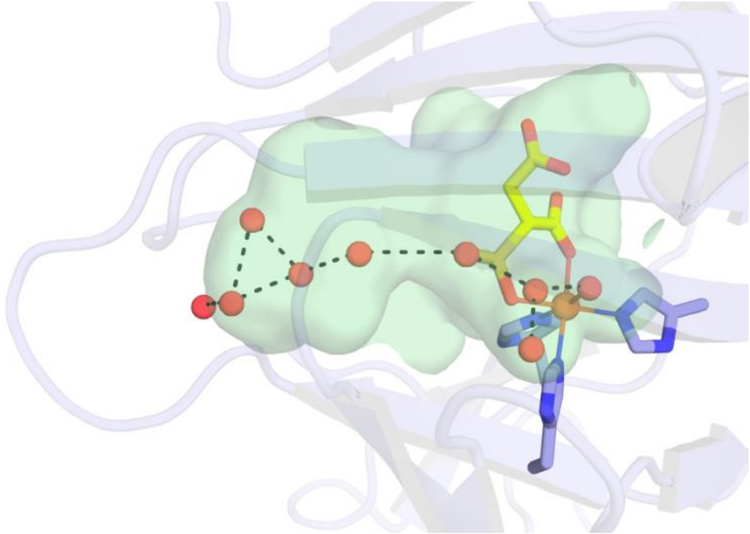

**Figure S11. DFT cluster models of $^{5}Re_S$ and $^{5}Re_R$ as studied in this work**

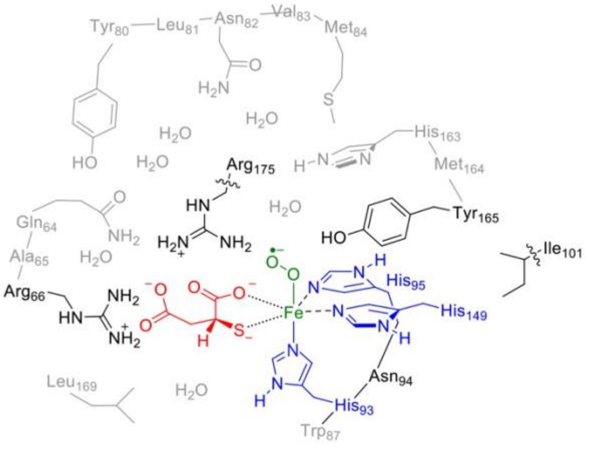

**$^{5}Re_S$**

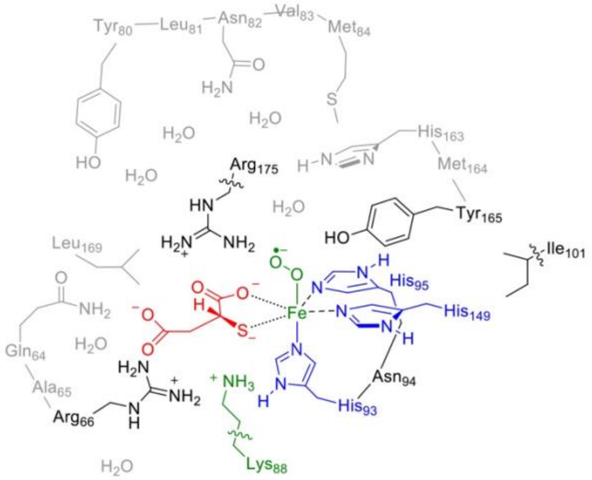

**$^{5}Re_R$**

**Figure S12. Overlays of the optimized geometries of $^{5}Re_S$ and $^{5}Re_R$ with the PDB files of 9NYV and 9NYW**

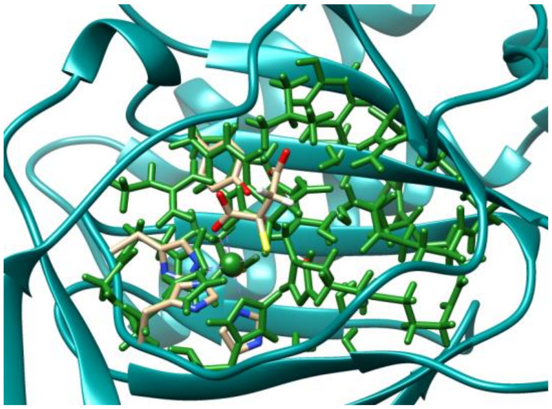

**Overlay 9NYW pdb and $^{5}Re_R$**

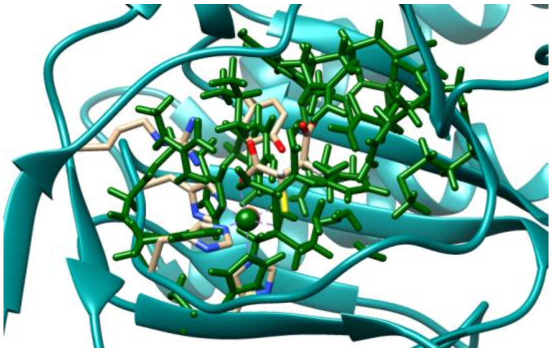

**Overlay 9NYV pdb and $^{5}Re_S$**

**Figure S13. UB3LYP-GD3BJ/BS1 calculated constraint geometry scan starting from $^{7}$Res on the right-hand-side**

Values are in kcal mol$^{-1}$ for the S−O$_d$ bond shortening. As shown, the scan does not lead to a local minimum and continuously increases in energy. Consequently, the oxygen atom transfer reaction cannot happen on the septet spin state surface for (*S*)-MS.

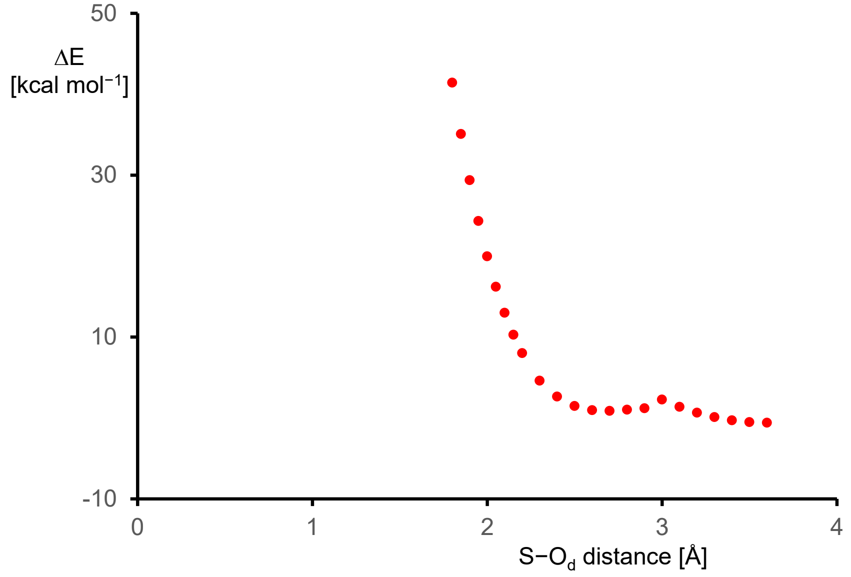

**Figure S14. UB3LYP-GD3BJ/BS1 calculated constraint geometry scan starting from $^{5}$Res on the right-hand-side**

Values are in kcal mol$^{-1}$ for the S−O$_d$ or S−O$_p$ bond shortening. As shown, the proximal oxygen atom attack is well higher in energy than distal oxygen atom attack. Consequently, the proximal oxygen atom attack mechanism and formation of the persulfenate intermediate is not part of the lowest energy mechanism.

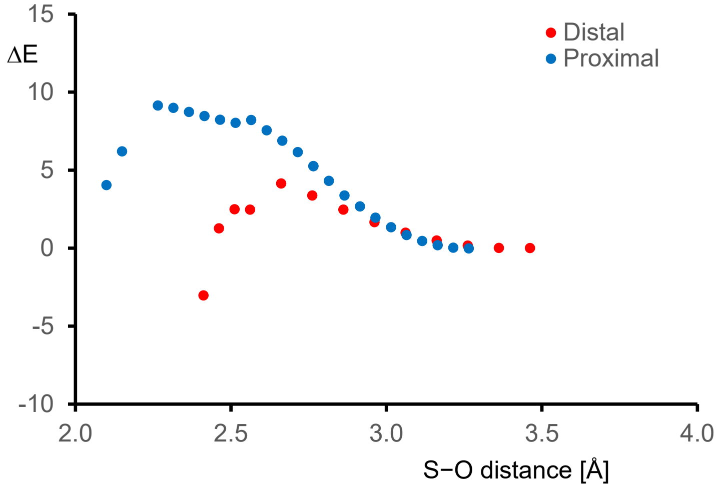

**Figure S15. RMSD profile of the 200 ns MD simulation on MSDO with (*S*)-MS bound**

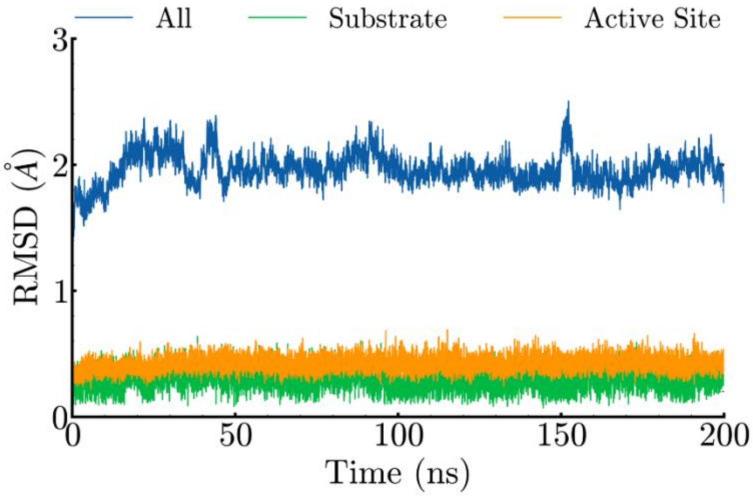

**Figure S16. RMSF pattern of the 200 ns MD simulation on MSDO with (*S*)-MS bound**

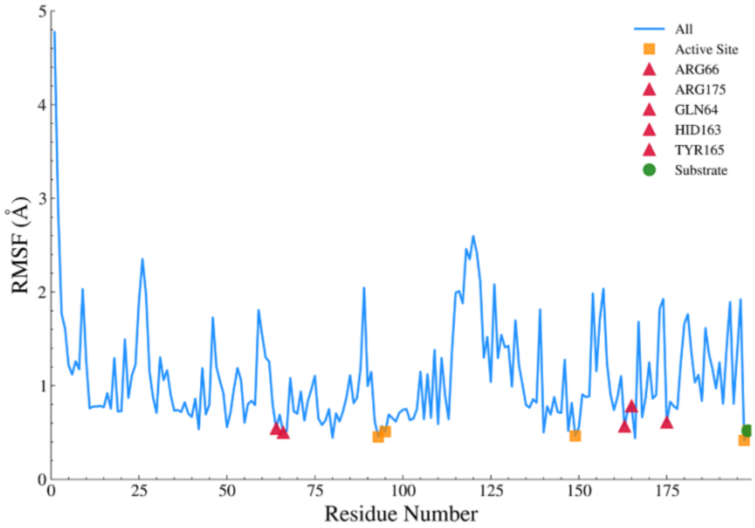

**Figure S17. Distance distribution between (*S*)-MS and amino acid residues for the 200 ns MD simulation**

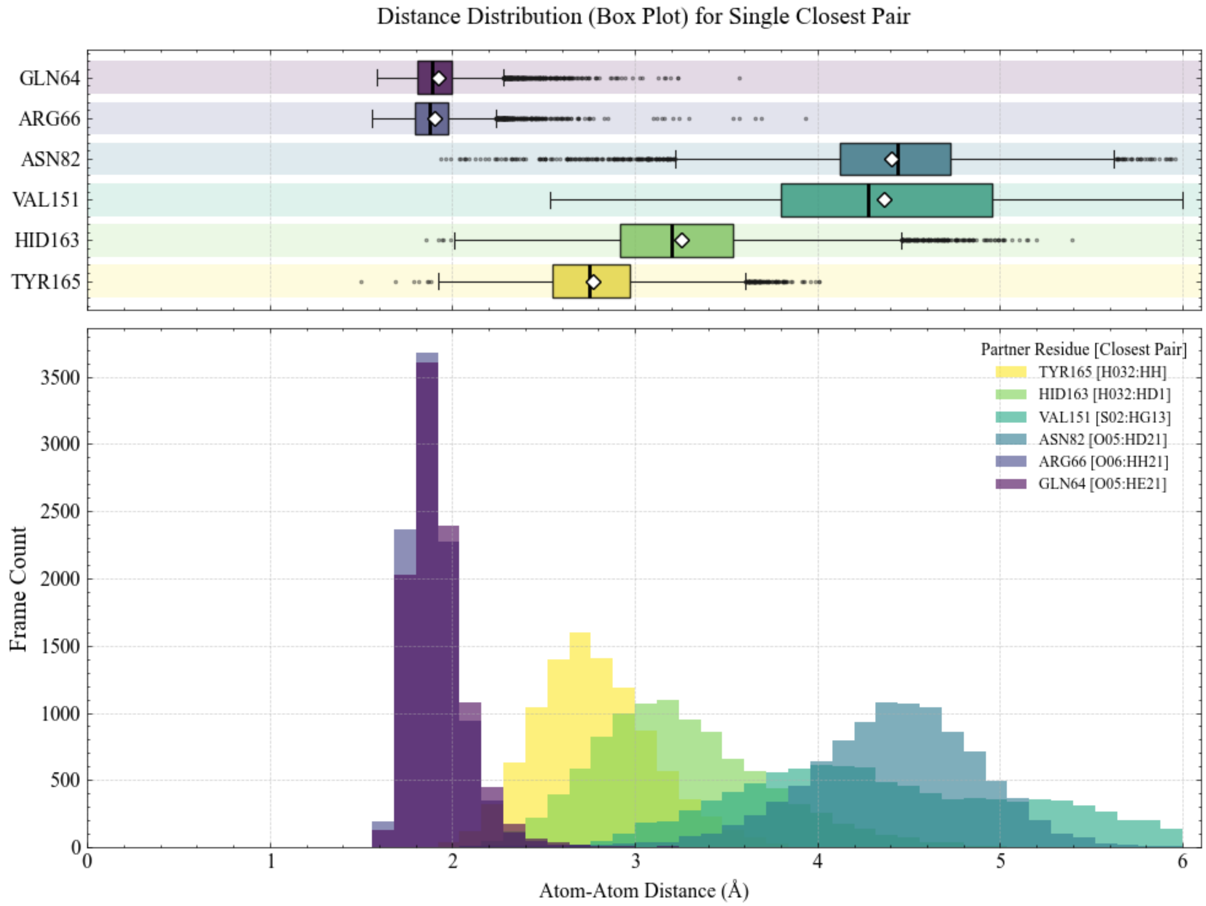

**Figure S18. (*S*)-MS ligand fingerprint map**

Thresholds over 10% were obtained from the MD simulation on (*S*)-MS bound MSDO.

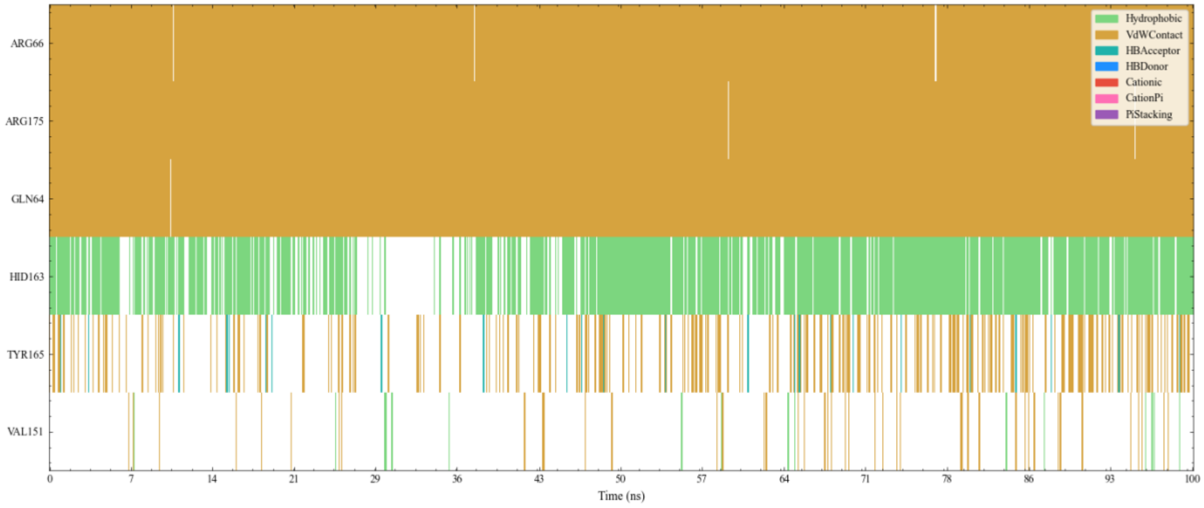

**Figure S19. (*S*)-MS substrate ligand heat map**

Threshold over 10% was obtained from the MD simulation on (*S*)-MS-bound MSDO.

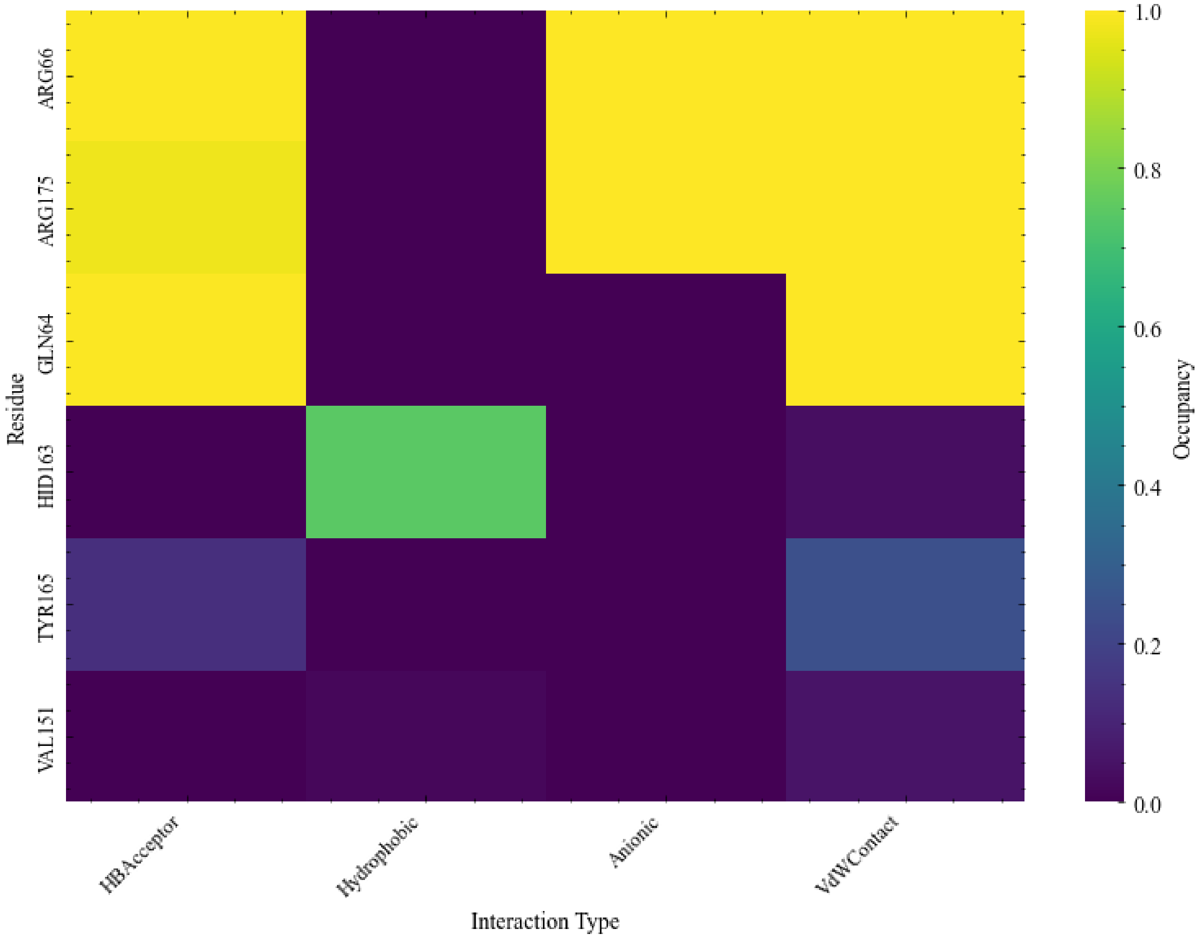

**Figure S20. RMSD profile of the 200 ns MD simulation on MSDO with (*R*)-MS bound**

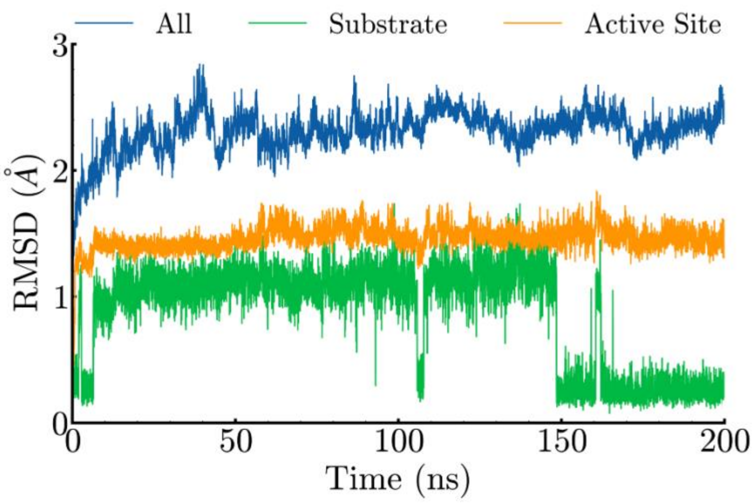

**Figure S21. RMSF pattern of the 200 ns MD simulation on MSDO with (*R*)-MS bound**

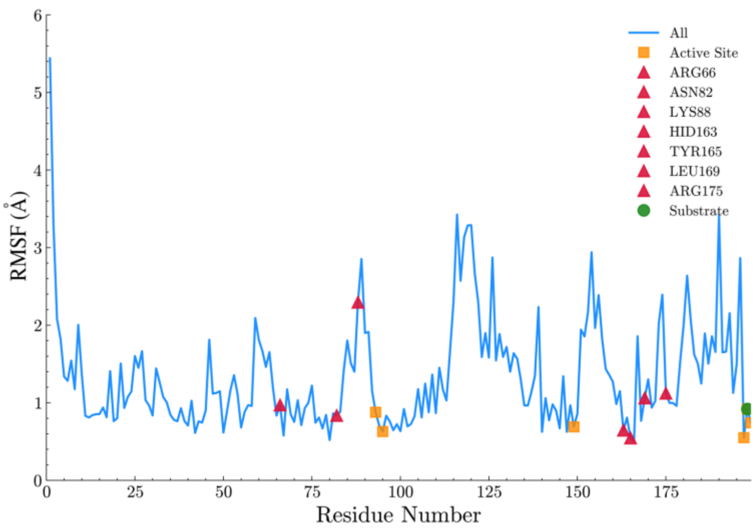

**Figure S22. Distance distribution between (*R*)-MS and amino acid residues for the 200 ns MD simulation**

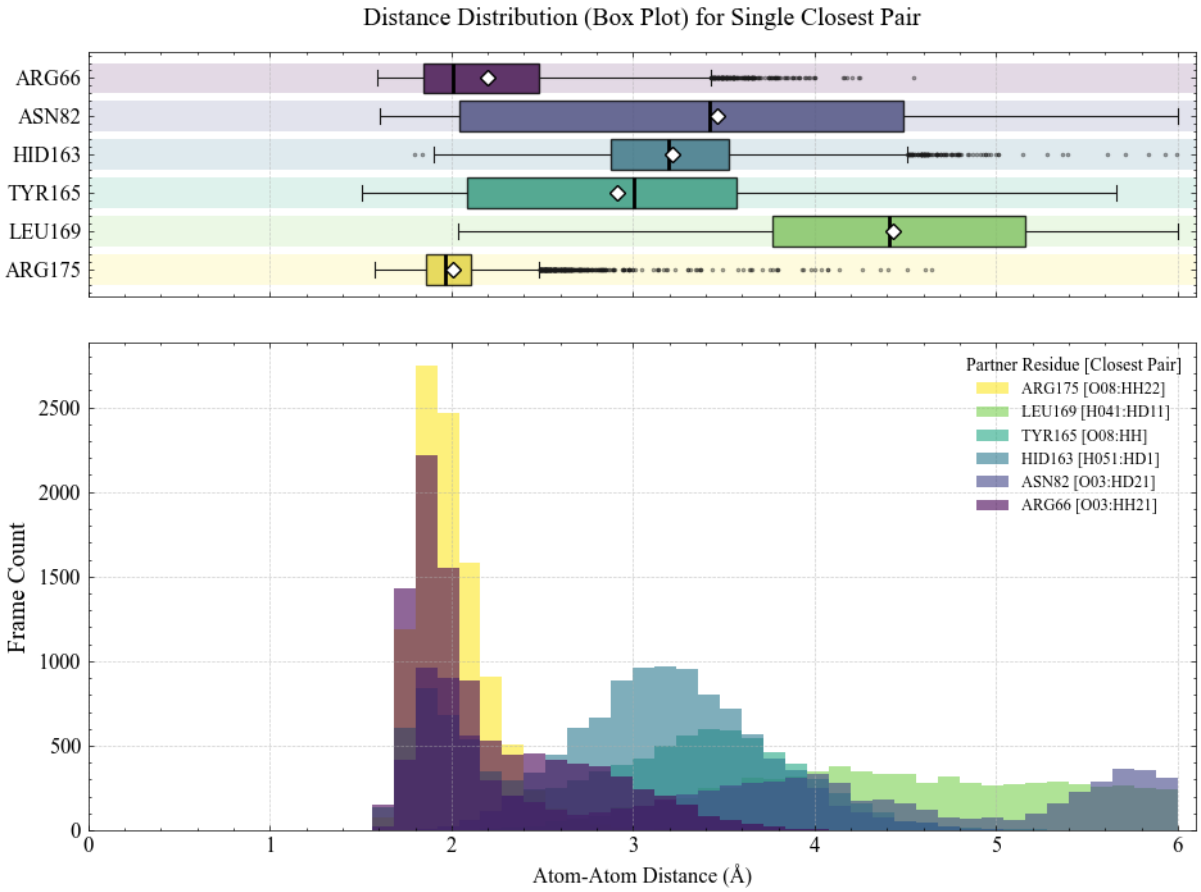

**Figure S23. (*R*)-MS substrate ligand heat map**

Threshold over 10% as obtained from the MD simulation of MSDO with (*R*)-MS bound.

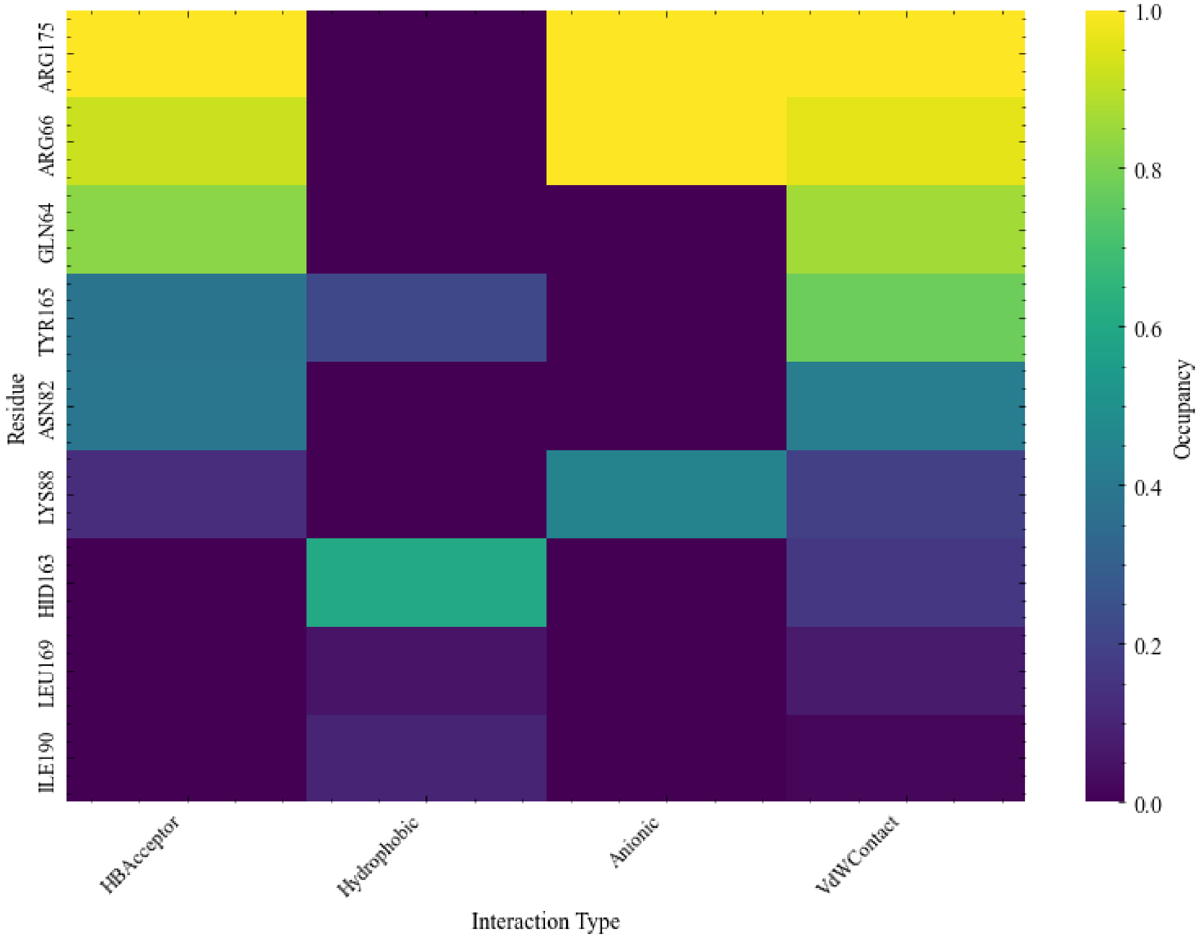

**Figure S24. (*R*)-MS ligand fingerprint map**

Threshold over 10% was obtained from the MD simulation on (*R*)-MS bound MSDO.

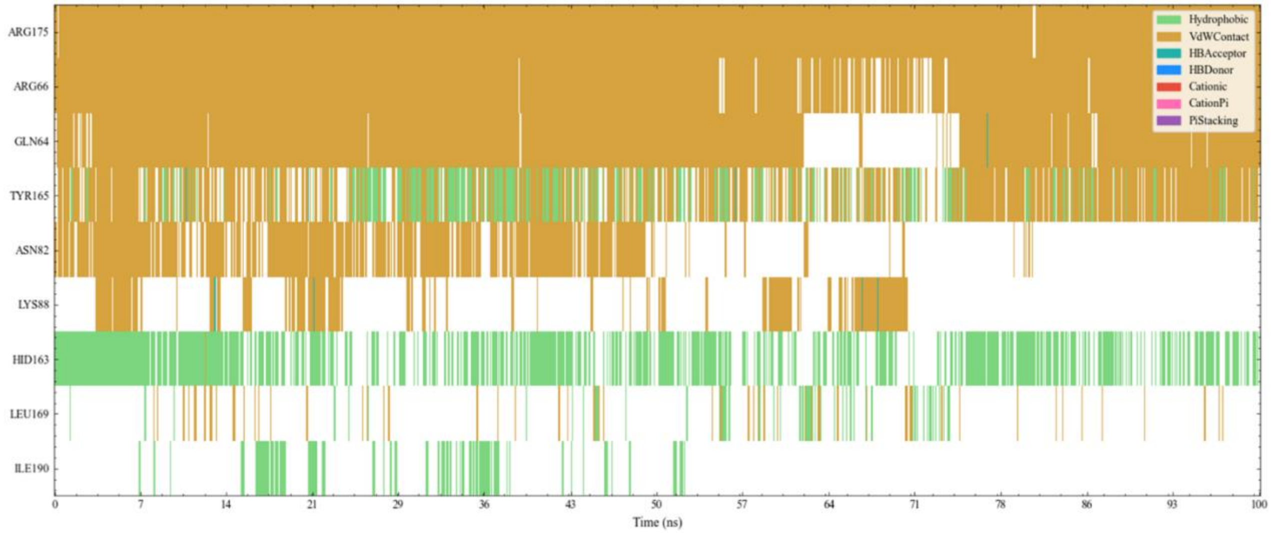

**Figure S25. Pairwise 2D RMSD heat map ($C_\alpha$ only) derived from the MD trajectory of (*S*)-MS-bound MSDO**

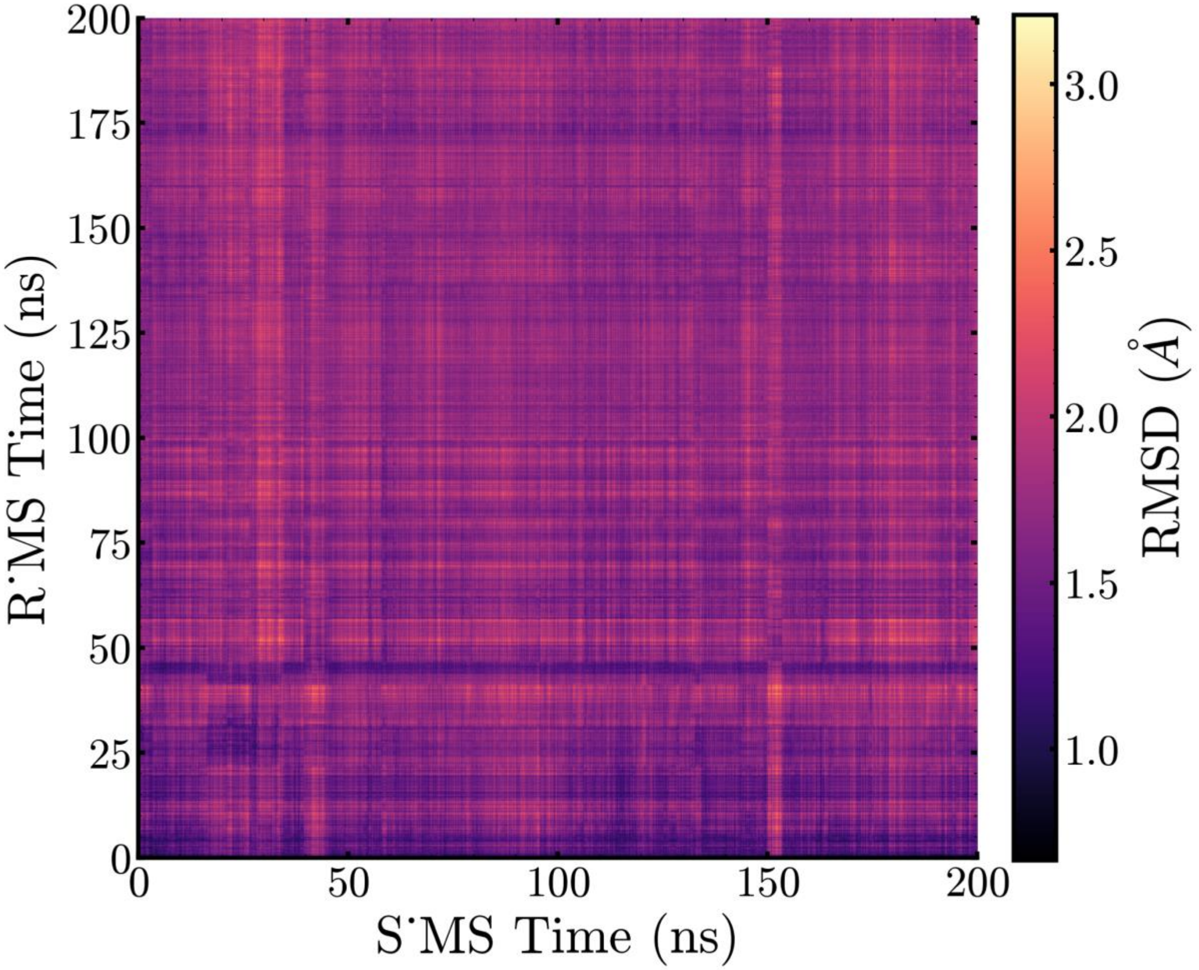

**Figure S26. Pairwise 2D RMSD heat map ($C_\alpha$ only) derived from the MD trajectory of (*R*)-MS-bound MSDO**

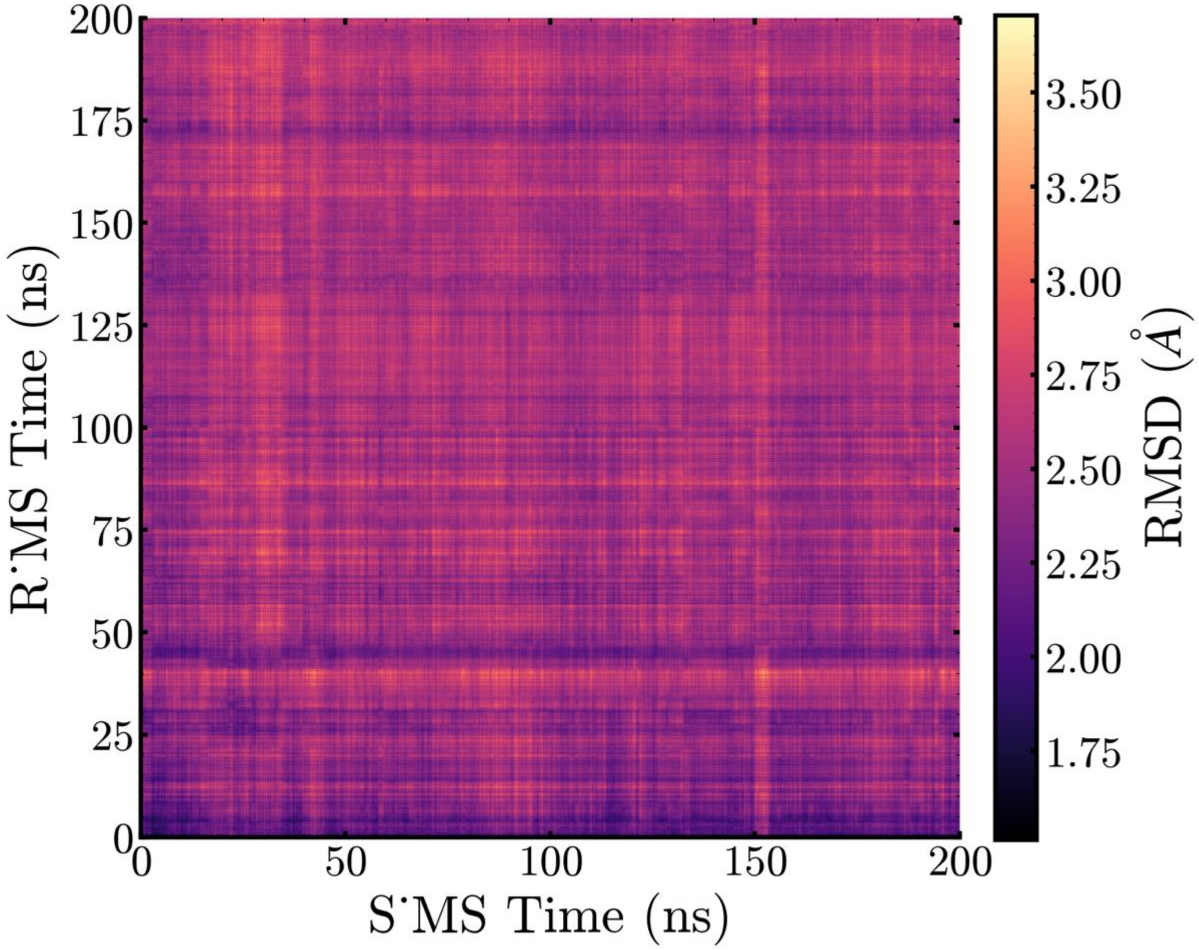

**Figure S27. UB3LYP-GD3BJ/BS1 optimized geometries of the Re$_S$ and TS1$_S$ structures calculated in various spin states**

Bond lengths are in Å, while the imaginary frequency in the transition state is in cm$^{-1}$.

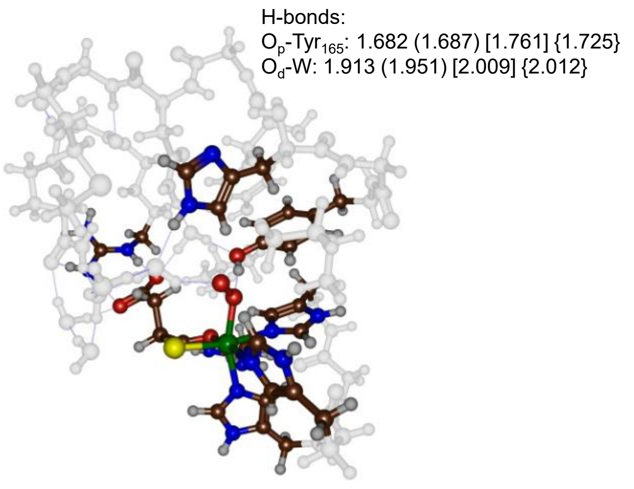

**$^5$Re$_S$ ($^7$Re$_S$) [$^3$Re$_S$] {$^1$Re$_S$}**

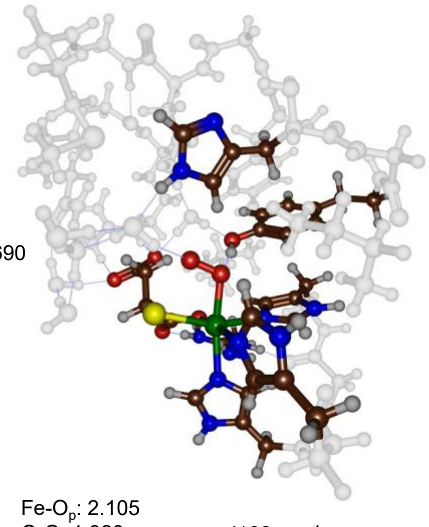

**$^5$TS1$_S$**

**Figure S28. UB3LYP-GD3BJ/BS1 optimized geometries of the IM1S and TS2S structures calculated in various spin states**

Bond lengths are in Å, while the imaginary frequency in the transition state is in cm$^{-1}$.

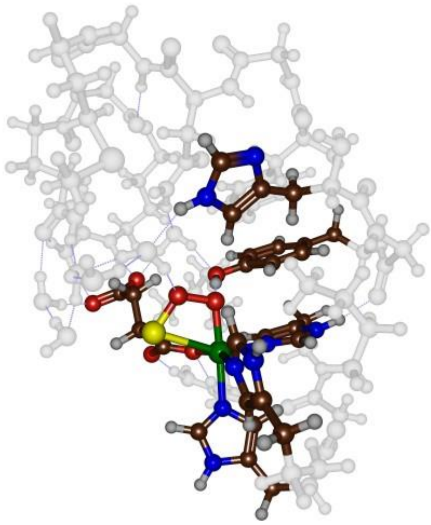

Fe-O$_p$: 2.031 [1.895] {1.955}
O-O: 1.488 [1.489] {1.492}
S-O$_d$: 1.681 [1.705] {1.690}
Fe-S: 2.744 [2.355] {2.605}
Fe-O: 2.049 [2.233] {2.097}
Fe-N$_{ax}$: 2.219 [2.064] {2.033}

**$^5$IM1$_S$ [$^3$IM1$_S$] {$^1$IM1$_S$}**

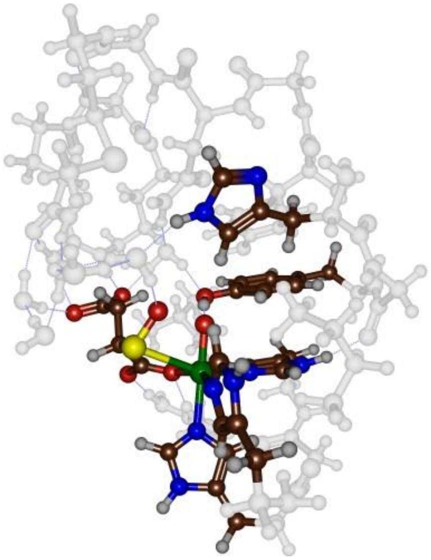

Fe-O$_p$: 1.827
O-O: 1.872
S-O$_d$: 1.592
Fe-S: 2.767
Fe-O: 2.059
Fe-N$_{ax}$: 2.232

i447 cm$^{-1}$

**$^5$TS2$_S$**

**Figure S29. UB3LYP-GD3BJ/BS1 optimized geometries of the IM2S and TS3S structures calculated in various spin states**

Bond lengths are in Å, while the imaginary frequency in the transition state is in $cm^{-1}$.

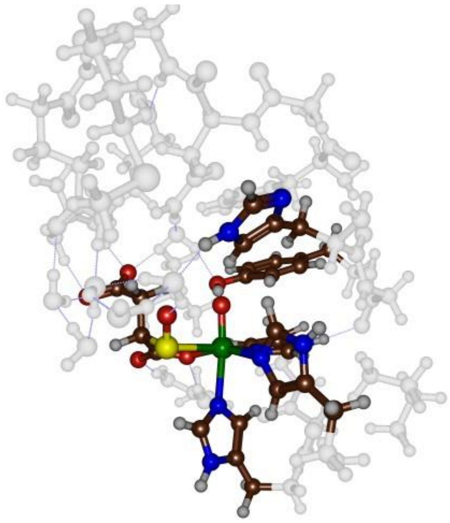

Fe-$O_p$: 1.711 [1.639] {1.640}
O-O: 2.997 [3.028] {2.993}
S-$O_d$: 1.522 [1.525] {1.533}
Fe-S: 2.552 [2.451] {2.369}
Fe-O: 2.090 [1.969] {1.942}
Fe-$N_{ax}$: 2.295 [2.205] {2.326}

**$^5$IM2$_S$ [$^3$IM2$_S$] {$^1$IM2$_S$}**

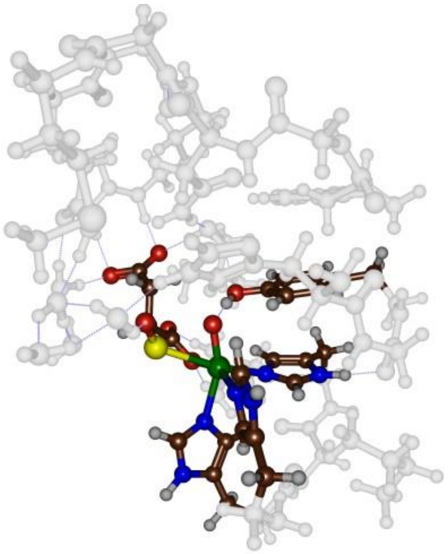

Fe-$O_p$: 1.763
S-$O_p$: 2.339
S-$O_d$: 1.512
Fe-S: 2.611
Fe-O: 2.092
Fe-$N_{ax}$: 2.246

i305 $cm^{-1}$

**$^5$TS3$_S$**

**Figure S30. UB3LYP-GD3BJ/BS1 optimized geometries of the Prods structures calculated in various spin states**

Bond lengths are in Å.

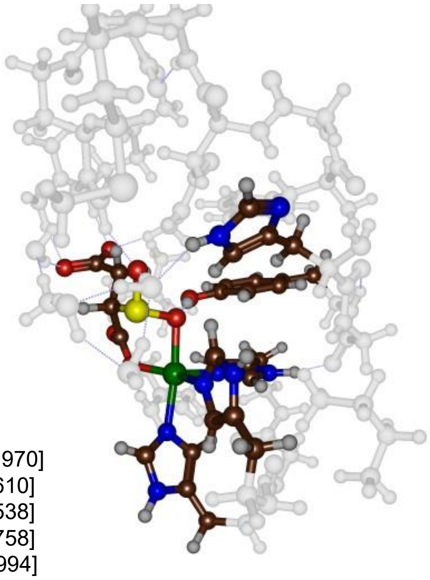

Fe-$O_p$: 2.110 [1.970]
S-$O_p$: 1.598 [1.610]
S-$O_d$: 1.540 [1.538]
Fe-S: 2.881 [2.758]
Fe-O: 2.012 [1.994]
Fe-$N_{ax}$: 2.203 [2.001]

**$^5$Prod$_S$ [$^3$Prod$_S$]**

**Figure S31. UB3LYP-GD3BJ/BS1 optimized geometries of the $Re_R$ and $TS1_R$ structures calculated in various spin states**

Bond lengths are in Å, while the imaginary frequency in the transition state is in $cm^{-1}$.

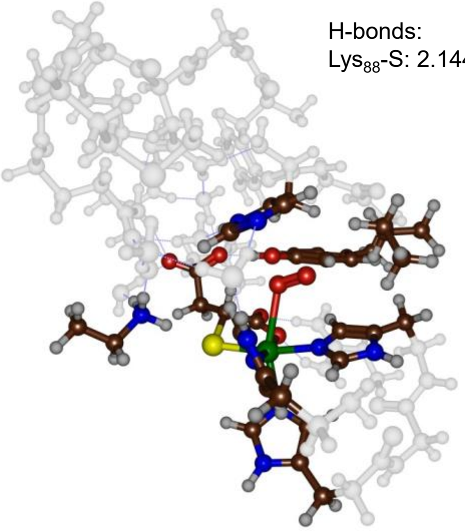

**$^5Re_R$ ($^7Re_R$) [$^3Re_R$] {$^1Re_R$}**

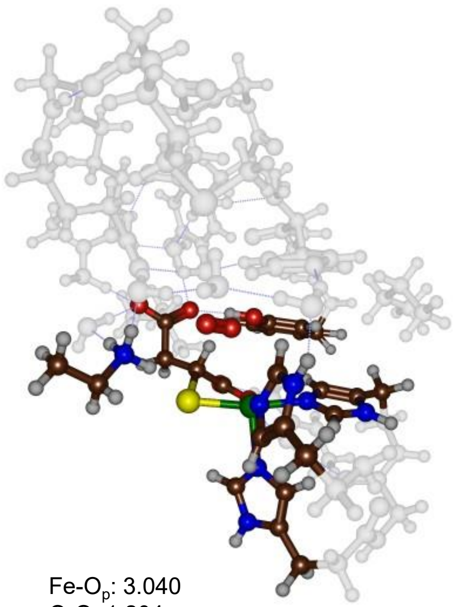

**$^5TS1_R$**

**Figure S32. UB3LYP-GD3BJ/BS1 optimized geometries of the IM1$_R$ and TS2$_R$ structures calculated in various spin states**

Bond lengths are in Å, while the imaginary frequency in the transition state is in cm$^{-1}$.

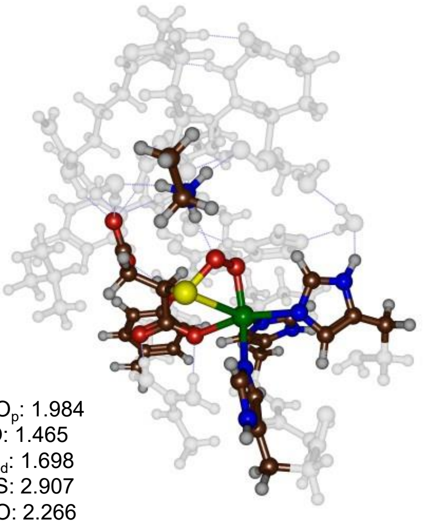

Fe-O$_p$: 1.984
O-O: 1.465
S-O$_d$: 1.698
Fe-S: 2.907
Fe-O: 2.266
Fe-N$_{ax}$: 2.139

**$^5$IM1$_R$**

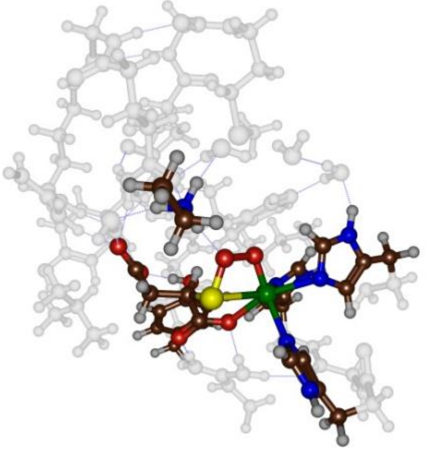

Fe-O$_p$: 1.783
O-O: 1.785
S-O$_d$: 1.612
Fe-S: 2.921
Fe-O: 2.191
Fe-N$_{ax}$: 2.156

i439 cm$^{-1}$

**$^5$TS2$_R$**

**Figure S33. UB3LYP-GD3BJ/BS1 optimized geometries of the IM2R and TS3R structures calculated in various spin states**

Bond lengths are in Å, while the imaginary frequency in the transition state is in $cm^{-1}$.

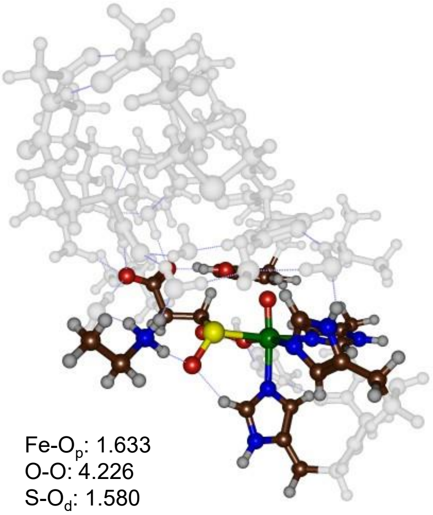

Fe-$O_p$: 1.633
O-O: 4.226
S-$O_d$: 1.580
Fe-S: 2.501
Fe-O: 2.091
Fe-$N_{ax}$: 2.101

**$^5$IM2$_R$**

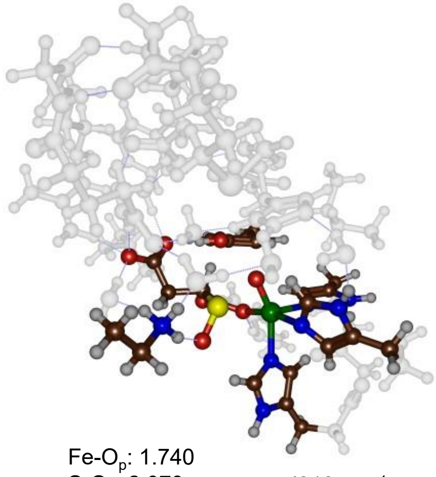

Fe-$O_p$: 1.740
S-$O_p$: 2.079
S-$O_d$: 1.566
Fe-S: 2.592
Fe-O: 2.112
Fe-$N_{ax}$: 2.174

i310 $cm^{-1}$

**$^5$TS3$_R$**

**Figure S34. UB3LYP-GD3BJ/BS1 optimized geometries of the Prod$_R$ structures calculated in various spin states**

Bond lengths are in Å.

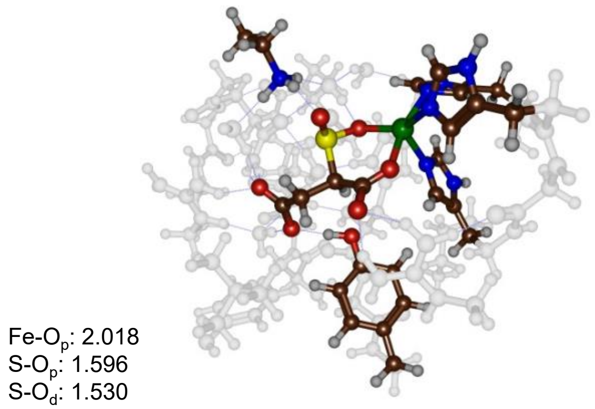

$^5$Prod$_R$

### Cartesian Coordinates

**$^{5}Re_{S}$:**

| | | | |
|---|---|---|---|
| 6 | 14.923650000 | 8.801732000 | 13.438714000 |
| 6 | 13.763248000 | 9.235527000 | 12.555776000 |
| 8 | 12.698929000 | 9.682998000 | 13.004111000 |
| 6 | 14.493956000 | 8.340224000 | 14.831569000 |
| 6 | 13.591242000 | 7.101007000 | 14.798662000 |
| 6 | 13.454288000 | 6.456725000 | 16.170203000 |
| 8 | 13.482608000 | 7.148452000 | 17.209040000 |
| 7 | 13.297057000 | 5.128254000 | 16.179208000 |
| 1 | 15.493502000 | 8.019858000 | 12.923167000 |
| 1 | 15.391538000 | 8.122162000 | 15.420021000 |
| 1 | 13.978795000 | 9.153446000 | 15.350347000 |
| 1 | 13.972110000 | 6.363944000 | 14.082937000 |
| 1 | 12.593178000 | 7.376598000 | 14.437468000 |
| 1 | 13.274570000 | 4.570266000 | 17.036540000 |
| 1 | 13.340492000 | 4.607090000 | 15.306856000 |
| 7 | 13.993706000 | 9.136213000 | 11.222197000 |
| 6 | 12.905322000 | 9.411419000 | 10.309813000 |
| 6 | 11.728766000 | 8.466117000 | 10.595798000 |
| 8 | 11.939973000 | 7.287314000 | 10.898176000 |
| 1 | 14.771693000 | 8.549421000 | 10.909028000 |
| 1 | 12.595784000 | 10.458025000 | 10.396575000 |
| 7 | 10.489540000 | 8.985620000 | 10.477324000 |
| 6 | 9.345799000 | 8.299997000 | 11.087114000 |
| 6 | 9.371858000 | 8.565548000 | 12.595134000 |
| 6 | 8.146617000 | 8.099309000 | 13.374587000 |
| 6 | 8.145465000 | 8.667705000 | 14.799817000 |
| 7 | 9.167193000 | 8.103417000 | 15.677915000 |
| 6 | 10.308965000 | 8.677392000 | 16.057088000 |
| 7 | 10.860138000 | 9.699709000 | 15.346853000 |
| 7 | 10.891722000 | 8.290163000 | 17.195466000 |
| 1 | 10.430677000 | 9.995074000 | 10.427002000 |
| 1 | 9.423143000 | 7.233325000 | 10.867856000 |
| 1 | 10.274574000 | 8.089698000 | 12.994079000 |
| 1 | 9.484432000 | 9.651186000 | 12.738377000 |
| 1 | 7.235082000 | 8.458811000 | 12.876836000 |
| 1 | 8.090260000 | 7.004664000 | 13.405242000 |
| 1 | 8.277669000 | 9.754752000 | 14.773807000 |
| 1 | 7.181733000 | 8.463868000 | 15.278435000 |
| 1 | 8.943039000 | 7.195426000 | 16.132023000 |
| 1 | 11.705342000 | 10.093662000 | 15.739058000 |
| 1 | 10.916568000 | 9.622356000 | 14.336975000 |
| 1 | 10.439757000 | 7.535827000 | 17.740378000 |
| 1 | 11.911339000 | 8.282385000 | 17.249270000 |
| 7 | 5.922211000 | 1.648281000 | 8.271601000 |
| 6 | 6.770551000 | 2.833317000 | 8.119203000 |
| 6 | 8.135502000 | 2.514817000 | 7.505901000 |
| 8 | 8.867053000 | 1.635175000 | 7.975674000 |
| 6 | 7.046151000 | 3.499889000 | 9.481334000 |
| 6 | 5.963579000 | 4.414452000 | 10.010932000 |
| 6 | 4.629054000 | 4.357255000 | 9.587709000 |
| 6 | 6.304119000 | 5.343063000 | 11.005442000 |
| 6 | 3.657769000 | 5.196220000 | 10.135390000 |
| 6 | 5.339659000 | 6.180518000 | 11.562663000 |
| 6 | 4.014040000 | 6.110743000 | 11.127108000 |
| 8 | 3.023840000 | 6.906600000 | 11.654245000 |
| 1 | 6.457030000 | 0.906047000 | 8.730581000 |
| 1 | 6.243204000 | 3.545383000 | 7.471126000 |
| 1 | 7.231918000 | 2.696388000 | 10.205749000 |
| 1 | 7.980841000 | 4.067676000 | 9.437088000 |
| 1 | 4.338915000 | 3.620374000 | 8.845610000 |
| 1 | 7.334415000 | 5.395766000 | 11.345812000 |
| 1 | 2.624331000 | 5.150654000 | 9.806407000 |
| 1 | 5.618987000 | 6.887763000 | 12.340746000 |
| 1 | 3.433689000 | 7.530506000 | 12.273184000 |
| 7 | 8.523865000 | 3.272881000 | 6.449388000 |
| 6 | 9.895798000 | 3.220116000 | 5.948362000 |
| 6 | 10.809541000 | 4.115829000 | 6.794307000 |
| 8 | 11.205200000 | 5.206050000 | 6.393788000 |
| 1 | 7.967479000 | 4.081473000 | 6.206661000 |
| 1 | 10.219094000 | 2.177185000 | 5.978904000 |
| 7 | 11.083595000 | 3.641218000 | 8.036643000 |
| 6 | 11.726328000 | 4.516996000 | 8.995283000 |
| 6 | 13.172029000 | 4.830869000 | 8.579088000 |
| 8 | 13.951371000 | 4.012710000 | 8.090033000 |
| 6 | 11.694752000 | 3.921816000 | 10.406846000 |
| 6 | 10.489049000 | 4.407464000 | 11.216375000 |
| 8 | 9.576250000 | 5.061167000 | 10.699724000 |
| 7 | 10.522130000 | 4.098506000 | 12.527619000 |
| 1 | 10.672112000 | 2.763421000 | 8.335760000 |
| 1 | 11.174017000 | 5.459995000 | 9.014036000 |
| 1 | 12.598150000 | 4.208512000 | 10.956232000 |
| 1 | 11.695284000 | 2.825994000 | 10.371182000 |
| 1 | 11.186820000 | 3.413232000 | 12.861559000 |
| 1 | 9.696054000 | 4.295453000 | 13.111482000 |
| 7 | 13.513109000 | 6.119070000 | 8.852044000 |
| 6 | 14.841302000 | 6.622287000 | 8.575677000 |
| 6 | 15.788129000 | 6.293304000 | 9.726805000 |
| 8 | 16.044042000 | 7.108792000 | 10.625300000 |
| 1 | 12.931898000 | 6.624873000 | 9.521000000 |
| 1 | 15.187974000 | 6.171553000 | 7.642311000 |
| 7 | 16.258544000 | 5.027857000 | 9.687165000 |
| 6 | 17.084661000 | 4.447837000 | 10.732599000 |
| 6 | 16.297187000 | 3.531894000 | 11.678829000 |
| 6 | 15.263071000 | 4.289465000 | 12.510262000 |
| 16 | 14.311864000 | 3.199046000 | 13.637570000 |
| 6 | 15.563484000 | 2.885448000 | 14.929561000 |
| 1 | 15.827712000 | 4.416336000 | 8.992984000 |
| 1 | 17.533833000 | 5.278995000 | 11.282058000 |
| 1 | 15.791235000 | 2.750286000 | 11.099389000 |
| 1 | 17.012715000 | 3.026871000 | 12.339110000 |
| 1 | 15.734084000 | 5.083551000 | 13.100154000 |
| 1 | 14.520479000 | 4.767965000 | 11.865002000 |
| 1 | 15.107800000 | 2.220708000 | 15.667360000 |
| 1 | 15.847240000 | 3.817682000 | 15.426598000 |
| 1 | 16.449121000 | 2.401099000 | 14.510348000 |
| 6 | 3.160868000 | -1.880374000 | 21.834499000 |
| 6 | 2.684023000 | -1.783615000 | 20.389729000 |
| 8 | 3.335976000 | -2.187180000 | 19.425216000 |
| 6 | 3.694584000 | -0.552856000 | 22.424834000 |
| 6 | 4.984942000 | -0.100604000 | 21.814370000 |
| 7 | 5.902031000 | 0.722093000 | 22.456778000 |
| 6 | 5.535068000 | -0.338124000 | 20.583062000 |
| 6 | 6.948902000 | 0.942646000 | 21.617825000 |
| 7 | 6.752563000 | 0.307409000 | 20.475755000 |
| 1 | 2.320833000 | -2.210505000 | 22.456638000 |
| 1 | 2.935600000 | 0.235931000 | 22.318437000 |
| 1 | 3.832749000 | -0.700500000 | 23.502890000 |
| 1 | 5.129792000 | -0.922385000 | 19.778466000 |
| 1 | 7.807328000 | 1.550557000 | 21.861716000 |
| 1 | 5.808771000 | 1.103926000 | 23.387945000 |
| 7 | 1.444030000 | -1.236294000 | 20.218832000 |
| 6 | 0.936226000 | -0.976697000 | 18.896305000 |
| 6 | 1.545786000 | 0.285993000 | 18.264097000 |
| 8 | 2.083278000 | 1.149072000 | 18.967890000 |
| 1 | 0.993724000 | -0.787489000 | 21.003280000 |
| 1 | 1.150796000 | -1.845277000 | 18.267767000 |
| 7 | 1.392625000 | 0.378188000 | 16.928578000 |
| 6 | 1.981697000 | 1.421642000 | 16.090541000 |
| 6 | 3.181540000 | 0.893099000 | 15.284103000 |
| 6 | 4.245023000 | 0.325288000 | 16.163541000 |
| 7 | 4.417804000 | -1.032051000 | 16.390320000 |
| 6 | 5.210963000 | 0.933275000 | 16.922408000 |
| 6 | 5.470478000 | -1.203120000 | 17.227566000 |
| 7 | 5.963483000 | -0.024514000 | 17.573486000 |
| 1 | 0.961348000 | -0.404569000 | 16.452857000 |

| | | | |
|---|---|---|---|
| 1 | 2.300923000 | 2.226348000 | 16.753655000 |
| 1 | 2.852086000 | 0.140765000 | 14.558025000 |
| 1 | 3.600663000 | 1.712998000 | 14.692387000 |
| 1 | 5.449658000 | 1.977212000 | 17.011425000 |
| 1 | 5.829743000 | -2.164308000 | 17.557596000 |
| 1 | 3.904727000 | -1.777024000 | 15.941191000 |
| 6 | 9.146931000 | -3.841036000 | 13.481665000 |
| 6 | 7.678261000 | -3.403030000 | 13.567522000 |
| 6 | 10.133858000 | -2.835552000 | 14.081908000 |
| 6 | 7.150527000 | -3.280198000 | 14.998863000 |
| 1 | 9.262964000 | -4.820482000 | 13.968885000 |
| 1 | 7.038457000 | -4.106499000 | 13.022908000 |
| 1 | 7.557373000 | -2.436786000 | 13.063160000 |
| 1 | 9.965407000 | -2.701055000 | 15.156106000 |
| 1 | 10.032188000 | -1.848996000 | 13.617051000 |
| 1 | 11.169283000 | -3.166811000 | 13.947409000 |
| 1 | 6.074151000 | -3.076810000 | 14.986014000 |
| 1 | 7.636937000 | -2.467072000 | 15.547832000 |
| 1 | 7.305621000 | -4.210661000 | 15.562346000 |
| 7 | 5.276874000 | -4.446242000 | 20.696502000 |
| 6 | 6.600072000 | -4.949077000 | 21.060179000 |
| 6 | 7.633883000 | -5.121308000 | 19.923540000 |
| 6 | 8.058208000 | -3.807044000 | 19.347055000 |
| 7 | 8.907220000 | -3.663459000 | 18.255260000 |
| 6 | 7.745271000 | -2.522841000 | 19.709510000 |
| 6 | 9.068736000 | -2.343592000 | 17.992797000 |
| 7 | 8.363368000 | -1.625071000 | 18.854469000 |
| 1 | 5.318474000 | -3.602794000 | 20.129641000 |
| 1 | 6.462648000 | -5.918310000 | 21.549277000 |
| 1 | 7.209110000 | -5.759289000 | 19.134919000 |
| 1 | 8.509928000 | -5.656021000 | 20.317778000 |
| 1 | 7.130266000 | -2.184596000 | 20.524141000 |
| 1 | 9.668782000 | -1.931665000 | 17.194775000 |
| 1 | 9.334667000 | -4.415498000 | 17.732087000 |
| 7 | 9.910820000 | -1.046504000 | 9.061796000 |
| 6 | 9.041190000 | -1.124822000 | 10.235250000 |
| 6 | 7.710960000 | -1.722298000 | 9.798902000 |
| 8 | 6.902950000 | -1.124511000 | 9.076197000 |
| 6 | 8.813826000 | 0.218698000 | 10.979789000 |
| 6 | 10.017464000 | 0.570490000 | 11.793686000 |
| 7 | 11.294941000 | 0.603122000 | 11.248495000 |
| 6 | 10.093890000 | 0.793518000 | 13.148434000 |
| 6 | 12.113419000 | 0.836874000 | 12.260855000 |
| 7 | 11.432812000 | 0.975683000 | 13.434898000 |
| 1 | 9.506578000 | -0.355255000 | 8.429025000 |
| 1 | 9.513030000 | -1.819357000 | 10.942943000 |
| 1 | 8.601270000 | 0.977165000 | 10.221698000 |
| 1 | 7.942939000 | 0.155456000 | 11.641127000 |
| 1 | 9.345822000 | 0.851288000 | 13.916567000 |
| 1 | 13.186806000 | 0.933633000 | 12.198255000 |
| 1 | 11.817478000 | 1.097749000 | 14.376992000 |
| 7 | 7.463975000 | -2.972784000 | 10.263282000 |
| 6 | 6.252779000 | -3.712419000 | 9.946358000 |
| 6 | 5.042157000 | -3.184126000 | 10.731790000 |
| 8 | 4.687739000 | -3.683223000 | 11.802356000 |
| 1 | 8.150630000 | -3.407613000 | 10.859202000 |
| 1 | 6.069503000 | -3.648628000 | 8.868973000 |
| 7 | 4.440357000 | -2.129193000 | 10.133275000 |
| 6 | 3.318069000 | -1.412610000 | 10.712764000 |
| 6 | 3.597637000 | 0.081097000 | 10.874096000 |
| 6 | 4.635403000 | 0.475119000 | 11.905798000 |
| 6 | 5.258287000 | -0.414830000 | 12.787896000 |
| 6 | 4.983003000 | 1.831912000 | 11.983892000 |
| 6 | 6.187253000 | 0.043522000 | 13.730451000 |
| 6 | 5.900707000 | 2.302193000 | 12.913982000 |
| 6 | 6.496172000 | 1.401819000 | 13.802795000 |
| 8 | 7.361749000 | 1.912093000 | 14.727978000 |
| 1 | 4.982686000 | -1.672799000 | 9.401436000 |
| 1 | 3.092144000 | -1.892657000 | 11.668457000 |
| 1 | 2.644280000 | 0.572373000 | 11.118115000 |

| | | | |
|---|---|---|---|
| 1 | 3.902881000 | 0.502190000 | 9.905584000 |
| 1 | 5.038808000 | -1.477882000 | 12.751399000 |
| 1 | 4.532596000 | 2.537134000 | 11.294749000 |
| 1 | 6.665202000 | -0.649120000 | 14.414675000 |
| 1 | 6.170624000 | 3.352212000 | 12.944941000 |
| 1 | 7.676387000 | 1.226781000 | 15.373059000 |
| 6 | 4.724073000 | 6.810813000 | 15.959413000 |
| 6 | 4.977063000 | 5.485431000 | 15.232972000 |
| 6 | 5.547009000 | 4.440722000 | 16.193497000 |
| 6 | 3.701056000 | 4.964745000 | 14.561088000 |
| 1 | 3.971290000 | 6.686299000 | 16.752897000 |
| 1 | 5.641321000 | 7.182481000 | 16.431312000 |
| 1 | 5.734056000 | 5.656055000 | 14.457542000 |
| 1 | 5.854021000 | 3.544102000 | 15.652152000 |
| 1 | 4.786041000 | 4.155022000 | 16.931811000 |
| 1 | 6.426650000 | 4.812719000 | 16.726674000 |
| 1 | 3.906845000 | 4.056114000 | 13.985019000 |
| 1 | 2.939815000 | 4.724627000 | 15.319780000 |
| 1 | 3.263379000 | 5.692952000 | 13.871075000 |
| 6 | 2.034241000 | 5.215727000 | 19.079809000 |
| 7 | 2.731081000 | 3.943903000 | 19.168395000 |
| 6 | 4.041136000 | 3.823621000 | 19.438118000 |
| 7 | 4.821975000 | 4.897257000 | 19.580225000 |
| 7 | 4.562060000 | 2.600708000 | 19.632460000 |
| 1 | 2.254565000 | 5.841152000 | 19.951737000 |
| 1 | 2.298260000 | 5.765211000 | 18.167064000 |
| 1 | 2.229214000 | 3.071107000 | 19.023645000 |
| 1 | 4.528029000 | 5.777762000 | 19.184132000 |
| 1 | 5.842717000 | 4.733860000 | 19.687482000 |
| 1 | 3.953237000 | 1.810114000 | 19.450367000 |
| 1 | 5.553815000 | 2.473483000 | 19.390421000 |
| 26 | 7.883823000 | 0.467507000 | 18.567255000 |
| 6 | 9.458032000 | 3.128160000 | 19.042788000 |
| 6 | 9.858587000 | 3.673487000 | 17.655420000 |
| 6 | 9.536528000 | 5.173972000 | 17.552532000 |
| 6 | 7.951385000 | 3.287676000 | 19.199653000 |
| 8 | 10.059744000 | 5.945579000 | 18.396471000 |
| 8 | 8.762213000 | 5.553749000 | 16.623012000 |
| 8 | 7.490588000 | 4.254737000 | 19.814384000 |
| 8 | 7.211042000 | 2.382444000 | 18.617416000 |
| 16 | 9.923592000 | 1.355917000 | 19.251568000 |
| 1 | 9.981666000 | 3.697609000 | 19.810322000 |
| 1 | 10.937398000 | 3.552836000 | 17.530599000 |
| 1 | 9.347800000 | 3.119291000 | 16.864767000 |
| 8 | 8.404740000 | 0.290333000 | 16.564721000 |
| 8 | 14.561683000 | 0.521598000 | 17.415556000 |
| 1 | 13.728790000 | 0.413817000 | 16.898242000 |
| 1 | 15.272637000 | 0.180001000 | 16.854562000 |
| 8 | 12.161501000 | 1.060459000 | 16.235747000 |
| 1 | 11.342590000 | 0.649028000 | 16.573984000 |
| 1 | 12.330856000 | 1.841692000 | 16.825504000 |
| 8 | 12.430172000 | 3.552826000 | 20.323485000 |
| 1 | 12.560631000 | 4.533454000 | 20.132098000 |
| 1 | 13.021450000 | 3.340993000 | 21.060668000 |
| 8 | 12.487097000 | 6.069056000 | 19.619035000 |
| 1 | 11.568380000 | 6.016180000 | 19.267898000 |
| 1 | 13.011504000 | 6.388832000 | 18.853133000 |
| 1 | 7.027059000 | -4.281745000 | 21.818952000 |
| 1 | 4.768845000 | -5.134309000 | 20.144771000 |
| 1 | -0.150778000 | -0.853790000 | 18.948987000 |
| 1 | 1.214109000 | 1.818598000 | 15.418428000 |
| 1 | 0.961071000 | 5.020082000 | 19.060851000 |
| 1 | 4.348675000 | 7.586605000 | 15.280222000 |
| 1 | 17.892607000 | 3.878232000 | 10.260459000 |
| 1 | 14.805894000 | 7.706864000 | 8.452015000 |
| 1 | 9.918380000 | 3.584214000 | 4.920948000 |
| 1 | 5.698815000 | 1.275161000 | 7.348920000 |
| 1 | 10.791062000 | -0.640158000 | 9.384503000 |
| 1 | 3.927560000 | -2.660031000 | 21.868545000 |
| 1 | 8.430449000 | 8.686405000 | 10.626967000 |

| | | | |
|---|---|---|---|
| 1 | 13.257002000 | 9.249134000 | 9.286290000 |
| 1 | 15.599583000 | 9.662454000 | 13.532818000 |
| 1 | 2.438072000 | -1.538945000 | 10.068726000 |
| 1 | 6.401883000 | -4.756384000 | 10.223449000 |
| 1 | 9.418744000 | -4.007661000 | 12.428623000 |
| 8 | 8.241723000 | 4.542619000 | 14.162345000 |
| 1 | 8.458822000 | 4.882903000 | 15.062995000 |
| 1 | 7.962377000 | 3.621921000 | 14.324907000 |
| 8 | 13.241275000 | 2.882375000 | 17.898332000 |
| 1 | 12.934442000 | 2.998351000 | 18.844756000 |
| 1 | 13.984220000 | 2.241041000 | 17.915317000 |
| 8 | 9.610672000 | -0.045250000 | 16.150591000 |

**$^{5}$TS1$_{S}$:**

| | | | |
|---|---|---|---|
| 6 | 14.850930000 | 9.030892000 | 13.389022000 |
| 6 | 13.673244000 | 9.406425000 | 12.503132000 |
| 8 | 12.597657000 | 9.827599000 | 12.950130000 |
| 6 | 14.428823000 | 8.494996000 | 14.757934000 |
| 6 | 13.582544000 | 7.219203000 | 14.665373000 |
| 6 | 13.447048000 | 6.529553000 | 16.014440000 |
| 8 | 13.423913000 | 7.191996000 | 17.072494000 |
| 7 | 13.355595000 | 5.195215000 | 15.988023000 |
| 1 | 15.482908000 | 8.306466000 | 12.863108000 |
| 1 | 15.329110000 | 8.294697000 | 15.348574000 |
| 1 | 13.871946000 | 9.262274000 | 15.302537000 |
| 1 | 14.012316000 | 6.522989000 | 13.936310000 |
| 1 | 12.581159000 | 7.460703000 | 14.288855000 |
| 1 | 13.348320000 | 4.619775000 | 16.835814000 |
| 1 | 13.434584000 | 4.693084000 | 15.106954000 |
| 7 | 13.893907000 | 9.277523000 | 11.170859000 |
| 6 | 12.784186000 | 9.477524000 | 10.265037000 |
| 6 | 11.647997000 | 8.499141000 | 10.603636000 |
| 8 | 11.908197000 | 7.345171000 | 10.958615000 |
| 1 | 14.697487000 | 8.721496000 | 10.866168000 |
| 1 | 12.434573000 | 10.513795000 | 10.318035000 |
| 7 | 10.388774000 | 8.963455000 | 10.467762000 |
| 6 | 9.271388000 | 8.267311000 | 11.114224000 |
| 6 | 9.294610000 | 8.601075000 | 12.609330000 |
| 6 | 8.069279000 | 8.172201000 | 13.410113000 |
| 6 | 8.077884000 | 8.790270000 | 14.815338000 |
| 7 | 9.085419000 | 8.231220000 | 15.711278000 |
| 6 | 10.253415000 | 8.773192000 | 16.053285000 |
| 7 | 10.815986000 | 9.773456000 | 15.322993000 |
| 7 | 10.856237000 | 8.377397000 | 17.179629000 |
| 1 | 10.290840000 | 9.966051000 | 10.366761000 |
| 1 | 9.383665000 | 7.195118000 | 10.941549000 |
| 1 | 10.197689000 | 8.145079000 | 13.029790000 |
| 1 | 9.406188000 | 9.692118000 | 12.702903000 |
| 1 | 7.158769000 | 8.520252000 | 12.902701000 |
| 1 | 8.006571000 | 7.079589000 | 13.479983000 |
| 1 | 8.233733000 | 9.872828000 | 14.753102000 |
| 1 | 7.109509000 | 8.624187000 | 15.298725000 |
| 1 | 8.844035000 | 7.337206000 | 16.183421000 |
| 1 | 11.683183000 | 10.145444000 | 15.686808000 |
| 1 | 10.831923000 | 9.705407000 | 14.311530000 |
| 1 | 10.397747000 | 7.642495000 | 17.742934000 |
| 1 | 11.876601000 | 8.331143000 | 17.195635000 |
| 7 | 6.063891000 | 1.601775000 | 8.327348000 |
| 6 | 6.944181000 | 2.760382000 | 8.154094000 |
| 6 | 8.322964000 | 2.382944000 | 7.609986000 |
| 8 | 9.016258000 | 1.515187000 | 8.152750000 |
| 6 | 7.188109000 | 3.478224000 | 9.494672000 |
| 6 | 6.093478000 | 4.408013000 | 9.970169000 |
| 6 | 4.781371000 | 4.374889000 | 9.481102000 |
| 6 | 6.398978000 | 5.325815000 | 10.986248000 |
| 6 | 3.797819000 | 5.227456000 | 9.985034000 |
| 6 | 5.421980000 | 6.175456000 | 11.500692000 |
| 6 | 4.118474000 | 6.130313000 | 10.999127000 |
| 8 | 3.116687000 | 6.939606000 | 11.480793000 |
| 1 | 6.565599000 | 0.874229000 | 8.842983000 |
| 1 | 6.458500000 | 3.454575000 | 7.455798000 |
| 1 | 7.362518000 | 2.703564000 | 10.252964000 |
| 1 | 8.123168000 | 4.045811000 | 9.448714000 |
| 1 | 4.513628000 | 3.646017000 | 8.723257000 |
| 1 | 7.411205000 | 5.358168000 | 11.379454000 |
| 1 | 2.781358000 | 5.200254000 | 9.605125000 |
| 1 | 5.673308000 | 6.870202000 | 12.298995000 |
| 1 | 3.504829000 | 7.554636000 | 12.122097000 |
| 7 | 8.768544000 | 3.076999000 | 6.531551000 |
| 6 | 10.167043000 | 2.992063000 | 6.116093000 |
| 6 | 11.021140000 | 3.945370000 | 6.959305000 |
| 8 | 11.401309000 | 5.029147000 | 6.526448000 |
| 1 | 8.242479000 | 3.886952000 | 6.231449000 |
| 1 | 10.483739000 | 1.953793000 | 6.234227000 |
| 7 | 11.254538000 | 3.539598000 | 8.234442000 |
| 6 | 11.840879000 | 4.484819000 | 9.163922000 |
| 6 | 13.271595000 | 4.854896000 | 8.738791000 |
| 8 | 14.089333000 | 4.061094000 | 8.273115000 |
| 6 | 11.821088000 | 3.953067000 | 10.600119000 |
| 6 | 10.584961000 | 4.416732000 | 11.377733000 |
| 8 | 9.660619000 | 5.026120000 | 10.830331000 |
| 7 | 10.611345000 | 4.147771000 | 12.698917000 |
| 1 | 10.856760000 | 2.667193000 | 8.565713000 |
| 1 | 11.242594000 | 5.399416000 | 9.138663000 |
| 1 | 12.699472000 | 4.314907000 | 11.144382000 |
| 1 | 11.877288000 | 2.858707000 | 10.613998000 |
| 1 | 11.286246000 | 3.484096000 | 13.055561000 |
| 1 | 9.770676000 | 4.339068000 | 13.264386000 |
| 7 | 13.549301000 | 6.167053000 | 8.964207000 |
| 6 | 14.834670000 | 6.734142000 | 8.618124000 |
| 6 | 15.841406000 | 6.506417000 | 9.741737000 |
| 8 | 16.057593000 | 7.359968000 | 10.614656000 |
| 1 | 12.946144000 | 6.674936000 | 9.611623000 |
| 1 | 15.171909000 | 6.267819000 | 7.688929000 |
| 7 | 16.410127000 | 5.281564000 | 9.711053000 |
| 6 | 17.285308000 | 4.782076000 | 10.758380000 |
| 6 | 16.579937000 | 3.783426000 | 11.685839000 |
| 6 | 15.437055000 | 4.426474000 | 12.469853000 |
| 16 | 14.509164000 | 3.239264000 | 13.513622000 |
| 6 | 15.719839000 | 2.973543000 | 14.854177000 |
| 1 | 15.995135000 | 4.621390000 | 9.052910000 |
| 1 | 17.639553000 | 5.651828000 | 11.317328000 |
| 1 | 16.185128000 | 2.947857000 | 11.095508000 |
| 1 | 17.324952000 | 3.365464000 | 12.373662000 |
| 1 | 15.801554000 | 5.243654000 | 13.102215000 |
| 1 | 14.698831000 | 4.854410000 | 11.785803000 |
| 1 | 15.269333000 | 2.283633000 | 15.570895000 |
| 1 | 15.938635000 | 3.914534000 | 15.367516000 |
| 1 | 16.643462000 | 2.535519000 | 14.467697000 |
| 6 | 3.264100000 | -1.967110000 | 21.758641000 |
| 6 | 2.764492000 | -1.840786000 | 20.324072000 |
| 8 | 3.396327000 | -2.231291000 | 19.341594000 |
| 6 | 3.795018000 | -0.649637000 | 22.370723000 |
| 6 | 5.078442000 | -0.181770000 | 21.758644000 |
| 7 | 5.943910000 | 0.716096000 | 22.370189000 |
| 6 | 5.670039000 | -0.470193000 | 20.557487000 |
| 6 | 6.999810000 | 0.931477000 | 21.540349000 |
| 7 | 6.862111000 | 0.220816000 | 20.434996000 |
| 1 | 2.435693000 | -2.319130000 | 22.384311000 |
| 1 | 3.031737000 | 0.137851000 | 22.285339000 |
| 1 | 3.941514000 | -0.817300000 | 23.445091000 |
| 1 | 5.303901000 | -1.112122000 | 19.777293000 |
| 1 | 7.813050000 | 1.605899000 | 21.763671000 |
| 1 | 5.808916000 | 1.153234000 | 23.271230000 |
| 7 | 1.522672000 | -1.285896000 | 20.186765000 |
| 6 | 0.988522000 | -1.002611000 | 18.880277000 |
| 6 | 1.572470000 | 0.279119000 | 18.262882000 |
| 8 | 2.133861000 | 1.124567000 | 18.968726000 |
| 1 | 1.093180000 | -0.844569000 | 20.986958000 |
| 1 | 1.204068000 | -1.854661000 | 18.229809000 |

| | | | |
|---|---|---|---|
| 7 | 1.371240000 | 0.406118000 | 16.935971000 |
| 6 | 1.957957000 | 1.453563000 | 16.101721000 |
| 6 | 3.130846000 | 0.914988000 | 15.262969000 |
| 6 | 4.199564000 | 0.311332000 | 16.112133000 |
| 7 | 4.361838000 | -1.052796000 | 16.298390000 |
| 6 | 5.193971000 | 0.888999000 | 16.857690000 |
| 6 | 5.436431000 | -1.256950000 | 17.101662000 |
| 7 | 5.955907000 | -0.093314000 | 17.459697000 |
| 1 | 0.928571000 | -0.367614000 | 16.455914000 |
| 1 | 2.305195000 | 2.242412000 | 16.769771000 |
| 1 | 2.770086000 | 0.179260000 | 14.534466000 |
| 1 | 3.555788000 | 1.733855000 | 14.673499000 |
| 1 | 5.450566000 | 1.927734000 | 16.962354000 |
| 1 | 5.783341000 | -2.230389000 | 17.407258000 |
| 1 | 3.834326000 | -1.782332000 | 15.840281000 |
| 6 | 8.427156000 | -4.113660000 | 14.023901000 |
| 6 | 7.000582000 | -4.685174000 | 14.002818000 |
| 6 | 8.490640000 | -2.584634000 | 13.973175000 |
| 6 | 6.096733000 | -4.179336000 | 15.130269000 |
| 1 | 8.959066000 | -4.476614000 | 14.919107000 |
| 1 | 7.054229000 | -5.781053000 | 14.041705000 |
| 1 | 6.509646000 | -4.428005000 | 13.058235000 |
| 1 | 8.082380000 | -2.132581000 | 14.881601000 |
| 1 | 7.903828000 | -2.194880000 | 13.136221000 |
| 1 | 9.515360000 | -2.215787000 | 13.856349000 |
| 1 | 5.107842000 | -4.645201000 | 15.064044000 |
| 1 | 5.959155000 | -3.095844000 | 15.056295000 |
| 1 | 6.512040000 | -4.399333000 | 16.123665000 |
| 7 | 5.324205000 | -4.520870000 | 20.516182000 |
| 6 | 6.643723000 | -5.079437000 | 20.801650000 |
| 6 | 7.614457000 | -5.239234000 | 19.609052000 |
| 6 | 8.018115000 | -3.915947000 | 19.037914000 |
| 7 | 8.668649000 | -3.754112000 | 17.820569000 |
| 6 | 7.844267000 | -2.637822000 | 19.505852000 |
| 6 | 8.849412000 | -2.428895000 | 17.591278000 |
| 7 | 8.348846000 | -1.725659000 | 18.595286000 |
| 1 | 5.377600000 | -3.660062000 | 19.976743000 |
| 1 | 6.498650000 | -6.061473000 | 21.262309000 |
| 1 | 7.139183000 | -5.852516000 | 18.829913000 |
| 1 | 8.503283000 | -5.793766000 | 19.942830000 |
| 1 | 7.392159000 | -2.310631000 | 20.426054000 |
| 1 | 9.307404000 | -2.013080000 | 16.709138000 |
| 1 | 8.949446000 | -4.495305000 | 17.192901000 |
| 7 | 9.941271000 | -1.185426000 | 9.264020000 |
| 6 | 9.084842000 | -1.198303000 | 10.450185000 |
| 6 | 7.728565000 | -1.749155000 | 10.035729000 |
| 8 | 6.957238000 | -1.144736000 | 9.281145000 |
| 6 | 8.915560000 | 0.172201000 | 11.161709000 |
| 6 | 10.154514000 | 0.522098000 | 11.922101000 |
| 7 | 11.407505000 | 0.530058000 | 11.322123000 |
| 6 | 10.289640000 | 0.782890000 | 13.266439000 |
| 6 | 12.269860000 | 0.784191000 | 12.292253000 |
| 7 | 11.642075000 | 0.964754000 | 13.488496000 |
| 1 | 9.552030000 | -0.497925000 | 8.617826000 |
| 1 | 9.534624000 | -1.894190000 | 11.170429000 |
| 1 | 8.696975000 | 0.913421000 | 10.388284000 |
| 1 | 8.065183000 | 0.149908000 | 11.851379000 |
| 1 | 9.568326000 | 0.904259000 | 14.054659000 |
| 1 | 13.340901000 | 0.867909000 | 12.182024000 |
| 1 | 12.079698000 | 1.086352000 | 14.408628000 |
| 7 | 7.405439000 | -2.964642000 | 10.553114000 |
| 6 | 6.187319000 | -3.669957000 | 10.179459000 |
| 6 | 4.955953000 | -3.082226000 | 10.886454000 |
| 8 | 4.534535000 | -3.543200000 | 11.951327000 |
| 1 | 8.094170000 | -3.449800000 | 11.107374000 |
| 1 | 6.065998000 | -3.622315000 | 9.092174000 |
| 7 | 4.419176000 | -2.024598000 | 10.237999000 |
| 6 | 3.314104000 | -1.240235000 | 10.764060000 |
| 6 | 3.662462000 | 0.242039000 | 10.898075000 |
| 6 | 4.692268000 | 0.609485000 | 11.947699000 |
| 6 | 5.277283000 | -0.295997000 | 12.839237000 |
| 6 | 5.077446000 | 1.955893000 | 12.032352000 |
| 6 | 6.208397000 | 0.135650000 | 13.790597000 |
| 6 | 6.001864000 | 2.398173000 | 12.969935000 |
| 6 | 6.566640000 | 1.481324000 | 13.863281000 |
| 8 | 7.439070000 | 1.964275000 | 14.795208000 |
| 1 | 5.012933000 | -1.605080000 | 9.524270000 |
| 1 | 3.036491000 | -1.684216000 | 11.723669000 |
| 1 | 2.728255000 | 0.783945000 | 11.105198000 |
| 1 | 4.012161000 | 0.624439000 | 9.928304000 |
| 1 | 5.019818000 | -1.350358000 | 12.812076000 |
| 1 | 4.653302000 | 2.675134000 | 11.340926000 |
| 1 | 6.647501000 | -0.569959000 | 14.486020000 |
| 1 | 6.300087000 | 3.440088000 | 13.002368000 |
| 1 | 7.783627000 | 1.262958000 | 15.410182000 |
| 6 | 4.770494000 | 6.833300000 | 15.940301000 |
| 6 | 4.983772000 | 5.507972000 | 15.201198000 |
| 6 | 5.467985000 | 4.421978000 | 16.163204000 |
| 6 | 3.711718000 | 5.055942000 | 14.475300000 |
| 1 | 3.965177000 | 6.744225000 | 16.685542000 |
| 1 | 5.682248000 | 7.137775000 | 16.467812000 |
| 1 | 5.777250000 | 5.654471000 | 14.457838000 |
| 1 | 5.743519000 | 3.519023000 | 15.615827000 |
| 1 | 4.670859000 | 4.163962000 | 16.874212000 |
| 1 | 6.348600000 | 4.742593000 | 16.727042000 |
| 1 | 3.896675000 | 4.142087000 | 13.900367000 |
| 1 | 2.909531000 | 4.849102000 | 15.200217000 |
| 1 | 3.336579000 | 5.808143000 | 13.774183000 |
| 6 | 2.163680000 | 5.187736000 | 19.121685000 |
| 7 | 2.838667000 | 3.903002000 | 19.184643000 |
| 6 | 4.149174000 | 3.758058000 | 19.441569000 |
| 7 | 4.947655000 | 4.817241000 | 19.591611000 |
| 7 | 4.653076000 | 2.525005000 | 19.615118000 |
| 1 | 2.386619000 | 5.789218000 | 20.009809000 |
| 1 | 2.445627000 | 5.754124000 | 18.224849000 |
| 1 | 2.320400000 | 3.040360000 | 19.036641000 |
| 1 | 4.666096000 | 5.706837000 | 19.206785000 |
| 1 | 5.968178000 | 4.638982000 | 19.694776000 |
| 1 | 4.031622000 | 1.745079000 | 19.429989000 |
| 1 | 5.640689000 | 2.392087000 | 19.356793000 |
| 26 | 7.845585000 | 0.383255000 | 18.441414000 |
| 6 | 9.537129000 | 3.124678000 | 18.861094000 |
| 6 | 9.865936000 | 3.814121000 | 17.516978000 |
| 6 | 9.476671000 | 5.301167000 | 17.549359000 |
| 6 | 8.034216000 | 3.228409000 | 19.111655000 |
| 8 | 10.012284000 | 6.040656000 | 18.412741000 |
| 8 | 8.641518000 | 5.706328000 | 16.682774000 |
| 8 | 7.607852000 | 4.167517000 | 19.797201000 |
| 8 | 7.273140000 | 2.338393000 | 18.549372000 |
| 16 | 10.102440000 | 1.380008000 | 18.893018000 |
| 1 | 10.085118000 | 3.641381000 | 19.651897000 |
| 1 | 10.945405000 | 3.749249000 | 17.352066000 |
| 1 | 9.360752000 | 3.306872000 | 16.694516000 |
| 8 | 8.637063000 | 0.286767000 | 16.493390000 |
| 8 | 14.957773000 | 0.740818000 | 17.532044000 |
| 1 | 14.165136000 | 0.491571000 | 17.002714000 |
| 1 | 15.724449000 | 0.450368000 | 17.017409000 |
| 8 | 12.578558000 | 0.948138000 | 16.199910000 |
| 1 | 11.746364000 | 0.549016000 | 16.518746000 |
| 1 | 12.658374000 | 1.791390000 | 16.724026000 |
| 8 | 12.472565000 | 3.581775000 | 20.153993000 |
| 1 | 12.576608000 | 4.570542000 | 19.978589000 |
| 1 | 13.072772000 | 3.376174000 | 20.886036000 |
| 8 | 12.496473000 | 6.107914000 | 19.507467000 |
| 1 | 11.559498000 | 6.083409000 | 19.202289000 |
| 1 | 12.992238000 | 6.416097000 | 18.718217000 |
| 1 | 7.126490000 | -4.454185000 | 21.562823000 |
| 1 | 4.772157000 | -5.169232000 | 19.958337000 |
| 1 | -0.099105000 | -0.895406000 | 18.953078000 |
| 1 | 1.182861000 | 1.873355000 | 15.452546000 |

| | | | |
|---|---|---|---|
| 1 | 1.087713000 | 5.010489000 | 19.087809000 |
| 1 | 4.483215000 | 7.641368000 | 15.255608000 |
| 1 | 18.153245000 | 4.302198000 | 10.292753000 |
| 1 | 14.730734000 | 7.809181000 | 8.455444000 |
| 1 | 10.252890000 | 3.289848000 | 5.070788000 |
| 1 | 5.871081000 | 1.188237000 | 7.415018000 |
| 1 | 10.837193000 | -0.795077000 | 9.563001000 |
| 1 | 4.036888000 | -2.741353000 | 21.763846000 |
| 1 | 8.341938000 | 8.601809000 | 10.641989000 |
| 1 | 13.128020000 | 9.291260000 | 9.242997000 |
| 1 | 15.460256000 | 9.934567000 | 13.524681000 |
| 1 | 2.450161000 | -1.342141000 | 10.094974000 |
| 1 | 6.282244000 | -4.712497000 | 10.483988000 |
| 1 | 8.987869000 | -4.532612000 | 13.176227000 |
| 8 | 8.275022000 | 4.600098000 | 14.242575000 |
| 1 | 8.418042000 | 4.991868000 | 15.138325000 |
| 1 | 8.010983000 | 3.678889000 | 14.429140000 |
| 8 | 13.391379000 | 2.983939000 | 17.745207000 |
| 1 | 13.028916000 | 3.084562000 | 18.674161000 |
| 1 | 14.199598000 | 2.430932000 | 17.825761000 |
| 8 | 9.844742000 | 0.837146000 | 16.454088000 |

**$^{5}$IM1$_{S}$:**

| | | | |
|---|---|---|---|
| 6 | 14.705496000 | 8.938492000 | 13.186091000 |
| 6 | 13.538540000 | 9.227335000 | 12.256256000 |
| 8 | 12.465122000 | 9.705103000 | 12.649099000 |
| 6 | 14.257724000 | 8.405015000 | 14.549139000 |
| 6 | 13.402072000 | 7.131973000 | 14.451091000 |
| 6 | 13.266717000 | 6.467084000 | 15.810026000 |
| 8 | 13.097324000 | 7.147537000 | 16.841395000 |
| 7 | 13.365138000 | 5.131043000 | 15.836461000 |
| 1 | 15.394395000 | 8.237099000 | 12.704204000 |
| 1 | 15.150099000 | 8.202237000 | 15.151504000 |
| 1 | 13.696022000 | 9.172767000 | 15.086961000 |
| 1 | 13.831669000 | 6.431931000 | 13.725810000 |
| 1 | 12.403395000 | 7.378796000 | 14.072358000 |
| 1 | 13.385141000 | 4.595589000 | 16.708589000 |
| 1 | 13.527077000 | 4.597935000 | 14.986711000 |
| 7 | 13.755938000 | 8.933390000 | 10.951234000 |
| 6 | 12.647583000 | 9.020242000 | 10.024060000 |
| 6 | 11.505233000 | 8.103010000 | 10.494777000 |
| 8 | 11.758147000 | 6.983223000 | 10.948563000 |
| 1 | 14.580869000 | 8.376621000 | 10.713923000 |
| 1 | 12.307519000 | 10.057374000 | 9.936568000 |
| 7 | 10.253155000 | 8.588608000 | 10.368579000 |
| 6 | 9.143850000 | 8.026279000 | 11.146562000 |
| 6 | 9.192938000 | 8.623862000 | 12.560233000 |
| 6 | 7.908173000 | 8.533102000 | 13.378980000 |
| 6 | 8.013814000 | 9.347987000 | 14.677669000 |
| 7 | 8.922736000 | 8.775123000 | 15.664399000 |
| 6 | 10.171795000 | 9.146721000 | 15.929386000 |
| 7 | 10.857803000 | 9.990319000 | 15.113546000 |
| 7 | 10.754913000 | 8.744688000 | 17.068339000 |
| 1 | 10.183888000 | 9.578237000 | 10.166048000 |
| 1 | 9.253076000 | 6.940122000 | 11.160373000 |
| 1 | 10.013605000 | 8.133503000 | 13.094858000 |
| 1 | 9.463433000 | 9.685687000 | 12.458581000 |
| 1 | 7.069707000 | 8.944859000 | 12.799081000 |
| 1 | 7.677512000 | 7.486803000 | 13.607028000 |
| 1 | 8.337502000 | 10.372251000 | 14.463616000 |
| 1 | 7.031281000 | 9.407913000 | 15.158898000 |
| 1 | 8.583812000 | 7.950548000 | 16.199457000 |
| 1 | 11.769662000 | 10.272772000 | 15.445823000 |
| 1 | 10.843924000 | 9.857089000 | 14.108192000 |
| 1 | 10.214802000 | 8.145009000 | 17.709465000 |
| 1 | 11.747703000 | 8.505188000 | 17.027852000 |
| 7 | 6.059251000 | 1.303152000 | 8.301764000 |
| 6 | 7.034930000 | 2.394837000 | 8.216258000 |
| 6 | 8.409799000 | 1.903243000 | 7.764927000 |
| 8 | 8.979811000 | 0.964661000 | 8.332079000 |
| 6 | 7.220038000 | 3.081258000 | 9.586985000 |
| 6 | 6.214295000 | 4.160049000 | 9.929980000 |
| 6 | 4.919262000 | 4.201289000 | 9.399020000 |
| 6 | 6.573005000 | 5.132356000 | 10.873967000 |
| 6 | 3.998252000 | 5.167879000 | 9.807639000 |
| 6 | 5.662596000 | 6.100294000 | 11.290492000 |
| 6 | 4.365840000 | 6.114320000 | 10.767622000 |
| 8 | 3.420417000 | 7.024432000 | 11.170497000 |
| 1 | 6.474142000 | 0.528977000 | 8.826839000 |
| 1 | 6.656837000 | 3.130012000 | 7.494698000 |
| 1 | 7.184970000 | 2.294533000 | 10.353053000 |
| 1 | 8.223859000 | 3.512161000 | 9.660495000 |
| 1 | 4.611674000 | 3.445424000 | 8.683443000 |
| 1 | 7.564958000 | 5.104630000 | 11.313679000 |
| 1 | 2.992073000 | 5.193949000 | 9.400762000 |
| 1 | 5.956896000 | 6.831429000 | 12.039891000 |
| 1 | 3.815339000 | 7.593600000 | 11.849173000 |
| 7 | 8.988048000 | 2.563002000 | 6.727769000 |
| 6 | 10.402080000 | 2.361069000 | 6.418548000 |
| 6 | 11.240606000 | 3.289804000 | 7.299725000 |
| 8 | 11.662033000 | 4.365334000 | 6.881924000 |
| 1 | 8.565813000 | 3.427530000 | 6.415139000 |
| 1 | 10.629829000 | 1.307952000 | 6.592997000 |
| 7 | 11.390089000 | 2.897959000 | 8.591416000 |
| 6 | 11.902175000 | 3.870492000 | 9.541663000 |
| 6 | 13.346593000 | 4.252492000 | 9.176270000 |
| 8 | 14.210205000 | 3.445534000 | 8.826781000 |
| 6 | 11.778742000 | 3.381265000 | 10.995289000 |
| 6 | 10.664456000 | 4.127200000 | 11.731011000 |
| 8 | 9.695446000 | 4.576104000 | 11.115262000 |
| 7 | 10.813737000 | 4.259330000 | 13.066186000 |
| 1 | 10.975895000 | 2.029903000 | 8.912610000 |
| 1 | 11.293803000 | 4.772742000 | 9.444125000 |
| 1 | 12.724760000 | 3.493448000 | 11.531515000 |
| 1 | 11.548515000 | 2.311761000 | 11.018148000 |
| 1 | 11.566740000 | 3.792016000 | 13.550558000 |
| 1 | 10.050325000 | 4.702551000 | 13.596609000 |
| 7 | 13.575019000 | 5.585800000 | 9.281983000 |
| 6 | 14.808536000 | 6.174979000 | 8.807837000 |
| 6 | 15.867466000 | 6.132049000 | 9.903240000 |
| 8 | 16.049013000 | 7.084740000 | 10.675010000 |
| 1 | 12.906792000 | 6.159156000 | 9.799520000 |
| 1 | 15.132676000 | 5.621721000 | 7.922228000 |
| 7 | 16.512013000 | 4.946705000 | 9.974333000 |
| 6 | 17.382649000 | 4.579288000 | 11.078717000 |
| 6 | 16.721311000 | 3.545608000 | 12.000818000 |
| 6 | 15.446655000 | 4.087846000 | 12.644159000 |
| 16 | 14.529883000 | 2.854562000 | 13.643122000 |
| 6 | 15.669847000 | 2.680451000 | 15.060264000 |
| 1 | 16.103838000 | 4.201112000 | 9.408500000 |
| 1 | 17.616204000 | 5.499908000 | 11.619294000 |
| 1 | 16.476410000 | 2.643535000 | 11.427163000 |
| 1 | 17.445449000 | 3.251744000 | 12.769661000 |
| 1 | 15.662558000 | 4.960486000 | 13.271544000 |
| 1 | 14.742317000 | 4.409682000 | 11.871947000 |
| 1 | 15.226454000 | 1.961047000 | 15.750176000 |
| 1 | 15.788697000 | 3.635515000 | 15.580154000 |
| 1 | 16.643615000 | 2.308695000 | 14.733139000 |
| 6 | 3.685530000 | -3.131474000 | 20.888403000 |
| 6 | 3.302389000 | -2.886974000 | 19.434041000 |
| 8 | 3.982708000 | -3.292572000 | 18.493069000 |
| 6 | 3.968181000 | -1.839510000 | 21.684537000 |
| 6 | 5.064429000 | -1.006361000 | 21.094108000 |
| 7 | 5.880297000 | -0.180491000 | 21.856518000 |
| 6 | 5.496336000 | -0.822367000 | 19.802939000 |
| 6 | 6.754162000 | 0.450647000 | 21.027948000 |
| 7 | 6.545820000 | 0.074125000 | 19.780332000 |
| 1 | 2.869941000 | -3.671509000 | 21.384431000 |
| 1 | 3.053425000 | -1.234939000 | 21.772468000 |
| 1 | 4.235801000 | -2.125874000 | 22.708853000 |

| | | | | | | | |
|---|---|---|---|---|---|---|---|
| 1 | 5.151644000 | -1.281074000 | 18.892929000 | 6 | 5.286387000 | -3.297035000 | 11.276227000 |
| 1 | 7.500457000 | 1.156632000 | 21.360837000 | 6 | 4.330260000 | -2.234748000 | 11.842893000 |
| 1 | 5.824040000 | -0.057576000 | 22.858204000 | 8 | 3.946192000 | -2.279490000 | 13.024597000 |
| 7 | 2.150576000 | -2.173153000 | 19.242334000 | 1 | 7.281968000 | -3.340153000 | 12.005117000 |
| 6 | 1.747589000 | -1.738112000 | 17.932129000 | 1 | 5.029983000 | -3.518636000 | 10.235165000 |
| 6 | 1.772701000 | -0.215250000 | 17.768883000 | 7 | 3.987514000 | -1.294807000 | 10.946761000 |
| 8 | 2.161626000 | 0.536033000 | 18.672257000 | 6 | 3.137179000 | -0.142263000 | 11.217230000 |
| 1 | 1.698101000 | -1.750619000 | 20.040413000 | 6 | 3.851064000 | 1.200675000 | 11.007815000 |
| 1 | 2.457583000 | -2.180991000 | 17.225939000 | 6 | 4.808537000 | 1.613758000 | 12.103301000 |
| 7 | 1.316588000 | 0.205601000 | 16.568161000 | 6 | 5.739716000 | 0.737984000 | 12.674859000 |
| 6 | 1.618311000 | 1.515109000 | 15.994925000 | 6 | 4.819787000 | 2.943300000 | 12.544790000 |
| 6 | 2.745507000 | 1.403403000 | 14.948650000 | 6 | 6.620592000 | 1.152697000 | 13.670849000 |
| 6 | 3.956085000 | 0.741715000 | 15.521617000 | 6 | 5.709837000 | 3.379736000 | 13.517529000 |
| 7 | 4.295357000 | -0.580322000 | 15.276293000 | 6 | 6.597947000 | 2.477629000 | 14.108381000 |
| 6 | 4.876453000 | 1.191369000 | 16.433863000 | 8 | 7.415249000 | 2.937548000 | 15.093873000 |
| 6 | 5.402486000 | -0.867792000 | 16.003036000 | 1 | 4.555499000 | -1.276949000 | 10.099793000 |
| 7 | 5.773524000 | 0.180472000 | 16.729738000 | 1 | 2.771746000 | -0.250938000 | 12.240686000 |
| 1 | 1.076838000 | -0.520398000 | 15.902405000 | 1 | 3.079262000 | 1.973039000 | 10.910597000 |
| 1 | 1.928305000 | 2.163159000 | 16.815137000 | 1 | 4.373947000 | 1.183893000 | 10.039737000 |
| 1 | 2.387707000 | 0.840281000 | 14.079802000 | 1 | 5.794252000 | -0.287709000 | 12.338842000 |
| 1 | 3.000070000 | 2.401387000 | 14.580538000 | 1 | 4.146682000 | 3.662126000 | 12.088843000 |
| 1 | 4.992298000 | 2.175151000 | 16.852776000 | 1 | 7.315768000 | 0.454819000 | 14.123753000 |
| 1 | 5.892011000 | -1.824817000 | 15.977675000 | 1 | 5.745258000 | 4.422164000 | 13.804755000 |
| 1 | 3.973120000 | -1.155646000 | 14.486354000 | 1 | 7.951452000 | 2.185428000 | 15.493597000 |
| 6 | 8.065214000 | -3.925749000 | 14.333045000 | 6 | 5.303971000 | 7.100928000 | 17.109352000 |
| 6 | 6.661700000 | -4.385347000 | 14.771116000 | 6 | 5.070348000 | 6.205498000 | 15.887381000 |
| 6 | 8.232414000 | -2.402472000 | 14.343638000 | 6 | 4.400419000 | 4.893370000 | 16.311300000 |
| 6 | 6.494398000 | -4.411236000 | 16.294678000 | 6 | 4.251725000 | 6.923538000 | 14.807005000 |
| 1 | 8.817319000 | -4.387474000 | 14.988763000 | 1 | 4.348488000 | 7.349474000 | 17.596853000 |
| 1 | 6.460616000 | -5.390857000 | 14.380081000 | 1 | 5.964660000 | 6.610684000 | 17.829880000 |
| 1 | 5.904636000 | -3.727169000 | 14.321688000 | 1 | 6.054091000 | 5.957821000 | 15.471285000 |
| 1 | 8.066170000 | -1.986229000 | 15.338571000 | 1 | 4.200089000 | 4.247413000 | 15.452167000 |
| 1 | 7.508353000 | -1.924904000 | 13.676683000 | 1 | 3.442551000 | 5.089599000 | 16.815888000 |
| 1 | 9.229939000 | -2.089474000 | 14.017690000 | 1 | 5.051289000 | 4.345331000 | 16.998273000 |
| 1 | 5.453536000 | -4.564662000 | 16.595381000 | 1 | 4.103106000 | 6.287896000 | 13.926113000 |
| 1 | 6.831271000 | -3.479416000 | 16.761030000 | 1 | 3.260513000 | 7.207733000 | 15.185025000 |
| 1 | 7.097221000 | -5.216442000 | 16.731442000 | 1 | 4.757055000 | 7.842727000 | 14.482379000 |
| 7 | 6.514416000 | -4.957372000 | 19.868333000 | 6 | 1.578332000 | 4.567747000 | 19.431884000 |
| 6 | 7.903672000 | -5.009118000 | 20.318174000 | 7 | 2.433325000 | 3.397837000 | 19.324873000 |
| 6 | 9.012067000 | -4.840216000 | 19.252376000 | 6 | 3.768904000 | 3.429200000 | 19.471472000 |
| 6 | 9.016732000 | -3.463702000 | 18.667418000 | 7 | 4.424169000 | 4.578968000 | 19.661795000 |
| 7 | 9.828629000 | -3.053127000 | 17.616107000 | 7 | 4.448016000 | 2.273716000 | 19.492306000 |
| 6 | 8.257428000 | -2.362830000 | 18.963918000 | 1 | 1.769757000 | 5.106406000 | 20.366324000 |
| 6 | 9.537748000 | -1.761132000 | 17.315686000 | 1 | 1.717205000 | 5.254599000 | 18.587263000 |
| 7 | 8.577279000 | -1.320037000 | 18.112749000 | 1 | 2.033812000 | 2.485334000 | 19.115762000 |
| 1 | 6.317902000 | -4.148025000 | 19.284295000 | 1 | 3.978933000 | 5.443584000 | 19.389426000 |
| 1 | 8.053758000 | -5.970827000 | 20.818595000 | 1 | 5.463852000 | 4.567910000 | 19.593092000 |
| 1 | 8.872401000 | -5.585347000 | 18.456242000 | 1 | 3.934074000 | 1.426972000 | 19.270464000 |
| 1 | 9.985340000 | -5.056905000 | 19.715791000 | 1 | 5.432126000 | 2.304458000 | 19.190836000 |
| 1 | 7.497538000 | -2.252580000 | 19.716840000 | 26 | 7.582519000 | 0.592240000 | 17.888011000 |
| 1 | 10.001482000 | -1.186407000 | 16.529874000 | 6 | 9.215672000 | 3.535589000 | 18.561975000 |
| 1 | 10.521547000 | -3.620464000 | 17.147831000 | 6 | 9.649073000 | 4.455871000 | 17.411165000 |
| 7 | 9.417170000 | -1.867828000 | 9.435494000 | 6 | 9.186184000 | 5.910000000 | 17.596167000 |
| 6 | 8.709197000 | -1.677287000 | 10.697659000 | 6 | 7.689631000 | 3.522657000 | 18.760433000 |
| 6 | 7.227443000 | -1.977133000 | 10.483145000 | 8 | 9.685744000 | 6.591898000 | 18.526773000 |
| 8 | 6.539527000 | -1.417511000 | 9.621690000 | 8 | 8.360953000 | 6.357437000 | 16.742834000 |
| 6 | 8.839081000 | -0.259217000 | 11.320078000 | 8 | 7.184362000 | 4.509015000 | 19.317286000 |
| 6 | 10.185052000 | 0.002015000 | 11.915243000 | 8 | 7.010065000 | 2.503077000 | 18.355199000 |
| 7 | 11.363713000 | -0.192773000 | 11.209369000 | 16 | 9.964515000 | 1.863188000 | 18.376940000 |
| 6 | 10.476439000 | 0.454099000 | 13.186020000 | 1 | 9.677627000 | 3.901950000 | 19.485303000 |
| 6 | 12.339318000 | 0.130444000 | 12.043861000 | 1 | 10.743349000 | 4.459629000 | 17.379062000 |
| 7 | 11.854015000 | 0.535962000 | 13.249239000 | 1 | 9.278344000 | 4.071684000 | 16.462439000 |
| 1 | 9.071516000 | -1.160332000 | 8.786042000 | 8 | 8.710466000 | 0.957136000 | 16.239253000 |
| 1 | 9.109748000 | -2.404418000 | 11.416569000 | 8 | 14.503035000 | 0.482420000 | 17.707601000 |
| 1 | 8.620878000 | 0.458336000 | 10.524125000 | 1 | 13.787160000 | 0.433539000 | 17.031574000 |
| 1 | 8.091523000 | -0.113293000 | 12.099148000 | 1 | 15.282853000 | 0.082507000 | 17.295818000 |
| 1 | 9.847819000 | 0.727742000 | 14.021641000 | 8 | 12.551647000 | 1.205629000 | 15.936459000 |
| 1 | 13.396407000 | 0.115436000 | 11.818826000 | 1 | 11.601150000 | 1.086099000 | 16.145532000 |
| 1 | 12.378110000 | 0.846524000 | 14.069054000 | 1 | 12.769723000 | 2.029222000 | 16.449756000 |
| 7 | 6.677949000 | -2.875992000 | 11.346206000 | 8 | 12.145760000 | 3.646325000 | 19.982028000 |

| | | | |
|---|---|---|---|
| 1 | 12.212766000 | 4.634500000 | 19.769899000 |
| 1 | 12.714505000 | 3.517464000 | 20.756369000 |
| 8 | 12.216418000 | 6.161570000 | 19.330195000 |
| 1 | 11.258866000 | 6.329698000 | 19.141944000 |
| 1 | 12.673899000 | 6.427871000 | 18.504064000 |
| 1 | 8.046562000 | -4.239621000 | 21.087340000 |
| 1 | 6.285049000 | -5.771766000 | 19.303161000 |
| 1 | 0.746541000 | -2.112843000 | 17.679283000 |
| 1 | 0.711814000 | 1.936325000 | 15.548140000 |
| 1 | 0.539301000 | 4.235487000 | 19.433752000 |
| 1 | 5.780627000 | 8.046942000 | 16.828741000 |
| 1 | 18.319571000 | 4.173761000 | 10.680574000 |
| 1 | 14.634259000 | 7.215570000 | 8.526796000 |
| 1 | 10.587195000 | 2.615774000 | 5.374576000 |
| 1 | 5.885409000 | 0.930446000 | 7.368081000 |
| 1 | 10.389647000 | -1.607496000 | 9.610840000 |
| 1 | 4.566553000 | -3.779842000 | 20.879707000 |
| 1 | 8.207790000 | 8.273466000 | 10.635301000 |
| 1 | 12.989439000 | 8.694098000 | 9.037420000 |
| 1 | 15.257398000 | 9.876830000 | 13.329617000 |
| 1 | 2.270387000 | -0.198747000 | 10.547121000 |
| 1 | 5.170374000 | -4.201409000 | 11.873645000 |
| 1 | 8.295965000 | -4.318558000 | 13.331480000 |
| 8 | 8.479281000 | 5.371267000 | 14.244752000 |
| 1 | 8.461384000 | 5.753943000 | 15.156691000 |
| 1 | 8.078567000 | 4.491684000 | 14.389463000 |
| 8 | 13.438220000 | 3.024997000 | 17.730658000 |
| 1 | 12.912056000 | 3.144785000 | 18.573856000 |
| 1 | 14.094708000 | 2.320765000 | 17.923419000 |
| 8 | 9.967102000 | 1.595464000 | 16.716913000 |

**$^{5}$TS2$_{S}$:**

| | | | |
|---|---|---|---|
| 6 | 14.683958000 | 8.961970000 | 13.170121000 |
| 6 | 13.508675000 | 9.249293000 | 12.250354000 |
| 8 | 12.434426000 | 9.716545000 | 12.653614000 |
| 6 | 14.250314000 | 8.414138000 | 14.531924000 |
| 6 | 13.400957000 | 7.137064000 | 14.430094000 |
| 6 | 13.285323000 | 6.458610000 | 15.783899000 |
| 8 | 13.116882000 | 7.128811000 | 16.822591000 |
| 7 | 13.400644000 | 5.124237000 | 15.799073000 |
| 1 | 15.374595000 | 8.269570000 | 12.677971000 |
| 1 | 15.149020000 | 8.211153000 | 15.124762000 |
| 1 | 13.689182000 | 9.173781000 | 15.081735000 |
| 1 | 13.825924000 | 6.447004000 | 13.692690000 |
| 1 | 12.396610000 | 7.381840000 | 14.065664000 |
| 1 | 13.436882000 | 4.582412000 | 16.667062000 |
| 1 | 13.559166000 | 4.598151000 | 14.944573000 |
| 7 | 13.719991000 | 8.966650000 | 10.941885000 |
| 6 | 12.605821000 | 9.053477000 | 10.021654000 |
| 6 | 11.472022000 | 8.125563000 | 10.491556000 |
| 8 | 11.733594000 | 7.002317000 | 10.931166000 |
| 1 | 14.546799000 | 8.416346000 | 10.695803000 |
| 1 | 12.258893000 | 10.089119000 | 9.943600000 |
| 7 | 10.216331000 | 8.606094000 | 10.380957000 |
| 6 | 9.116658000 | 8.026502000 | 11.159748000 |
| 6 | 9.172300000 | 8.604007000 | 12.581456000 |
| 6 | 7.893453000 | 8.494008000 | 13.407059000 |
| 6 | 8.000629000 | 9.294641000 | 14.714258000 |
| 7 | 8.920446000 | 8.719213000 | 15.689431000 |
| 6 | 10.168181000 | 9.099399000 | 15.949338000 |
| 7 | 10.842974000 | 9.952108000 | 15.133980000 |
| 7 | 10.761041000 | 8.696601000 | 17.082790000 |
| 1 | 10.140285000 | 9.598093000 | 10.193024000 |
| 1 | 9.233354000 | 6.941063000 | 11.158125000 |
| 1 | 9.998563000 | 8.109894000 | 13.103727000 |
| 1 | 9.436321000 | 9.668608000 | 12.493121000 |
| 1 | 7.048342000 | 8.906409000 | 12.837281000 |
| 1 | 7.672900000 | 7.443367000 | 13.624878000 |
| 1 | 8.314530000 | 10.323916000 | 14.509681000 |
| 1 | 7.020668000 | 9.341118000 | 15.202059000 |
| 1 | 8.589879000 | 7.891944000 | 16.224831000 |
| 1 | 11.754488000 | 10.240046000 | 15.462394000 |
| 1 | 10.822852000 | 9.827031000 | 14.127691000 |
| 1 | 10.228656000 | 8.092139000 | 17.725905000 |
| 1 | 11.755231000 | 8.463885000 | 17.034185000 |
| 7 | 6.066441000 | 1.303492000 | 8.301850000 |
| 6 | 7.040482000 | 2.396244000 | 8.212204000 |
| 6 | 8.415085000 | 1.904998000 | 7.759575000 |
| 8 | 8.990018000 | 0.972193000 | 8.331349000 |
| 6 | 7.226411000 | 3.085841000 | 9.580964000 |
| 6 | 6.218667000 | 4.162758000 | 9.923647000 |
| 6 | 4.923508000 | 4.201822000 | 9.392856000 |
| 6 | 6.575653000 | 5.135267000 | 10.868088000 |
| 6 | 4.000433000 | 5.166119000 | 9.802348000 |
| 6 | 5.663328000 | 6.101223000 | 11.285024000 |
| 6 | 4.366277000 | 6.112732000 | 10.762746000 |
| 8 | 3.419228000 | 7.020689000 | 11.166735000 |
| 1 | 6.483253000 | 0.532519000 | 8.830135000 |
| 1 | 6.660604000 | 3.129309000 | 7.489364000 |
| 1 | 7.194505000 | 2.300928000 | 10.348961000 |
| 1 | 8.229409000 | 3.519056000 | 9.652261000 |
| 1 | 4.617244000 | 3.445636000 | 8.677092000 |
| 1 | 7.567471000 | 5.108129000 | 11.308344000 |
| 1 | 2.994094000 | 5.190201000 | 9.395752000 |
| 1 | 5.956043000 | 6.832896000 | 12.034443000 |
| 1 | 3.813912000 | 7.591058000 | 11.844540000 |
| 7 | 8.986900000 | 2.558478000 | 6.714771000 |
| 6 | 10.400900000 | 2.360041000 | 6.402396000 |
| 6 | 11.237461000 | 3.293916000 | 7.280107000 |
| 8 | 11.652184000 | 4.371090000 | 6.859934000 |
| 1 | 8.561354000 | 3.420299000 | 6.399095000 |
| 1 | 10.632569000 | 1.308067000 | 6.578505000 |
| 7 | 11.392745000 | 2.903149000 | 8.571425000 |
| 6 | 11.896957000 | 3.879712000 | 9.521949000 |
| 6 | 13.335328000 | 4.278915000 | 9.151918000 |
| 8 | 14.206808000 | 3.482042000 | 8.798497000 |
| 6 | 11.785186000 | 3.385082000 | 10.975038000 |
| 6 | 10.678491000 | 4.130760000 | 11.722727000 |
| 8 | 9.701577000 | 4.577778000 | 11.118183000 |
| 7 | 10.846380000 | 4.267769000 | 13.055607000 |
| 1 | 10.976084000 | 2.036959000 | 8.894482000 |
| 1 | 11.277851000 | 4.775321000 | 9.429279000 |
| 1 | 12.735997000 | 3.493703000 | 11.503566000 |
| 1 | 11.553840000 | 2.315373000 | 10.996545000 |
| 1 | 11.590857000 | 3.778229000 | 13.531324000 |
| 1 | 10.082903000 | 4.697684000 | 13.596669000 |
| 7 | 13.549799000 | 5.614074000 | 9.260554000 |
| 6 | 14.775778000 | 6.217326000 | 8.784625000 |
| 6 | 15.839545000 | 6.181036000 | 9.875706000 |
| 8 | 16.020510000 | 7.135963000 | 10.644973000 |
| 1 | 12.876751000 | 6.180668000 | 9.779441000 |
| 1 | 15.102631000 | 5.670794000 | 7.895798000 |
| 7 | 16.490024000 | 4.999028000 | 9.945020000 |
| 6 | 17.370716000 | 4.636824000 | 11.043106000 |
| 6 | 16.731941000 | 3.581798000 | 11.956759000 |
| 6 | 15.452297000 | 4.094352000 | 12.614345000 |
| 16 | 14.570062000 | 2.835697000 | 13.614090000 |
| 6 | 15.730622000 | 2.680042000 | 15.016959000 |
| 1 | 16.084763000 | 4.251392000 | 9.379480000 |
| 1 | 17.589057000 | 5.556386000 | 11.591860000 |
| 1 | 16.498990000 | 2.682070000 | 11.374553000 |
| 1 | 17.465850000 | 3.291755000 | 12.717690000 |
| 1 | 15.656925000 | 4.966547000 | 13.246308000 |
| 1 | 14.735538000 | 4.407614000 | 11.850075000 |
| 1 | 15.314335000 | 1.947016000 | 15.709291000 |
| 1 | 15.834115000 | 3.635120000 | 15.540139000 |
| 1 | 16.708464000 | 2.332704000 | 14.675460000 |
| 6 | 3.653285000 | -3.090547000 | 20.868856000 |
| 6 | 3.254144000 | -2.858900000 | 19.416656000 |
| 8 | 3.923754000 | -3.273167000 | 18.472059000 |

| | | | |
|---|---|---|---|
| 6 | 3.954105000 | -1.792206000 | 21.647410000 |
| 6 | 5.048378000 | -0.971482000 | 21.036227000 |
| 7 | 5.858847000 | -0.122172000 | 21.778144000 |
| 6 | 5.480748000 | -0.818310000 | 19.741085000 |
| 6 | 6.728327000 | 0.494597000 | 20.934809000 |
| 7 | 6.523072000 | 0.085852000 | 19.696794000 |
| 1 | 2.840128000 | -3.619196000 | 21.380823000 |
| 1 | 3.044667000 | -1.180259000 | 21.739455000 |
| 1 | 4.231920000 | -2.069609000 | 22.671527000 |
| 1 | 5.139164000 | -1.304802000 | 18.844391000 |
| 1 | 7.466252000 | 1.216425000 | 21.251521000 |
| 1 | 5.799803000 | 0.027702000 | 22.776005000 |
| 7 | 2.099659000 | -2.147125000 | 19.231497000 |
| 6 | 1.686190000 | -1.718050000 | 17.922607000 |
| 6 | 1.730819000 | -0.196951000 | 17.746357000 |
| 8 | 2.133697000 | 0.556049000 | 18.642311000 |
| 1 | 1.658493000 | -1.715152000 | 20.030922000 |
| 1 | 2.380710000 | -2.177257000 | 17.211501000 |
| 7 | 1.275635000 | 0.221513000 | 16.544746000 |
| 6 | 1.584393000 | 1.526524000 | 15.965044000 |
| 6 | 2.706712000 | 1.403521000 | 14.915072000 |
| 6 | 3.916131000 | 0.736935000 | 15.484827000 |
| 7 | 4.256983000 | -0.583145000 | 15.230873000 |
| 6 | 4.839675000 | 1.183982000 | 16.395041000 |
| 6 | 5.369582000 | -0.872424000 | 15.945484000 |
| 7 | 5.739662000 | 0.172696000 | 16.677300000 |
| 1 | 1.022409000 | -0.504827000 | 15.884426000 |
| 1 | 1.901780000 | 2.175703000 | 16.781540000 |
| 1 | 2.342367000 | 0.839004000 | 14.049852000 |
| 1 | 2.966917000 | 2.398416000 | 14.541908000 |
| 1 | 4.954748000 | 2.165050000 | 16.820397000 |
| 1 | 5.867938000 | -1.824110000 | 15.904176000 |
| 1 | 3.932912000 | -1.158705000 | 14.441060000 |
| 6 | 8.059081000 | -3.926140000 | 14.357598000 |
| 6 | 6.656255000 | -4.375412000 | 14.806858000 |
| 6 | 8.235117000 | -2.404395000 | 14.356593000 |
| 6 | 6.491696000 | -4.369765000 | 16.331127000 |
| 1 | 8.812515000 | -4.389073000 | 15.011094000 |
| 1 | 6.454106000 | -5.388813000 | 14.437458000 |
| 1 | 5.898782000 | -3.727206000 | 14.343471000 |
| 1 | 8.069989000 | -1.976661000 | 15.346653000 |
| 1 | 7.515847000 | -1.928959000 | 13.683024000 |
| 1 | 9.234804000 | -2.099044000 | 14.030377000 |
| 1 | 5.451755000 | -4.518883000 | 16.637267000 |
| 1 | 6.828547000 | -3.428408000 | 16.778579000 |
| 1 | 7.096919000 | -5.164954000 | 16.782689000 |
| 7 | 6.486639000 | -4.946601000 | 19.902146000 |
| 6 | 7.868495000 | -4.998785000 | 20.375898000 |
| 6 | 8.992821000 | -4.833614000 | 19.326456000 |
| 6 | 8.997514000 | -3.461445000 | 18.731134000 |
| 7 | 9.804366000 | -3.059512000 | 17.672111000 |
| 6 | 8.234963000 | -2.360277000 | 19.017890000 |
| 6 | 9.506879000 | -1.773170000 | 17.356689000 |
| 7 | 8.546145000 | -1.327363000 | 18.151919000 |
| 1 | 6.308332000 | -4.144937000 | 19.301811000 |
| 1 | 8.009175000 | -5.958838000 | 20.882006000 |
| 1 | 8.867803000 | -5.584931000 | 18.533685000 |
| 1 | 9.959909000 | -5.042897000 | 19.805853000 |
| 1 | 7.476791000 | -2.246263000 | 19.772058000 |
| 1 | 9.963170000 | -1.200402000 | 16.565111000 |
| 1 | 10.497616000 | -3.629562000 | 17.207479000 |
| 7 | 9.448499000 | -1.847358000 | 9.471923000 |
| 6 | 8.725969000 | -1.652956000 | 10.725613000 |
| 6 | 7.247493000 | -1.957764000 | 10.496981000 |
| 8 | 6.563508000 | -1.396293000 | 9.633645000 |
| 6 | 8.847840000 | -0.231571000 | 11.343204000 |
| 6 | 10.182134000 | 0.025982000 | 11.965713000 |
| 7 | 11.371674000 | -0.160969000 | 11.275987000 |
| 6 | 10.453590000 | 0.461526000 | 13.247192000 |
| 6 | 12.333929000 | 0.150831000 | 12.130347000 |

| | | | |
|---|---|---|---|
| 7 | 11.830015000 | 0.543149000 | 13.332391000 |
| 1 | 9.109158000 | -1.142753000 | 8.815903000 |
| 1 | 9.120310000 | -2.376253000 | 11.451597000 |
| 1 | 8.649440000 | 0.481527000 | 10.537981000 |
| 1 | 8.083973000 | -0.077405000 | 12.103962000 |
| 1 | 9.814441000 | 0.721997000 | 14.079446000 |
| 1 | 13.394595000 | 0.137300000 | 11.922549000 |
| 1 | 12.345592000 | 0.825480000 | 14.170069000 |
| 7 | 6.693138000 | -2.863135000 | 11.351038000 |
| 6 | 5.305084000 | -3.292678000 | 11.259347000 |
| 6 | 4.335466000 | -2.239831000 | 11.820751000 |
| 8 | 3.937251000 | -2.295309000 | 12.997582000 |
| 1 | 7.297158000 | -3.344076000 | 11.998129000 |
| 1 | 5.062891000 | -3.508961000 | 10.213682000 |
| 7 | 3.998148000 | -1.294616000 | 10.928238000 |
| 6 | 3.142040000 | -0.146385000 | 11.200824000 |
| 6 | 3.852138000 | 1.200743000 | 11.006621000 |
| 6 | 4.811039000 | 1.605848000 | 12.104246000 |
| 6 | 5.729199000 | 0.720424000 | 12.681603000 |
| 6 | 4.841789000 | 2.937793000 | 12.538216000 |
| 6 | 6.617073000 | 1.128282000 | 13.673257000 |
| 6 | 5.741037000 | 3.367937000 | 13.506028000 |
| 6 | 6.618119000 | 2.456712000 | 14.099964000 |
| 8 | 7.449674000 | 2.907719000 | 15.077161000 |
| 1 | 4.577160000 | -1.266328000 | 10.088908000 |
| 1 | 2.768776000 | -0.263891000 | 12.220528000 |
| 1 | 3.077884000 | 1.971774000 | 10.917526000 |
| 1 | 4.374578000 | 1.195327000 | 10.038070000 |
| 1 | 5.768940000 | -0.308391000 | 12.354311000 |
| 1 | 4.177443000 | 3.663293000 | 12.079975000 |
| 1 | 7.302820000 | 0.425927000 | 14.132608000 |
| 1 | 5.791808000 | 4.411768000 | 13.786316000 |
| 1 | 7.951791000 | 2.141612000 | 15.497596000 |
| 6 | 5.317238000 | 7.069958000 | 17.125752000 |
| 6 | 5.096643000 | 6.177857000 | 15.898925000 |
| 6 | 4.439696000 | 4.856696000 | 16.314790000 |
| 6 | 4.273194000 | 6.891947000 | 14.819591000 |
| 1 | 4.357722000 | 7.305806000 | 17.611947000 |
| 1 | 5.981390000 | 6.583406000 | 17.845676000 |
| 1 | 6.083983000 | 5.941544000 | 15.484737000 |
| 1 | 4.254220000 | 4.211285000 | 15.451936000 |
| 1 | 3.476124000 | 5.039570000 | 16.813588000 |
| 1 | 5.093225000 | 4.315116000 | 17.004606000 |
| 1 | 4.135775000 | 6.259498000 | 13.934626000 |
| 1 | 3.277140000 | 7.161722000 | 15.195633000 |
| 1 | 4.767963000 | 7.819106000 | 14.501662000 |
| 6 | 1.568766000 | 4.585314000 | 19.430215000 |
| 7 | 2.417981000 | 3.413110000 | 19.303156000 |
| 6 | 3.753123000 | 3.435694000 | 19.453173000 |
| 7 | 4.413375000 | 4.578132000 | 19.667494000 |
| 7 | 4.427723000 | 2.277376000 | 19.452124000 |
| 1 | 1.763523000 | 5.107376000 | 20.373319000 |
| 1 | 1.711040000 | 5.285305000 | 18.596970000 |
| 1 | 2.014203000 | 2.505665000 | 19.080687000 |
| 1 | 3.978579000 | 5.450432000 | 19.403012000 |
| 1 | 5.453308000 | 4.555983000 | 19.616321000 |
| 1 | 3.909176000 | 1.435162000 | 19.223896000 |
| 1 | 5.412263000 | 2.306548000 | 19.152184000 |
| 26 | 7.559394000 | 0.581197000 | 17.782920000 |
| 6 | 9.218706000 | 3.498553000 | 18.625980000 |
| 6 | 9.655507000 | 4.391001000 | 17.455913000 |
| 6 | 9.205960000 | 5.850739000 | 17.626974000 |
| 6 | 7.689942000 | 3.484806000 | 18.802033000 |
| 8 | 9.716914000 | 6.537303000 | 18.548373000 |
| 8 | 8.374169000 | 6.295370000 | 16.778751000 |
| 8 | 7.177499000 | 4.448696000 | 19.389925000 |
| 8 | 7.011796000 | 2.485802000 | 18.341967000 |
| 16 | 9.947343000 | 1.810197000 | 18.455555000 |
| 1 | 9.668122000 | 3.884688000 | 19.545951000 |
| 1 | 10.749459000 | 4.382977000 | 17.415169000 |

| | | | |
|---|---|---|---|
| 1 | 9.274162000 | 3.992646000 | 16.517441000 |
| 8 | 8.551196000 | 0.849726000 | 16.271752000 |
| 8 | 14.694394000 | 0.525471000 | 17.734940000 |
| 1 | 13.960998000 | 0.423717000 | 17.083640000 |
| 1 | 15.473161000 | 0.126139000 | 17.320718000 |
| 8 | 12.660828000 | 1.105834000 | 16.002890000 |
| 1 | 11.730398000 | 0.983122000 | 16.292298000 |
| 1 | 12.883344000 | 1.964832000 | 16.451501000 |
| 8 | 12.207896000 | 3.608215000 | 19.918210000 |
| 1 | 12.270066000 | 4.601264000 | 19.724124000 |
| 1 | 12.759854000 | 3.469802000 | 20.702968000 |
| 8 | 12.264705000 | 6.132534000 | 19.314783000 |
| 1 | 11.303567000 | 6.292207000 | 19.138271000 |
| 1 | 12.711030000 | 6.403305000 | 18.483659000 |
| 1 | 7.999853000 | -4.226800000 | 21.144629000 |
| 1 | 6.263527000 | -5.768025000 | 19.344660000 |
| 1 | 0.676769000 | -2.080630000 | 17.686228000 |
| 1 | 0.679191000 | 1.952249000 | 15.520033000 |
| 1 | 0.528056000 | 4.258418000 | 19.426794000 |
| 1 | 5.783587000 | 8.022929000 | 16.851309000 |
| 1 | 18.313954000 | 4.252563000 | 10.638568000 |
| 1 | 14.589924000 | 7.257155000 | 8.508294000 |
| 1 | 10.582394000 | 2.613207000 | 5.357438000 |
| 1 | 5.893651000 | 0.926877000 | 7.369541000 |
| 1 | 10.418434000 | -1.584678000 | 9.658152000 |
| 1 | 4.529625000 | -3.745333000 | 20.857725000 |
| 1 | 8.175134000 | 8.274590000 | 10.659065000 |
| 1 | 12.944267000 | 8.736569000 | 9.030836000 |
| 1 | 15.229782000 | 9.903332000 | 13.317177000 |
| 1 | 2.280366000 | -0.200740000 | 10.524001000 |
| 1 | 5.186637000 | -4.201259000 | 11.849915000 |
| 1 | 8.279623000 | -4.325974000 | 13.356615000 |
| 8 | 8.510599000 | 5.346958000 | 14.263720000 |
| 1 | 8.488210000 | 5.711331000 | 15.182477000 |
| 1 | 8.120280000 | 4.460075000 | 14.392390000 |
| 8 | 13.542176000 | 3.025400000 | 17.687133000 |
| 1 | 13.012845000 | 3.134862000 | 18.529551000 |
| 1 | 14.230882000 | 2.355287000 | 17.888905000 |
| 8 | 10.154734000 | 1.590708000 | 16.892296000 |

**$^5$IM2$_S$:**

| | | | |
|---|---|---|---|
| 6 | 14.773746000 | 9.138737000 | 13.085915000 |
| 6 | 13.645204000 | 9.367814000 | 12.094679000 |
| 8 | 12.545724000 | 9.834662000 | 12.424095000 |
| 6 | 14.280108000 | 8.478142000 | 14.377742000 |
| 6 | 13.566982000 | 7.138471000 | 14.143320000 |
| 6 | 13.279723000 | 6.437070000 | 15.459987000 |
| 8 | 12.971804000 | 7.084437000 | 16.475856000 |
| 7 | 13.358283000 | 5.089460000 | 15.474735000 |
| 1 | 15.561459000 | 8.537463000 | 12.622566000 |
| 1 | 15.137538000 | 8.326444000 | 15.042659000 |
| 1 | 13.599512000 | 9.150652000 | 14.903506000 |
| 1 | 14.167069000 | 6.493391000 | 13.491753000 |
| 1 | 12.618415000 | 7.298894000 | 13.614991000 |
| 1 | 13.400979000 | 4.604154000 | 16.376412000 |
| 1 | 13.789181000 | 4.575593000 | 14.709244000 |
| 7 | 13.916007000 | 9.016710000 | 10.815339000 |
| 6 | 12.845079000 | 9.035457000 | 9.841366000 |
| 6 | 11.684172000 | 8.150321000 | 10.331282000 |
| 8 | 11.920492000 | 7.068142000 | 10.878510000 |
| 1 | 14.785756000 | 8.512078000 | 10.624332000 |
| 1 | 12.505712000 | 10.061784000 | 9.664613000 |
| 7 | 10.438516000 | 8.615809000 | 10.114064000 |
| 6 | 9.297334000 | 8.118318000 | 10.892488000 |
| 6 | 9.324179000 | 8.789932000 | 12.273973000 |
| 6 | 8.011064000 | 8.779950000 | 13.051061000 |
| 6 | 8.113416000 | 9.613092000 | 14.337933000 |
| 7 | 8.964084000 | 9.007121000 | 15.358560000 |
| 6 | 10.217704000 | 9.325910000 | 15.658793000 |
| 7 | 10.955422000 | 10.149598000 | 14.869349000 |
| 7 | 10.758936000 | 8.893433000 | 16.810148000 |
| 1 | 10.374905000 | 9.584240000 | 9.825709000 |
| 1 | 9.374020000 | 7.031471000 | 10.964979000 |
| 1 | 10.112226000 | 8.302886000 | 12.858509000 |
| 1 | 9.635899000 | 9.834448000 | 12.126101000 |
| 1 | 7.210996000 | 9.215150000 | 12.435566000 |
| 1 | 7.725826000 | 7.751404000 | 13.295595000 |
| 1 | 8.494458000 | 10.617232000 | 14.123573000 |
| 1 | 7.121026000 | 9.725729000 | 14.787868000 |
| 1 | 8.564537000 | 8.211315000 | 15.892272000 |
| 1 | 11.857746000 | 10.413186000 | 15.240015000 |
| 1 | 10.976646000 | 10.023967000 | 13.862387000 |
| 1 | 10.176016000 | 8.320341000 | 17.437626000 |
| 1 | 11.732229000 | 8.590729000 | 16.771085000 |
| 7 | 6.012080000 | 0.994134000 | 8.303154000 |
| 6 | 7.151704000 | 1.914533000 | 8.336306000 |
| 6 | 8.480823000 | 1.168287000 | 8.101028000 |
| 8 | 8.606675000 | -0.023603000 | 8.385040000 |
| 6 | 7.250381000 | 2.595187000 | 9.722465000 |
| 6 | 6.326800000 | 3.762101000 | 10.003162000 |
| 6 | 5.065532000 | 3.917522000 | 9.415625000 |
| 6 | 6.734124000 | 4.705893000 | 10.956345000 |
| 6 | 4.225005000 | 4.971267000 | 9.779887000 |
| 6 | 5.899816000 | 5.754109000 | 11.334736000 |
| 6 | 4.635434000 | 5.880272000 | 10.756965000 |
| 8 | 3.755186000 | 6.869427000 | 11.132362000 |
| 1 | 6.297966000 | 0.184356000 | 8.859724000 |
| 1 | 7.016598000 | 2.681754000 | 7.563370000 |
| 1 | 7.053003000 | 1.812029000 | 10.462822000 |
| 1 | 8.278922000 | 2.927556000 | 9.901967000 |
| 1 | 4.728340000 | 3.190308000 | 8.684277000 |
| 1 | 7.718731000 | 4.622271000 | 11.405481000 |
| 1 | 3.244282000 | 5.088607000 | 9.328648000 |
| 1 | 6.230704000 | 6.449472000 | 12.101996000 |
| 1 | 4.198797000 | 7.433656000 | 11.784551000 |
| 7 | 9.516182000 | 1.917152000 | 7.649446000 |
| 6 | 10.904350000 | 1.489593000 | 7.806955000 |
| 6 | 11.637575000 | 2.709181000 | 8.346929000 |
| 8 | 11.799295000 | 3.696125000 | 7.627415000 |
| 1 | 9.376499000 | 2.900976000 | 7.454894000 |
| 1 | 10.919194000 | 0.630908000 | 8.481195000 |
| 7 | 11.967200000 | 2.704764000 | 9.658471000 |
| 6 | 12.396322000 | 3.966718000 | 10.222495000 |
| 6 | 13.819654000 | 4.307471000 | 9.755054000 |
| 8 | 14.715978000 | 3.476124000 | 9.588373000 |
| 6 | 12.248401000 | 4.030788000 | 11.743561000 |
| 6 | 10.801628000 | 4.364281000 | 12.130847000 |
| 8 | 9.899865000 | 4.351169000 | 11.292075000 |
| 7 | 10.599652000 | 4.700477000 | 13.423030000 |
| 1 | 11.822667000 | 1.883246000 | 10.239533000 |
| 1 | 11.736322000 | 4.730671000 | 9.808680000 |
| 1 | 12.892719000 | 4.826381000 | 12.131595000 |
| 1 | 12.564613000 | 3.100097000 | 12.223265000 |
| 1 | 11.348472000 | 4.671311000 | 14.101939000 |
| 1 | 9.652774000 | 4.994489000 | 13.706826000 |
| 7 | 13.989179000 | 5.635536000 | 9.536357000 |
| 6 | 15.217298000 | 6.156217000 | 8.975166000 |
| 6 | 16.265179000 | 6.332204000 | 10.070093000 |
| 8 | 16.348662000 | 7.383720000 | 10.721210000 |
| 1 | 13.278927000 | 6.268615000 | 9.905815000 |
| 1 | 15.563718000 | 5.466267000 | 8.200794000 |
| 7 | 17.017564000 | 5.231535000 | 10.283683000 |
| 6 | 17.895189000 | 5.071151000 | 11.434478000 |
| 6 | 17.420736000 | 3.927799000 | 12.342429000 |
| 6 | 15.995283000 | 4.161356000 | 12.838483000 |
| 16 | 15.306138000 | 2.821644000 | 13.875688000 |
| 6 | 16.328631000 | 3.012055000 | 15.376579000 |
| 1 | 16.679423000 | 4.388454000 | 9.821974000 |
| 1 | 17.897134000 | 6.026369000 | 11.965867000 |
| 1 | 17.458991000 | 2.976765000 | 11.797480000 |

| | | | |
|---|---|---|---|
| 1 | 18.118165000 | 3.842701000 | 13.183654000 |
| 1 | 15.931671000 | 5.108146000 | 13.389106000 |
| 1 | 15.316701000 | 4.219922000 | 11.985454000 |
| 1 | 15.991446000 | 2.266409000 | 16.098627000 |
| 1 | 16.190729000 | 4.005344000 | 15.814393000 |
| 1 | 17.385724000 | 2.843210000 | 15.157633000 |
| 6 | 4.836473000 | -3.296994000 | 20.609242000 |
| 6 | 4.272710000 | -2.960201000 | 19.239199000 |
| 8 | 4.961953000 | -2.979585000 | 18.217211000 |
| 6 | 5.216477000 | -2.023391000 | 21.408968000 |
| 6 | 6.210580000 | -1.150759000 | 20.707793000 |
| 7 | 7.456990000 | -0.823411000 | 21.220962000 |
| 6 | 6.160650000 | -0.516914000 | 19.491090000 |
| 6 | 8.097716000 | -0.024820000 | 20.324768000 |
| 7 | 7.333048000 | 0.172343000 | 19.264803000 |
| 1 | 4.101570000 | -3.866403000 | 21.190067000 |
| 1 | 4.309999000 | -1.441486000 | 21.618548000 |
| 1 | 5.613055000 | -2.335124000 | 22.381428000 |
| 1 | 5.382684000 | -0.531830000 | 18.749087000 |
| 1 | 9.091944000 | 0.369676000 | 20.471291000 |
| 1 | 7.819269000 | -1.109840000 | 22.120095000 |
| 7 | 2.956848000 | -2.600328000 | 19.208646000 |
| 6 | 2.317377000 | -2.165415000 | 17.994674000 |
| 6 | 2.063314000 | -0.653677000 | 17.937725000 |
| 8 | 2.463956000 | 0.113971000 | 18.821129000 |
| 1 | 2.454683000 | -2.506599000 | 20.080018000 |
| 1 | 2.989093000 | -2.428819000 | 17.172585000 |
| 7 | 1.365020000 | -0.266683000 | 16.846739000 |
| 6 | 1.328222000 | 1.107522000 | 16.351103000 |
| 6 | 2.361600000 | 1.325019000 | 15.225744000 |
| 6 | 3.721393000 | 0.874838000 | 15.642428000 |
| 7 | 4.221418000 | -0.380023000 | 15.328728000 |
| 6 | 4.669993000 | 1.458559000 | 16.442560000 |
| 6 | 5.439406000 | -0.504118000 | 15.895997000 |
| 7 | 5.732776000 | 0.588002000 | 16.595560000 |
| 1 | 1.151904000 | -0.989116000 | 16.168107000 |
| 1 | 1.556464000 | 1.759554000 | 17.194178000 |
| 1 | 2.055818000 | 0.785869000 | 14.321729000 |
| 1 | 2.386454000 | 2.385202000 | 14.957171000 |
| 1 | 4.703048000 | 2.451455000 | 16.858958000 |
| 1 | 6.066536000 | -1.373578000 | 15.799863000 |
| 1 | 3.858769000 | -1.016213000 | 14.603769000 |
| 6 | 5.434239000 | -5.512976000 | 14.782691000 |
| 6 | 3.906425000 | -5.557167000 | 14.937631000 |
| 6 | 6.043483000 | -4.174271000 | 15.214809000 |
| 6 | 3.442746000 | -5.559784000 | 16.397874000 |
| 1 | 5.888215000 | -6.330906000 | 15.360718000 |
| 1 | 3.520419000 | -6.453269000 | 14.435676000 |
| 1 | 3.475276000 | -4.693041000 | 14.413114000 |
| 1 | 5.941843000 | -4.003455000 | 16.287221000 |
| 1 | 5.532245000 | -3.346887000 | 14.710512000 |
| 1 | 7.113420000 | -4.124283000 | 14.975528000 |
| 1 | 2.348464000 | -5.571343000 | 16.465722000 |
| 1 | 3.810008000 | -4.681759000 | 16.936826000 |
| 1 | 3.813736000 | -6.448480000 | 16.924344000 |
| 7 | 7.255614000 | -4.944839000 | 18.686756000 |
| 6 | 8.707119000 | -4.897931000 | 18.538910000 |
| 6 | 9.270507000 | -4.504447000 | 17.153972000 |
| 6 | 9.053104000 | -3.059310000 | 16.831214000 |
| 7 | 9.548870000 | -2.438991000 | 15.687889000 |
| 6 | 8.404505000 | -2.059783000 | 17.508866000 |
| 6 | 9.205414000 | -1.125283000 | 15.714178000 |
| 7 | 8.505582000 | -0.867486000 | 16.809606000 |
| 1 | 6.796357000 | -4.089285000 | 18.380244000 |
| 1 | 9.100459000 | -5.886360000 | 18.797292000 |
| 1 | 8.813561000 | -5.141971000 | 16.383487000 |
| 1 | 10.348462000 | -4.721349000 | 17.135926000 |
| 1 | 7.857244000 | -2.118088000 | 18.432911000 |
| 1 | 9.486676000 | -0.408406000 | 14.957320000 |
| 1 | 10.122363000 | -2.873384000 | 14.976796000 |

| | | | |
|---|---|---|---|
| 7 | 9.567796000 | -1.958447000 | 10.665872000 |
| 6 | 8.763269000 | -1.262938000 | 11.672352000 |
| 6 | 7.307091000 | -1.624755000 | 11.372309000 |
| 8 | 6.711943000 | -1.253018000 | 10.359824000 |
| 6 | 8.999591000 | 0.274019000 | 11.728973000 |
| 6 | 10.336155000 | 0.596533000 | 12.318879000 |
| 7 | 11.519888000 | 0.247941000 | 11.677753000 |
| 6 | 10.624545000 | 1.140132000 | 13.555487000 |
| 6 | 12.491902000 | 0.552582000 | 12.528789000 |
| 7 | 12.001802000 | 1.099863000 | 13.671072000 |
| 1 | 9.286107000 | -1.586991000 | 9.755015000 |
| 1 | 9.018706000 | -1.679488000 | 12.659395000 |
| 1 | 8.923883000 | 0.641873000 | 10.706493000 |
| 1 | 8.224528000 | 0.773468000 | 12.312416000 |
| 1 | 9.992163000 | 1.579927000 | 14.318288000 |
| 1 | 13.550650000 | 0.420443000 | 12.354065000 |
| 1 | 12.540348000 | 1.328026000 | 14.516486000 |
| 7 | 6.671441000 | -2.419092000 | 12.283947000 |
| 6 | 5.399242000 | -3.043252000 | 11.935547000 |
| 6 | 4.199753000 | -2.112798000 | 12.168614000 |
| 8 | 3.589878000 | -2.110183000 | 13.255891000 |
| 1 | 7.193102000 | -2.800894000 | 13.058688000 |
| 1 | 5.437327000 | -3.350609000 | 10.885865000 |
| 7 | 3.887156000 | -1.351398000 | 11.111473000 |
| 6 | 2.852181000 | -0.322296000 | 11.083030000 |
| 6 | 3.420038000 | 1.076240000 | 10.782937000 |
| 6 | 4.205602000 | 1.693231000 | 11.914925000 |
| 6 | 5.354318000 | 1.076789000 | 12.424616000 |
| 6 | 3.841561000 | 2.934060000 | 12.450936000 |
| 6 | 6.107646000 | 1.658727000 | 13.436999000 |
| 6 | 4.599292000 | 3.543215000 | 13.445646000 |
| 6 | 5.734658000 | 2.902424000 | 13.946196000 |
| 8 | 6.461317000 | 3.521826000 | 14.919971000 |
| 1 | 4.584507000 | -1.337940000 | 10.369837000 |
| 1 | 2.345420000 | -0.351817000 | 12.050350000 |
| 1 | 2.575956000 | 1.728165000 | 10.530138000 |
| 1 | 4.047283000 | 1.024744000 | 9.880640000 |
| 1 | 5.688675000 | 0.146339000 | 11.995796000 |
| 1 | 2.978595000 | 3.457723000 | 12.050553000 |
| 1 | 6.982996000 | 1.168608000 | 13.846569000 |
| 1 | 4.342092000 | 4.525599000 | 13.816981000 |
| 1 | 7.198619000 | 2.910035000 | 15.231685000 |
| 6 | 5.080496000 | 7.050543000 | 17.312083000 |
| 6 | 4.634720000 | 6.275358000 | 16.066581000 |
| 6 | 3.307642000 | 5.544648000 | 16.319354000 |
| 6 | 4.531446000 | 7.212472000 | 14.855082000 |
| 1 | 4.310649000 | 7.770633000 | 17.623433000 |
| 1 | 5.288939000 | 6.373756000 | 18.146960000 |
| 1 | 5.400776000 | 5.523457000 | 15.847153000 |
| 1 | 2.962898000 | 5.005154000 | 15.430118000 |
| 1 | 2.516871000 | 6.253331000 | 16.603341000 |
| 1 | 3.415127000 | 4.809992000 | 17.127323000 |
| 1 | 4.169281000 | 6.687257000 | 13.962634000 |
| 1 | 3.840061000 | 8.041969000 | 15.055021000 |
| 1 | 5.514969000 | 7.634814000 | 14.619888000 |
| 6 | 1.432391000 | 4.073844000 | 19.621723000 |
| 7 | 2.405085000 | 2.998776000 | 19.510948000 |
| 6 | 3.734951000 | 3.176857000 | 19.602992000 |
| 7 | 4.269481000 | 4.392275000 | 19.718535000 |
| 7 | 4.533830000 | 2.098410000 | 19.635231000 |
| 1 | 1.582203000 | 4.641882000 | 20.546553000 |
| 1 | 1.482021000 | 4.758773000 | 18.766164000 |
| 1 | 2.100284000 | 2.043703000 | 19.340520000 |
| 1 | 3.711863000 | 5.207877000 | 19.514841000 |
| 1 | 5.305536000 | 4.502062000 | 19.673189000 |
| 1 | 4.094055000 | 1.209150000 | 19.426478000 |
| 1 | 5.485571000 | 2.232295000 | 19.267393000 |
| 26 | 7.688969000 | 1.137174000 | 17.212793000 |
| 6 | 9.035187000 | 3.870971000 | 18.438278000 |
| 6 | 9.301241000 | 4.667983000 | 17.160250000 |

| | | | |
|---|---|---|---|
| 6 | 8.957858000 | 6.155427000 | 17.327379000 |
| 6 | 7.536838000 | 3.759487000 | 18.759414000 |
| 8 | 9.526584000 | 6.805775000 | 18.242432000 |
| 8 | 8.157708000 | 6.660171000 | 16.484229000 |
| 8 | 7.024951000 | 4.634694000 | 19.469751000 |
| 8 | 6.890111000 | 2.760790000 | 18.259364000 |
| 16 | 9.758723000 | 2.164456000 | 18.295076000 |
| 1 | 9.546278000 | 4.341481000 | 19.280827000 |
| 1 | 10.373337000 | 4.600531000 | 16.942371000 |
| 1 | 8.762797000 | 4.229847000 | 16.321695000 |
| 8 | 8.295828000 | 1.842590000 | 15.777432000 |
| 8 | 15.218870000 | 1.004103000 | 18.073660000 |
| 1 | 14.541605000 | 0.760052000 | 17.402107000 |
| 1 | 16.031562000 | 0.547612000 | 17.811851000 |
| 8 | 13.174142000 | 1.246006000 | 16.240122000 |
| 1 | 12.296207000 | 1.093335000 | 16.643345000 |
| 1 | 13.367388000 | 2.152851000 | 16.605962000 |
| 8 | 12.087367000 | 3.662807000 | 19.688537000 |
| 1 | 12.048529000 | 4.644848000 | 19.435901000 |
| 1 | 12.526055000 | 3.654407000 | 20.553339000 |
| 8 | 12.022431000 | 6.155737000 | 18.973023000 |
| 1 | 11.076017000 | 6.395259000 | 18.803881000 |
| 1 | 12.485377000 | 6.376880000 | 18.137698000 |
| 1 | 9.108346000 | -4.201949000 | 19.286484000 |
| 1 | 6.863272000 | -5.694175000 | 18.119853000 |
| 1 | 1.373673000 | -2.703218000 | 17.842054000 |
| 1 | 0.316238000 | 1.337090000 | 16.003091000 |
| 1 | 0.435152000 | 3.632595000 | 19.649654000 |
| 1 | 6.011370000 | 7.590845000 | 17.125041000 |
| 1 | 18.918325000 | 4.875178000 | 11.093133000 |
| 1 | 15.024058000 | 7.127964000 | 8.515505000 |
| 1 | 11.344744000 | 1.201236000 | 6.846822000 |
| 1 | 5.907313000 | 0.638054000 | 7.352645000 |
| 1 | 10.535564000 | -1.655562000 | 10.793489000 |
| 1 | 5.714329000 | -3.929429000 | 20.448976000 |
| 1 | 8.382202000 | 8.359699000 | 10.342705000 |
| 1 | 13.227199000 | 8.640858000 | 8.894877000 |
| 1 | 15.210886000 | 10.115786000 | 13.328360000 |
| 1 | 2.122167000 | -0.595357000 | 10.311349000 |
| 1 | 5.258907000 | -3.924687000 | 12.557191000 |
| 1 | 5.694407000 | -5.712348000 | 13.734341000 |
| 8 | 7.991779000 | 5.656557000 | 14.012618000 |
| 1 | 8.012579000 | 6.005226000 | 14.938362000 |
| 1 | 7.377884000 | 4.900133000 | 14.084572000 |
| 8 | 13.874162000 | 3.381856000 | 17.709061000 |
| 1 | 13.211655000 | 3.371964000 | 18.456195000 |
| 1 | 14.623038000 | 2.824478000 | 18.008439000 |
| 8 | 10.874015000 | 2.210198000 | 17.260816000 |

**$^{5}$TS3$_{S}$:**

| | | | |
|---|---|---|---|
| 6 | 14.889206000 | 9.036758000 | 13.055009000 |
| 6 | 13.786239000 | 9.305459000 | 12.044822000 |
| 8 | 12.693772000 | 9.798969000 | 12.358052000 |
| 6 | 14.350544000 | 8.418468000 | 14.349706000 |
| 6 | 13.582591000 | 7.106795000 | 14.127202000 |
| 6 | 13.259684000 | 6.435799000 | 15.451297000 |
| 8 | 12.938963000 | 7.106637000 | 16.447953000 |
| 7 | 13.324282000 | 5.088253000 | 15.494013000 |
| 1 | 15.654329000 | 8.394197000 | 12.609031000 |
| 1 | 15.192721000 | 8.240520000 | 15.027737000 |
| 1 | 13.693552000 | 9.128024000 | 14.856532000 |
| 1 | 14.159162000 | 6.428454000 | 13.488367000 |
| 1 | 12.644484000 | 7.300607000 | 13.591965000 |
| 1 | 13.353406000 | 4.624886000 | 16.406226000 |
| 1 | 13.767262000 | 4.557422000 | 14.747065000 |
| 7 | 14.071942000 | 8.961429000 | 10.766733000 |
| 6 | 13.018518000 | 9.014702000 | 9.775346000 |
| 6 | 11.826393000 | 8.159632000 | 10.242769000 |
| 8 | 12.024004000 | 7.072230000 | 10.794572000 |
| 1 | 14.931746000 | 8.437066000 | 10.583880000 |
| 1 | 12.709689000 | 10.050545000 | 9.597555000 |
| 7 | 10.596871000 | 8.655405000 | 10.002498000 |
| 6 | 9.430924000 | 8.182914000 | 10.759416000 |
| 6 | 9.451235000 | 8.844517000 | 12.145591000 |
| 6 | 8.124018000 | 8.860080000 | 12.898495000 |
| 6 | 8.207855000 | 9.719596000 | 14.169874000 |
| 7 | 9.020326000 | 9.125460000 | 15.227245000 |
| 6 | 10.276294000 | 9.410515000 | 15.547414000 |
| 7 | 11.060000000 | 10.189623000 | 14.757743000 |
| 7 | 10.778552000 | 8.987736000 | 16.721407000 |
| 1 | 10.561303000 | 9.625513000 | 9.714969000 |
| 1 | 9.476784000 | 7.093746000 | 10.825533000 |
| 1 | 10.219310000 | 8.338156000 | 12.740214000 |
| 1 | 9.786894000 | 9.882964000 | 12.007964000 |
| 1 | 7.340167000 | 9.290051000 | 12.259186000 |
| 1 | 7.819041000 | 7.840121000 | 13.154286000 |
| 1 | 8.609281000 | 10.712741000 | 13.941152000 |
| 1 | 7.206346000 | 9.856078000 | 14.591378000 |
| 1 | 8.585597000 | 8.356873000 | 15.774205000 |
| 1 | 11.962789000 | 10.430076000 | 15.142540000 |
| 1 | 11.096894000 | 10.045210000 | 13.754012000 |
| 1 | 10.166357000 | 8.447228000 | 17.348222000 |
| 1 | 11.739971000 | 8.646677000 | 16.706038000 |
| 7 | 5.890633000 | 1.030945000 | 8.353749000 |
| 6 | 7.043271000 | 1.934200000 | 8.404895000 |
| 6 | 8.366186000 | 1.173957000 | 8.170857000 |
| 8 | 8.475200000 | -0.025978000 | 8.428351000 |
| 6 | 7.144164000 | 2.596646000 | 9.798951000 |
| 6 | 6.206456000 | 3.743770000 | 10.108029000 |
| 6 | 4.919844000 | 3.868166000 | 9.569318000 |
| 6 | 6.627618000 | 4.698322000 | 11.044291000 |
| 6 | 4.064068000 | 4.893569000 | 9.975488000 |
| 6 | 5.780160000 | 5.721001000 | 11.460864000 |
| 6 | 4.487954000 | 5.809028000 | 10.940589000 |
| 8 | 3.595630000 | 6.770044000 | 11.357877000 |
| 1 | 6.175572000 | 0.202692000 | 8.880233000 |
| 1 | 6.923599000 | 2.712925000 | 7.641083000 |
| 1 | 6.970557000 | 1.797334000 | 10.528250000 |
| 1 | 8.168933000 | 2.945168000 | 9.968892000 |
| 1 | 4.578486000 | 3.139158000 | 8.842033000 |
| 1 | 7.634628000 | 4.647664000 | 11.447385000 |
| 1 | 3.063649000 | 4.986265000 | 9.563383000 |
| 1 | 6.127815000 | 6.428590000 | 12.208372000 |
| 1 | 4.050380000 | 7.344428000 | 11.993306000 |
| 7 | 9.414891000 | 1.922406000 | 7.753200000 |
| 6 | 10.797160000 | 1.475159000 | 7.901180000 |
| 6 | 11.557676000 | 2.697604000 | 8.396024000 |
| 8 | 11.691421000 | 3.674149000 | 7.657336000 |
| 1 | 9.288309000 | 2.909842000 | 7.568408000 |
| 1 | 10.811874000 | 0.634788000 | 8.598329000 |
| 7 | 11.943341000 | 2.702921000 | 9.692094000 |
| 6 | 12.408747000 | 3.962514000 | 10.231076000 |
| 6 | 13.851684000 | 4.244075000 | 9.783901000 |
| 8 | 14.724137000 | 3.379792000 | 9.660962000 |
| 6 | 12.229307000 | 4.071439000 | 11.746041000 |
| 6 | 10.782873000 | 4.449096000 | 12.093310000 |
| 8 | 9.900650000 | 4.465477000 | 11.234280000 |
| 7 | 10.562536000 | 4.801769000 | 13.379702000 |
| 1 | 11.832737000 | 1.882284000 | 10.282806000 |
| 1 | 11.782135000 | 4.734293000 | 9.782630000 |
| 1 | 12.882290000 | 4.863486000 | 12.126497000 |
| 1 | 12.515304000 | 3.147954000 | 12.257658000 |
| 1 | 11.272940000 | 4.681465000 | 14.089495000 |
| 1 | 9.605516000 | 5.066349000 | 13.651482000 |
| 7 | 14.064975000 | 5.557481000 | 9.520725000 |
| 6 | 15.314679000 | 6.025381000 | 8.961065000 |
| 6 | 16.356953000 | 6.205882000 | 10.060784000 |
| 8 | 16.458935000 | 7.271075000 | 10.686522000 |
| 1 | 13.367233000 | 6.222673000 | 9.856159000 |
| 1 | 15.650792000 | 5.303851000 | 8.211206000 |

| | | | |
|---|---|---|---|
| 7 | 17.086945000 | 5.096829000 | 10.305913000 |
| 6 | 17.960039000 | 4.952229000 | 11.462148000 |
| 6 | 17.460708000 | 3.848015000 | 12.403800000 |
| 6 | 16.037686000 | 4.122084000 | 12.885974000 |
| 16 | 15.326940000 | 2.825600000 | 13.961893000 |
| 6 | 16.337899000 | 3.053129000 | 15.465705000 |
| 1 | 16.734913000 | 4.248284000 | 9.865845000 |
| 1 | 17.982113000 | 5.922893000 | 11.964102000 |
| 1 | 17.483641000 | 2.879110000 | 11.890300000 |
| 1 | 18.153277000 | 3.777283000 | 13.250362000 |
| 1 | 15.986196000 | 5.087278000 | 13.404749000 |
| 1 | 15.362225000 | 4.161915000 | 12.029077000 |
| 1 | 16.002342000 | 2.318622000 | 16.199755000 |
| 1 | 16.190040000 | 4.053228000 | 15.883986000 |
| 1 | 17.397361000 | 2.887348000 | 15.256170000 |
| 6 | 4.982095000 | -3.554766000 | 20.664302000 |
| 6 | 4.409436000 | -3.207874000 | 19.300150000 |
| 8 | 5.085806000 | -3.244809000 | 18.269774000 |
| 6 | 5.338927000 | -2.279308000 | 21.474522000 |
| 6 | 6.263199000 | -1.363403000 | 20.736107000 |
| 7 | 7.582268000 | -1.125960000 | 21.092073000 |
| 6 | 6.084773000 | -0.658300000 | 19.570865000 |
| 6 | 8.139452000 | -0.305540000 | 20.159741000 |
| 7 | 7.253885000 | -0.017826000 | 19.222983000 |
| 1 | 4.261886000 | -4.148502000 | 21.239348000 |
| 1 | 4.418967000 | -1.737745000 | 21.725422000 |
| 1 | 5.785102000 | -2.585420000 | 22.426869000 |
| 1 | 5.212814000 | -0.599011000 | 18.941717000 |
| 1 | 9.160685000 | 0.044134000 | 20.194426000 |
| 1 | 8.043232000 | -1.480096000 | 21.919009000 |
| 7 | 3.106093000 | -2.806843000 | 19.286552000 |
| 6 | 2.447695000 | -2.396130000 | 18.074271000 |
| 6 | 2.092727000 | -0.905233000 | 18.030449000 |
| 8 | 2.490594000 | -0.103993000 | 18.882920000 |
| 1 | 2.605179000 | -2.731658000 | 20.160415000 |
| 1 | 3.144313000 | -2.600881000 | 17.255995000 |
| 7 | 1.312317000 | -0.581371000 | 16.970367000 |
| 6 | 1.159668000 | 0.776213000 | 16.456006000 |
| 6 | 2.164734000 | 1.078769000 | 15.321049000 |
| 6 | 3.552073000 | 0.683469000 | 15.700737000 |
| 7 | 4.078044000 | -0.558080000 | 15.379337000 |
| 6 | 4.495092000 | 1.283180000 | 16.496302000 |
| 6 | 5.299074000 | -0.661402000 | 15.944208000 |
| 7 | 5.574215000 | 0.433501000 | 16.646712000 |
| 1 | 1.113052000 | -1.328782000 | 16.315358000 |
| 1 | 1.329368000 | 1.454337000 | 17.289417000 |
| 1 | 1.866708000 | 0.548340000 | 14.409636000 |
| 1 | 2.131238000 | 2.145706000 | 15.080621000 |
| 1 | 4.510287000 | 2.275301000 | 16.914661000 |
| 1 | 5.942325000 | -1.518062000 | 15.835810000 |
| 1 | 3.728836000 | -1.200110000 | 14.651331000 |
| 6 | 5.406577000 | -5.643382000 | 14.633965000 |
| 6 | 3.889372000 | -5.652363000 | 14.874355000 |
| 6 | 6.085995000 | -4.358440000 | 15.121276000 |
| 6 | 3.508291000 | -5.724057000 | 16.356437000 |
| 1 | 5.865255000 | -6.513704000 | 15.124876000 |
| 1 | 3.448823000 | -6.508444000 | 14.347837000 |
| 1 | 3.459025000 | -4.749322000 | 14.420244000 |
| 1 | 6.052133000 | -4.261379000 | 16.208075000 |
| 1 | 5.578987000 | -3.481879000 | 14.702838000 |
| 1 | 7.142106000 | -4.325530000 | 14.823866000 |
| 1 | 2.419284000 | -5.713517000 | 16.485538000 |
| 1 | 3.927339000 | -4.886044000 | 16.921101000 |
| 1 | 3.883302000 | -6.649206000 | 16.812489000 |
| 7 | 7.402692000 | -5.162508000 | 18.633149000 |
| 6 | 8.836384000 | -4.996778000 | 18.419022000 |
| 6 | 9.309037000 | -4.567473000 | 17.009287000 |
| 6 | 9.043193000 | -3.123934000 | 16.712371000 |
| 7 | 9.659863000 | -2.431468000 | 15.674121000 |
| 6 | 8.265684000 | -2.182402000 | 17.337111000 |
| 6 | 9.263620000 | -1.132890000 | 15.710894000 |
| 7 | 8.414930000 | -0.953865000 | 16.711939000 |
| 1 | 6.857671000 | -4.344779000 | 18.366734000 |
| 1 | 9.318960000 | -5.949595000 | 18.660507000 |
| 1 | 8.831785000 | -5.212615000 | 16.257889000 |
| 1 | 10.390335000 | -4.750255000 | 16.930671000 |
| 1 | 7.594469000 | -2.302074000 | 18.169638000 |
| 1 | 9.613194000 | -0.372931000 | 15.027257000 |
| 1 | 10.336887000 | -2.812740000 | 15.026565000 |
| 7 | 9.553127000 | -1.802172000 | 10.779612000 |
| 6 | 8.771912000 | -1.103894000 | 11.804210000 |
| 6 | 7.311017000 | -1.443191000 | 11.511543000 |
| 8 | 6.687586000 | -0.955134000 | 10.567693000 |
| 6 | 9.026301000 | 0.428305000 | 11.875114000 |
| 6 | 10.376298000 | 0.723352000 | 12.449545000 |
| 7 | 11.543526000 | 0.362196000 | 11.785894000 |
| 6 | 10.693216000 | 1.245783000 | 13.688258000 |
| 6 | 12.534497000 | 0.641129000 | 12.623997000 |
| 7 | 12.071651000 | 1.181772000 | 13.780182000 |
| 1 | 9.265174000 | -1.424700000 | 9.872465000 |
| 1 | 9.032186000 | -1.535911000 | 12.782225000 |
| 1 | 8.940115000 | 0.817165000 | 10.860330000 |
| 1 | 8.264309000 | 0.928490000 | 12.477078000 |
| 1 | 10.082970000 | 1.680313000 | 14.470826000 |
| 1 | 13.588063000 | 0.496846000 | 12.429522000 |
| 1 | 12.631252000 | 1.395533000 | 14.615285000 |
| 7 | 6.707640000 | -2.350886000 | 12.333258000 |
| 6 | 5.448110000 | -2.966641000 | 11.922581000 |
| 6 | 4.224612000 | -2.094301000 | 12.243926000 |
| 8 | 3.575986000 | -2.264110000 | 13.296202000 |
| 1 | 7.262992000 | -2.828102000 | 13.028409000 |
| 1 | 5.489494000 | -3.169193000 | 10.847345000 |
| 7 | 3.934754000 | -1.185592000 | 11.305233000 |
| 6 | 2.861150000 | -0.199834000 | 11.372673000 |
| 6 | 3.369381000 | 1.231175000 | 11.123522000 |
| 6 | 4.154463000 | 1.837521000 | 12.262751000 |
| 6 | 5.354629000 | 1.265331000 | 12.702729000 |
| 6 | 3.734914000 | 3.022488000 | 12.878691000 |
| 6 | 6.111145000 | 1.845029000 | 13.713126000 |
| 6 | 4.481195000 | 3.617774000 | 13.893042000 |
| 6 | 5.676903000 | 3.029668000 | 14.310805000 |
| 8 | 6.402756000 | 3.644284000 | 15.291540000 |
| 1 | 4.648915000 | -1.059500000 | 10.589638000 |
| 1 | 2.389644000 | -0.299843000 | 12.352754000 |
| 1 | 2.498129000 | 1.860176000 | 10.909688000 |
| 1 | 3.975904000 | 1.231741000 | 10.206662000 |
| 1 | 5.724855000 | 0.376002000 | 12.217119000 |
| 1 | 2.828912000 | 3.512746000 | 12.533176000 |
| 1 | 7.036185000 | 1.395351000 | 14.055518000 |
| 1 | 4.163016000 | 4.547667000 | 14.348449000 |
| 1 | 7.170725000 | 3.054932000 | 15.555158000 |
| 6 | 5.011534000 | 7.554965000 | 17.384586000 |
| 6 | 4.402284000 | 6.711107000 | 16.258539000 |
| 6 | 2.871188000 | 6.657625000 | 16.358350000 |
| 6 | 4.823696000 | 7.251676000 | 14.884831000 |
| 1 | 4.649930000 | 8.590620000 | 17.330132000 |
| 1 | 4.741948000 | 7.165462000 | 18.374560000 |
| 1 | 4.804155000 | 5.692111000 | 16.348722000 |
| 1 | 2.440174000 | 5.960329000 | 15.628451000 |
| 1 | 2.434749000 | 7.646232000 | 16.169121000 |
| 1 | 2.526688000 | 6.358352000 | 17.359429000 |
| 1 | 4.313404000 | 6.719390000 | 14.073047000 |
| 1 | 4.570199000 | 8.317466000 | 14.799150000 |
| 1 | 5.899291000 | 7.132871000 | 14.735685000 |
| 6 | 1.355605000 | 3.798398000 | 18.810478000 |
| 7 | 2.375705000 | 2.838412000 | 19.196945000 |
| 6 | 3.690444000 | 3.127729000 | 19.239983000 |
| 7 | 4.147676000 | 4.333416000 | 18.909675000 |
| 7 | 4.544405000 | 2.186585000 | 19.673153000 |
| 1 | 1.427048000 | 4.701248000 | 19.425505000 |

| | | | |
|---|---|---|---|
| 1 | 1.425279000 | 4.084057000 | 17.752223000 |
| 1 | 2.149223000 | 1.845618000 | 19.213828000 |
| 1 | 3.542483000 | 4.999673000 | 18.453732000 |
| 1 | 5.171369000 | 4.533026000 | 18.880808000 |
| 1 | 4.182189000 | 1.244966000 | 19.751946000 |
| 1 | 5.521741000 | 2.280412000 | 19.355413000 |
| 26 | 7.537132000 | 0.989150000 | 17.235411000 |
| 6 | 8.991875000 | 3.929980000 | 18.111738000 |
| 6 | 9.336005000 | 4.846549000 | 16.944290000 |
| 6 | 8.940818000 | 6.307562000 | 17.200950000 |
| 6 | 7.488585000 | 3.805602000 | 18.408882000 |
| 8 | 9.479409000 | 6.921762000 | 18.157067000 |
| 8 | 8.143712000 | 6.841429000 | 16.370131000 |
| 8 | 6.864102000 | 4.836824000 | 18.674490000 |
| 8 | 6.975682000 | 2.618492000 | 18.421951000 |
| 16 | 9.734944000 | 2.247966000 | 17.870852000 |
| 1 | 9.477258000 | 4.304464000 | 19.020080000 |
| 1 | 10.422889000 | 4.822289000 | 16.810385000 |
| 1 | 8.875072000 | 4.476715000 | 16.031797000 |
| 8 | 8.352537000 | 1.956037000 | 16.006842000 |
| 8 | 15.262864000 | 1.101493000 | 18.261568000 |
| 1 | 14.607314000 | 0.830033000 | 17.579696000 |
| 1 | 16.092668000 | 0.664165000 | 18.021604000 |
| 8 | 13.294729000 | 1.284726000 | 16.338219000 |
| 1 | 12.378644000 | 1.192078000 | 16.671994000 |
| 1 | 13.491710000 | 2.197240000 | 16.687596000 |
| 8 | 11.931105000 | 3.657982000 | 19.610580000 |
| 1 | 11.907345000 | 4.640097000 | 19.355991000 |
| 1 | 12.285811000 | 3.656479000 | 20.513338000 |
| 8 | 11.927907000 | 6.161028000 | 18.920963000 |
| 1 | 10.995054000 | 6.444072000 | 18.736376000 |
| 1 | 12.420216000 | 6.395351000 | 18.106567000 |
| 1 | 9.212031000 | -4.265683000 | 19.146047000 |
| 1 | 7.045930000 | -5.936962000 | 18.076750000 |
| 1 | 1.545210000 | -2.995622000 | 17.900595000 |
| 1 | 0.130146000 | 0.913231000 | 16.110855000 |
| 1 | 0.376767000 | 3.349328000 | 18.985818000 |
| 1 | 6.102349000 | 7.555363000 | 17.317632000 |
| 1 | 18.978887000 | 4.724207000 | 11.127918000 |
| 1 | 15.153861000 | 6.988357000 | 8.471260000 |
| 1 | 11.217768000 | 1.151950000 | 6.942881000 |
| 1 | 5.767756000 | 0.710346000 | 7.393031000 |
| 1 | 10.527937000 | -1.518971000 | 10.897766000 |
| 1 | 5.872298000 | -4.166060000 | 20.492115000 |
| 1 | 8.532055000 | 8.454825000 | 10.197121000 |
| 1 | 13.405696000 | 8.613894000 | 8.833387000 |
| 1 | 15.369588000 | 9.995371000 | 13.289706000 |
| 1 | 2.111170000 | -0.456689000 | 10.614122000 |
| 1 | 5.328823000 | -3.906756000 | 12.456714000 |
| 1 | 5.598014000 | -5.773999000 | 13.560258000 |
| 8 | 7.921279000 | 5.698018000 | 13.984438000 |
| 1 | 7.977979000 | 6.123328000 | 14.880231000 |
| 1 | 7.297650000 | 4.966554000 | 14.142363000 |
| 8 | 13.903143000 | 3.458523000 | 17.791760000 |
| 1 | 13.187093000 | 3.416992000 | 18.485247000 |
| 1 | 14.644515000 | 2.921032000 | 18.141620000 |
| 8 | 10.935612000 | 2.374921000 | 16.960562000 |

**$^{5}$Prod$_{S}$:**

| | | | |
|---|---|---|---|
| 6 | 15.130131000 | 8.730492000 | 13.173785000 |
| 6 | 14.207520000 | 9.131086000 | 12.035330000 |
| 8 | 13.182805000 | 9.802427000 | 12.215898000 |
| 6 | 14.330833000 | 7.992426000 | 14.254645000 |
| 6 | 13.655708000 | 6.716748000 | 13.727872000 |
| 6 | 12.718866000 | 6.145342000 | 14.770154000 |
| 8 | 11.859737000 | 6.876139000 | 15.302011000 |
| 7 | 12.859184000 | 4.855352000 | 15.111286000 |
| 1 | 15.949509000 | 8.107488000 | 12.804580000 |
| 1 | 14.991804000 | 7.745050000 | 15.094097000 |
| 1 | 13.548245000 | 8.649965000 | 14.639251000 |
| 1 | 14.402989000 | 5.985837000 | 13.405394000 |
| 1 | 13.047011000 | 6.951185000 | 12.846179000 |
| 1 | 12.158776000 | 4.417019000 | 15.709287000 |
| 1 | 13.524296000 | 4.227031000 | 14.669533000 |
| 7 | 14.559972000 | 8.671902000 | 10.807529000 |
| 6 | 13.618199000 | 8.817426000 | 9.715067000 |
| 6 | 12.295375000 | 8.127942000 | 10.080751000 |
| 8 | 12.312306000 | 7.008965000 | 10.607511000 |
| 1 | 15.269991000 | 7.937727000 | 10.759917000 |
| 1 | 13.464643000 | 9.876935000 | 9.487616000 |
| 7 | 11.158120000 | 8.789366000 | 9.805148000 |
| 6 | 9.898444000 | 8.412802000 | 10.456536000 |
| 6 | 9.878643000 | 8.990148000 | 11.879375000 |
| 6 | 8.494496000 | 9.076172000 | 12.517998000 |
| 6 | 8.503019000 | 9.948667000 | 13.780183000 |
| 7 | 9.162054000 | 9.338538000 | 14.933215000 |
| 6 | 10.380262000 | 9.588653000 | 15.391116000 |
| 7 | 11.301820000 | 10.250645000 | 14.645874000 |
| 7 | 10.684636000 | 9.283027000 | 16.669163000 |
| 1 | 11.259449000 | 9.769651000 | 9.573302000 |
| 1 | 9.820746000 | 7.323584000 | 10.463599000 |
| 1 | 10.551521000 | 8.388930000 | 12.502070000 |
| 1 | 10.306024000 | 10.002459000 | 11.828005000 |
| 1 | 7.793192000 | 9.539732000 | 11.809412000 |
| 1 | 8.106751000 | 8.079398000 | 12.754362000 |
| 1 | 8.984099000 | 10.912053000 | 13.578722000 |
| 1 | 7.474324000 | 10.153001000 | 14.095796000 |
| 1 | 8.583189000 | 8.685516000 | 15.495788000 |
| 1 | 12.170325000 | 10.473359000 | 15.111578000 |
| 1 | 11.430724000 | 10.028554000 | 13.663663000 |
| 1 | 9.972996000 | 8.787277000 | 17.225877000 |
| 1 | 11.615755000 | 8.917313000 | 16.831352000 |
| 7 | 5.572884000 | 1.388527000 | 8.500155000 |
| 6 | 6.686535000 | 2.334146000 | 8.380233000 |
| 6 | 7.999843000 | 1.596466000 | 8.066563000 |
| 8 | 8.152295000 | 0.416498000 | 8.387144000 |
| 6 | 6.901859000 | 3.101042000 | 9.706334000 |
| 6 | 5.924057000 | 4.211002000 | 10.035223000 |
| 6 | 4.606341000 | 4.252653000 | 9.563747000 |
| 6 | 6.342718000 | 5.219968000 | 10.914019000 |
| 6 | 3.713222000 | 5.234638000 | 9.995642000 |
| 6 | 5.459518000 | 6.203595000 | 11.352749000 |
| 6 | 4.133085000 | 6.196615000 | 10.915852000 |
| 8 | 3.207585000 | 7.105621000 | 11.369490000 |
| 1 | 5.941111000 | 0.576099000 | 9.003593000 |
| 1 | 6.463100000 | 3.053024000 | 7.582268000 |
| 1 | 6.863729000 | 2.354434000 | 10.510352000 |
| 1 | 7.916010000 | 3.516095000 | 9.737934000 |
| 1 | 4.265298000 | 3.489866000 | 8.873095000 |
| 1 | 7.375235000 | 5.242576000 | 11.251073000 |
| 1 | 2.688434000 | 5.258141000 | 9.637282000 |
| 1 | 5.808131000 | 6.962908000 | 12.048033000 |
| 1 | 3.657085000 | 7.720631000 | 11.969989000 |
| 7 | 8.988768000 | 2.328558000 | 7.496336000 |
| 6 | 10.386832000 | 1.920855000 | 7.610626000 |
| 6 | 11.098796000 | 3.058259000 | 8.336451000 |
| 8 | 11.248150000 | 4.138773000 | 7.767856000 |
| 1 | 8.840107000 | 3.318681000 | 7.338900000 |
| 1 | 10.409494000 | 0.969767000 | 8.144953000 |
| 7 | 11.430049000 | 2.832648000 | 9.632409000 |
| 6 | 12.045742000 | 3.910393000 | 10.385896000 |
| 6 | 13.567892000 | 3.855218000 | 10.181698000 |
| 8 | 14.249243000 | 2.836686000 | 10.353894000 |
| 6 | 11.642361000 | 3.933121000 | 11.856731000 |
| 6 | 10.228122000 | 4.498109000 | 12.041265000 |
| 8 | 9.623525000 | 5.033900000 | 11.107633000 |
| 7 | 9.709710000 | 4.399815000 | 13.285124000 |
| 1 | 11.401423000 | 1.892412000 | 10.023258000 |
| 1 | 11.666802000 | 4.828230000 | 9.940947000 |
| 1 | 12.324701000 | 4.602770000 | 12.390568000 |

| | | | |
|---|---|---|---|
| 1 | 11.735897000 | 2.956715000 | 12.334053000 |
| 1 | 10.227430000 | 3.999302000 | 14.059233000 |
| 1 | 8.871695000 | 4.952792000 | 13.496667000 |
| 7 | 14.080728000 | 5.025041000 | 9.732403000 |
| 6 | 15.451905000 | 5.152003000 | 9.295567000 |
| 6 | 16.404082000 | 5.345739000 | 10.473425000 |
| 8 | 16.450676000 | 6.417172000 | 11.092028000 |
| 1 | 13.499277000 | 5.856378000 | 9.842622000 |
| 1 | 15.717012000 | 4.266862000 | 8.710185000 |
| 7 | 17.150413000 | 4.259576000 | 10.774432000 |
| 6 | 17.955529000 | 4.150453000 | 11.984696000 |
| 6 | 17.357597000 | 3.132688000 | 12.966259000 |
| 6 | 15.895914000 | 3.457779000 | 13.267700000 |
| 16 | 15.090134000 | 2.372011000 | 14.493208000 |
| 6 | 15.916460000 | 2.930459000 | 16.025407000 |
| 1 | 16.861166000 | 3.395904000 | 10.329259000 |
| 1 | 17.984373000 | 5.149662000 | 12.425459000 |
| 1 | 17.426181000 | 2.118666000 | 12.552805000 |
| 1 | 17.959331000 | 3.142596000 | 13.882375000 |
| 1 | 15.808479000 | 4.496457000 | 13.607779000 |
| 1 | 15.291664000 | 3.346535000 | 12.363375000 |
| 1 | 15.424450000 | 2.438235000 | 16.867747000 |
| 1 | 15.799855000 | 4.013236000 | 16.139039000 |
| 1 | 16.977926000 | 2.670550000 | 16.017928000 |
| 6 | 4.953144000 | -3.960226000 | 20.330986000 |
| 6 | 4.583795000 | -3.516144000 | 18.922817000 |
| 8 | 5.417030000 | -3.395934000 | 18.021128000 |
| 6 | 4.969570000 | -2.769963000 | 21.326324000 |
| 6 | 5.899100000 | -1.673764000 | 20.909909000 |
| 7 | 7.107207000 | -1.396661000 | 21.533945000 |
| 6 | 5.842001000 | -0.798526000 | 19.852839000 |
| 6 | 7.722382000 | -0.388437000 | 20.861186000 |
| 7 | 6.980549000 | -0.012593000 | 19.831715000 |
| 1 | 4.232334000 | -4.704751000 | 20.689379000 |
| 1 | 3.958391000 | -2.357941000 | 21.425320000 |
| 1 | 5.247243000 | -3.146771000 | 22.316671000 |
| 1 | 5.071501000 | -0.699572000 | 19.104110000 |
| 1 | 8.684646000 | 0.021018000 | 21.131417000 |
| 1 | 7.461323000 | -1.855259000 | 22.362672000 |
| 7 | 3.263974000 | -3.235609000 | 18.729972000 |
| 6 | 2.758206000 | -2.702271000 | 17.491649000 |
| 6 | 2.342700000 | -1.227830000 | 17.590436000 |
| 8 | 2.613839000 | -0.536984000 | 18.578558000 |
| 1 | 2.641692000 | -3.260363000 | 19.525394000 |
| 1 | 3.564282000 | -2.789351000 | 16.757889000 |
| 7 | 1.657158000 | -0.790309000 | 16.506433000 |
| 6 | 1.430755000 | 0.628007000 | 16.256114000 |
| 6 | 2.560572000 | 1.258802000 | 15.411225000 |
| 6 | 3.911188000 | 0.877924000 | 15.910666000 |
| 7 | 4.474887000 | -0.348718000 | 15.604289000 |
| 6 | 4.799389000 | 1.480401000 | 16.762296000 |
| 6 | 5.660141000 | -0.449740000 | 16.241193000 |
| 7 | 5.880975000 | 0.645880000 | 16.960241000 |
| 1 | 1.641185000 | -1.395598000 | 15.691124000 |
| 1 | 1.366967000 | 1.107948000 | 17.230719000 |
| 1 | 2.462971000 | 0.934613000 | 14.372669000 |
| 1 | 2.457500000 | 2.347627000 | 15.410444000 |
| 1 | 4.787646000 | 2.480390000 | 17.157477000 |
| 1 | 6.305156000 | -1.311179000 | 16.200973000 |
| 1 | 4.086357000 | -0.984368000 | 14.894708000 |
| 6 | 7.289404000 | -4.989531000 | 14.342081000 |
| 6 | 6.106642000 | -5.947137000 | 14.545204000 |
| 6 | 7.170733000 | -3.696111000 | 15.153592000 |
| 6 | 5.928664000 | -6.422637000 | 15.990531000 |
| 1 | 8.223360000 | -5.502545000 | 14.611942000 |
| 1 | 6.238507000 | -6.815065000 | 13.887825000 |
| 1 | 5.182432000 | -5.451061000 | 14.213285000 |
| 1 | 7.227220000 | -3.871010000 | 16.228514000 |
| 1 | 6.199986000 | -3.214081000 | 14.978310000 |
| 1 | 7.967569000 | -2.984217000 | 14.901368000 |
| 1 | 5.117509000 | -7.154742000 | 16.066521000 |
| 1 | 5.693794000 | -5.591642000 | 16.663522000 |
| 1 | 6.843794000 | -6.910618000 | 16.352378000 |
| 7 | 7.865631000 | -5.036646000 | 18.885158000 |
| 6 | 9.302096000 | -4.795491000 | 18.985016000 |
| 6 | 10.005496000 | -4.220736000 | 17.735681000 |
| 6 | 9.617690000 | -2.800252000 | 17.470669000 |
| 7 | 10.142275000 | -2.036219000 | 16.434553000 |
| 6 | 8.730625000 | -1.964936000 | 18.098242000 |
| 6 | 9.584115000 | -0.803326000 | 16.470505000 |
| 7 | 8.718767000 | -0.730520000 | 17.470999000 |
| 1 | 7.352473000 | -4.243530000 | 18.504073000 |
| 1 | 9.782805000 | -5.743156000 | 19.246946000 |
| 1 | 9.766893000 | -4.847924000 | 16.864151000 |
| 1 | 11.094314000 | -4.284831000 | 17.875983000 |
| 1 | 8.085449000 | -2.170569000 | 18.933957000 |
| 1 | 9.820547000 | -0.011549000 | 15.783011000 |
| 1 | 10.846168000 | -2.329442000 | 15.770472000 |
| 7 | 9.016346000 | -1.801829000 | 10.342624000 |
| 6 | 8.370003000 | -1.155729000 | 11.480704000 |
| 6 | 6.868097000 | -1.411992000 | 11.341744000 |
| 8 | 6.210019000 | -1.090622000 | 10.348893000 |
| 6 | 8.698620000 | 0.359833000 | 11.638841000 |
| 6 | 10.092688000 | 0.546963000 | 12.144416000 |
| 7 | 11.212074000 | 0.269827000 | 11.371867000 |
| 6 | 10.490739000 | 0.894706000 | 13.416998000 |
| 6 | 12.257699000 | 0.436818000 | 12.173914000 |
| 7 | 11.871718000 | 0.814520000 | 13.420098000 |
| 1 | 8.672332000 | -1.338087000 | 9.499474000 |
| 1 | 8.713527000 | -1.657712000 | 12.397571000 |
| 1 | 8.562877000 | 0.826536000 | 10.662579000 |
| 1 | 8.009365000 | 0.841154000 | 12.335404000 |
| 1 | 9.938490000 | 1.296165000 | 14.252917000 |
| 1 | 13.293974000 | 0.346987000 | 11.881707000 |
| 1 | 12.451785000 | 0.959855000 | 14.251484000 |
| 7 | 6.269697000 | -2.035236000 | 12.398626000 |
| 6 | 4.930989000 | -2.606387000 | 12.287958000 |
| 6 | 3.797884000 | -1.587696000 | 12.491825000 |
| 8 | 3.142757000 | -1.563055000 | 13.557697000 |
| 1 | 6.851811000 | -2.359617000 | 13.154934000 |
| 1 | 4.827136000 | -3.072831000 | 11.302033000 |
| 7 | 3.589854000 | -0.782239000 | 11.447972000 |
| 6 | 2.647693000 | 0.331891000 | 11.407044000 |
| 6 | 3.347179000 | 1.680015000 | 11.153792000 |
| 6 | 4.195771000 | 2.185881000 | 12.296642000 |
| 6 | 5.365433000 | 1.519095000 | 12.688204000 |
| 6 | 3.855216000 | 3.353291000 | 12.989526000 |
| 6 | 6.151188000 | 1.973872000 | 13.742971000 |
| 6 | 4.631811000 | 3.824197000 | 14.045839000 |
| 6 | 5.779941000 | 3.127706000 | 14.437278000 |
| 8 | 6.475633000 | 3.588732000 | 15.508495000 |
| 1 | 4.285008000 | -0.849394000 | 10.701870000 |
| 1 | 2.095868000 | 0.324934000 | 12.348680000 |
| 1 | 2.568213000 | 2.416954000 | 10.931622000 |
| 1 | 3.953772000 | 1.592439000 | 10.242092000 |
| 1 | 5.675920000 | 0.639868000 | 12.142447000 |
| 1 | 2.981122000 | 3.920546000 | 12.683218000 |
| 1 | 7.039668000 | 1.435061000 | 14.055977000 |
| 1 | 4.355890000 | 4.725550000 | 14.579398000 |
| 1 | 7.258966000 | 3.010333000 | 15.703771000 |
| 6 | 4.681857000 | 7.734355000 | 17.484372000 |
| 6 | 4.229518000 | 6.827664000 | 16.333902000 |
| 6 | 2.759352000 | 6.409468000 | 16.487360000 |
| 6 | 4.433926000 | 7.522038000 | 14.978918000 |
| 1 | 4.079698000 | 8.651985000 | 17.516767000 |
| 1 | 4.575922000 | 7.232192000 | 18.454352000 |
| 1 | 4.864683000 | 5.930123000 | 16.357524000 |
| 1 | 2.474005000 | 5.639772000 | 15.758624000 |
| 1 | 2.095237000 | 7.269182000 | 16.335513000 |
| 1 | 2.535384000 | 6.031062000 | 17.495419000 |

| | | | |
|---|---|---|---|
| 1 | 4.061402000 | 6.908785000 | 14.148269000 |
| 1 | 3.888154000 | 8.474740000 | 14.958637000 |
| 1 | 5.492754000 | 7.727355000 | 14.804033000 |
| 6 | 1.603945000 | 3.237301000 | 18.821846000 |
| 7 | 2.740793000 | 2.461514000 | 19.294400000 |
| 6 | 3.968756000 | 3.001297000 | 19.429576000 |
| 7 | 4.220947000 | 4.257781000 | 19.056459000 |
| 7 | 4.946608000 | 2.287312000 | 19.998090000 |
| 1 | 1.514280000 | 4.160088000 | 19.401967000 |
| 1 | 1.676264000 | 3.493777000 | 17.755698000 |
| 1 | 2.687929000 | 1.446237000 | 19.202869000 |
| 1 | 3.594863000 | 4.721305000 | 18.413298000 |
| 1 | 5.209115000 | 4.581638000 | 19.040451000 |
| 1 | 4.762113000 | 1.330568000 | 20.262062000 |
| 1 | 5.927834000 | 2.527950000 | 19.790887000 |
| 26 | 7.653599000 | 1.037591000 | 18.015421000 |
| 6 | 9.010756000 | 4.333549000 | 17.767824000 |
| 6 | 8.848905000 | 5.309499000 | 16.618124000 |
| 6 | 8.541087000 | 6.752964000 | 16.998771000 |
| 6 | 7.701045000 | 4.009339000 | 18.492765000 |
| 8 | 9.153348000 | 7.286778000 | 17.960548000 |
| 8 | 7.761059000 | 7.374382000 | 16.205913000 |
| 8 | 6.863169000 | 4.903013000 | 18.623681000 |
| 8 | 7.548800000 | 2.801796000 | 18.977600000 |
| 16 | 9.795343000 | 2.762421000 | 17.157328000 |
| 1 | 9.727063000 | 4.706948000 | 18.505895000 |
| 1 | 9.794581000 | 5.362170000 | 16.067553000 |
| 1 | 8.089611000 | 4.955057000 | 15.929704000 |
| 8 | 8.550115000 | 2.086799000 | 16.418681000 |
| 8 | 11.874638000 | -0.132602000 | 18.635130000 |
| 1 | 11.868641000 | 0.189616000 | 17.702323000 |
| 1 | 12.439008000 | -0.920568000 | 18.604142000 |
| 8 | 12.427224000 | 1.062957000 | 16.168076000 |
| 1 | 11.829495000 | 1.850695000 | 16.102703000 |
| 1 | 13.132662000 | 1.415302000 | 16.776707000 |
| 8 | 12.061983000 | 4.090903000 | 18.937639000 |
| 1 | 12.033291000 | 5.014951000 | 18.541374000 |
| 1 | 12.190648000 | 4.243230000 | 19.886200000 |
| 8 | 11.753794000 | 6.555430000 | 18.018108000 |
| 1 | 10.801305000 | 6.769856000 | 18.191826000 |
| 1 | 11.798331000 | 6.583510000 | 17.034704000 |
| 1 | 9.484930000 | -4.114789000 | 19.826539000 |
| 1 | 7.675361000 | -5.819149000 | 18.263388000 |
| 1 | 1.912175000 | -3.299787000 | 17.131849000 |
| 1 | 0.466565000 | 0.748471000 | 15.753549000 |
| 1 | 0.698236000 | 2.649452000 | 18.978302000 |
| 1 | 5.734256000 | 8.003529000 | 17.362761000 |
| 1 | 18.980228000 | 3.867809000 | 11.718131000 |
| 1 | 15.531763000 | 6.030266000 | 8.648983000 |
| 1 | 10.839055000 | 1.794851000 | 6.622038000 |
| 1 | 5.343856000 | 1.035252000 | 7.570824000 |
| 1 | 10.011809000 | -1.578883000 | 10.390890000 |
| 1 | 5.937469000 | -4.433792000 | 20.276663000 |
| 1 | 9.077970000 | 8.817341000 | 9.855032000 |
| 1 | 14.041218000 | 8.337807000 | 8.826883000 |
| 1 | 15.563344000 | 9.644050000 | 13.598360000 |
| 1 | 1.931185000 | 0.141396000 | 10.599021000 |
| 1 | 4.819406000 | -3.373205000 | 13.054962000 |
| 1 | 7.376695000 | -4.751145000 | 13.273720000 |
| 8 | 7.437019000 | 6.146389000 | 13.815956000 |
| 1 | 7.479569000 | 6.548048000 | 14.723899000 |
| 1 | 6.648755000 | 5.582095000 | 13.806690000 |
| 8 | 13.735642000 | 2.005365000 | 18.315409000 |
| 1 | 13.223443000 | 2.839029000 | 18.470024000 |
| 1 | 13.212796000 | 1.323196000 | 18.785509000 |
| 8 | 10.766313000 | 3.214846000 | 16.050449000 |

**$^{7}Re_{S}$:**

| | | | |
|---|---|---|---|
| 6 | 14.925014000 | 8.791916000 | 13.451910000 |
| 6 | 13.767286000 | 9.242934000 | 12.574076000 |
| 8 | 12.706639000 | 9.693594000 | 13.028229000 |
| 6 | 14.491379000 | 8.317122000 | 14.839131000 |
| 6 | 13.577345000 | 7.086922000 | 14.791058000 |
| 6 | 13.429204000 | 6.429623000 | 16.155353000 |
| 8 | 13.474075000 | 7.109030000 | 17.201444000 |
| 7 | 13.242431000 | 5.104693000 | 16.148423000 |
| 1 | 15.489884000 | 8.012170000 | 12.927685000 |
| 1 | 15.386813000 | 8.083976000 | 15.425058000 |
| 1 | 13.983600000 | 9.128726000 | 15.367623000 |
| 1 | 13.952920000 | 6.353445000 | 14.068865000 |
| 1 | 12.582909000 | 7.375237000 | 14.429649000 |
| 1 | 13.208611000 | 4.534400000 | 16.997241000 |
| 1 | 13.281149000 | 4.594612000 | 15.269560000 |
| 7 | 13.994936000 | 9.154605000 | 11.239440000 |
| 6 | 12.905290000 | 9.441452000 | 10.332229000 |
| 6 | 11.726991000 | 8.496590000 | 10.613334000 |
| 8 | 11.936123000 | 7.313621000 | 10.900312000 |
| 1 | 14.769098000 | 8.566714000 | 10.918776000 |
| 1 | 12.599105000 | 10.488087000 | 10.429334000 |
| 7 | 10.489248000 | 9.022463000 | 10.507370000 |
| 6 | 9.345081000 | 8.337419000 | 11.117109000 |
| 6 | 9.373398000 | 8.599962000 | 12.625673000 |
| 6 | 8.145577000 | 8.140578000 | 13.405311000 |
| 6 | 8.149891000 | 8.705004000 | 14.832290000 |
| 7 | 9.167901000 | 8.130343000 | 15.707859000 |
| 6 | 10.316707000 | 8.691185000 | 16.084459000 |
| 7 | 10.875837000 | 9.710975000 | 15.376824000 |
| 7 | 10.899893000 | 8.292790000 | 17.219092000 |
| 1 | 10.435443000 | 10.032777000 | 10.469216000 |
| 1 | 9.420782000 | 7.271202000 | 10.894894000 |
| 1 | 10.273342000 | 8.117781000 | 13.023429000 |
| 1 | 9.493300000 | 9.684607000 | 12.770753000 |
| 1 | 7.236155000 | 8.508504000 | 12.909684000 |
| 1 | 8.081843000 | 7.046333000 | 13.433082000 |
| 1 | 8.290140000 | 9.791161000 | 14.809451000 |
| 1 | 7.184962000 | 8.506846000 | 15.311091000 |
| 1 | 8.937004000 | 7.223440000 | 16.161222000 |
| 1 | 11.726324000 | 10.094354000 | 15.768103000 |
| 1 | 10.928802000 | 9.637600000 | 14.366456000 |
| 1 | 10.442063000 | 7.541410000 | 17.763548000 |
| 1 | 11.919368000 | 8.272185000 | 17.267014000 |
| 7 | 5.926388000 | 1.640801000 | 8.305571000 |
| 6 | 6.764520000 | 2.830465000 | 8.134340000 |
| 6 | 8.123678000 | 2.519477000 | 7.504004000 |
| 8 | 8.863502000 | 1.640018000 | 7.960261000 |
| 6 | 7.054264000 | 3.508850000 | 9.487999000 |
| 6 | 5.973868000 | 4.423166000 | 10.023373000 |
| 6 | 4.635159000 | 4.357564000 | 9.615022000 |
| 6 | 6.321469000 | 5.361096000 | 11.006828000 |
| 6 | 3.666799000 | 5.198124000 | 10.165618000 |
| 6 | 5.359977000 | 6.199710000 | 11.567286000 |
| 6 | 4.030387000 | 6.122435000 | 11.145381000 |
| 8 | 3.043036000 | 6.919987000 | 11.675438000 |
| 1 | 6.471551000 | 0.903100000 | 8.759283000 |
| 1 | 6.222964000 | 3.534556000 | 7.489353000 |
| 1 | 7.253583000 | 2.712250000 | 10.216433000 |
| 1 | 7.985381000 | 4.081165000 | 9.427180000 |
| 1 | 4.339367000 | 3.613643000 | 8.882185000 |
| 1 | 7.355136000 | 5.420906000 | 11.335504000 |
| 1 | 2.630192000 | 5.146447000 | 9.847715000 |
| 1 | 5.645393000 | 6.913845000 | 12.336868000 |
| 1 | 3.456898000 | 7.549096000 | 12.286430000 |
| 7 | 8.498060000 | 3.287140000 | 6.449073000 |
| 6 | 9.867809000 | 3.252011000 | 5.940641000 |
| 6 | 10.777844000 | 4.149844000 | 6.788240000 |
| 8 | 11.159837000 | 5.246799000 | 6.392881000 |
| 1 | 7.936454000 | 4.096543000 | 6.221052000 |
| 1 | 10.201543000 | 2.212085000 | 5.961200000 |
| 7 | 11.066172000 | 3.669557000 | 8.025410000 |
| 6 | 11.712740000 | 4.545670000 | 8.981083000 |

| | | | |
|---|---|---|---|
| 6 | 13.155277000 | 4.865571000 | 8.559089000 |
| 8 | 13.934593000 | 4.051291000 | 8.063273000 |
| 6 | 11.690776000 | 3.948592000 | 10.391735000 |
| 6 | 10.487980000 | 4.430030000 | 11.208196000 |
| 8 | 9.580858000 | 5.099051000 | 10.701404000 |
| 7 | 10.518356000 | 4.101938000 | 12.514725000 |
| 1 | 10.671377000 | 2.783003000 | 8.321037000 |
| 1 | 11.157132000 | 5.486599000 | 9.004463000 |
| 1 | 12.595425000 | 4.238694000 | 10.937185000 |
| 1 | 11.694983000 | 2.853060000 | 10.354287000 |
| 1 | 11.178907000 | 3.410026000 | 12.842715000 |
| 1 | 9.704674000 | 4.321304000 | 13.107492000 |
| 7 | 13.495348000 | 6.152945000 | 8.836820000 |
| 6 | 14.822942000 | 6.658607000 | 8.560588000 |
| 6 | 15.773587000 | 6.322788000 | 9.706808000 |
| 8 | 16.032576000 | 7.132426000 | 10.609655000 |
| 1 | 12.914993000 | 6.657102000 | 9.507707000 |
| 1 | 15.167715000 | 6.214332000 | 7.623439000 |
| 7 | 16.243963000 | 5.057538000 | 9.658760000 |
| 6 | 17.071099000 | 4.472051000 | 10.700420000 |
| 6 | 16.282922000 | 3.554115000 | 11.644358000 |
| 6 | 15.251460000 | 4.311291000 | 12.479432000 |
| 16 | 14.284879000 | 3.216830000 | 13.589508000 |
| 6 | 15.521644000 | 2.879503000 | 14.889357000 |
| 1 | 15.810470000 | 4.449663000 | 8.963001000 |
| 1 | 17.522831000 | 5.300268000 | 11.252193000 |
| 1 | 15.774153000 | 2.776525000 | 11.061967000 |
| 1 | 16.997651000 | 3.044301000 | 12.301867000 |
| 1 | 15.725844000 | 5.095100000 | 13.080206000 |
| 1 | 14.516068000 | 4.802955000 | 11.835897000 |
| 1 | 15.053316000 | 2.208209000 | 15.613316000 |
| 1 | 15.805965000 | 3.803079000 | 15.401817000 |
| 1 | 16.408373000 | 2.394687000 | 14.473024000 |
| 6 | 3.038179000 | -1.852655000 | 21.821485000 |
| 6 | 2.596459000 | -1.763205000 | 20.365589000 |
| 8 | 3.273925000 | -2.162630000 | 19.417190000 |
| 6 | 3.562380000 | -0.520844000 | 22.409685000 |
| 6 | 4.873699000 | -0.084594000 | 21.833116000 |
| 7 | 5.760156000 | 0.763277000 | 22.486040000 |
| 6 | 5.474900000 | -0.358790000 | 20.633520000 |
| 6 | 6.838574000 | 0.964484000 | 21.684170000 |
| 7 | 6.692596000 | 0.291135000 | 20.556124000 |
| 1 | 2.182463000 | -2.173507000 | 22.426739000 |
| 1 | 2.812617000 | 0.271391000 | 22.269058000 |
| 1 | 3.667363000 | -0.654079000 | 23.493410000 |
| 1 | 5.104699000 | -0.969317000 | 19.830815000 |
| 1 | 7.681302000 | 1.588792000 | 21.941445000 |
| 1 | 5.625936000 | 1.174439000 | 23.399566000 |
| 7 | 1.357249000 | -1.223592000 | 20.164332000 |
| 6 | 0.890242000 | -0.952649000 | 18.828959000 |
| 6 | 1.546821000 | 0.295994000 | 18.216952000 |
| 8 | 2.060480000 | 1.158080000 | 18.940047000 |
| 1 | 0.889863000 | -0.770857000 | 20.936487000 |
| 1 | 1.098613000 | -1.825488000 | 18.204239000 |
| 7 | 1.462712000 | 0.378736000 | 16.875206000 |
| 6 | 2.098642000 | 1.412815000 | 16.060594000 |
| 6 | 3.302887000 | 0.858010000 | 15.280619000 |
| 6 | 4.346670000 | 0.284802000 | 16.180427000 |
| 7 | 4.502986000 | -1.071965000 | 16.420553000 |
| 6 | 5.312011000 | 0.888971000 | 16.943191000 |
| 6 | 5.543789000 | -1.247037000 | 17.272620000 |
| 7 | 6.046634000 | -0.070839000 | 17.612243000 |
| 1 | 1.043965000 | -0.401040000 | 16.383705000 |
| 1 | 2.421323000 | 2.203803000 | 16.738207000 |
| 1 | 2.972932000 | 0.103166000 | 14.557469000 |
| 1 | 3.742173000 | 1.664812000 | 14.685988000 |
| 1 | 5.559531000 | 1.931335000 | 17.028325000 |
| 1 | 5.885484000 | -2.209376000 | 17.618175000 |
| 1 | 3.981291000 | -1.815302000 | 15.978766000 |
| 6 | 9.221940000 | -3.872586000 | 13.440588000 |
| 6 | 7.765841000 | -3.396569000 | 13.538389000 |
| 6 | 10.240636000 | -2.905102000 | 14.050328000 |
| 6 | 7.252094000 | -3.262088000 | 14.973443000 |
| 1 | 9.312766000 | -4.861931000 | 13.912761000 |
| 1 | 7.104470000 | -4.083109000 | 12.998294000 |
| 1 | 7.665738000 | -2.426322000 | 13.036767000 |
| 1 | 10.091736000 | -2.790805000 | 15.130067000 |
| 1 | 10.155857000 | -1.906969000 | 13.607232000 |
| 1 | 11.265227000 | -3.259609000 | 13.893720000 |
| 1 | 6.179648000 | -3.039199000 | 14.970629000 |
| 1 | 7.759413000 | -2.457183000 | 15.514794000 |
| 1 | 7.394781000 | -4.195104000 | 15.536409000 |
| 7 | 5.211376000 | -4.371148000 | 20.793386000 |
| 6 | 6.516009000 | -4.903343000 | 21.182202000 |
| 6 | 7.558040000 | -5.108255000 | 20.059325000 |
| 6 | 8.000680000 | -3.809507000 | 19.461653000 |
| 7 | 8.787198000 | -3.699123000 | 18.320323000 |
| 6 | 7.759351000 | -2.514069000 | 19.838606000 |
| 6 | 8.986238000 | -2.387177000 | 18.045730000 |
| 7 | 8.361615000 | -1.641049000 | 18.945596000 |
| 1 | 5.289260000 | -3.534080000 | 20.220674000 |
| 1 | 6.349165000 | -5.864797000 | 21.677397000 |
| 1 | 7.131897000 | -5.754004000 | 19.277945000 |
| 1 | 8.424105000 | -5.646992000 | 20.469789000 |
| 1 | 7.202410000 | -2.150904000 | 20.683881000 |
| 1 | 9.540372000 | -1.993460000 | 17.205380000 |
| 1 | 9.154135000 | -4.467280000 | 17.775121000 |
| 7 | 9.908159000 | -1.052972000 | 9.000198000 |
| 6 | 9.056171000 | -1.150624000 | 10.184713000 |
| 6 | 7.724886000 | -1.756730000 | 9.765385000 |
| 8 | 6.902441000 | -1.161108000 | 9.057046000 |
| 6 | 8.822654000 | 0.184508000 | 10.942884000 |
| 6 | 10.031947000 | 0.543114000 | 11.743591000 |
| 7 | 11.296828000 | 0.620063000 | 11.175217000 |
| 6 | 10.129931000 | 0.731829000 | 13.102407000 |
| 6 | 12.129966000 | 0.846544000 | 12.177386000 |
| 7 | 11.469564000 | 0.938807000 | 13.366794000 |
| 1 | 9.486798000 | -0.362217000 | 8.378254000 |
| 1 | 9.545543000 | -1.844986000 | 10.880639000 |
| 1 | 8.594242000 | 0.946942000 | 10.193475000 |
| 1 | 7.960902000 | 0.107212000 | 11.614124000 |
| 1 | 9.397990000 | 0.742236000 | 13.888663000 |
| 1 | 13.199229000 | 0.969883000 | 12.096458000 |
| 1 | 11.860116000 | 1.051924000 | 14.307217000 |
| 7 | 7.492492000 | -3.008737000 | 10.233140000 |
| 6 | 6.274701000 | -3.750073000 | 9.947501000 |
| 6 | 5.085159000 | -3.219696000 | 10.762955000 |
| 8 | 4.757025000 | -3.715706000 | 11.843278000 |
| 1 | 8.189966000 | -3.439061000 | 10.819766000 |
| 1 | 6.064206000 | -3.687686000 | 8.875062000 |
| 7 | 4.469611000 | -2.165255000 | 10.177943000 |
| 6 | 3.365280000 | -1.445527000 | 10.786493000 |
| 6 | 3.647730000 | 0.049307000 | 10.925578000 |
| 6 | 4.705269000 | 0.460094000 | 11.930554000 |
| 6 | 5.352004000 | -0.411718000 | 12.813596000 |
| 6 | 5.039878000 | 1.821164000 | 11.985464000 |
| 6 | 6.288157000 | 0.068976000 | 13.739077000 |
| 6 | 5.962357000 | 2.313440000 | 12.898497000 |
| 6 | 6.579118000 | 1.432312000 | 13.792306000 |
| 8 | 7.446938000 | 1.972819000 | 14.697429000 |
| 1 | 4.993128000 | -1.710894000 | 9.431150000 |
| 1 | 3.169889000 | -1.917721000 | 11.752812000 |
| 1 | 2.699456000 | 0.542610000 | 11.184939000 |
| 1 | 3.930548000 | 0.460460000 | 9.946102000 |
| 1 | 5.144849000 | -1.477824000 | 12.792217000 |
| 1 | 4.572464000 | 2.512332000 | 11.293452000 |
| 1 | 6.786635000 | -0.610709000 | 14.421796000 |
| 1 | 6.217727000 | 3.367481000 | 12.914258000 |
| 1 | 7.767448000 | 1.322517000 | 15.372782000 |
| 6 | 4.736851000 | 6.783291000 | 15.938771000 |

| | | | |
|---|---|---|---|
| 6 | 4.986014000 | 5.453246000 | 15.219485000 |
| 6 | 5.546685000 | 4.407548000 | 16.184641000 |
| 6 | 3.710358000 | 4.937671000 | 14.543076000 |
| 1 | 3.957221000 | 6.676048000 | 16.708594000 |
| 1 | 5.647503000 | 7.139558000 | 16.434716000 |
| 1 | 5.748507000 | 5.618174000 | 14.448406000 |
| 1 | 5.851915000 | 3.510180000 | 15.643251000 |
| 1 | 4.782790000 | 4.123677000 | 16.920671000 |
| 1 | 6.426422000 | 4.777463000 | 16.719370000 |
| 1 | 3.914871000 | 4.030167000 | 13.964698000 |
| 1 | 2.947083000 | 4.697260000 | 15.299580000 |
| 1 | 3.276290000 | 5.669264000 | 13.854540000 |
| 6 | 2.041474000 | 5.227375000 | 19.037098000 |
| 7 | 2.728103000 | 3.950893000 | 19.137236000 |
| 6 | 4.033420000 | 3.822382000 | 19.425588000 |
| 7 | 4.819152000 | 4.891777000 | 19.575837000 |
| 7 | 4.543149000 | 2.596944000 | 19.630629000 |
| 1 | 2.250460000 | 5.850977000 | 19.913120000 |
| 1 | 2.326028000 | 5.775249000 | 18.129820000 |
| 1 | 2.221508000 | 3.081427000 | 18.989074000 |
| 1 | 4.538780000 | 5.770507000 | 19.165758000 |
| 1 | 5.838216000 | 4.724012000 | 19.697350000 |
| 1 | 3.931090000 | 1.809276000 | 19.447148000 |
| 1 | 5.539538000 | 2.463046000 | 19.409501000 |
| 26 | 7.891624000 | 0.450711000 | 18.704909000 |
| 6 | 9.453621000 | 3.139880000 | 19.059152000 |
| 6 | 9.827883000 | 3.688777000 | 17.666164000 |
| 6 | 9.519778000 | 5.192515000 | 17.575451000 |
| 6 | 7.949231000 | 3.289100000 | 19.238789000 |
| 8 | 10.052680000 | 5.954051000 | 18.422482000 |
| 8 | 8.748321000 | 5.585246000 | 16.648426000 |
| 8 | 7.488384000 | 4.266887000 | 19.836597000 |
| 8 | 7.208312000 | 2.363761000 | 18.689485000 |
| 16 | 9.950233000 | 1.377533000 | 19.276749000 |
| 1 | 9.984143000 | 3.716595000 | 19.817170000 |
| 1 | 10.902012000 | 3.556891000 | 17.515672000 |
| 1 | 9.294018000 | 3.144485000 | 16.884043000 |
| 8 | 8.512608000 | 0.449170000 | 16.609561000 |
| 8 | 14.494827000 | 0.469665000 | 17.335763000 |
| 1 | 13.656452000 | 0.343661000 | 16.832088000 |
| 1 | 15.204029000 | 0.150519000 | 16.759489000 |
| 8 | 12.079762000 | 0.967303000 | 16.170207000 |
| 1 | 11.246675000 | 0.540286000 | 16.448341000 |
| 1 | 12.215812000 | 1.742141000 | 16.775924000 |
| 8 | 12.446866000 | 3.508604000 | 20.281047000 |
| 1 | 12.577518000 | 4.490297000 | 20.095338000 |
| 1 | 13.065123000 | 3.283952000 | 20.991769000 |
| 8 | 12.502220000 | 6.034843000 | 19.615984000 |
| 1 | 11.573916000 | 6.007667000 | 19.288590000 |
| 1 | 13.013169000 | 6.353663000 | 18.840460000 |
| 1 | 6.947411000 | -4.239601000 | 21.941580000 |
| 1 | 4.697840000 | -5.049309000 | 20.234274000 |
| 1 | -0.194267000 | -0.802818000 | 18.854514000 |
| 1 | 1.359815000 | 1.834720000 | 15.371445000 |
| 1 | 0.967599000 | 5.038840000 | 18.998117000 |
| 1 | 4.395569000 | 7.563986000 | 15.247125000 |
| 1 | 17.877035000 | 3.902624000 | 10.224732000 |
| 1 | 14.786376000 | 7.743926000 | 8.444278000 |
| 1 | 9.881527000 | 3.624126000 | 4.915945000 |
| 1 | 5.691904000 | 1.261464000 | 7.388246000 |
| 1 | 10.787210000 | -0.636357000 | 9.312749000 |
| 1 | 3.802178000 | -2.633800000 | 21.880905000 |
| 1 | 8.429765000 | 8.726278000 | 10.658874000 |
| 1 | 13.253976000 | 9.287252000 | 9.306441000 |
| 1 | 15.606562000 | 9.646972000 | 13.556374000 |
| 1 | 2.465613000 | -1.578196000 | 10.171471000 |
| 1 | 6.431401000 | -4.793653000 | 10.221856000 |
| 1 | 9.484061000 | -4.030402000 | 12.383700000 |
| 8 | 8.260234000 | 4.618515000 | 14.161590000 |
| 1 | 8.472428000 | 4.946586000 | 15.067779000 |
| 1 | 7.981488000 | 3.695660000 | 14.313617000 |
| 8 | 13.145964000 | 2.815643000 | 17.823674000 |
| 1 | 12.872178000 | 2.940048000 | 18.779180000 |
| 1 | 13.894360000 | 2.180018000 | 17.826161000 |
| 8 | 9.498037000 | -0.250249000 | 16.098820000 |

**$^3Re_S$:**

| | | | |
|---|---|---|---|
| 6 | 14.870344000 | 8.861816000 | 13.453349000 |
| 6 | 13.704358000 | 9.274414000 | 12.567685000 |
| 8 | 12.633503000 | 9.708507000 | 13.013625000 |
| 6 | 14.444651000 | 8.386790000 | 14.842963000 |
| 6 | 13.562384000 | 7.132655000 | 14.802084000 |
| 6 | 13.452506000 | 6.468243000 | 16.166573000 |
| 8 | 13.463284000 | 7.147093000 | 17.213828000 |
| 7 | 13.342292000 | 5.134935000 | 16.160716000 |
| 1 | 15.457908000 | 8.093040000 | 12.938108000 |
| 1 | 15.344264000 | 8.180960000 | 15.432610000 |
| 1 | 13.915115000 | 9.189282000 | 15.363979000 |
| 1 | 13.948721000 | 6.412689000 | 14.071844000 |
| 1 | 12.555877000 | 7.395695000 | 14.455510000 |
| 1 | 13.345807000 | 4.568957000 | 17.012500000 |
| 1 | 13.390123000 | 4.623166000 | 15.283273000 |
| 7 | 13.936069000 | 9.171389000 | 11.234755000 |
| 6 | 12.843978000 | 9.426887000 | 10.321151000 |
| 6 | 11.680464000 | 8.466743000 | 10.611922000 |
| 8 | 11.907911000 | 7.291544000 | 10.916373000 |
| 1 | 14.724835000 | 8.597971000 | 10.923731000 |
| 1 | 12.519784000 | 10.469471000 | 10.402875000 |
| 7 | 10.434476000 | 8.970248000 | 10.494340000 |
| 6 | 9.300126000 | 8.274366000 | 11.110329000 |
| 6 | 9.323210000 | 8.552740000 | 12.616127000 |
| 6 | 8.101549000 | 8.083769000 | 13.399270000 |
| 6 | 8.103950000 | 8.651269000 | 14.824893000 |
| 7 | 9.134416000 | 8.092385000 | 15.695310000 |
| 6 | 10.273253000 | 8.674588000 | 16.070188000 |
| 7 | 10.805965000 | 9.711711000 | 15.365812000 |
| 7 | 10.870098000 | 8.281404000 | 17.198842000 |
| 1 | 10.363976000 | 9.978907000 | 10.442433000 |
| 1 | 9.391828000 | 7.207145000 | 10.899037000 |
| 1 | 10.229427000 | 8.087481000 | 13.019552000 |
| 1 | 9.427620000 | 9.640297000 | 12.750671000 |
| 1 | 7.187677000 | 8.441638000 | 12.904579000 |
| 1 | 8.048208000 | 6.988989000 | 13.430208000 |
| 1 | 8.228490000 | 9.739266000 | 14.799449000 |
| 1 | 7.143991000 | 8.440097000 | 15.308046000 |
| 1 | 8.921595000 | 7.177207000 | 16.142530000 |
| 1 | 11.649780000 | 10.110373000 | 15.756363000 |
| 1 | 10.859876000 | 9.641486000 | 14.355219000 |
| 1 | 10.431686000 | 7.511926000 | 17.735647000 |
| 1 | 11.890119000 | 8.283735000 | 17.244881000 |
| 7 | 5.948853000 | 1.641183000 | 8.244668000 |
| 6 | 6.808283000 | 2.820631000 | 8.108410000 |
| 6 | 8.181095000 | 2.490850000 | 7.519052000 |
| 8 | 8.903552000 | 1.616481000 | 8.012204000 |
| 6 | 7.064776000 | 3.481838000 | 9.476452000 |
| 6 | 5.978929000 | 4.400924000 | 9.990978000 |
| 6 | 4.650233000 | 4.349843000 | 9.549100000 |
| 6 | 6.310034000 | 5.328861000 | 10.989372000 |
| 6 | 3.675642000 | 5.194607000 | 10.081977000 |
| 6 | 5.342086000 | 6.171955000 | 11.531953000 |
| 6 | 4.022465000 | 6.108669000 | 11.077480000 |
| 8 | 3.029048000 | 6.910272000 | 11.589063000 |
| 1 | 6.471110000 | 0.896495000 | 8.714223000 |
| 1 | 6.297977000 | 3.537729000 | 7.452327000 |
| 1 | 7.234998000 | 2.674794000 | 10.200771000 |
| 1 | 8.002969000 | 4.044775000 | 9.449096000 |
| 1 | 4.367139000 | 3.614130000 | 8.803083000 |
| 1 | 7.335647000 | 5.376628000 | 11.344356000 |
| 1 | 2.646799000 | 5.154230000 | 9.738243000 |
| 1 | 5.614107000 | 6.877921000 | 12.313552000 |

| | | | |
|---|---|---|---|
| 1 | 3.433150000 | 7.534923000 | 12.211026000 |
| 7 | 8.587062000 | 3.231774000 | 6.457108000 |
| 6 | 9.968008000 | 3.171719000 | 5.981988000 |
| 6 | 10.866056000 | 4.076718000 | 6.834573000 |
| 8 | 11.258113000 | 5.167567000 | 6.432116000 |
| 1 | 8.038769000 | 4.040727000 | 6.197357000 |
| 1 | 10.289561000 | 2.128934000 | 6.030062000 |
| 7 | 11.131694000 | 3.611074000 | 8.082474000 |
| 6 | 11.754991000 | 4.502779000 | 9.039892000 |
| 6 | 13.197119000 | 4.838454000 | 8.627751000 |
| 8 | 13.992572000 | 4.030712000 | 8.147410000 |
| 6 | 11.721760000 | 3.918758000 | 10.456186000 |
| 6 | 10.500935000 | 4.394017000 | 11.250143000 |
| 8 | 9.584166000 | 5.029610000 | 10.718528000 |
| 7 | 10.525438000 | 4.098974000 | 12.564758000 |
| 1 | 10.713814000 | 2.737980000 | 8.386393000 |
| 1 | 11.187841000 | 5.437145000 | 9.049123000 |
| 1 | 12.615073000 | 4.226400000 | 11.010508000 |
| 1 | 11.739948000 | 2.822635000 | 10.431177000 |
| 1 | 11.200236000 | 3.431494000 | 12.913855000 |
| 1 | 9.690822000 | 4.291954000 | 13.137910000 |
| 7 | 13.513828000 | 6.135038000 | 8.889180000 |
| 6 | 14.829707000 | 6.662743000 | 8.600368000 |
| 6 | 15.790081000 | 6.363721000 | 9.748250000 |
| 8 | 16.026549000 | 7.188796000 | 10.643230000 |
| 1 | 12.921198000 | 6.639870000 | 9.548628000 |
| 1 | 15.179992000 | 6.211034000 | 7.668806000 |
| 7 | 16.294879000 | 5.111497000 | 9.711099000 |
| 6 | 17.133306000 | 4.556175000 | 10.760206000 |
| 6 | 16.367626000 | 3.612000000 | 11.696651000 |
| 6 | 15.301309000 | 4.335201000 | 12.518222000 |
| 16 | 14.342723000 | 3.209626000 | 13.603009000 |
| 6 | 15.580204000 | 2.859632000 | 14.899301000 |
| 1 | 15.874265000 | 4.485229000 | 9.024011000 |
| 1 | 17.550052000 | 5.400450000 | 11.315119000 |
| 1 | 15.892174000 | 2.816529000 | 11.110267000 |
| 1 | 17.092630000 | 3.128553000 | 12.362634000 |
| 1 | 15.743692000 | 5.127984000 | 13.131651000 |
| 1 | 14.564835000 | 4.810059000 | 11.863248000 |
| 1 | 15.119356000 | 2.168869000 | 15.609276000 |
| 1 | 15.852548000 | 3.776000000 | 15.430786000 |
| 1 | 16.472945000 | 2.393276000 | 14.475237000 |
| 6 | 3.231938000 | -1.792907000 | 21.922221000 |
| 6 | 2.714120000 | -1.701396000 | 20.491383000 |
| 8 | 3.330557000 | -2.124772000 | 19.511401000 |
| 6 | 3.844173000 | -0.480555000 | 22.473144000 |
| 6 | 5.120714000 | -0.079056000 | 21.799733000 |
| 7 | 6.168696000 | 0.573882000 | 22.437376000 |
| 6 | 5.539576000 | -0.222269000 | 20.504089000 |
| 6 | 7.163583000 | 0.788759000 | 21.534998000 |
| 7 | 6.806162000 | 0.309970000 | 20.356777000 |
| 1 | 2.398896000 | -2.074754000 | 22.576840000 |
| 1 | 3.109602000 | 0.334215000 | 22.396799000 |
| 1 | 4.027704000 | -0.625887000 | 23.544334000 |
| 1 | 5.020507000 | -0.681574000 | 19.684196000 |
| 1 | 8.104948000 | 1.271251000 | 21.753229000 |
| 1 | 6.189910000 | 0.856374000 | 23.407618000 |
| 7 | 1.479904000 | -1.132960000 | 20.348877000 |
| 6 | 0.937352000 | -0.885989000 | 19.037225000 |
| 6 | 1.554656000 | 0.351219000 | 18.364011000 |
| 8 | 2.113749000 | 1.223233000 | 19.038262000 |
| 1 | 1.061334000 | -0.664144000 | 21.139222000 |
| 1 | 1.111803000 | -1.770956000 | 18.419254000 |
| 7 | 1.385697000 | 0.413633000 | 17.027948000 |
| 6 | 1.955943000 | 1.444748000 | 16.162649000 |
| 6 | 3.115574000 | 0.898860000 | 15.309601000 |
| 6 | 4.203686000 | 0.312669000 | 16.146042000 |
| 7 | 4.367643000 | -1.048115000 | 16.356703000 |
| 6 | 5.204288000 | 0.905043000 | 16.870883000 |
| 6 | 5.450092000 | -1.241953000 | 17.149078000 |
| 7 | 5.971467000 | -0.070416000 | 17.480348000 |
| 1 | 0.930470000 | -0.369335000 | 16.575756000 |
| 1 | 2.313109000 | 2.246400000 | 16.810005000 |
| 1 | 2.747301000 | 0.152029000 | 14.596117000 |
| 1 | 3.523854000 | 1.714092000 | 14.703500000 |
| 1 | 5.459050000 | 1.945092000 | 16.965057000 |
| 1 | 5.807700000 | -2.208033000 | 17.464681000 |
| 1 | 3.826706000 | -1.782827000 | 15.923594000 |
| 6 | 9.165947000 | -3.852645000 | 13.446930000 |
| 6 | 7.702766000 | -3.396814000 | 13.534483000 |
| 6 | 10.165732000 | -2.859968000 | 14.046841000 |
| 6 | 7.178001000 | -3.271477000 | 14.966665000 |
| 1 | 9.269893000 | -4.833214000 | 13.934192000 |
| 1 | 7.054206000 | -4.091533000 | 12.988886000 |
| 1 | 7.592636000 | -2.428283000 | 13.031608000 |
| 1 | 9.998917000 | -2.723652000 | 15.120903000 |
| 1 | 10.075633000 | -1.872299000 | 13.581869000 |
| 1 | 11.196694000 | -3.204493000 | 13.911613000 |
| 1 | 6.103094000 | -3.060590000 | 14.955669000 |
| 1 | 7.671374000 | -2.463304000 | 15.517092000 |
| 1 | 7.328167000 | -4.202662000 | 15.529803000 |
| 7 | 5.150175000 | -4.460138000 | 20.751112000 |
| 6 | 6.482598000 | -4.938692000 | 21.112614000 |
| 6 | 7.497246000 | -5.166385000 | 19.968754000 |
| 6 | 7.938325000 | -3.882194000 | 19.339286000 |
| 7 | 8.880892000 | -3.789983000 | 18.322590000 |
| 6 | 7.577170000 | -2.583553000 | 19.589398000 |
| 6 | 9.049575000 | -2.482285000 | 17.998731000 |
| 7 | 8.261420000 | -1.724652000 | 18.744814000 |
| 1 | 5.169677000 | -3.634651000 | 20.156968000 |
| 1 | 6.357720000 | -5.882649000 | 21.652291000 |
| 1 | 7.053599000 | -5.829533000 | 19.211638000 |
| 1 | 8.371092000 | -5.697947000 | 20.372134000 |
| 1 | 6.887417000 | -2.207024000 | 20.323395000 |
| 1 | 9.724381000 | -2.119445000 | 17.240005000 |
| 1 | 9.371010000 | -4.564176000 | 17.895820000 |
| 7 | 9.950948000 | -1.054733000 | 9.099807000 |
| 6 | 9.060593000 | -1.129660000 | 10.258031000 |
| 6 | 7.735480000 | -1.722562000 | 9.799178000 |
| 8 | 6.930440000 | -1.114551000 | 9.081887000 |
| 6 | 8.825577000 | 0.213814000 | 10.999270000 |
| 6 | 10.014657000 | 0.561155000 | 11.836620000 |
| 7 | 11.304660000 | 0.573429000 | 11.323004000 |
| 6 | 10.062127000 | 0.801392000 | 13.189880000 |
| 6 | 12.101098000 | 0.810380000 | 12.352554000 |
| 7 | 11.394908000 | 0.972066000 | 13.508120000 |
| 1 | 9.563838000 | -0.358293000 | 8.462188000 |
| 1 | 9.517686000 | -1.826567000 | 10.972718000 |
| 1 | 8.628717000 | 0.973427000 | 10.238232000 |
| 1 | 7.942026000 | 0.152842000 | 11.643915000 |
| 1 | 9.290625000 | 0.887602000 | 13.930772000 |
| 1 | 13.176637000 | 0.894932000 | 12.313992000 |
| 1 | 11.759069000 | 1.099765000 | 14.459021000 |
| 7 | 7.489168000 | -2.982605000 | 10.236865000 |
| 6 | 6.282270000 | -3.717646000 | 9.894980000 |
| 6 | 5.062795000 | -3.200537000 | 10.674383000 |
| 8 | 4.707817000 | -3.703452000 | 11.742836000 |
| 1 | 8.177308000 | -3.431824000 | 10.820239000 |
| 1 | 6.110787000 | -3.640116000 | 8.816516000 |
| 7 | 4.452690000 | -2.151791000 | 10.072840000 |
| 6 | 3.313878000 | -1.452937000 | 10.641829000 |
| 6 | 3.569800000 | 0.044155000 | 10.811243000 |
| 6 | 4.592729000 | 0.449180000 | 11.853428000 |
| 6 | 5.214628000 | -0.436299000 | 12.740520000 |
| 6 | 4.928119000 | 1.809097000 | 11.933331000 |
| 6 | 6.133650000 | 0.029584000 | 13.689096000 |
| 6 | 5.835982000 | 2.286653000 | 12.869465000 |
| 6 | 6.431319000 | 1.389202000 | 13.759296000 |
| 8 | 7.289780000 | 1.910797000 | 14.689852000 |
| 1 | 4.997454000 | -1.684856000 | 9.349951000 |

| | | | |
|---|---|---|---|
| 1 | 3.083220000 | -1.939432000 | 11.593055000 |
| 1 | 2.607478000 | 0.520853000 | 11.048321000 |
| 1 | 3.878456000 | 0.473108000 | 9.847324000 |
| 1 | 5.004812000 | -1.501145000 | 12.701680000 |
| 1 | 4.479295000 | 2.510388000 | 11.239112000 |
| 1 | 6.612319000 | -0.660703000 | 14.375913000 |
| 1 | 6.100738000 | 3.337901000 | 12.901499000 |
| 1 | 7.637633000 | 1.226016000 | 15.301796000 |
| 6 | 4.672791000 | 6.808581000 | 15.933716000 |
| 6 | 4.931681000 | 5.489534000 | 15.197536000 |
| 6 | 5.487553000 | 4.433072000 | 16.153111000 |
| 6 | 3.661656000 | 4.980091000 | 14.505777000 |
| 1 | 3.889328000 | 6.684610000 | 16.696918000 |
| 1 | 5.578806000 | 7.161543000 | 16.440611000 |
| 1 | 5.698466000 | 5.666165000 | 14.433066000 |
| 1 | 5.784455000 | 3.536828000 | 15.605774000 |
| 1 | 4.722668000 | 4.149964000 | 16.888458000 |
| 1 | 6.371205000 | 4.790071000 | 16.689958000 |
| 1 | 3.870032000 | 4.076183000 | 13.923140000 |
| 1 | 2.891417000 | 4.736037000 | 15.253698000 |
| 1 | 3.234151000 | 5.716328000 | 13.817836000 |
| 6 | 2.078782000 | 5.318859000 | 19.163213000 |
| 7 | 2.769830000 | 4.041167000 | 19.189387000 |
| 6 | 4.088442000 | 3.906767000 | 19.415834000 |
| 7 | 4.883664000 | 4.971809000 | 19.537136000 |
| 7 | 4.603693000 | 2.679780000 | 19.583633000 |
| 1 | 2.343766000 | 5.922732000 | 20.037992000 |
| 1 | 2.305270000 | 5.887921000 | 18.252570000 |
| 1 | 2.251892000 | 3.174516000 | 19.075118000 |
| 1 | 4.592900000 | 5.855157000 | 19.145259000 |
| 1 | 5.903905000 | 4.787013000 | 19.627564000 |
| 1 | 3.979096000 | 1.894844000 | 19.439140000 |
| 1 | 5.589760000 | 2.540190000 | 19.317254000 |
| 26 | 7.789569000 | 0.418819000 | 18.449991000 |
| 6 | 9.459411000 | 3.077241000 | 18.982143000 |
| 6 | 9.879253000 | 3.653772000 | 17.615172000 |
| 6 | 9.543190000 | 5.149012000 | 17.525701000 |
| 6 | 7.954816000 | 3.262753000 | 19.128277000 |
| 8 | 10.069352000 | 5.921354000 | 18.368773000 |
| 8 | 8.748817000 | 5.533771000 | 16.613408000 |
| 8 | 7.509770000 | 4.224443000 | 19.766617000 |
| 8 | 7.204046000 | 2.397702000 | 18.507827000 |
| 16 | 9.876469000 | 1.290407000 | 19.183303000 |
| 1 | 9.973339000 | 3.619923000 | 19.774915000 |
| 1 | 10.961392000 | 3.544591000 | 17.509133000 |
| 1 | 9.389345000 | 3.106289000 | 16.806590000 |
| 8 | 8.512608000 | 0.340028000 | 16.547643000 |
| 8 | 14.575144000 | 0.448797000 | 17.354283000 |
| 1 | 13.713884000 | 0.380078000 | 16.878539000 |
| 1 | 15.242383000 | 0.084005000 | 16.755385000 |
| 8 | 12.140788000 | 1.095121000 | 16.287830000 |
| 1 | 11.348939000 | 0.736690000 | 16.733383000 |
| 1 | 12.404982000 | 1.884173000 | 16.833884000 |
| 8 | 12.448996000 | 3.446106000 | 20.254412000 |
| 1 | 12.552174000 | 4.433955000 | 20.077376000 |
| 1 | 13.014877000 | 3.251577000 | 21.016053000 |
| 8 | 12.481291000 | 5.976170000 | 19.602006000 |
| 1 | 11.560717000 | 5.950870000 | 19.249067000 |
| 1 | 13.007380000 | 6.326843000 | 18.851525000 |
| 1 | 6.921112000 | -4.231947000 | 21.827923000 |
| 1 | 4.639334000 | -5.172918000 | 20.234466000 |
| 1 | -0.144181000 | -0.735781000 | 19.121685000 |
| 1 | 1.169087000 | 1.851785000 | 15.519192000 |
| 1 | 1.004296000 | 5.130898000 | 19.188102000 |
| 1 | 4.332259000 | 7.598248000 | 15.252112000 |
| 1 | 17.963359000 | 4.015174000 | 10.292756000 |
| 1 | 14.771073000 | 7.745258000 | 8.467429000 |
| 1 | 10.009786000 | 3.524256000 | 4.951163000 |
| 1 | 5.740568000 | 1.269413000 | 7.317898000 |
| 1 | 10.829527000 | -0.658616000 | 9.439041000 |
| 1 | 3.963665000 | -2.605677000 | 21.945029000 |
| 1 | 8.379469000 | 8.644360000 | 10.647237000 |
| 1 | 13.197957000 | 9.264435000 | 9.298432000 |
| 1 | 15.527972000 | 9.735962000 | 13.553214000 |
| 1 | 2.443427000 | -1.589247000 | 9.987032000 |
| 1 | 6.428265000 | -4.765136000 | 10.160237000 |
| 1 | 9.434856000 | -4.022282000 | 12.393679000 |
| 8 | 8.213329000 | 4.546754000 | 14.155606000 |
| 1 | 8.432439000 | 4.870932000 | 15.063388000 |
| 1 | 7.911366000 | 3.631788000 | 14.302071000 |
| 8 | 13.383564000 | 2.875544000 | 17.856206000 |
| 1 | 13.051656000 | 2.962453000 | 18.797685000 |
| 1 | 14.100719000 | 2.205763000 | 17.864575000 |
| 8 | 9.641492000 | -0.167810000 | 16.183426000 |

**$^{3}$IM1$_{S}$:**

| | | | |
|---|---|---|---|
| 6 | 15.371811000 | 7.797898000 | 13.537036000 |
| 6 | 14.415272000 | 8.433425000 | 12.538808000 |
| 8 | 13.487593000 | 9.178512000 | 12.880923000 |
| 6 | 14.678183000 | 7.420190000 | 14.848003000 |
| 6 | 13.544250000 | 6.406557000 | 14.662658000 |
| 6 | 12.931530000 | 5.993277000 | 15.986847000 |
| 8 | 12.940339000 | 6.758385000 | 16.971956000 |
| 7 | 12.353668000 | 4.775817000 | 16.036807000 |
| 1 | 15.859499000 | 6.930724000 | 13.079294000 |
| 1 | 15.423795000 | 7.014583000 | 15.541348000 |
| 1 | 14.277517000 | 8.319328000 | 15.322544000 |
| 1 | 13.889575000 | 5.516290000 | 14.126654000 |
| 1 | 12.748903000 | 6.837544000 | 14.040167000 |
| 1 | 11.948556000 | 4.491781000 | 16.927242000 |
| 1 | 12.576448000 | 4.076493000 | 15.333562000 |
| 7 | 14.653650000 | 8.134464000 | 11.238994000 |
| 6 | 13.710951000 | 8.571123000 | 10.234111000 |
| 6 | 12.301932000 | 8.046745000 | 10.566962000 |
| 8 | 12.152971000 | 6.941105000 | 11.096106000 |
| 1 | 15.350404000 | 7.418191000 | 11.018836000 |
| 1 | 13.704785000 | 9.664385000 | 10.159969000 |
| 7 | 11.267773000 | 8.842045000 | 10.228922000 |
| 6 | 9.937039000 | 8.622252000 | 10.808773000 |
| 6 | 9.951278000 | 9.122499000 | 12.257762000 |
| 6 | 8.592143000 | 9.277080000 | 12.935835000 |
| 6 | 8.718452000 | 10.026824000 | 14.271856000 |
| 7 | 9.349385000 | 9.256072000 | 15.342243000 |
| 6 | 10.619275000 | 9.300286000 | 15.735554000 |
| 7 | 11.577064000 | 9.908436000 | 14.991199000 |
| 7 | 10.965427000 | 8.796867000 | 16.930648000 |
| 1 | 11.494142000 | 9.796053000 | 9.977337000 |
| 1 | 9.703049000 | 7.556666000 | 10.758282000 |
| 1 | 10.582802000 | 8.432323000 | 12.827830000 |
| 1 | 10.451385000 | 10.103450000 | 12.257390000 |
| 1 | 7.923782000 | 9.866350000 | 12.292688000 |
| 1 | 8.114670000 | 8.303266000 | 13.092708000 |
| 1 | 9.282008000 | 10.956912000 | 14.136870000 |
| 1 | 7.724187000 | 10.297321000 | 14.640987000 |
| 1 | 8.727972000 | 8.628465000 | 15.883241000 |
| 1 | 12.514828000 | 9.900131000 | 15.368081000 |
| 1 | 11.573928000 | 9.806893000 | 13.984488000 |
| 1 | 10.242681000 | 8.335512000 | 17.500638000 |
| 1 | 11.880318000 | 8.350994000 | 17.000792000 |
| 7 | 5.581409000 | 1.550432000 | 8.438153000 |
| 6 | 6.631528000 | 2.573406000 | 8.355632000 |
| 6 | 7.980658000 | 1.929882000 | 8.031891000 |
| 8 | 8.301495000 | 0.862789000 | 8.557322000 |
| 6 | 6.817237000 | 3.312640000 | 9.697621000 |
| 6 | 5.768387000 | 4.336337000 | 10.079988000 |
| 6 | 4.465004000 | 4.344585000 | 9.569456000 |
| 6 | 6.106356000 | 5.299605000 | 11.040584000 |
| 6 | 3.518951000 | 5.265289000 | 10.021623000 |
| 6 | 5.166012000 | 6.210143000 | 11.515239000 |
| 6 | 3.861343000 | 6.180907000 | 11.018251000 |

| | | | |
|---|---|---|---|
| 8 | 2.877403000 | 7.017536000 | 11.489767000 |
| 1 | 5.968313000 | 0.761853000 | 8.963452000 |
| 1 | 6.354345000 | 3.295290000 | 7.577457000 |
| 1 | 6.866278000 | 2.543355000 | 10.478890000 |
| 1 | 7.799272000 | 3.802008000 | 9.705829000 |
| 1 | 4.175864000 | 3.600815000 | 8.835202000 |
| 1 | 7.122373000 | 5.346893000 | 11.419435000 |
| 1 | 2.505477000 | 5.265408000 | 9.631754000 |
| 1 | 5.453449000 | 6.913441000 | 12.293790000 |
| 1 | 3.270640000 | 7.599209000 | 12.158874000 |
| 7 | 8.809076000 | 2.593445000 | 7.184644000 |
| 6 | 10.223292000 | 2.239183000 | 7.113851000 |
| 6 | 11.015483000 | 3.154363000 | 8.046434000 |
| 8 | 11.611556000 | 4.137292000 | 7.609524000 |
| 1 | 8.557183000 | 3.534932000 | 6.910909000 |
| 1 | 10.303235000 | 1.187824000 | 7.395433000 |
| 7 | 10.914089000 | 2.886075000 | 9.378490000 |
| 6 | 11.581110000 | 3.813027000 | 10.276286000 |
| 6 | 13.106906000 | 3.700549000 | 10.069253000 |
| 8 | 13.711460000 | 2.631160000 | 9.925743000 |
| 6 | 11.111688000 | 3.690018000 | 11.730541000 |
| 6 | 9.970150000 | 4.671945000 | 12.031201000 |
| 8 | 9.419279000 | 5.322864000 | 11.143157000 |
| 7 | 9.631379000 | 4.783387000 | 13.338801000 |
| 1 | 10.682857000 | 1.945942000 | 9.702585000 |
| 1 | 11.291233000 | 4.809371000 | 9.942155000 |
| 1 | 11.938396000 | 3.893878000 | 12.420999000 |
| 1 | 10.761201000 | 2.680299000 | 11.956170000 |
| 1 | 9.947346000 | 4.098955000 | 14.012811000 |
| 1 | 8.809367000 | 5.352356000 | 13.583000000 |
| 7 | 13.720121000 | 4.904261000 | 9.975638000 |
| 6 | 15.000867000 | 5.053101000 | 9.314862000 |
| 6 | 16.136598000 | 4.920759000 | 10.320311000 |
| 8 | 16.542788000 | 5.892375000 | 10.972422000 |
| 1 | 13.187160000 | 5.732119000 | 10.244086000 |
| 1 | 15.076108000 | 4.299411000 | 8.525422000 |
| 7 | 16.588332000 | 3.654437000 | 10.468520000 |
| 6 | 17.408944000 | 3.251842000 | 11.602329000 |
| 6 | 16.561038000 | 2.541169000 | 12.667885000 |
| 6 | 15.398519000 | 3.417636000 | 13.130484000 |
| 16 | 14.216194000 | 2.584704000 | 14.241141000 |
| 6 | 15.175503000 | 2.576333000 | 15.795710000 |
| 1 | 15.994359000 | 2.943053000 | 10.046613000 |
| 1 | 17.863793000 | 4.161808000 | 12.000279000 |
| 1 | 16.159847000 | 1.607467000 | 12.254690000 |
| 1 | 17.204429000 | 2.268218000 | 13.512901000 |
| 1 | 15.763110000 | 4.331412000 | 13.612801000 |
| 1 | 14.795978000 | 3.720533000 | 12.271866000 |
| 1 | 14.517757000 | 2.239726000 | 16.599551000 |
| 1 | 15.515222000 | 3.590269000 | 16.028245000 |
| 1 | 16.038943000 | 1.910135000 | 15.721972000 |
| 6 | 4.656683000 | -2.591457000 | 21.315586000 |
| 6 | 4.180573000 | -2.818205000 | 19.889072000 |
| 8 | 4.890598000 | -3.302617000 | 19.009331000 |
| 6 | 4.915766000 | -1.091580000 | 21.596311000 |
| 6 | 5.911957000 | -0.458769000 | 20.673919000 |
| 7 | 7.028582000 | 0.222905000 | 21.130556000 |
| 6 | 5.952312000 | -0.315096000 | 19.308029000 |
| 6 | 7.687409000 | 0.751987000 | 20.069076000 |
| 7 | 7.048021000 | 0.442074000 | 18.950604000 |
| 1 | 3.902506000 | -2.955439000 | 22.023114000 |
| 1 | 3.975027000 | -0.528699000 | 21.535354000 |
| 1 | 5.263750000 | -0.993764000 | 22.630479000 |
| 1 | 5.272659000 | -0.682902000 | 18.559364000 |
| 1 | 8.592443000 | 1.337250000 | 20.150188000 |
| 1 | 7.308159000 | 0.306112000 | 22.097961000 |
| 7 | 2.900553000 | -2.397597000 | 19.643747000 |
| 6 | 2.392498000 | -2.258767000 | 18.303521000 |
| 6 | 2.049390000 | -0.800436000 | 17.981658000 |
| 8 | 2.260578000 | 0.105566000 | 18.803174000 |

| | | | |
|---|---|---|---|
| 1 | 2.431714000 | -1.859153000 | 20.359425000 |
| 1 | 3.175514000 | -2.612905000 | 17.624521000 |
| 7 | 1.496287000 | -0.610898000 | 16.769928000 |
| 6 | 1.385340000 | 0.694098000 | 16.121887000 |
| 6 | 2.370342000 | 0.798402000 | 14.945593000 |
| 6 | 3.775029000 | 0.505846000 | 15.361559000 |
| 7 | 4.436413000 | -0.646256000 | 14.977805000 |
| 6 | 4.662984000 | 1.192432000 | 16.150784000 |
| 6 | 5.685799000 | -0.617040000 | 15.491226000 |
| 7 | 5.845905000 | 0.480622000 | 16.225053000 |
| 1 | 1.426290000 | -1.419826000 | 16.162350000 |
| 1 | 1.609102000 | 1.445039000 | 16.880889000 |
| 1 | 2.085121000 | 0.095810000 | 14.155406000 |
| 1 | 2.311749000 | 1.801933000 | 14.513102000 |
| 1 | 4.571033000 | 2.159216000 | 16.612824000 |
| 1 | 6.445739000 | -1.361564000 | 15.319079000 |
| 1 | 4.054443000 | -1.321532000 | 14.300957000 |
| 6 | 11.017104000 | -3.195226000 | 13.369974000 |
| 6 | 9.685465000 | -3.505254000 | 14.070525000 |
| 6 | 12.125898000 | -2.749429000 | 14.327816000 |
| 6 | 9.735590000 | -4.729866000 | 14.988130000 |
| 1 | 11.346733000 | -4.082802000 | 12.813147000 |
| 1 | 8.918942000 | -3.681702000 | 13.301076000 |
| 1 | 9.358545000 | -2.623694000 | 14.638644000 |
| 1 | 12.382797000 | -3.532498000 | 15.050786000 |
| 1 | 11.814384000 | -1.864150000 | 14.891340000 |
| 1 | 13.036895000 | -2.492855000 | 13.775931000 |
| 1 | 8.764345000 | -4.913477000 | 15.460473000 |
| 1 | 10.468075000 | -4.601088000 | 15.791133000 |
| 1 | 10.014820000 | -5.628603000 | 14.425332000 |
| 7 | 7.710489000 | -3.377388000 | 20.376141000 |
| 6 | 9.011048000 | -3.972303000 | 20.082785000 |
| 6 | 9.333928000 | -4.140848000 | 18.573441000 |
| 6 | 9.238342000 | -2.852972000 | 17.823434000 |
| 7 | 10.325455000 | -2.057880000 | 17.498924000 |
| 6 | 8.141737000 | -2.127858000 | 17.409432000 |
| 6 | 9.864790000 | -0.924077000 | 16.914902000 |
| 7 | 8.543762000 | -0.926993000 | 16.848527000 |
| 1 | 7.674973000 | -2.442164000 | 19.972895000 |
| 1 | 9.062587000 | -4.957466000 | 20.560099000 |
| 1 | 8.624133000 | -4.857197000 | 18.142969000 |
| 1 | 10.331905000 | -4.581307000 | 18.453796000 |
| 1 | 7.098571000 | -2.391485000 | 17.506470000 |
| 1 | 10.515146000 | -0.141371000 | 16.562479000 |
| 1 | 11.317908000 | -2.229438000 | 17.630789000 |
| 7 | 8.888289000 | -1.811392000 | 9.663887000 |
| 6 | 8.235251000 | -1.615569000 | 10.947361000 |
| 6 | 6.744746000 | -1.894024000 | 10.785505000 |
| 8 | 6.081101000 | -1.474927000 | 9.832757000 |
| 6 | 8.385043000 | -0.204314000 | 11.594122000 |
| 6 | 9.795599000 | 0.118676000 | 11.960502000 |
| 7 | 10.807603000 | 0.250990000 | 11.018742000 |
| 6 | 10.320386000 | 0.280843000 | 13.223200000 |
| 6 | 11.919323000 | 0.477078000 | 11.704805000 |
| 7 | 11.673391000 | 0.500308000 | 13.043196000 |
| 1 | 8.461394000 | -1.163784000 | 9.002213000 |
| 1 | 8.650631000 | -2.342785000 | 11.655378000 |
| 1 | 7.992881000 | 0.529379000 | 10.885311000 |
| 1 | 7.778231000 | -0.168798000 | 12.505027000 |
| 1 | 9.846093000 | 0.261573000 | 14.191497000 |
| 1 | 12.881688000 | 0.710433000 | 11.271466000 |
| 1 | 12.332792000 | 0.735959000 | 13.777646000 |
| 7 | 6.154960000 | -2.590214000 | 11.803384000 |
| 6 | 4.796099000 | -3.106048000 | 11.671656000 |
| 6 | 3.699478000 | -2.058151000 | 11.926454000 |
| 8 | 3.103237000 | -2.018687000 | 13.021344000 |
| 1 | 6.760405000 | -3.024323000 | 12.486076000 |
| 1 | 4.669700000 | -3.529489000 | 10.668758000 |
| 7 | 3.435264000 | -1.258161000 | 10.887936000 |
| 6 | 2.427954000 | -0.202688000 | 10.883073000 |

| | | | |
|---|---|---|---|
| 6 | 3.011037000 | 1.194589000 | 10.620366000 |
| 6 | 3.850250000 | 1.778215000 | 11.731532000 |
| 6 | 5.091356000 | 1.235808000 | 12.086784000 |
| 6 | 3.447393000 | 2.956482000 | 12.373684000 |
| 6 | 5.898789000 | 1.840067000 | 13.048587000 |
| 6 | 4.246927000 | 3.578960000 | 13.325031000 |
| 6 | 5.476375000 | 3.015348000 | 13.669874000 |
| 8 | 6.245367000 | 3.648970000 | 14.607430000 |
| 1 | 4.104303000 | -1.304174000 | 10.119076000 |
| 1 | 1.913258000 | -0.246469000 | 11.844249000 |
| 1 | 2.164502000 | 1.863337000 | 10.425397000 |
| 1 | 3.607522000 | 1.179594000 | 9.698183000 |
| 1 | 5.455375000 | 0.354209000 | 11.574364000 |
| 1 | 2.511995000 | 3.427103000 | 12.086251000 |
| 1 | 6.865005000 | 1.426996000 | 13.310866000 |
| 1 | 3.951745000 | 4.519676000 | 13.768242000 |
| 1 | 6.988831000 | 3.049789000 | 14.882966000 |
| 6 | 4.640652000 | 6.942785000 | 17.282252000 |
| 6 | 4.120458000 | 6.164389000 | 16.068080000 |
| 6 | 3.337583000 | 4.922815000 | 16.514170000 |
| 6 | 3.283404000 | 7.061437000 | 15.147915000 |
| 1 | 3.802408000 | 7.318794000 | 17.888096000 |
| 1 | 5.275586000 | 6.315923000 | 17.916214000 |
| 1 | 4.991002000 | 5.818876000 | 15.500357000 |
| 1 | 2.926374000 | 4.370236000 | 15.663157000 |
| 1 | 2.495901000 | 5.204332000 | 17.163875000 |
| 1 | 3.995132000 | 4.248277000 | 17.069487000 |
| 1 | 2.956432000 | 6.530692000 | 14.245123000 |
| 1 | 2.384188000 | 7.425891000 | 15.661900000 |
| 1 | 3.863821000 | 7.938549000 | 14.835198000 |
| 6 | 1.103537000 | 4.010892000 | 19.963015000 |
| 7 | 2.085341000 | 3.003229000 | 19.600612000 |
| 6 | 3.411997000 | 3.221766000 | 19.575487000 |
| 7 | 3.930138000 | 4.426309000 | 19.834305000 |
| 7 | 4.236688000 | 2.191117000 | 19.349380000 |
| 1 | 1.374384000 | 4.492259000 | 20.908919000 |
| 1 | 0.999705000 | 4.779025000 | 19.185849000 |
| 1 | 1.786491000 | 2.071518000 | 19.328009000 |
| 1 | 3.351580000 | 5.245567000 | 19.720155000 |
| 1 | 4.953955000 | 4.552725000 | 19.683727000 |
| 1 | 3.804132000 | 1.303167000 | 19.117980000 |
| 1 | 5.134126000 | 2.409211000 | 18.883397000 |
| 26 | 7.505630000 | 1.158815000 | 17.070149000 |
| 6 | 8.550735000 | 4.075665000 | 17.987287000 |
| 6 | 8.613698000 | 5.057426000 | 16.815986000 |
| 6 | 8.552732000 | 6.530547000 | 17.229539000 |
| 6 | 7.093654000 | 3.873926000 | 18.427656000 |
| 8 | 9.300936000 | 6.911515000 | 18.172164000 |
| 8 | 7.843044000 | 7.289983000 | 16.509715000 |
| 8 | 6.658792000 | 4.648931000 | 19.301482000 |
| 8 | 6.427390000 | 2.956340000 | 17.841165000 |
| 16 | 9.392311000 | 2.494810000 | 17.519486000 |
| 1 | 9.126808000 | 4.447177000 | 18.832912000 |
| 1 | 9.570185000 | 4.955108000 | 16.292291000 |
| 1 | 7.827523000 | 4.829761000 | 16.100814000 |
| 8 | 8.121599000 | 1.835522000 | 15.410324000 |
| 8 | 12.954651000 | -0.927494000 | 17.262678000 |
| 1 | 12.647141000 | -0.340012000 | 16.537479000 |
| 1 | 13.800482000 | -1.287632000 | 16.954871000 |
| 8 | 11.978088000 | 1.276691000 | 15.832361000 |
| 1 | 11.108875000 | 1.661039000 | 15.603599000 |
| 1 | 12.228608000 | 1.658933000 | 16.722634000 |
| 8 | 11.155428000 | 2.978372000 | 20.013021000 |
| 1 | 11.270892000 | 3.881077000 | 19.601984000 |
| 1 | 11.482578000 | 3.082191000 | 20.920120000 |
| 8 | 11.390947000 | 5.323441000 | 18.748046000 |
| 1 | 10.583350000 | 5.896519000 | 18.619314000 |
| 1 | 12.128498000 | 5.878362000 | 18.410498000 |
| 1 | 9.783848000 | -3.350559000 | 20.551468000 |
| 1 | 6.969240000 | -3.881582000 | 19.887429000 |
| 1 | 1.506140000 | -2.886648000 | 18.141757000 |
| 1 | 0.356382000 | 0.847876000 | 15.780126000 |
| 1 | 0.136767000 | 3.521961000 | 20.091090000 |
| 1 | 5.249134000 | 7.798667000 | 16.976699000 |
| 1 | 18.211558000 | 2.593187000 | 11.254165000 |
| 1 | 15.053138000 | 6.043509000 | 8.857216000 |
| 1 | 10.595084000 | 2.385090000 | 6.098407000 |
| 1 | 5.414796000 | 1.179330000 | 7.502259000 |
| 1 | 9.850061000 | -1.483427000 | 9.758829000 |
| 1 | 5.582290000 | -3.150986000 | 21.461570000 |
| 1 | 9.211366000 | 9.172120000 | 10.201203000 |
| 1 | 14.025275000 | 8.172732000 | 9.264176000 |
| 1 | 16.164798000 | 8.527675000 | 13.747227000 |
| 1 | 1.696239000 | -0.439614000 | 10.100596000 |
| 1 | 4.655624000 | -3.895532000 | 12.410517000 |
| 1 | 10.857249000 | -2.404351000 | 12.629415000 |
| 8 | 7.248659000 | 6.262629000 | 14.038514000 |
| 1 | 7.351724000 | 6.629482000 | 14.950158000 |
| 1 | 6.740288000 | 5.434536000 | 14.144922000 |
| 8 | 12.890349000 | 1.778100000 | 18.256956000 |
| 1 | 12.276622000 | 2.129465000 | 18.951539000 |
| 1 | 12.975585000 | 0.815385000 | 18.392266000 |
| 8 | 9.382917000 | 2.515329000 | 15.814585000 |

**$^{3}$TS2$_{S}$:**

| | | | |
|---|---|---|---|
| 6 | 15.388813000 | 7.822073000 | 13.537753000 |
| 6 | 14.428091000 | 8.452352000 | 12.540092000 |
| 8 | 13.500331000 | 9.197211000 | 12.882383000 |
| 6 | 14.698538000 | 7.445298000 | 14.850827000 |
| 6 | 13.565501000 | 6.430618000 | 14.667448000 |
| 6 | 12.950318000 | 6.019720000 | 15.991235000 |
| 8 | 12.958715000 | 6.785982000 | 16.975885000 |
| 7 | 12.370898000 | 4.803966000 | 16.041122000 |
| 1 | 15.877890000 | 6.955048000 | 13.081150000 |
| 1 | 15.445765000 | 7.041090000 | 15.543224000 |
| 1 | 14.297686000 | 8.344516000 | 15.325176000 |
| 1 | 13.911626000 | 5.539396000 | 14.133529000 |
| 1 | 12.771072000 | 6.860505000 | 14.043057000 |
| 1 | 11.954522000 | 4.523515000 | 16.927729000 |
| 1 | 12.581568000 | 4.106815000 | 15.332272000 |
| 7 | 14.663202000 | 8.148627000 | 11.240669000 |
| 6 | 13.717964000 | 8.580869000 | 10.236528000 |
| 6 | 12.310473000 | 8.053327000 | 10.570654000 |
| 8 | 12.164301000 | 6.951756000 | 11.109073000 |
| 1 | 15.358208000 | 7.430378000 | 11.021511000 |
| 1 | 13.708394000 | 9.674019000 | 10.161239000 |
| 7 | 11.273714000 | 8.840677000 | 10.222065000 |
| 6 | 9.941656000 | 8.616699000 | 10.797272000 |
| 6 | 9.948750000 | 9.121128000 | 12.244640000 |
| 6 | 8.587965000 | 9.262528000 | 12.922164000 |
| 6 | 8.707155000 | 10.015244000 | 14.257218000 |
| 7 | 9.344426000 | 9.251803000 | 15.329047000 |
| 6 | 10.614530000 | 9.304780000 | 15.721204000 |
| 7 | 11.567769000 | 9.917927000 | 14.975201000 |
| 7 | 10.964389000 | 8.805207000 | 16.916699000 |
| 1 | 11.495562000 | 9.793549000 | 9.962356000 |
| 1 | 9.712987000 | 7.549860000 | 10.749787000 |
| 1 | 10.587054000 | 8.438829000 | 12.816612000 |
| 1 | 10.438378000 | 10.107424000 | 12.242133000 |
| 1 | 7.913504000 | 9.843926000 | 12.278251000 |
| 1 | 8.120451000 | 8.284040000 | 13.080212000 |
| 1 | 9.262097000 | 10.950316000 | 14.120782000 |
| 1 | 7.710394000 | 10.277267000 | 14.625666000 |
| 1 | 8.726655000 | 8.622622000 | 15.872471000 |
| 1 | 12.507510000 | 9.911155000 | 15.347244000 |
| 1 | 11.560085000 | 9.819632000 | 13.968367000 |
| 1 | 10.244695000 | 8.338907000 | 17.486591000 |
| 1 | 11.883118000 | 8.367459000 | 16.989201000 |
| 7 | 5.584502000 | 1.552887000 | 8.438285000 |
| 6 | 6.634268000 | 2.576324000 | 8.357411000 |

| | | | |
|---|---|---|---|
| 6 | 7.984193000 | 1.933379000 | 8.035872000 |
| 8 | 8.304864000 | 0.866680000 | 8.562079000 |
| 6 | 6.817778000 | 3.315620000 | 9.699526000 |
| 6 | 5.766989000 | 4.337306000 | 10.081700000 |
| 6 | 4.464649000 | 4.345592000 | 9.568571000 |
| 6 | 6.102252000 | 5.298597000 | 11.045179000 |
| 6 | 3.516639000 | 5.263883000 | 10.021611000 |
| 6 | 5.159798000 | 6.206324000 | 11.521024000 |
| 6 | 3.855907000 | 6.176461000 | 11.022077000 |
| 8 | 2.869723000 | 7.009371000 | 11.495601000 |
| 1 | 5.970942000 | 0.764752000 | 8.964593000 |
| 1 | 6.358080000 | 3.298114000 | 7.578796000 |
| 1 | 6.868007000 | 2.546354000 | 10.480789000 |
| 1 | 7.798997000 | 3.806678000 | 9.708256000 |
| 1 | 4.177804000 | 3.603568000 | 8.831662000 |
| 1 | 7.117713000 | 5.346503000 | 11.425500000 |
| 1 | 2.503860000 | 5.263784000 | 9.629933000 |
| 1 | 5.445211000 | 6.907435000 | 12.302221000 |
| 1 | 3.261167000 | 7.589675000 | 12.166955000 |
| 7 | 8.813343000 | 2.597238000 | 7.189543000 |
| 6 | 10.227886000 | 2.243680000 | 7.121527000 |
| 6 | 11.017312000 | 3.159571000 | 8.055662000 |
| 8 | 11.612525000 | 4.143437000 | 7.619718000 |
| 1 | 8.561847000 | 3.538961000 | 6.916227000 |
| 1 | 10.307795000 | 1.192431000 | 7.403477000 |
| 7 | 10.914795000 | 2.890565000 | 9.387595000 |
| 6 | 11.583053000 | 3.816404000 | 10.285758000 |
| 6 | 13.108417000 | 3.705219000 | 10.075546000 |
| 8 | 13.713147000 | 2.636538000 | 9.928024000 |
| 6 | 11.116179000 | 3.690918000 | 11.739745000 |
| 6 | 9.955767000 | 4.651939000 | 12.037076000 |
| 8 | 9.425258000 | 5.328294000 | 11.155175000 |
| 7 | 9.579064000 | 4.717900000 | 13.336416000 |
| 1 | 10.685728000 | 1.949556000 | 9.710732000 |
| 1 | 11.291809000 | 4.813525000 | 9.954860000 |
| 1 | 11.938631000 | 3.921214000 | 12.427210000 |
| 1 | 10.791363000 | 2.675009000 | 11.974497000 |
| 1 | 9.882245000 | 4.020239000 | 14.002960000 |
| 1 | 8.764426000 | 5.296565000 | 13.580307000 |
| 7 | 13.720717000 | 4.909608000 | 9.983437000 |
| 6 | 14.999970000 | 5.061130000 | 9.320375000 |
| 6 | 16.137307000 | 4.929192000 | 10.323967000 |
| 8 | 16.545748000 | 5.901009000 | 10.974318000 |
| 1 | 13.188496000 | 5.736792000 | 10.255280000 |
| 1 | 15.074860000 | 4.308094000 | 8.530289000 |
| 7 | 16.587086000 | 3.662280000 | 10.473099000 |
| 6 | 17.406298000 | 3.259370000 | 11.607723000 |
| 6 | 16.556369000 | 2.550100000 | 12.672440000 |
| 6 | 15.393309000 | 3.427030000 | 13.133143000 |
| 16 | 14.214009000 | 2.593676000 | 14.247007000 |
| 6 | 15.178817000 | 2.589230000 | 15.798101000 |
| 1 | 15.991494000 | 2.951524000 | 10.052269000 |
| 1 | 17.861629000 | 4.169067000 | 12.005735000 |
| 1 | 16.155392000 | 1.616245000 | 12.259394000 |
| 1 | 17.198476000 | 2.277406000 | 13.518505000 |
| 1 | 15.757875000 | 4.342022000 | 13.613153000 |
| 1 | 14.790073000 | 3.727701000 | 12.274240000 |
| 1 | 14.527201000 | 2.244882000 | 16.603453000 |
| 1 | 15.511667000 | 3.605130000 | 16.031918000 |
| 1 | 16.046841000 | 1.929610000 | 15.719300000 |
| 6 | 4.648765000 | -2.567777000 | 21.316213000 |
| 6 | 4.158137000 | -2.800429000 | 19.895809000 |
| 8 | 4.861754000 | -3.281352000 | 19.008973000 |
| 6 | 4.920502000 | -1.067487000 | 21.582924000 |
| 6 | 5.919715000 | -0.449735000 | 20.653716000 |
| 7 | 7.042853000 | 0.225956000 | 21.103115000 |
| 6 | 5.956694000 | -0.314407000 | 19.286942000 |
| 6 | 7.702419000 | 0.744873000 | 20.037597000 |
| 7 | 7.056626000 | 0.433346000 | 18.922889000 |
| 1 | 3.898254000 | -2.920411000 | 22.033281000 |
| 1 | 3.984048000 | -0.498061000 | 21.517814000 |
| 1 | 5.271112000 | -0.963086000 | 22.615546000 |
| 1 | 5.272402000 | -0.681425000 | 18.542309000 |
| 1 | 8.611520000 | 1.324447000 | 20.113081000 |
| 1 | 7.325685000 | 0.313355000 | 22.069222000 |
| 7 | 2.872825000 | -2.387861000 | 19.664107000 |
| 6 | 2.350865000 | -2.251116000 | 18.329167000 |
| 6 | 2.033116000 | -0.789200000 | 17.995759000 |
| 8 | 2.265261000 | 0.119982000 | 18.807959000 |
| 1 | 2.409365000 | -1.851705000 | 20.384955000 |
| 1 | 3.118092000 | -2.627969000 | 17.644643000 |
| 7 | 1.476496000 | -0.600949000 | 16.785448000 |
| 6 | 1.378472000 | 0.699946000 | 16.127120000 |
| 6 | 2.357551000 | 0.783401000 | 14.944398000 |
| 6 | 3.762381000 | 0.484259000 | 15.355425000 |
| 7 | 4.420146000 | -0.667310000 | 14.964010000 |
| 6 | 4.655825000 | 1.166883000 | 16.142329000 |
| 6 | 5.672323000 | -0.640939000 | 15.471190000 |
| 7 | 5.837596000 | 0.453591000 | 16.207437000 |
| 1 | 1.388169000 | -1.414058000 | 16.186019000 |
| 1 | 1.615655000 | 1.454329000 | 16.878610000 |
| 1 | 2.062365000 | 0.076055000 | 14.161982000 |
| 1 | 2.304733000 | 1.783433000 | 14.502842000 |
| 1 | 4.567875000 | 2.132802000 | 16.607271000 |
| 1 | 6.430640000 | -1.385664000 | 15.291786000 |
| 1 | 4.034977000 | -1.339571000 | 14.285783000 |
| 6 | 11.030292000 | -3.187623000 | 13.374850000 |
| 6 | 9.697724000 | -3.508054000 | 14.068853000 |
| 6 | 12.130965000 | -2.733427000 | 14.338150000 |
| 6 | 9.752113000 | -4.734584000 | 14.983737000 |
| 1 | 11.369270000 | -4.072612000 | 12.819520000 |
| 1 | 8.935959000 | -3.687809000 | 13.295586000 |
| 1 | 9.362588000 | -2.629922000 | 14.637647000 |
| 1 | 12.387238000 | -3.512856000 | 15.065263000 |
| 1 | 11.811182000 | -1.847767000 | 14.896198000 |
| 1 | 13.044042000 | -2.473701000 | 13.791219000 |
| 1 | 8.780233000 | -4.925634000 | 15.451863000 |
| 1 | 10.480812000 | -4.603409000 | 15.789797000 |
| 1 | 10.039047000 | -5.630121000 | 14.419738000 |
| 7 | 7.687051000 | -3.390903000 | 20.366707000 |
| 6 | 8.983665000 | -3.995991000 | 20.076518000 |
| 6 | 9.311019000 | -4.163527000 | 18.567879000 |
| 6 | 9.228599000 | -2.872509000 | 17.821706000 |
| 7 | 10.322325000 | -2.081658000 | 17.509195000 |
| 6 | 8.138860000 | -2.139198000 | 17.404527000 |
| 6 | 9.872582000 | -0.941934000 | 16.928698000 |
| 7 | 8.551983000 | -0.937693000 | 16.853557000 |
| 1 | 7.658664000 | -2.456606000 | 19.960792000 |
| 1 | 9.025252000 | -4.982450000 | 20.552010000 |
| 1 | 8.597103000 | -4.872804000 | 18.132536000 |
| 1 | 10.305834000 | -4.611665000 | 18.450890000 |
| 1 | 7.093493000 | -2.396711000 | 17.493312000 |
| 1 | 10.529842000 | -0.161544000 | 16.583356000 |
| 1 | 11.313115000 | -2.258403000 | 17.648146000 |
| 7 | 8.891573000 | -1.804405000 | 9.677697000 |
| 6 | 8.234392000 | -1.603549000 | 10.958648000 |
| 6 | 6.744271000 | -1.879255000 | 10.789624000 |
| 8 | 6.085373000 | -1.453777000 | 9.836446000 |
| 6 | 8.387254000 | -0.191612000 | 11.603254000 |
| 6 | 9.798882000 | 0.125836000 | 11.971107000 |
| 7 | 10.812175000 | 0.252284000 | 11.029795000 |
| 6 | 10.323826000 | 0.285595000 | 13.234101000 |
| 6 | 11.924742000 | 0.472210000 | 11.716482000 |
| 7 | 11.678058000 | 0.497981000 | 13.054647000 |
| 1 | 8.469900000 | -1.156070000 | 9.013360000 |
| 1 | 8.644867000 | -2.330897000 | 11.669391000 |
| 1 | 7.999097000 | 0.542238000 | 10.892366000 |
| 1 | 7.779106000 | -0.152281000 | 12.513176000 |
| 1 | 9.849125000 | 0.271850000 | 14.202329000 |
| 1 | 12.888938000 | 0.699128000 | 11.283855000 |

| | | | |
|---|---|---|---|
| 1 | 12.339362000 | 0.725462000 | 13.790201000 |
| 7 | 6.148605000 | -2.581834000 | 11.799878000 |
| 6 | 4.791947000 | -3.100275000 | 11.652969000 |
| 6 | 3.690518000 | -2.056873000 | 11.905534000 |
| 8 | 3.087895000 | -2.024101000 | 12.997045000 |
| 1 | 6.750825000 | -3.025856000 | 12.479102000 |
| 1 | 4.674989000 | -3.517908000 | 10.646493000 |
| 7 | 3.428865000 | -1.253859000 | 10.868691000 |
| 6 | 2.419011000 | -0.200778000 | 10.863609000 |
| 6 | 3.000353000 | 1.197869000 | 10.604413000 |
| 6 | 3.833727000 | 1.780266000 | 11.720449000 |
| 6 | 5.073786000 | 1.238241000 | 12.079888000 |
| 6 | 3.426047000 | 2.955672000 | 12.364613000 |
| 6 | 5.874141000 | 1.838818000 | 13.049664000 |
| 6 | 4.218870000 | 3.574779000 | 13.323999000 |
| 6 | 5.445348000 | 3.009192000 | 13.675518000 |
| 8 | 6.205455000 | 3.636274000 | 14.625781000 |
| 1 | 4.102856000 | -1.293775000 | 10.103949000 |
| 1 | 1.903129000 | -0.246924000 | 11.824098000 |
| 1 | 2.153538000 | 1.865499000 | 10.406929000 |
| 1 | 3.601119000 | 1.185133000 | 9.684951000 |
| 1 | 5.442691000 | 0.360129000 | 11.565136000 |
| 1 | 2.491961000 | 3.426600000 | 12.073529000 |
| 1 | 6.839399000 | 1.425852000 | 13.315684000 |
| 1 | 3.919687000 | 4.513392000 | 13.769114000 |
| 1 | 6.941028000 | 3.033250000 | 14.905780000 |
| 6 | 4.637274000 | 6.935061000 | 17.281211000 |
| 6 | 4.106980000 | 6.158668000 | 16.070201000 |
| 6 | 3.344321000 | 4.905979000 | 16.520019000 |
| 6 | 3.247120000 | 7.051683000 | 15.167338000 |
| 1 | 3.804516000 | 7.297550000 | 17.902770000 |
| 1 | 5.289518000 | 6.311527000 | 17.900673000 |
| 1 | 4.972716000 | 5.826184000 | 15.487827000 |
| 1 | 2.924176000 | 4.355732000 | 15.671887000 |
| 1 | 2.511159000 | 5.174048000 | 17.186044000 |
| 1 | 4.017986000 | 4.233401000 | 17.058204000 |
| 1 | 2.917412000 | 6.523753000 | 14.263799000 |
| 1 | 2.348955000 | 7.399682000 | 15.694410000 |
| 1 | 3.811714000 | 7.939277000 | 14.855295000 |
| 6 | 1.133239000 | 4.029958000 | 19.981246000 |
| 7 | 2.107418000 | 3.019312000 | 19.606862000 |
| 6 | 3.434858000 | 3.231850000 | 19.572837000 |
| 7 | 3.960691000 | 4.432255000 | 19.834671000 |
| 7 | 4.252706000 | 2.198508000 | 19.334735000 |
| 1 | 1.413193000 | 4.505493000 | 20.927460000 |
| 1 | 1.027551000 | 4.802316000 | 19.208530000 |
| 1 | 1.802088000 | 2.089906000 | 19.333724000 |
| 1 | 3.385354000 | 5.255008000 | 19.729850000 |
| 1 | 4.983959000 | 4.553945000 | 19.675920000 |
| 1 | 3.814253000 | 1.313971000 | 19.101553000 |
| 1 | 5.151372000 | 2.412911000 | 18.868834000 |
| 26 | 7.520701000 | 1.135909000 | 17.047220000 |
| 6 | 8.576800000 | 4.072311000 | 17.969583000 |
| 6 | 8.627626000 | 5.055557000 | 16.799461000 |
| 6 | 8.560035000 | 6.527282000 | 17.215585000 |
| 6 | 7.122116000 | 3.862244000 | 18.413656000 |
| 8 | 9.309844000 | 6.911123000 | 18.155846000 |
| 8 | 7.842697000 | 7.284645000 | 16.500798000 |
| 8 | 6.683613000 | 4.640413000 | 19.282933000 |
| 8 | 6.459377000 | 2.937056000 | 17.834810000 |
| 16 | 9.428516000 | 2.499692000 | 17.496913000 |
| 1 | 9.149704000 | 4.447985000 | 18.815426000 |
| 1 | 9.581018000 | 4.958600000 | 16.269060000 |
| 1 | 7.839385000 | 4.823724000 | 16.087849000 |
| 8 | 8.058805000 | 1.765675000 | 15.416915000 |
| 8 | 12.958284000 | -0.971945000 | 17.308101000 |
| 1 | 12.661504000 | -0.387099000 | 16.576309000 |
| 1 | 13.807019000 | -1.335368000 | 17.012425000 |
| 8 | 12.012638000 | 1.227476000 | 15.851900000 |
| 1 | 11.152434000 | 1.631485000 | 15.621233000 |
| 1 | 12.274310000 | 1.619234000 | 16.734652000 |
| 8 | 11.187142000 | 2.985145000 | 20.003926000 |
| 1 | 11.297779000 | 3.887136000 | 19.590706000 |
| 1 | 11.516023000 | 3.093655000 | 20.909874000 |
| 8 | 11.408917000 | 5.338598000 | 18.744440000 |
| 1 | 10.597408000 | 5.905051000 | 18.611338000 |
| 1 | 12.143812000 | 5.900200000 | 18.411718000 |
| 1 | 9.759812000 | -3.381615000 | 20.549365000 |
| 1 | 6.942221000 | -3.890957000 | 19.879329000 |
| 1 | 1.449894000 | -2.862245000 | 18.185766000 |
| 1 | 0.349505000 | 0.863696000 | 15.790029000 |
| 1 | 0.165021000 | 3.545154000 | 20.113965000 |
| 1 | 5.230730000 | 7.800100000 | 16.971776000 |
| 1 | 18.208666000 | 2.599731000 | 11.260792000 |
| 1 | 15.049914000 | 6.051964000 | 8.863431000 |
| 1 | 10.601588000 | 2.389686000 | 6.106806000 |
| 1 | 5.420078000 | 1.181376000 | 7.502159000 |
| 1 | 9.854323000 | -1.480200000 | 9.775783000 |
| 1 | 5.571635000 | -3.132973000 | 21.458354000 |
| 1 | 9.215767000 | 9.161279000 | 10.185244000 |
| 1 | 14.032523000 | 8.182493000 | 9.266656000 |
| 1 | 16.180030000 | 8.554707000 | 13.744557000 |
| 1 | 1.688949000 | -0.437917000 | 10.079654000 |
| 1 | 4.647018000 | -3.894429000 | 12.385972000 |
| 1 | 10.867972000 | -2.397852000 | 12.633572000 |
| 8 | 7.225410000 | 6.245676000 | 14.040564000 |
| 1 | 7.340050000 | 6.617230000 | 14.949095000 |
| 1 | 6.710449000 | 5.423383000 | 14.156375000 |
| 8 | 12.925274000 | 1.751100000 | 18.272931000 |
| 1 | 12.307671000 | 2.119818000 | 18.955080000 |
| 1 | 12.994559000 | 0.789248000 | 18.421616000 |
| 8 | 9.453662000 | 2.523512000 | 15.823953000 |

**$^{3}$IM2$_{S}$:**

| | | | |
|---|---|---|---|
| 6 | 14.559061000 | 9.361971000 | 13.094715000 |
| 6 | 13.401054000 | 9.476964000 | 12.118652000 |
| 8 | 12.285055000 | 9.904650000 | 12.447887000 |
| 6 | 14.148373000 | 8.623214000 | 14.373597000 |
| 6 | 13.606470000 | 7.213051000 | 14.111037000 |
| 6 | 13.364229000 | 6.462955000 | 15.411672000 |
| 8 | 13.088842000 | 7.067955000 | 16.461051000 |
| 7 | 13.434781000 | 5.113478000 | 15.369954000 |
| 1 | 15.406700000 | 8.861152000 | 12.618657000 |
| 1 | 15.012856000 | 8.565862000 | 15.043392000 |
| 1 | 13.386084000 | 9.198812000 | 14.901282000 |
| 1 | 14.297490000 | 6.650299000 | 13.472223000 |
| 1 | 12.661634000 | 7.262118000 | 13.552457000 |
| 1 | 13.503443000 | 4.595489000 | 16.253321000 |
| 1 | 13.848401000 | 4.629298000 | 14.578000000 |
| 7 | 13.661138000 | 9.063524000 | 10.856274000 |
| 6 | 12.575314000 | 8.993416000 | 9.900429000 |
| 6 | 11.461526000 | 8.079073000 | 10.441704000 |
| 8 | 11.750693000 | 7.008206000 | 10.987684000 |
| 1 | 14.547978000 | 8.584131000 | 10.678221000 |
| 1 | 12.190486000 | 9.996650000 | 9.689008000 |
| 7 | 10.196398000 | 8.509854000 | 10.269364000 |
| 6 | 9.098223000 | 8.000718000 | 11.099605000 |
| 6 | 9.156117000 | 8.706301000 | 12.462638000 |
| 6 | 7.877760000 | 8.668376000 | 13.294233000 |
| 6 | 8.029210000 | 9.490860000 | 14.582697000 |
| 7 | 8.955568000 | 8.888357000 | 15.536273000 |
| 6 | 10.201499000 | 9.263804000 | 15.798447000 |
| 7 | 10.853677000 | 10.171637000 | 15.021869000 |
| 7 | 10.815561000 | 8.803048000 | 16.900870000 |
| 1 | 10.099974000 | 9.474225000 | 9.976470000 |
| 1 | 9.208716000 | 6.919206000 | 11.195400000 |
| 1 | 9.984025000 | 8.261889000 | 13.025532000 |
| 1 | 9.425875000 | 9.756883000 | 12.282412000 |
| 1 | 7.044241000 | 9.095541000 | 12.718652000 |
| 1 | 7.619568000 | 7.633514000 | 13.542704000 |

| | | | |
|---|---|---|---|
| 1 | 8.372800000 | 10.506632000 | 14.361967000 |
| 1 | 7.061864000 | 9.571999000 | 15.090992000 |
| 1 | 8.629065000 | 8.034853000 | 16.032875000 |
| 1 | 11.744606000 | 10.480861000 | 15.385732000 |
| 1 | 10.870570000 | 10.067701000 | 14.010440000 |
| 1 | 10.291028000 | 8.159155000 | 17.513005000 |
| 1 | 11.802133000 | 8.559811000 | 16.814011000 |
| 7 | 6.093572000 | 0.949066000 | 8.240938000 |
| 6 | 7.236753000 | 1.863282000 | 8.304623000 |
| 6 | 8.567612000 | 1.114640000 | 8.093340000 |
| 8 | 8.695178000 | -0.068073000 | 8.411484000 |
| 6 | 7.312090000 | 2.528138000 | 9.699167000 |
| 6 | 6.408897000 | 3.713142000 | 9.969933000 |
| 6 | 5.179116000 | 3.921623000 | 9.334417000 |
| 6 | 6.803124000 | 4.619775000 | 10.963929000 |
| 6 | 4.357453000 | 4.994161000 | 9.687631000 |
| 6 | 5.987069000 | 5.685878000 | 11.331786000 |
| 6 | 4.754476000 | 5.867756000 | 10.701964000 |
| 8 | 3.891589000 | 6.878228000 | 11.061674000 |
| 1 | 6.361659000 | 0.137267000 | 8.803277000 |
| 1 | 7.120759000 | 2.637517000 | 7.535569000 |
| 1 | 7.078420000 | 1.741843000 | 10.425619000 |
| 1 | 8.341653000 | 2.835427000 | 9.910919000 |
| 1 | 4.849206000 | 3.221833000 | 8.573279000 |
| 1 | 7.759979000 | 4.484163000 | 11.458017000 |
| 1 | 3.400893000 | 5.153292000 | 9.198985000 |
| 1 | 6.303369000 | 6.349666000 | 12.132783000 |
| 1 | 4.322049000 | 7.409746000 | 11.749152000 |
| 7 | 9.601718000 | 1.853279000 | 7.620494000 |
| 6 | 10.991393000 | 1.440197000 | 7.802853000 |
| 6 | 11.699618000 | 2.667214000 | 8.358934000 |
| 8 | 11.846500000 | 3.664858000 | 7.650298000 |
| 1 | 9.459810000 | 2.832715000 | 7.406019000 |
| 1 | 11.002135000 | 0.580586000 | 8.475759000 |
| 7 | 12.027860000 | 2.656499000 | 9.670291000 |
| 6 | 12.394388000 | 3.931199000 | 10.250702000 |
| 6 | 13.784657000 | 4.360109000 | 9.759441000 |
| 8 | 14.720700000 | 3.583595000 | 9.554110000 |
| 6 | 12.290585000 | 3.951192000 | 11.777107000 |
| 6 | 10.833605000 | 4.153630000 | 12.209527000 |
| 8 | 9.908466000 | 3.988315000 | 11.413623000 |
| 7 | 10.642033000 | 4.537056000 | 13.489696000 |
| 1 | 11.842497000 | 1.845360000 | 10.254304000 |
| 1 | 11.684257000 | 4.669246000 | 9.872496000 |
| 1 | 12.883517000 | 4.787014000 | 12.162680000 |
| 1 | 12.695187000 | 3.036998000 | 12.220984000 |
| 1 | 11.419250000 | 4.687178000 | 14.119351000 |
| 1 | 9.688560000 | 4.811044000 | 13.769156000 |
| 7 | 13.879784000 | 5.701280000 | 9.576617000 |
| 6 | 15.078197000 | 6.296399000 | 9.024786000 |
| 6 | 16.124119000 | 6.485371000 | 10.119806000 |
| 8 | 16.152078000 | 7.513157000 | 10.812698000 |
| 1 | 13.146157000 | 6.285717000 | 9.979074000 |
| 1 | 15.449255000 | 5.647756000 | 8.226683000 |
| 7 | 16.937540000 | 5.420754000 | 10.284177000 |
| 6 | 17.836595000 | 5.263869000 | 11.419063000 |
| 6 | 17.440084000 | 4.059546000 | 12.284802000 |
| 6 | 16.007837000 | 4.188693000 | 12.799147000 |
| 16 | 15.419433000 | 2.799140000 | 13.833458000 |
| 6 | 16.444097000 | 3.050605000 | 15.323236000 |
| 1 | 16.639021000 | 4.578787000 | 9.794110000 |
| 1 | 17.790373000 | 6.195204000 | 11.989574000 |
| 1 | 17.530363000 | 3.133189000 | 11.704760000 |
| 1 | 18.149155000 | 3.985706000 | 13.117187000 |
| 1 | 15.889926000 | 5.123487000 | 13.361549000 |
| 1 | 15.317040000 | 4.211578000 | 11.954247000 |
| 1 | 16.145938000 | 2.297410000 | 16.054726000 |
| 1 | 16.263989000 | 4.041026000 | 15.752557000 |
| 1 | 17.506539000 | 2.929032000 | 15.098687000 |
| 6 | 4.745749000 | -3.211372000 | 20.680794000 |

| | | | |
|---|---|---|---|
| 6 | 4.166609000 | -2.884926000 | 19.313485000 |
| 8 | 4.830433000 | -2.965570000 | 18.278134000 |
| 6 | 5.140245000 | -1.936343000 | 21.469215000 |
| 6 | 6.139829000 | -1.081827000 | 20.753431000 |
| 7 | 7.368119000 | -0.715550000 | 21.284303000 |
| 6 | 6.113409000 | -0.505246000 | 19.508971000 |
| 6 | 8.027385000 | 0.045870000 | 20.373069000 |
| 7 | 7.287417000 | 0.183918000 | 19.284010000 |
| 1 | 4.015041000 | -3.774550000 | 21.272972000 |
| 1 | 4.241699000 | -1.343076000 | 21.681802000 |
| 1 | 5.541819000 | -2.244183000 | 22.440928000 |
| 1 | 5.360284000 | -0.557354000 | 18.744478000 |
| 1 | 9.016479000 | 0.450380000 | 20.523126000 |
| 1 | 7.710383000 | -0.958678000 | 22.203870000 |
| 7 | 2.870099000 | -2.460181000 | 19.302527000 |
| 6 | 2.215101000 | -2.055110000 | 18.086570000 |
| 6 | 2.007783000 | -0.539882000 | 17.963173000 |
| 8 | 2.429420000 | 0.255293000 | 18.811852000 |
| 1 | 2.386998000 | -2.331468000 | 20.180047000 |
| 1 | 2.856669000 | -2.377217000 | 17.261685000 |
| 7 | 1.319999000 | -0.187418000 | 16.853955000 |
| 6 | 1.348073000 | 1.151539000 | 16.267507000 |
| 6 | 2.396582000 | 1.235092000 | 15.139193000 |
| 6 | 3.734595000 | 0.771313000 | 15.610618000 |
| 7 | 4.239092000 | -0.485820000 | 15.318420000 |
| 6 | 4.657938000 | 1.350900000 | 16.443863000 |
| 6 | 5.438910000 | -0.608694000 | 15.931571000 |
| 7 | 5.712421000 | 0.481321000 | 16.638000000 |
| 1 | 1.092819000 | -0.939457000 | 16.212824000 |
| 1 | 1.600262000 | 1.849384000 | 17.066383000 |
| 1 | 2.077534000 | 0.627206000 | 14.284368000 |
| 1 | 2.461998000 | 2.265598000 | 14.776568000 |
| 1 | 4.675542000 | 2.346369000 | 16.853627000 |
| 1 | 6.063521000 | -1.482322000 | 15.861421000 |
| 1 | 3.891282000 | -1.114791000 | 14.581432000 |
| 6 | 5.279418000 | -5.510529000 | 14.799487000 |
| 6 | 3.748723000 | -5.495589000 | 14.927814000 |
| 6 | 5.929400000 | -4.191300000 | 15.231869000 |
| 6 | 3.260588000 | -5.459483000 | 16.379714000 |
| 1 | 5.692900000 | -6.339877000 | 15.391213000 |
| 1 | 3.338442000 | -6.384253000 | 14.431904000 |
| 1 | 3.359629000 | -4.624270000 | 14.382867000 |
| 1 | 5.817432000 | -4.008183000 | 16.301992000 |
| 1 | 5.455664000 | -3.351877000 | 14.711564000 |
| 1 | 7.003427000 | -4.179690000 | 15.006575000 |
| 1 | 2.165654000 | -5.426842000 | 16.428106000 |
| 1 | 3.653277000 | -4.589754000 | 16.914821000 |
| 1 | 3.587008000 | -6.354761000 | 16.924257000 |
| 7 | 7.187274000 | -4.848620000 | 18.720041000 |
| 6 | 8.636003000 | -4.817003000 | 18.544873000 |
| 6 | 9.181715000 | -4.430196000 | 17.150333000 |
| 6 | 8.979702000 | -2.981752000 | 16.831894000 |
| 7 | 9.468621000 | -2.360791000 | 15.688114000 |
| 6 | 8.359880000 | -1.977804000 | 17.529715000 |
| 6 | 9.153439000 | -1.040370000 | 15.729240000 |
| 7 | 8.476661000 | -0.781876000 | 16.836793000 |
| 1 | 6.725905000 | -3.992856000 | 18.417789000 |
| 1 | 9.022916000 | -5.809826000 | 18.795887000 |
| 1 | 8.705443000 | -5.061659000 | 16.387064000 |
| 1 | 10.256538000 | -4.659981000 | 17.115910000 |
| 1 | 7.828829000 | -2.037618000 | 18.461708000 |
| 1 | 9.435508000 | -0.322530000 | 14.976865000 |
| 1 | 10.027661000 | -2.795402000 | 14.965939000 |
| 7 | 9.587977000 | -1.975794000 | 10.728646000 |
| 6 | 8.777975000 | -1.269403000 | 11.723149000 |
| 6 | 7.325256000 | -1.622770000 | 11.396850000 |
| 8 | 6.758522000 | -1.261313000 | 10.364580000 |
| 6 | 9.031366000 | 0.265532000 | 11.785409000 |
| 6 | 10.382103000 | 0.563974000 | 12.356537000 |
| 7 | 11.548049000 | 0.199108000 | 11.692097000 |

| | | | |
|---|---|---|---|
| 6 | 10.705262000 | 1.091544000 | 13.591201000 |
| 6 | 12.540879000 | 0.478032000 | 12.528020000 |
| 7 | 12.082424000 | 1.024892000 | 13.682520000 |
| 1 | 9.314669000 | -1.610415000 | 9.812814000 |
| 1 | 9.013514000 | -1.688699000 | 12.714247000 |
| 1 | 8.949236000 | 0.642437000 | 10.766793000 |
| 1 | 8.269428000 | 0.771064000 | 12.382871000 |
| 1 | 10.104911000 | 1.552396000 | 14.363995000 |
| 1 | 13.593313000 | 0.324824000 | 12.333221000 |
| 1 | 12.637960000 | 1.231383000 | 14.523890000 |
| 7 | 6.663513000 | -2.404695000 | 12.300878000 |
| 6 | 5.403224000 | -3.037647000 | 11.923765000 |
| 6 | 4.192883000 | -2.108375000 | 12.102717000 |
| 8 | 3.538504000 | -2.099593000 | 13.163013000 |
| 1 | 7.164745000 | -2.776666000 | 13.093783000 |
| 1 | 5.475024000 | -3.362824000 | 10.881110000 |
| 7 | 3.922227000 | -1.354694000 | 11.028115000 |
| 6 | 2.897253000 | -0.317538000 | 10.956378000 |
| 6 | 3.488354000 | 1.075412000 | 10.672713000 |
| 6 | 4.236867000 | 1.687816000 | 11.832112000 |
| 6 | 5.349299000 | 1.054406000 | 12.399055000 |
| 6 | 3.869310000 | 2.935817000 | 12.346747000 |
| 6 | 6.056323000 | 1.620780000 | 13.452426000 |
| 6 | 4.582300000 | 3.530666000 | 13.383145000 |
| 6 | 5.671648000 | 2.865392000 | 13.949509000 |
| 8 | 6.344913000 | 3.465499000 | 14.976561000 |
| 1 | 4.649469000 | -1.346919000 | 10.315587000 |
| 1 | 2.353298000 | -0.338473000 | 11.903421000 |
| 1 | 2.659268000 | 1.733988000 | 10.389193000 |
| 1 | 4.145366000 | 1.016749000 | 9.792378000 |
| 1 | 5.688787000 | 0.117807000 | 11.989715000 |
| 1 | 3.038841000 | 3.475197000 | 11.901425000 |
| 1 | 6.896272000 | 1.110232000 | 13.909047000 |
| 1 | 4.323332000 | 4.520308000 | 13.734126000 |
| 1 | 7.028441000 | 2.833100000 | 15.340936000 |
| 6 | 5.263498000 | 7.125524000 | 17.086976000 |
| 6 | 4.712625000 | 6.381219000 | 15.864957000 |
| 6 | 3.541395000 | 5.474730000 | 16.269013000 |
| 6 | 4.295049000 | 7.358249000 | 14.758288000 |
| 1 | 4.478249000 | 7.727190000 | 17.567188000 |
| 1 | 5.679509000 | 6.428357000 | 17.821790000 |
| 1 | 5.514515000 | 5.745440000 | 15.475750000 |
| 1 | 3.101403000 | 4.963800000 | 15.406591000 |
| 1 | 2.744039000 | 6.063966000 | 16.745731000 |
| 1 | 3.874892000 | 4.705105000 | 16.973265000 |
| 1 | 3.914391000 | 6.828967000 | 13.875319000 |
| 1 | 3.503566000 | 8.036981000 | 15.102600000 |
| 1 | 5.147417000 | 7.972660000 | 14.443743000 |
| 6 | 1.582957000 | 4.265443000 | 19.503337000 |
| 7 | 2.512604000 | 3.151907000 | 19.404038000 |
| 6 | 3.846142000 | 3.276957000 | 19.502547000 |
| 7 | 4.432272000 | 4.471043000 | 19.601454000 |
| 7 | 4.600304000 | 2.166403000 | 19.562097000 |
| 1 | 1.771794000 | 4.848702000 | 20.411206000 |
| 1 | 1.644263000 | 4.926788000 | 18.630603000 |
| 1 | 2.169809000 | 2.207581000 | 19.240919000 |
| 1 | 3.921694000 | 5.302997000 | 19.346053000 |
| 1 | 5.470864000 | 4.523583000 | 19.569363000 |
| 1 | 4.122076000 | 1.291406000 | 19.376192000 |
| 1 | 5.548698000 | 2.249153000 | 19.177365000 |
| 26 | 7.734575000 | 1.072463000 | 17.316112000 |
| 6 | 9.204798000 | 3.648089000 | 18.412245000 |
| 6 | 9.532085000 | 4.453078000 | 17.151704000 |
| 6 | 9.176709000 | 5.935964000 | 17.335748000 |
| 6 | 7.700000000 | 3.630231000 | 18.687757000 |
| 8 | 9.751262000 | 6.587506000 | 18.246497000 |
| 8 | 8.340072000 | 6.435273000 | 16.526124000 |
| 8 | 7.210589000 | 4.506175000 | 19.409606000 |
| 8 | 6.984015000 | 2.702958000 | 18.124426000 |
| 16 | 9.836720000 | 1.902553000 | 18.264655000 |
| 1 | 9.722649000 | 4.084212000 | 19.268159000 |
| 1 | 10.610272000 | 4.377182000 | 16.975259000 |
| 1 | 9.016177000 | 4.031714000 | 16.289539000 |
| 8 | 8.165855000 | 1.716866000 | 15.871983000 |
| 8 | 15.378818000 | 1.070506000 | 18.022246000 |
| 1 | 14.701471000 | 0.773849000 | 17.371237000 |
| 1 | 16.197706000 | 0.611783000 | 17.784815000 |
| 8 | 13.301909000 | 1.148974000 | 16.219591000 |
| 1 | 12.433903000 | 0.983362000 | 16.644243000 |
| 1 | 13.467770000 | 2.080399000 | 16.529856000 |
| 8 | 12.220893000 | 3.566129000 | 19.599086000 |
| 1 | 12.241134000 | 4.565319000 | 19.420079000 |
| 1 | 12.691339000 | 3.456013000 | 20.439843000 |
| 8 | 12.265400000 | 6.091561000 | 19.007321000 |
| 1 | 11.310297000 | 6.281614000 | 18.824908000 |
| 1 | 12.713680000 | 6.308276000 | 18.163776000 |
| 1 | 9.059715000 | -4.126436000 | 19.285153000 |
| 1 | 6.776594000 | -5.598309000 | 18.167010000 |
| 1 | 1.250308000 | -2.566402000 | 17.980332000 |
| 1 | 0.349249000 | 1.402035000 | 15.896553000 |
| 1 | 0.570415000 | 3.863396000 | 19.559213000 |
| 1 | 6.081927000 | 7.795681000 | 16.809995000 |
| 1 | 18.864963000 | 5.141625000 | 11.059499000 |
| 1 | 14.838335000 | 7.272073000 | 8.596693000 |
| 1 | 11.451976000 | 1.159288000 | 6.850023000 |
| 1 | 6.007929000 | 0.596157000 | 7.287304000 |
| 1 | 10.555885000 | -1.674814000 | 10.859674000 |
| 1 | 5.619586000 | -3.847720000 | 20.514035000 |
| 1 | 8.157256000 | 8.198363000 | 10.576886000 |
| 1 | 12.960074000 | 8.577755000 | 8.964424000 |
| 1 | 14.878675000 | 10.378665000 | 13.354946000 |
| 1 | 2.195344000 | -0.587069000 | 10.157893000 |
| 1 | 5.246074000 | -3.909045000 | 12.555471000 |
| 1 | 5.549843000 | -5.726295000 | 13.757005000 |
| 8 | 8.030139000 | 5.479195000 | 14.060973000 |
| 1 | 8.110105000 | 5.813398000 | 14.989677000 |
| 1 | 7.388725000 | 4.748311000 | 14.153829000 |
| 8 | 13.944579000 | 3.378623000 | 17.567156000 |
| 1 | 13.292320000 | 3.336625000 | 18.323396000 |
| 1 | 14.725706000 | 2.868051000 | 17.867910000 |
| 8 | 10.897231000 | 1.876734000 | 17.169050000 |

**$^3Prod_S$:**

| | | | |
|---|---|---|---|
| 6 | 15.142461000 | 8.770845000 | 13.214558000 |
| 6 | 14.188252000 | 9.172060000 | 12.102925000 |
| 8 | 13.164000000 | 9.836594000 | 12.313210000 |
| 6 | 14.374698000 | 8.023750000 | 14.312388000 |
| 6 | 13.669957000 | 6.760727000 | 13.794462000 |
| 6 | 12.768427000 | 6.180597000 | 14.862400000 |
| 8 | 11.954156000 | 6.914405000 | 15.455562000 |
| 7 | 12.888998000 | 4.876159000 | 15.154985000 |
| 1 | 15.954260000 | 8.153024000 | 12.820741000 |
| 1 | 15.062712000 | 7.760182000 | 15.124612000 |
| 1 | 13.612044000 | 8.681772000 | 14.733431000 |
| 1 | 14.397938000 | 6.027506000 | 13.434494000 |
| 1 | 13.029611000 | 7.011617000 | 12.940140000 |
| 1 | 12.209251000 | 4.441547000 | 15.778124000 |
| 1 | 13.518128000 | 4.246704000 | 14.663637000 |
| 7 | 14.508139000 | 8.716446000 | 10.866343000 |
| 6 | 13.542765000 | 8.863955000 | 9.795462000 |
| 6 | 12.221496000 | 8.187377000 | 10.193995000 |
| 8 | 12.236522000 | 7.068591000 | 10.722071000 |
| 1 | 15.244488000 | 8.009662000 | 10.790825000 |
| 1 | 13.390793000 | 9.923288000 | 9.565027000 |
| 7 | 11.087583000 | 8.863347000 | 9.941053000 |
| 6 | 9.837909000 | 8.519736000 | 10.629418000 |
| 6 | 9.867112000 | 9.122537000 | 12.041784000 |
| 6 | 8.500842000 | 9.287945000 | 12.702704000 |
| 6 | 8.587886000 | 10.144568000 | 13.973220000 |
| 7 | 9.234099000 | 9.476767000 | 15.100384000 |

| | | | |
|---|---|---|---|
| 6 | 10.469453000 | 9.659690000 | 15.543488000 |
| 7 | 11.406043000 | 10.307458000 | 14.803912000 |
| 7 | 10.784108000 | 9.293673000 | 16.802769000 |
| 1 | 11.199996000 | 9.840087000 | 9.699457000 |
| 1 | 9.740756000 | 7.432072000 | 10.653071000 |
| 1 | 10.523931000 | 8.504492000 | 12.665018000 |
| 1 | 10.339083000 | 10.112792000 | 11.966202000 |
| 1 | 7.818903000 | 9.801114000 | 12.009954000 |
| 1 | 8.056202000 | 8.313997000 | 12.934324000 |
| 1 | 9.124521000 | 11.078133000 | 13.772344000 |
| 1 | 7.580701000 | 10.409132000 | 14.312641000 |
| 1 | 8.641476000 | 8.818608000 | 15.644622000 |
| 1 | 12.286794000 | 10.483326000 | 15.266728000 |
| 1 | 11.517514000 | 10.109280000 | 13.813795000 |
| 1 | 10.061979000 | 8.808301000 | 17.354653000 |
| 1 | 11.700633000 | 8.878971000 | 16.926011000 |
| 7 | 5.623418000 | 1.212563000 | 8.537128000 |
| 6 | 6.747680000 | 2.144037000 | 8.407961000 |
| 6 | 8.057148000 | 1.385170000 | 8.122798000 |
| 8 | 8.194057000 | 0.210163000 | 8.468457000 |
| 6 | 6.959957000 | 2.934182000 | 9.720520000 |
| 6 | 5.985559000 | 4.050727000 | 10.036641000 |
| 6 | 4.677993000 | 4.112477000 | 9.540746000 |
| 6 | 6.399062000 | 5.046267000 | 10.933051000 |
| 6 | 3.791416000 | 5.105649000 | 9.961319000 |
| 6 | 5.522561000 | 6.038421000 | 11.363140000 |
| 6 | 4.206488000 | 6.054607000 | 10.896999000 |
| 8 | 3.284586000 | 6.973346000 | 11.339545000 |
| 1 | 5.978976000 | 0.409099000 | 9.063108000 |
| 1 | 6.538775000 | 2.848659000 | 7.593482000 |
| 1 | 6.920092000 | 2.201817000 | 10.536616000 |
| 1 | 7.975073000 | 3.347700000 | 9.745003000 |
| 1 | 4.338148000 | 3.357557000 | 8.840814000 |
| 1 | 7.425452000 | 5.055924000 | 11.287764000 |
| 1 | 2.774198000 | 5.146259000 | 9.583525000 |
| 1 | 5.869288000 | 6.783489000 | 12.074714000 |
| 1 | 3.733396000 | 7.582944000 | 11.946004000 |
| 7 | 9.063498000 | 2.099347000 | 7.561798000 |
| 6 | 10.458540000 | 1.687743000 | 7.702966000 |
| 6 | 11.176596000 | 2.871123000 | 8.343576000 |
| 8 | 11.360815000 | 3.896857000 | 7.691044000 |
| 1 | 8.918002000 | 3.081244000 | 7.358444000 |
| 1 | 10.476189000 | 0.778298000 | 8.305950000 |
| 7 | 11.462600000 | 2.764361000 | 9.666386000 |
| 6 | 12.062316000 | 3.908070000 | 10.325745000 |
| 6 | 13.575412000 | 3.931868000 | 10.051358000 |
| 8 | 14.295699000 | 2.927398000 | 10.081263000 |
| 6 | 11.721288000 | 3.996002000 | 11.811797000 |
| 6 | 10.311108000 | 4.567768000 | 12.023307000 |
| 8 | 9.665771000 | 5.042460000 | 11.085440000 |
| 7 | 9.846463000 | 4.550623000 | 13.292435000 |
| 1 | 11.450785000 | 1.856108000 | 10.123380000 |
| 1 | 11.620427000 | 4.784392000 | 9.853131000 |
| 1 | 12.421196000 | 4.690535000 | 12.288290000 |
| 1 | 11.836941000 | 3.038546000 | 12.324669000 |
| 1 | 10.381201000 | 4.171385000 | 14.064017000 |
| 1 | 9.008215000 | 5.102657000 | 13.507611000 |
| 7 | 14.032896000 | 5.164391000 | 9.727377000 |
| 6 | 15.379621000 | 5.378948000 | 9.248252000 |
| 6 | 16.368902000 | 5.492261000 | 10.404735000 |
| 8 | 16.475262000 | 6.538330000 | 11.059621000 |
| 1 | 13.428265000 | 5.961293000 | 9.935588000 |
| 1 | 15.646200000 | 4.558193000 | 8.576548000 |
| 7 | 17.066438000 | 4.361280000 | 10.650490000 |
| 6 | 17.896996000 | 4.173473000 | 11.833517000 |
| 6 | 17.293214000 | 3.133286000 | 12.787444000 |
| 6 | 15.845116000 | 3.479996000 | 13.127647000 |
| 16 | 15.049113000 | 2.394555000 | 14.360815000 |
| 6 | 15.902101000 | 2.932742000 | 15.886002000 |
| 1 | 16.716905000 | 3.529943000 | 10.184499000 |

| | | | |
|---|---|---|---|
| 1 | 17.970317000 | 5.150822000 | 12.316556000 |
| 1 | 17.330164000 | 2.133849000 | 12.336641000 |
| 1 | 17.911880000 | 3.096169000 | 13.691548000 |
| 1 | 15.782606000 | 4.516453000 | 13.479535000 |
| 1 | 15.219291000 | 3.388836000 | 12.235914000 |
| 1 | 15.429078000 | 2.424022000 | 16.729447000 |
| 1 | 15.782145000 | 4.012747000 | 16.019918000 |
| 1 | 16.964059000 | 2.676981000 | 15.854570000 |
| 6 | 5.081500000 | -3.805390000 | 20.227124000 |
| 6 | 4.695980000 | -3.441875000 | 18.799774000 |
| 8 | 5.529117000 | -3.289750000 | 17.902893000 |
| 6 | 5.063001000 | -2.566499000 | 21.163726000 |
| 6 | 5.989145000 | -1.481715000 | 20.710156000 |
| 7 | 7.129556000 | -1.083150000 | 21.392551000 |
| 6 | 5.991311000 | -0.739594000 | 19.555858000 |
| 6 | 7.771397000 | -0.137479000 | 20.655449000 |
| 7 | 7.106552000 | 0.075917000 | 19.529542000 |
| 1 | 4.383892000 | -4.551701000 | 20.624951000 |
| 1 | 4.043631000 | -2.166346000 | 21.221686000 |
| 1 | 5.328155000 | -2.890700000 | 22.175835000 |
| 1 | 5.284778000 | -0.750820000 | 18.745589000 |
| 1 | 8.694806000 | 0.341085000 | 20.944471000 |
| 1 | 7.424888000 | -1.421834000 | 22.297966000 |
| 7 | 3.361108000 | -3.268249000 | 18.585409000 |
| 6 | 2.839616000 | -2.737662000 | 17.351260000 |
| 6 | 2.412724000 | -1.266606000 | 17.470636000 |
| 8 | 2.656922000 | -0.601107000 | 18.484185000 |
| 1 | 2.742847000 | -3.284208000 | 19.384559000 |
| 1 | 3.633547000 | -2.817649000 | 16.603830000 |
| 7 | 1.746781000 | -0.806107000 | 16.385984000 |
| 6 | 1.479009000 | 0.614000000 | 16.198346000 |
| 6 | 2.609543000 | 1.335536000 | 15.429020000 |
| 6 | 3.968711000 | 0.957551000 | 15.908606000 |
| 7 | 4.536975000 | -0.259234000 | 15.575491000 |
| 6 | 4.884689000 | 1.568161000 | 16.726902000 |
| 6 | 5.748024000 | -0.346181000 | 16.165063000 |
| 7 | 5.982631000 | 0.747504000 | 16.879946000 |
| 1 | 1.774305000 | -1.373345000 | 15.542718000 |
| 1 | 1.357785000 | 1.040312000 | 17.192247000 |
| 1 | 2.533706000 | 1.101609000 | 14.364433000 |
| 1 | 2.476192000 | 2.417982000 | 15.514988000 |
| 1 | 4.874392000 | 2.564794000 | 17.131792000 |
| 1 | 6.398406000 | -1.203215000 | 16.098992000 |
| 1 | 4.124318000 | -0.914399000 | 14.902111000 |
| 6 | 6.636997000 | -5.319611000 | 14.331724000 |
| 6 | 5.292097000 | -5.807881000 | 14.888297000 |
| 6 | 7.229107000 | -4.139183000 | 15.108336000 |
| 6 | 5.379745000 | -6.384257000 | 16.305103000 |
| 1 | 7.352745000 | -6.152706000 | 14.336723000 |
| 1 | 4.885056000 | -6.568442000 | 14.210482000 |
| 1 | 4.572098000 | -4.975376000 | 14.884393000 |
| 1 | 7.499216000 | -4.414066000 | 16.129219000 |
| 1 | 6.498316000 | -3.325485000 | 15.207186000 |
| 1 | 8.132651000 | -3.749891000 | 14.621900000 |
| 1 | 4.414493000 | -6.792568000 | 16.624479000 |
| 1 | 5.669564000 | -5.616804000 | 17.028093000 |
| 1 | 6.112784000 | -7.199839000 | 16.349780000 |
| 7 | 8.027486000 | -4.889412000 | 18.761764000 |
| 6 | 9.468347000 | -4.650381000 | 18.803826000 |
| 6 | 10.121055000 | -4.064652000 | 17.532105000 |
| 6 | 9.681847000 | -2.656972000 | 17.277965000 |
| 7 | 10.080122000 | -1.898301000 | 16.182574000 |
| 6 | 8.874279000 | -1.819026000 | 18.000885000 |
| 6 | 9.531926000 | -0.664120000 | 16.279845000 |
| 7 | 8.790013000 | -0.586140000 | 17.375510000 |
| 1 | 7.503440000 | -4.104186000 | 18.377430000 |
| 1 | 9.958904000 | -5.599665000 | 19.039739000 |
| 1 | 9.882495000 | -4.701631000 | 16.669101000 |
| 1 | 11.214577000 | -4.094554000 | 17.647460000 |
| 1 | 8.344508000 | -2.023374000 | 18.913498000 |

| | | | |
|---|---|---|---|
| 1 | 9.682882000 | 0.122959000 | 15.564767000 |
| 1 | 10.705505000 | -2.191182000 | 15.443988000 |
| 7 | 9.042212000 | -1.970882000 | 10.562427000 |
| 6 | 8.386565000 | -1.236854000 | 11.641988000 |
| 6 | 6.886859000 | -1.504134000 | 11.503040000 |
| 8 | 6.242273000 | -1.225370000 | 10.488796000 |
| 6 | 8.712545000 | 0.286451000 | 11.673445000 |
| 6 | 10.104541000 | 0.518102000 | 12.164392000 |
| 7 | 11.224144000 | 0.132360000 | 11.438702000 |
| 6 | 10.500676000 | 1.001077000 | 13.392586000 |
| 6 | 12.265598000 | 0.362636000 | 12.230266000 |
| 7 | 11.879375000 | 0.890082000 | 13.420272000 |
| 1 | 8.705748000 | -1.570551000 | 9.684056000 |
| 1 | 8.721052000 | -1.664544000 | 12.599364000 |
| 1 | 8.577985000 | 0.661809000 | 10.659458000 |
| 1 | 8.019603000 | 0.824525000 | 12.321892000 |
| 1 | 9.955412000 | 1.504839000 | 14.176497000 |
| 1 | 13.302093000 | 0.197658000 | 11.972010000 |
| 1 | 12.450990000 | 1.062367000 | 14.253457000 |
| 7 | 6.274045000 | -2.086402000 | 12.576027000 |
| 6 | 4.942101000 | -2.673579000 | 12.453572000 |
| 6 | 3.794760000 | -1.662304000 | 12.599544000 |
| 8 | 3.123336000 | -1.608115000 | 13.655174000 |
| 1 | 6.851097000 | -2.402373000 | 13.340770000 |
| 1 | 4.864207000 | -3.170969000 | 11.480310000 |
| 7 | 3.594467000 | -0.894579000 | 11.527163000 |
| 6 | 2.644581000 | 0.210835000 | 11.438806000 |
| 6 | 3.337241000 | 1.551544000 | 11.131648000 |
| 6 | 4.189860000 | 2.101659000 | 12.250588000 |
| 6 | 5.369843000 | 1.460109000 | 12.649277000 |
| 6 | 3.844242000 | 3.288487000 | 12.907439000 |
| 6 | 6.166589000 | 1.962230000 | 13.674210000 |
| 6 | 4.632137000 | 3.806318000 | 13.930789000 |
| 6 | 5.794957000 | 3.138622000 | 14.324668000 |
| 8 | 6.514658000 | 3.662298000 | 15.355364000 |
| 1 | 4.301449000 | -0.980238000 | 10.794328000 |
| 1 | 2.093422000 | 0.238234000 | 12.380770000 |
| 1 | 2.553756000 | 2.276480000 | 10.887003000 |
| 1 | 3.942385000 | 1.436093000 | 10.221726000 |
| 1 | 5.681721000 | 0.567129000 | 12.128577000 |
| 1 | 2.957399000 | 3.832828000 | 12.597099000 |
| 1 | 7.061970000 | 1.441954000 | 13.997523000 |
| 1 | 4.358361000 | 4.726573000 | 14.430848000 |
| 1 | 7.264310000 | 3.068440000 | 15.612091000 |
| 6 | 4.636691000 | 7.608641000 | 17.486299000 |
| 6 | 4.183277000 | 6.725672000 | 16.317730000 |
| 6 | 2.707339000 | 6.321913000 | 16.449486000 |
| 6 | 4.410019000 | 7.442930000 | 14.977980000 |
| 1 | 4.039616000 | 8.528930000 | 17.531523000 |
| 1 | 4.524999000 | 7.092363000 | 18.447973000 |
| 1 | 4.804740000 | 5.818383000 | 16.331394000 |
| 1 | 2.421464000 | 5.572733000 | 15.699959000 |
| 1 | 2.054228000 | 7.192471000 | 16.312147000 |
| 1 | 2.469069000 | 5.919260000 | 17.444974000 |
| 1 | 4.023660000 | 6.858173000 | 14.133209000 |
| 1 | 3.889243000 | 8.409586000 | 14.976327000 |
| 1 | 5.474249000 | 7.624287000 | 14.808565000 |
| 6 | 1.561056000 | 3.147239000 | 18.826436000 |
| 7 | 2.715011000 | 2.368794000 | 19.249303000 |
| 6 | 3.933315000 | 2.918381000 | 19.414593000 |
| 7 | 4.165123000 | 4.199754000 | 19.114263000 |
| 7 | 4.922807000 | 2.189103000 | 19.943777000 |
| 1 | 1.465313000 | 4.041987000 | 19.448094000 |
| 1 | 1.615526000 | 3.451361000 | 17.771994000 |
| 1 | 2.675269000 | 1.355782000 | 19.127264000 |
| 1 | 3.533610000 | 4.682454000 | 18.490745000 |
| 1 | 5.149970000 | 4.531892000 | 19.099157000 |
| 1 | 4.753852000 | 1.205786000 | 20.105133000 |
| 1 | 5.898239000 | 2.449220000 | 19.719861000 |
| 26 | 7.772767000 | 1.097769000 | 17.943329000 |

| | | | |
|---|---|---|---|
| 6 | 8.969806000 | 4.369860000 | 17.820574000 |
| 6 | 8.806594000 | 5.360830000 | 16.684562000 |
| 6 | 8.566028000 | 6.814724000 | 17.075377000 |
| 6 | 7.650825000 | 4.005293000 | 18.511269000 |
| 8 | 9.185093000 | 7.313217000 | 18.050656000 |
| 8 | 7.828371000 | 7.474495000 | 16.272893000 |
| 8 | 6.803434000 | 4.890603000 | 18.650290000 |
| 8 | 7.494669000 | 2.788535000 | 18.962221000 |
| 16 | 9.792406000 | 2.823597000 | 17.203302000 |
| 1 | 9.661494000 | 4.750139000 | 18.577917000 |
| 1 | 9.736921000 | 5.382563000 | 16.104898000 |
| 1 | 8.014774000 | 5.038656000 | 16.018556000 |
| 8 | 8.544863000 | 2.133389000 | 16.455928000 |
| 8 | 11.886623000 | -0.129700000 | 18.553021000 |
| 1 | 11.876692000 | 0.209019000 | 17.626170000 |
| 1 | 12.386209000 | -0.958423000 | 18.494351000 |
| 8 | 12.433353000 | 1.163326000 | 16.144807000 |
| 1 | 11.822969000 | 1.938924000 | 16.075241000 |
| 1 | 13.127375000 | 1.507129000 | 16.773285000 |
| 8 | 12.024913000 | 3.988945000 | 18.989909000 |
| 1 | 12.006412000 | 4.930051000 | 18.633042000 |
| 1 | 12.108556000 | 4.098712000 | 19.949480000 |
| 8 | 11.747676000 | 6.475243000 | 18.143582000 |
| 1 | 10.802181000 | 6.724817000 | 18.304423000 |
| 1 | 11.819054000 | 6.535176000 | 17.162692000 |
| 1 | 9.686557000 | -3.976808000 | 19.642517000 |
| 1 | 7.813387000 | -5.689476000 | 18.170812000 |
| 1 | 1.992806000 | -3.339438000 | 17.002369000 |
| 1 | 0.530520000 | 0.724876000 | 15.664836000 |
| 1 | 0.665878000 | 2.539631000 | 18.967502000 |
| 1 | 5.690468000 | 7.877040000 | 17.374080000 |
| 1 | 18.905382000 | 3.870824000 | 11.528986000 |
| 1 | 15.408582000 | 6.313401000 | 8.681841000 |
| 1 | 10.910417000 | 1.486415000 | 6.726860000 |
| 1 | 5.401527000 | 0.839676000 | 7.613786000 |
| 1 | 10.037108000 | -1.742354000 | 10.601106000 |
| 1 | 6.079938000 | -4.250476000 | 20.194494000 |
| 1 | 9.010404000 | 8.929769000 | 10.041552000 |
| 1 | 13.941282000 | 8.377247000 | 8.900012000 |
| 1 | 15.583157000 | 9.683462000 | 13.633110000 |
| 1 | 1.929369000 | -0.017068000 | 10.639470000 |
| 1 | 4.822527000 | -3.418143000 | 13.239783000 |
| 1 | 6.515916000 | -5.040764000 | 13.277012000 |
| 8 | 7.529031000 | 6.243550000 | 13.867046000 |
| 1 | 7.603439000 | 6.675055000 | 14.758954000 |
| 1 | 6.824494000 | 5.584909000 | 13.968441000 |
| 8 | 13.779131000 | 1.995742000 | 18.308344000 |
| 1 | 13.241359000 | 2.803705000 | 18.507991000 |
| 1 | 13.278691000 | 1.275326000 | 18.745247000 |
| 8 | 10.742381000 | 3.294869000 | 16.089307000 |

**$^1Re_S$:**

| | | | |
|---|---|---|---|
| 6 | 14.927160000 | 8.736387000 | 13.481319000 |
| 6 | 13.776713000 | 9.177400000 | 12.588974000 |
| 8 | 12.714282000 | 9.638163000 | 13.028026000 |
| 6 | 14.488074000 | 8.294293000 | 14.877408000 |
| 6 | 13.569381000 | 7.065214000 | 14.857034000 |
| 6 | 13.458474000 | 6.418507000 | 16.229703000 |
| 8 | 13.470177000 | 7.110586000 | 17.268029000 |
| 7 | 13.350006000 | 5.084896000 | 16.240401000 |
| 1 | 15.488771000 | 7.942160000 | 12.975659000 |
| 1 | 15.382481000 | 8.070834000 | 15.468750000 |
| 1 | 13.983064000 | 9.119381000 | 15.387441000 |
| 1 | 13.926743000 | 6.328126000 | 14.129309000 |
| 1 | 12.566847000 | 7.354465000 | 14.520280000 |
| 1 | 13.348303000 | 4.528715000 | 17.096725000 |
| 1 | 13.387319000 | 4.563208000 | 15.368214000 |
| 7 | 14.015581000 | 9.069855000 | 11.257651000 |
| 6 | 12.939184000 | 9.357452000 | 10.334704000 |
| 6 | 11.748041000 | 8.427570000 | 10.611076000 |

| | | | |
|---|---|---|---|
| 8 | 11.941933000 | 7.242737000 | 10.900609000 |
| 1 | 14.791853000 | 8.477094000 | 10.951758000 |
| 1 | 12.642875000 | 10.408244000 | 10.416941000 |
| 7 | 10.517481000 | 8.968800000 | 10.499536000 |
| 6 | 9.361551000 | 8.301628000 | 11.107098000 |
| 6 | 9.378955000 | 8.581192000 | 12.612799000 |
| 6 | 8.141462000 | 8.137738000 | 13.385968000 |
| 6 | 8.147526000 | 8.697514000 | 14.814577000 |
| 7 | 9.166792000 | 8.120152000 | 15.687064000 |
| 6 | 10.311179000 | 8.686588000 | 16.069462000 |
| 7 | 10.864620000 | 9.713615000 | 15.365611000 |
| 7 | 10.893894000 | 8.287698000 | 17.203130000 |
| 1 | 10.477514000 | 9.979720000 | 10.460307000 |
| 1 | 9.429234000 | 7.232308000 | 10.897022000 |
| 1 | 10.271757000 | 8.096425000 | 13.023380000 |
| 1 | 9.506306000 | 9.666436000 | 12.747338000 |
| 1 | 7.239282000 | 8.520672000 | 12.887946000 |
| 1 | 8.062331000 | 7.044654000 | 13.412097000 |
| 1 | 8.288267000 | 9.783689000 | 14.796329000 |
| 1 | 7.182647000 | 8.497472000 | 15.293162000 |
| 1 | 8.944875000 | 7.205411000 | 16.130296000 |
| 1 | 11.710845000 | 10.100722000 | 15.762640000 |
| 1 | 10.928993000 | 9.633650000 | 14.356252000 |
| 1 | 10.438877000 | 7.530840000 | 17.745057000 |
| 1 | 11.913211000 | 8.276460000 | 17.259773000 |
| 7 | 5.928328000 | 1.602088000 | 8.282235000 |
| 6 | 6.783796000 | 2.778159000 | 8.103416000 |
| 6 | 8.150875000 | 2.440945000 | 7.503056000 |
| 8 | 8.866539000 | 1.551885000 | 7.979037000 |
| 6 | 7.059277000 | 3.478227000 | 9.449309000 |
| 6 | 5.984262000 | 4.415786000 | 9.954524000 |
| 6 | 4.652488000 | 4.370519000 | 9.522810000 |
| 6 | 6.330438000 | 5.356187000 | 10.936506000 |
| 6 | 3.688749000 | 5.230752000 | 10.051323000 |
| 6 | 5.373771000 | 6.213898000 | 11.475625000 |
| 6 | 4.050040000 | 6.154614000 | 11.032391000 |
| 8 | 3.066722000 | 6.970505000 | 11.541357000 |
| 1 | 6.452576000 | 0.870029000 | 8.768539000 |
| 1 | 6.262524000 | 3.476347000 | 7.435596000 |
| 1 | 7.236271000 | 2.691900000 | 10.194329000 |
| 1 | 7.999173000 | 4.036585000 | 9.394083000 |
| 1 | 4.357012000 | 3.628501000 | 8.787764000 |
| 1 | 7.358775000 | 5.401383000 | 11.283369000 |
| 1 | 2.657229000 | 5.194449000 | 9.715249000 |
| 1 | 5.658874000 | 6.928420000 | 12.244848000 |
| 1 | 3.481475000 | 7.601791000 | 12.149472000 |
| 7 | 8.561382000 | 3.195971000 | 6.452778000 |
| 6 | 9.941042000 | 3.133023000 | 5.974748000 |
| 6 | 10.847381000 | 4.027520000 | 6.829706000 |
| 8 | 11.251796000 | 5.114238000 | 6.428462000 |
| 1 | 8.018185000 | 4.012255000 | 6.205372000 |
| 1 | 10.257198000 | 2.088291000 | 6.015545000 |
| 7 | 11.106590000 | 3.559369000 | 8.078284000 |
| 6 | 11.741857000 | 4.443946000 | 9.034374000 |
| 6 | 13.194912000 | 4.746089000 | 8.635531000 |
| 8 | 13.977165000 | 3.919909000 | 8.164568000 |
| 6 | 11.682120000 | 3.875546000 | 10.457048000 |
| 6 | 10.477061000 | 4.408001000 | 11.238089000 |
| 8 | 9.575381000 | 5.047279000 | 10.685589000 |
| 7 | 10.496453000 | 4.157610000 | 12.562080000 |
| 1 | 10.688458000 | 2.685848000 | 8.379893000 |
| 1 | 11.195007000 | 5.390190000 | 9.028824000 |
| 1 | 12.585454000 | 4.152401000 | 11.011526000 |
| 1 | 11.657267000 | 2.779174000 | 10.444540000 |
| 1 | 11.147538000 | 3.476814000 | 12.930298000 |
| 1 | 9.663136000 | 4.385969000 | 13.124142000 |
| 7 | 13.537809000 | 6.035282000 | 8.899499000 |
| 6 | 14.868841000 | 6.533372000 | 8.627768000 |
| 6 | 15.808493000 | 6.212802000 | 9.787142000 |
| 8 | 16.060445000 | 7.035024000 | 10.680564000 |

| | | | |
|---|---|---|---|
| 1 | 12.947281000 | 6.556599000 | 9.548441000 |
| 1 | 15.221217000 | 6.074382000 | 7.700511000 |
| 7 | 16.277897000 | 4.946590000 | 9.760538000 |
| 6 | 17.095782000 | 4.374040000 | 10.816466000 |
| 6 | 16.299128000 | 3.464860000 | 11.761786000 |
| 6 | 15.267095000 | 4.231892000 | 12.587091000 |
| 16 | 14.262933000 | 3.146364000 | 13.671716000 |
| 6 | 15.484187000 | 2.723983000 | 14.961858000 |
| 1 | 15.848694000 | 4.329652000 | 9.070066000 |
| 1 | 17.541146000 | 5.208837000 | 11.363463000 |
| 1 | 15.790006000 | 2.686274000 | 11.181045000 |
| 1 | 17.008686000 | 2.956395000 | 12.425782000 |
| 1 | 15.744389000 | 5.005284000 | 13.199191000 |
| 1 | 14.547726000 | 4.737419000 | 11.935851000 |
| 1 | 14.991134000 | 2.043054000 | 15.659575000 |
| 1 | 15.794846000 | 3.618383000 | 15.509164000 |
| 1 | 16.356160000 | 2.225176000 | 14.531623000 |
| 6 | 3.447589000 | -2.059842000 | 21.891728000 |
| 6 | 3.038532000 | -1.911964000 | 20.432306000 |
| 8 | 3.730233000 | -2.318081000 | 19.495941000 |
| 6 | 4.047175000 | -0.769169000 | 22.508002000 |
| 6 | 5.295011000 | -0.299731000 | 21.826159000 |
| 7 | 6.500999000 | -0.055672000 | 22.471427000 |
| 6 | 5.547347000 | -0.029466000 | 20.507301000 |
| 6 | 7.421560000 | 0.336471000 | 21.551549000 |
| 7 | 6.864404000 | 0.351978000 | 20.353121000 |
| 1 | 2.571298000 | -2.346139000 | 22.485242000 |
| 1 | 3.295698000 | 0.031232000 | 22.490203000 |
| 1 | 4.255330000 | -0.967862000 | 23.565174000 |
| 1 | 4.895151000 | -0.118814000 | 19.658284000 |
| 1 | 8.450369000 | 0.592001000 | 21.757420000 |
| 1 | 6.664962000 | -0.130796000 | 23.465972000 |
| 7 | 1.836393000 | -1.300078000 | 20.214606000 |
| 6 | 1.317061000 | -1.098151000 | 18.886895000 |
| 6 | 1.644986000 | 0.281820000 | 18.291997000 |
| 8 | 2.157798000 | 1.174342000 | 18.975547000 |
| 1 | 1.341809000 | -0.894347000 | 20.995849000 |
| 1 | 1.752454000 | -1.872059000 | 18.248955000 |
| 7 | 1.295904000 | 0.423364000 | 16.994797000 |
| 6 | 1.749513000 | 1.516393000 | 16.134010000 |
| 6 | 2.911977000 | 1.068643000 | 15.222941000 |
| 6 | 4.020465000 | 0.469823000 | 16.021421000 |
| 7 | 4.185387000 | -0.898338000 | 16.196512000 |
| 6 | 4.988071000 | 1.044469000 | 16.801791000 |
| 6 | 5.244128000 | -1.105069000 | 17.021965000 |
| 7 | 5.737918000 | 0.056709000 | 17.412960000 |
| 1 | 0.909805000 | -0.389424000 | 16.529797000 |
| 1 | 2.080018000 | 2.322818000 | 16.789602000 |
| 1 | 2.557361000 | 0.351529000 | 14.472787000 |
| 1 | 3.282642000 | 1.932398000 | 14.661766000 |
| 1 | 5.221407000 | 2.085030000 | 16.940186000 |
| 1 | 5.602422000 | -2.077839000 | 17.312101000 |
| 1 | 3.720582000 | -1.617815000 | 15.660199000 |
| 6 | 9.264119000 | -3.665394000 | 13.619863000 |
| 6 | 7.786656000 | -3.254573000 | 13.688709000 |
| 6 | 10.221804000 | -2.643956000 | 14.239999000 |
| 6 | 7.248147000 | -3.119314000 | 15.115023000 |
| 1 | 9.390568000 | -4.644080000 | 14.105462000 |
| 1 | 7.165375000 | -3.981189000 | 13.152466000 |
| 1 | 7.649576000 | -2.299167000 | 13.166823000 |
| 1 | 10.031971000 | -2.520587000 | 15.312005000 |
| 1 | 10.102628000 | -1.659562000 | 13.775788000 |
| 1 | 11.265546000 | -2.953432000 | 14.116603000 |
| 1 | 6.164625000 | -2.960901000 | 15.090460000 |
| 1 | 7.701018000 | -2.275901000 | 15.646388000 |
| 1 | 7.437116000 | -4.028648000 | 15.701668000 |
| 7 | 5.402829000 | -4.658826000 | 20.623761000 |
| 6 | 6.817422000 | -4.885012000 | 20.897111000 |
| 6 | 7.786732000 | -4.956507000 | 19.691094000 |
| 6 | 8.107799000 | -3.612932000 | 19.113079000 |

| | | | |
|---|---|---|---|
| 7 | 9.262930000 | -3.349814000 | 18.387743000 |
| 6 | 7.466005000 | -2.403517000 | 19.190491000 |
| 6 | 9.293319000 | -2.035590000 | 18.058474000 |
| 7 | 8.211154000 | -1.438515000 | 18.530817000 |
| 1 | 5.215670000 | -3.826374000 | 20.068578000 |
| 1 | 6.895648000 | -5.828098000 | 21.448100000 |
| 1 | 7.361999000 | -5.619761000 | 18.923642000 |
| 1 | 8.723124000 | -5.428104000 | 20.020405000 |
| 1 | 6.538560000 | -2.154705000 | 19.676880000 |
| 1 | 10.067187000 | -1.547186000 | 17.490526000 |
| 1 | 9.973368000 | -4.025580000 | 18.142972000 |
| 7 | 9.896273000 | -1.104183000 | 9.141848000 |
| 6 | 9.004549000 | -1.144664000 | 10.300861000 |
| 6 | 7.691686000 | -1.783370000 | 9.871372000 |
| 8 | 6.869925000 | -1.222344000 | 9.135023000 |
| 6 | 8.745470000 | 0.224685000 | 10.983711000 |
| 6 | 9.911524000 | 0.598060000 | 11.841026000 |
| 7 | 11.219294000 | 0.543098000 | 11.375280000 |
| 6 | 9.916731000 | 0.907012000 | 13.179893000 |
| 6 | 11.985210000 | 0.804136000 | 12.421326000 |
| 7 | 11.242960000 | 1.048971000 | 13.540032000 |
| 1 | 9.509520000 | -0.429247000 | 8.481573000 |
| 1 | 9.475179000 | -1.800225000 | 11.044676000 |
| 1 | 8.570177000 | 0.954515000 | 10.189437000 |
| 1 | 7.844638000 | 0.187253000 | 11.605891000 |
| 1 | 9.123374000 | 1.045335000 | 13.889295000 |
| 1 | 13.063588000 | 0.851724000 | 12.418988000 |
| 1 | 11.577141000 | 1.184718000 | 14.500411000 |
| 7 | 7.480630000 | -3.030198000 | 10.364094000 |
| 6 | 6.291478000 | -3.811771000 | 10.069470000 |
| 6 | 5.065252000 | -3.285779000 | 10.831253000 |
| 8 | 4.734386000 | -3.733687000 | 11.931544000 |
| 1 | 8.178767000 | -3.429168000 | 10.971643000 |
| 1 | 6.110252000 | -3.793777000 | 8.989920000 |
| 7 | 4.420481000 | -2.292857000 | 10.173989000 |
| 6 | 3.268338000 | -1.591797000 | 10.710994000 |
| 6 | 3.489293000 | -0.082488000 | 10.794880000 |
| 6 | 4.500546000 | 0.408950000 | 11.811066000 |
| 6 | 5.148391000 | -0.403923000 | 12.748138000 |
| 6 | 4.793645000 | 1.780722000 | 11.811576000 |
| 6 | 6.046984000 | 0.144367000 | 13.672713000 |
| 6 | 5.674070000 | 2.339903000 | 12.727429000 |
| 6 | 6.291043000 | 1.516889000 | 13.672373000 |
| 8 | 7.123591000 | 2.124418000 | 14.570608000 |
| 1 | 4.949801000 | -1.850684000 | 9.424257000 |
| 1 | 3.054395000 | -2.029900000 | 11.689308000 |
| 1 | 2.515587000 | 0.384294000 | 11.003826000 |
| 1 | 3.787469000 | 0.297191000 | 9.807404000 |
| 1 | 4.977022000 | -1.476490000 | 12.765787000 |
| 1 | 4.331115000 | 2.425450000 | 11.073200000 |
| 1 | 6.550605000 | -0.491272000 | 14.393986000 |
| 1 | 5.902600000 | 3.399862000 | 12.702959000 |
| 1 | 7.430114000 | 1.517595000 | 15.282167000 |
| 6 | 4.601506000 | 6.950474000 | 15.932469000 |
| 6 | 4.920272000 | 5.637366000 | 15.208974000 |
| 6 | 5.495693000 | 4.605224000 | 16.179910000 |
| 6 | 3.678151000 | 5.081735000 | 14.501701000 |
| 1 | 3.813986000 | 6.798280000 | 16.686833000 |
| 1 | 5.485330000 | 7.344128000 | 16.448737000 |
| 1 | 5.692514000 | 5.834170000 | 14.454704000 |
| 1 | 5.827309000 | 3.713886000 | 15.644560000 |
| 1 | 4.728243000 | 4.305167000 | 16.905152000 |
| 1 | 6.360019000 | 4.991867000 | 16.728162000 |
| 1 | 3.921570000 | 4.174390000 | 13.938507000 |
| 1 | 2.901089000 | 4.828909000 | 15.239819000 |
| 1 | 3.243506000 | 5.795061000 | 13.794189000 |
| 6 | 1.914129000 | 5.341204000 | 19.141503000 |
| 7 | 2.643181000 | 4.083848000 | 19.160460000 |
| 6 | 3.965643000 | 3.988422000 | 19.383736000 |
| 7 | 4.731092000 | 5.076034000 | 19.506717000 |
| 7 | 4.511157000 | 2.776069000 | 19.545453000 |
| 1 | 2.144112000 | 5.938160000 | 20.030768000 |
| 1 | 2.141314000 | 5.930802000 | 18.244311000 |
| 1 | 2.151157000 | 3.202645000 | 19.046092000 |
| 1 | 4.415252000 | 5.950189000 | 19.113117000 |
| 1 | 5.753167000 | 4.913221000 | 19.562671000 |
| 1 | 3.901761000 | 1.978702000 | 19.404152000 |
| 1 | 5.500191000 | 2.651214000 | 19.297819000 |
| 26 | 7.736676000 | 0.577869000 | 18.412548000 |
| 6 | 9.377767000 | 3.121446000 | 19.015933000 |
| 6 | 9.877651000 | 3.686144000 | 17.670145000 |
| 6 | 9.555012000 | 5.181761000 | 17.550404000 |
| 6 | 7.876391000 | 3.327774000 | 19.086363000 |
| 8 | 10.056192000 | 5.960893000 | 18.403861000 |
| 8 | 8.795300000 | 5.558396000 | 16.607599000 |
| 8 | 7.386551000 | 4.320099000 | 19.626144000 |
| 8 | 7.136941000 | 2.425080000 | 18.470027000 |
| 16 | 9.742662000 | 1.320648000 | 19.264932000 |
| 1 | 9.862638000 | 3.663469000 | 19.826916000 |
| 1 | 10.963201000 | 3.570970000 | 17.636526000 |
| 1 | 9.436768000 | 3.138292000 | 16.834765000 |
| 8 | 8.257188000 | 0.735647000 | 16.579011000 |
| 8 | 14.317623000 | 0.258831000 | 17.362978000 |
| 1 | 13.446492000 | 0.283366000 | 16.900630000 |
| 1 | 14.950222000 | -0.088585000 | 16.717694000 |
| 8 | 11.942163000 | 1.132756000 | 16.336293000 |
| 1 | 11.102635000 | 0.865481000 | 16.758376000 |
| 1 | 12.278488000 | 1.890177000 | 16.888042000 |
| 8 | 12.383825000 | 3.409441000 | 20.284183000 |
| 1 | 12.502237000 | 4.394510000 | 20.102494000 |
| 1 | 12.926088000 | 3.218108000 | 21.063560000 |
| 8 | 12.465312000 | 5.940799000 | 19.640337000 |
| 1 | 11.543257000 | 5.948154000 | 19.289235000 |
| 1 | 12.998612000 | 6.298464000 | 18.898079000 |
| 1 | 7.169528000 | -4.097794000 | 21.575330000 |
| 1 | 5.006296000 | -5.446150000 | 20.114845000 |
| 1 | 0.229095000 | -1.233533000 | 18.892089000 |
| 1 | 0.906984000 | 1.881753000 | 15.538817000 |
| 1 | 0.845789000 | 5.120570000 | 19.141528000 |
| 1 | 4.237563000 | 7.721996000 | 15.242469000 |
| 1 | 17.906965000 | 3.800456000 | 10.354864000 |
| 1 | 14.835857000 | 7.616933000 | 8.494872000 |
| 1 | 9.982791000 | 3.491906000 | 4.946090000 |
| 1 | 5.713293000 | 1.202747000 | 7.368695000 |
| 1 | 10.773402000 | -0.697766000 | 9.472161000 |
| 1 | 4.166639000 | -2.882781000 | 21.935789000 |
| 1 | 8.453843000 | 8.693211000 | 10.636135000 |
| 1 | 13.298621000 | 9.188504000 | 9.314969000 |
| 1 | 15.614490000 | 9.588736000 | 13.569446000 |
| 1 | 2.398087000 | -1.785177000 | 10.070357000 |
| 1 | 6.466236000 | -4.840594000 | 10.385744000 |
| 1 | 9.552539000 | -3.819250000 | 12.569372000 |
| 8 | 8.176477000 | 4.719411000 | 14.107896000 |
| 1 | 8.426881000 | 4.957930000 | 15.032788000 |
| 1 | 7.788133000 | 3.828853000 | 14.191087000 |
| 8 | 13.348358000 | 2.786581000 | 17.908770000 |
| 1 | 13.010784000 | 2.889737000 | 18.846237000 |
| 1 | 13.996198000 | 2.049579000 | 17.913036000 |
| 8 | 9.330671000 | 0.131712000 | 16.149315000 |

**$^{1}$IM1$_{S}$:**

| | | | |
|---|---|---|---|
| 6 | 15.230043000 | 9.112242000 | 13.317600000 |
| 6 | 14.048895000 | 9.468044000 | 12.427040000 |
| 8 | 12.998675000 | 9.955470000 | 12.865967000 |
| 6 | 14.787694000 | 8.522999000 | 14.659383000 |
| 6 | 13.971242000 | 7.240517000 | 14.490588000 |
| 6 | 13.602498000 | 6.594274000 | 15.812939000 |
| 8 | 13.516169000 | 7.261283000 | 16.868241000 |
| 7 | 13.352708000 | 5.280122000 | 15.784758000 |
| 1 | 15.896865000 | 8.422842000 | 12.789025000 |

| | | | |
|---|---|---|---|
| 1 | 15.672439000 | 8.320960000 | 15.272660000 |
| 1 | 14.195508000 | 9.259008000 | 15.209300000 |
| 1 | 14.511326000 | 6.516967000 | 13.869634000 |
| 1 | 13.039647000 | 7.455842000 | 13.950253000 |
| 1 | 12.974230000 | 4.844400000 | 16.622909000 |
| 1 | 13.448164000 | 4.719423000 | 14.943255000 |
| 7 | 14.231436000 | 9.230217000 | 11.104246000 |
| 6 | 13.120399000 | 9.429972000 | 10.201177000 |
| 6 | 11.928549000 | 8.551000000 | 10.615249000 |
| 8 | 12.118046000 | 7.418182000 | 11.071531000 |
| 1 | 15.021717000 | 8.645419000 | 10.820051000 |
| 1 | 12.831817000 | 10.486325000 | 10.182652000 |
| 7 | 10.700265000 | 9.069200000 | 10.420974000 |
| 6 | 9.526172000 | 8.488954000 | 11.083678000 |
| 6 | 9.554078000 | 8.893828000 | 12.562459000 |
| 6 | 8.263629000 | 8.678879000 | 13.349220000 |
| 6 | 8.329577000 | 9.357684000 | 14.727138000 |
| 7 | 9.230482000 | 8.713103000 | 15.679345000 |
| 6 | 10.479017000 | 9.059444000 | 15.984892000 |
| 7 | 11.185060000 | 9.933364000 | 15.223255000 |
| 7 | 11.039611000 | 8.593367000 | 17.109689000 |
| 1 | 10.656130000 | 10.060352000 | 10.220253000 |
| 1 | 9.553274000 | 7.403834000 | 10.961199000 |
| 1 | 10.378563000 | 8.345397000 | 13.031173000 |
| 1 | 9.807222000 | 9.964553000 | 12.599542000 |
| 1 | 7.421121000 | 9.128600000 | 12.804429000 |
| 1 | 8.044799000 | 7.611516000 | 13.465524000 |
| 1 | 8.628463000 | 10.406988000 | 14.622431000 |
| 1 | 7.338169000 | 9.344929000 | 15.192510000 |
| 1 | 8.842945000 | 7.916945000 | 16.218376000 |
| 1 | 12.121715000 | 10.150833000 | 15.534681000 |
| 1 | 11.117404000 | 9.897949000 | 14.214274000 |
| 1 | 10.504868000 | 7.936030000 | 17.695878000 |
| 1 | 12.044386000 | 8.407800000 | 17.092183000 |
| 7 | 5.732344000 | 1.163270000 | 8.157611000 |
| 6 | 6.791919000 | 2.171083000 | 8.089326000 |
| 6 | 8.128847000 | 1.548958000 | 7.644639000 |
| 8 | 8.338844000 | 0.340799000 | 7.753892000 |
| 6 | 7.007322000 | 2.830254000 | 9.475801000 |
| 6 | 6.042816000 | 3.933003000 | 9.858702000 |
| 6 | 4.713027000 | 3.970144000 | 9.420280000 |
| 6 | 6.471115000 | 4.912052000 | 10.765382000 |
| 6 | 3.824798000 | 4.937917000 | 9.891684000 |
| 6 | 5.588960000 | 5.873136000 | 11.254876000 |
| 6 | 4.257794000 | 5.881120000 | 10.828053000 |
| 8 | 3.337573000 | 6.784830000 | 11.300678000 |
| 1 | 6.141931000 | 0.331217000 | 8.593783000 |
| 1 | 6.498592000 | 2.948114000 | 7.371904000 |
| 1 | 6.934692000 | 2.030436000 | 10.223755000 |
| 1 | 8.028018000 | 3.222221000 | 9.552955000 |
| 1 | 4.360417000 | 3.208480000 | 8.732560000 |
| 1 | 7.502089000 | 4.919492000 | 11.108531000 |
| 1 | 2.792530000 | 4.965401000 | 9.556302000 |
| 1 | 5.938968000 | 6.604283000 | 11.981841000 |
| 1 | 3.777436000 | 7.354714000 | 11.950313000 |
| 7 | 9.079965000 | 2.413542000 | 7.215263000 |
| 6 | 10.499570000 | 2.067605000 | 7.236646000 |
| 6 | 11.201469000 | 3.267893000 | 7.858773000 |
| 8 | 11.319835000 | 4.311974000 | 7.219360000 |
| 1 | 8.865074000 | 3.401308000 | 7.156040000 |
| 1 | 10.611243000 | 1.144150000 | 7.809053000 |
| 7 | 11.547826000 | 3.169575000 | 9.166176000 |
| 6 | 12.078935000 | 4.357406000 | 9.799599000 |
| 6 | 13.506202000 | 4.644245000 | 9.301302000 |
| 8 | 14.296919000 | 3.774093000 | 8.931624000 |
| 6 | 11.995140000 | 4.309708000 | 11.324446000 |
| 6 | 10.606482000 | 4.722309000 | 11.829974000 |
| 8 | 9.724084000 | 5.097214000 | 11.052822000 |
| 7 | 10.454278000 | 4.700373000 | 13.169118000 |
| 1 | 11.527067000 | 2.276548000 | 9.648331000 |

| | | | |
|---|---|---|---|
| 1 | 11.450239000 | 5.187537000 | 9.472422000 |
| 1 | 12.704561000 | 5.026350000 | 11.749132000 |
| 1 | 12.267186000 | 3.325245000 | 11.712545000 |
| 1 | 11.145751000 | 4.282465000 | 13.782210000 |
| 1 | 9.554925000 | 4.999217000 | 13.567158000 |
| 7 | 13.800449000 | 5.970639000 | 9.293695000 |
| 6 | 15.065236000 | 6.462245000 | 8.786048000 |
| 6 | 16.148702000 | 6.339283000 | 9.854237000 |
| 8 | 16.381897000 | 7.256547000 | 10.654352000 |
| 1 | 13.208846000 | 6.589235000 | 9.849405000 |
| 1 | 15.323652000 | 5.888231000 | 7.892220000 |
| 7 | 16.753071000 | 5.130757000 | 9.865208000 |
| 6 | 17.620215000 | 4.679789000 | 10.943316000 |
| 6 | 16.910905000 | 3.659386000 | 11.845638000 |
| 6 | 15.676423000 | 4.265094000 | 12.509062000 |
| 16 | 14.698875000 | 3.103564000 | 13.526925000 |
| 6 | 15.828503000 | 2.848171000 | 14.938103000 |
| 1 | 16.288425000 | 4.418458000 | 9.302034000 |
| 1 | 17.915969000 | 5.567671000 | 11.507679000 |
| 1 | 16.610677000 | 2.786298000 | 11.253654000 |
| 1 | 17.620879000 | 3.306992000 | 12.603055000 |
| 1 | 15.957454000 | 5.124795000 | 13.127785000 |
| 1 | 14.980069000 | 4.625005000 | 11.748226000 |
| 1 | 15.269210000 | 2.271430000 | 15.679603000 |
| 1 | 16.108003000 | 3.812833000 | 15.373453000 |
| 1 | 16.724344000 | 2.300762000 | 14.635279000 |
| 6 | 3.565235000 | -2.707028000 | 21.277962000 |
| 6 | 3.007489000 | -2.551015000 | 19.869384000 |
| 8 | 3.635500000 | -2.854964000 | 18.856936000 |
| 6 | 3.954988000 | -1.361059000 | 21.931380000 |
| 6 | 5.073443000 | -0.653130000 | 21.230873000 |
| 7 | 5.980701000 | 0.162764000 | 21.891657000 |
| 6 | 5.459351000 | -0.611740000 | 19.914525000 |
| 6 | 6.867092000 | 0.652274000 | 20.985134000 |
| 7 | 6.577750000 | 0.190299000 | 19.780984000 |
| 1 | 2.809028000 | -3.185995000 | 21.910962000 |
| 1 | 3.076432000 | -0.701911000 | 21.983519000 |
| 1 | 4.247176000 | -1.561109000 | 22.969367000 |
| 1 | 5.034313000 | -1.114030000 | 19.064222000 |
| 1 | 7.680703000 | 1.316989000 | 21.232103000 |
| 1 | 5.976070000 | 0.373736000 | 22.879742000 |
| 7 | 1.739419000 | -2.040589000 | 19.801579000 |
| 6 | 1.162775000 | -1.675418000 | 18.534534000 |
| 6 | 1.551786000 | -0.257756000 | 18.081116000 |
| 8 | 1.973850000 | 0.581800000 | 18.888250000 |
| 1 | 1.325296000 | -1.656536000 | 20.638815000 |
| 1 | 1.498106000 | -2.406072000 | 17.794145000 |
| 7 | 1.332374000 | -0.023638000 | 16.773996000 |
| 6 | 1.729462000 | 1.181471000 | 16.051742000 |
| 6 | 2.826357000 | 0.864610000 | 15.022361000 |
| 6 | 4.049318000 | 0.306360000 | 15.673428000 |
| 7 | 4.530728000 | -0.973306000 | 15.462073000 |
| 6 | 4.901157000 | 0.875753000 | 16.583989000 |
| 6 | 5.648521000 | -1.130018000 | 16.211031000 |
| 7 | 5.894620000 | -0.027460000 | 16.912949000 |
| 1 | 1.050959000 | -0.814392000 | 16.207222000 |
| 1 | 2.091213000 | 1.896961000 | 16.791300000 |
| 1 | 2.440650000 | 0.155462000 | 14.281571000 |
| 1 | 3.079062000 | 1.775376000 | 14.470275000 |
| 1 | 4.914178000 | 1.879275000 | 16.967497000 |
| 1 | 6.238941000 | -2.028177000 | 16.224965000 |
| 1 | 4.250304000 | -1.600065000 | 14.698120000 |
| 6 | 8.640772000 | -3.757280000 | 14.240941000 |
| 6 | 7.273807000 | -4.410514000 | 14.534479000 |
| 6 | 8.622344000 | -2.229712000 | 14.345587000 |
| 6 | 7.034377000 | -4.668991000 | 16.025575000 |
| 1 | 9.391982000 | -4.170655000 | 14.928626000 |
| 1 | 7.197015000 | -5.367619000 | 14.002332000 |
| 1 | 6.467318000 | -3.775390000 | 14.143825000 |
| 1 | 8.279107000 | -1.892351000 | 15.324344000 |

| | | | |
|---|---|---|---|
| 1 | 7.945573000 | -1.797281000 | 13.602561000 |
| 1 | 9.612758000 | -1.793700000 | 14.181751000 |
| 1 | 6.016846000 | -5.029873000 | 16.211971000 |
| 1 | 7.183211000 | -3.768844000 | 16.629631000 |
| 1 | 7.733632000 | -5.424963000 | 16.403190000 |
| 7 | 6.073898000 | -4.890035000 | 20.007389000 |
| 6 | 7.436948000 | -4.937855000 | 20.531938000 |
| 6 | 8.605091000 | -4.856250000 | 19.522309000 |
| 6 | 8.699999000 | -3.510662000 | 18.877143000 |
| 7 | 9.666565000 | -3.142974000 | 17.951733000 |
| 6 | 7.907500000 | -2.401535000 | 19.001812000 |
| 6 | 9.437057000 | -1.864648000 | 17.553327000 |
| 7 | 8.364626000 | -1.388558000 | 18.172511000 |
| 1 | 5.908552000 | -4.093729000 | 19.396328000 |
| 1 | 7.538099000 | -5.870585000 | 21.095848000 |
| 1 | 8.480180000 | -5.634383000 | 18.755590000 |
| 1 | 9.543118000 | -5.085369000 | 20.048238000 |
| 1 | 7.049639000 | -2.262815000 | 19.632990000 |
| 1 | 10.059834000 | -1.348896000 | 16.839425000 |
| 1 | 10.427274000 | -3.720692000 | 17.622509000 |
| 7 | 9.473349000 | -1.928002000 | 9.445383000 |
| 6 | 8.729315000 | -1.550993000 | 10.644525000 |
| 6 | 7.282511000 | -2.020368000 | 10.489354000 |
| 8 | 6.502722000 | -1.551959000 | 9.654892000 |
| 6 | 8.732337000 | -0.032289000 | 10.972935000 |
| 6 | 10.005576000 | 0.435129000 | 11.594187000 |
| 7 | 11.241985000 | 0.341711000 | 10.961212000 |
| 6 | 10.174202000 | 0.973577000 | 12.848221000 |
| 6 | 12.127672000 | 0.805266000 | 11.829507000 |
| 7 | 11.533944000 | 1.187337000 | 12.998896000 |
| 1 | 9.086534000 | -1.396496000 | 8.663540000 |
| 1 | 9.175895000 | -2.085385000 | 11.492388000 |
| 1 | 8.539555000 | 0.485118000 | 10.033611000 |
| 1 | 7.914900000 | 0.218041000 | 11.651907000 |
| 1 | 9.466821000 | 1.243638000 | 13.617065000 |
| 1 | 13.193025000 | 0.894010000 | 11.666452000 |
| 1 | 11.909129000 | 1.789119000 | 13.744279000 |
| 7 | 6.876356000 | -2.973667000 | 11.375998000 |
| 6 | 5.541027000 | -3.555124000 | 11.342635000 |
| 6 | 4.497184000 | -2.591200000 | 11.932379000 |
| 8 | 4.174405000 | -2.640997000 | 13.131111000 |
| 1 | 7.562069000 | -3.387594000 | 11.987233000 |
| 1 | 5.283561000 | -3.810492000 | 10.309574000 |
| 7 | 4.027101000 | -1.710499000 | 11.031952000 |
| 6 | 3.121902000 | -0.606363000 | 11.322956000 |
| 6 | 3.747459000 | 0.773160000 | 11.062956000 |
| 6 | 4.715390000 | 1.265531000 | 12.116819000 |
| 6 | 5.663975000 | 0.442084000 | 12.734711000 |
| 6 | 4.696166000 | 2.616387000 | 12.490819000 |
| 6 | 6.533593000 | 0.927744000 | 13.708866000 |
| 6 | 5.565864000 | 3.120608000 | 13.450033000 |
| 6 | 6.474362000 | 2.269493000 | 14.084360000 |
| 8 | 7.278820000 | 2.787522000 | 15.055388000 |
| 1 | 4.563921000 | -1.658612000 | 10.166044000 |
| 1 | 2.808556000 | -0.717434000 | 12.362998000 |
| 1 | 2.926949000 | 1.496109000 | 10.983378000 |
| 1 | 4.230552000 | 0.775998000 | 10.074640000 |
| 1 | 5.741040000 | -0.600377000 | 12.464843000 |
| 1 | 4.005868000 | 3.296731000 | 12.003707000 |
| 1 | 7.242961000 | 0.266870000 | 14.192906000 |
| 1 | 5.559101000 | 4.174915000 | 13.692331000 |
| 1 | 7.746973000 | 2.052066000 | 15.556873000 |
| 6 | 5.122667000 | 6.856001000 | 17.027472000 |
| 6 | 4.812942000 | 5.967262000 | 15.817188000 |
| 6 | 4.231076000 | 4.620162000 | 16.262768000 |
| 6 | 3.880684000 | 6.671994000 | 14.825002000 |
| 1 | 4.204441000 | 7.079949000 | 17.593334000 |
| 1 | 5.851009000 | 6.380054000 | 17.690692000 |
| 1 | 5.763510000 | 5.772085000 | 15.308349000 |
| 1 | 3.998188000 | 3.982748000 | 15.404350000 |

| | | | |
|---|---|---|---|
| 1 | 3.302538000 | 4.765331000 | 16.834210000 |
| 1 | 4.948852000 | 4.084258000 | 16.891250000 |
| 1 | 3.689088000 | 6.053679000 | 13.940567000 |
| 1 | 2.910460000 | 6.902616000 | 15.285744000 |
| 1 | 4.317433000 | 7.622115000 | 14.489580000 |
| 6 | 1.474662000 | 4.628526000 | 19.496824000 |
| 7 | 2.315132000 | 3.450149000 | 19.372882000 |
| 6 | 3.654041000 | 3.472474000 | 19.478335000 |
| 7 | 4.322608000 | 4.617302000 | 19.646849000 |
| 7 | 4.327330000 | 2.312814000 | 19.479354000 |
| 1 | 1.701168000 | 5.173729000 | 20.419601000 |
| 1 | 1.592494000 | 5.306646000 | 18.641923000 |
| 1 | 1.901253000 | 2.539602000 | 19.185630000 |
| 1 | 3.884895000 | 5.484194000 | 19.370250000 |
| 1 | 5.359980000 | 4.584375000 | 19.566266000 |
| 1 | 3.795631000 | 1.468960000 | 19.292721000 |
| 1 | 5.300175000 | 2.329788000 | 19.138808000 |
| 26 | 7.548658000 | 0.496697000 | 18.021409000 |
| 6 | 9.070022000 | 3.439202000 | 18.516821000 |
| 6 | 9.432586000 | 4.275749000 | 17.281851000 |
| 6 | 9.232759000 | 5.777429000 | 17.530565000 |
| 6 | 7.548232000 | 3.440784000 | 18.725157000 |
| 8 | 9.973053000 | 6.320753000 | 18.395553000 |
| 8 | 8.386758000 | 6.382372000 | 16.812565000 |
| 8 | 7.068943000 | 4.422384000 | 19.318360000 |
| 8 | 6.861181000 | 2.463900000 | 18.252597000 |
| 16 | 9.798877000 | 1.748993000 | 18.412217000 |
| 1 | 9.551631000 | 3.883965000 | 19.394059000 |
| 1 | 10.491707000 | 4.137268000 | 17.049859000 |
| 1 | 8.842254000 | 3.953124000 | 16.426588000 |
| 8 | 8.494016000 | 0.836371000 | 16.343807000 |
| 8 | 11.691628000 | -0.375308000 | 15.814151000 |
| 1 | 11.005533000 | 0.290723000 | 16.025089000 |
| 1 | 11.791942000 | -0.311304000 | 14.847978000 |
| 8 | 11.726612000 | 2.885772000 | 15.188516000 |
| 1 | 10.867526000 | 2.641204000 | 15.575546000 |
| 1 | 12.404810000 | 2.507687000 | 15.825815000 |
| 8 | 13.033179000 | 2.728588000 | 19.254068000 |
| 1 | 12.838304000 | 3.679441000 | 19.047307000 |
| 1 | 13.848496000 | 2.751459000 | 19.777420000 |
| 8 | 12.441220000 | 5.264542000 | 18.533453000 |
| 1 | 11.507450000 | 5.601139000 | 18.598024000 |
| 1 | 12.963152000 | 6.040591000 | 18.230315000 |
| 1 | 7.553274000 | -4.124877000 | 21.259873000 |
| 1 | 5.868298000 | -5.722122000 | 19.458695000 |
| 1 | 0.070460000 | -1.731717000 | 18.602068000 |
| 1 | 0.853507000 | 1.616081000 | 15.557271000 |
| 1 | 0.433160000 | 4.306376000 | 19.534662000 |
| 1 | 5.550554000 | 7.815681000 | 16.715121000 |
| 1 | 18.524551000 | 4.234692000 | 10.513858000 |
| 1 | 14.961169000 | 7.513442000 | 8.509264000 |
| 1 | 10.890293000 | 1.914431000 | 6.225833000 |
| 1 | 5.503710000 | 0.865985000 | 7.208835000 |
| 1 | 10.427522000 | -1.585347000 | 9.566172000 |
| 1 | 4.425286000 | -3.379652000 | 21.212928000 |
| 1 | 8.633959000 | 8.873314000 | 10.579234000 |
| 1 | 13.436785000 | 9.152325000 | 9.190825000 |
| 1 | 15.799104000 | 10.034274000 | 13.494371000 |
| 1 | 2.232279000 | -0.722642000 | 10.692131000 |
| 1 | 5.543063000 | -4.461905000 | 11.946967000 |
| 1 | 8.993230000 | -4.047087000 | 13.240123000 |
| 8 | 7.934606000 | 5.424915000 | 14.318493000 |
| 1 | 8.063130000 | 5.784835000 | 15.230268000 |
| 1 | 7.675181000 | 4.497799000 | 14.484862000 |
| 8 | 13.414734000 | 1.661502000 | 16.800470000 |
| 1 | 13.279138000 | 1.948135000 | 17.744039000 |
| 1 | 13.014831000 | 0.775256000 | 16.681210000 |
| 8 | 9.793527000 | 1.446106000 | 16.749698000 |

**$^{1}$IM2$_{S}$:**

| | | | |
|---|---|---|---|
| 6 | 14.564826000 | 9.370242000 | 13.114032000 |
| 6 | 13.407665000 | 9.487134000 | 12.137287000 |
| 8 | 12.291341000 | 9.914479000 | 12.466004000 |
| 6 | 14.154795000 | 8.624144000 | 14.388829000 |
| 6 | 13.616039000 | 7.214384000 | 14.119185000 |
| 6 | 13.379305000 | 6.455720000 | 15.416266000 |
| 8 | 13.127474000 | 7.054119000 | 16.474843000 |
| 7 | 13.427160000 | 5.105042000 | 15.361239000 |
| 1 | 15.414149000 | 8.873692000 | 12.636651000 |
| 1 | 15.018674000 | 8.565073000 | 15.059091000 |
| 1 | 13.391274000 | 9.195928000 | 14.918843000 |
| 1 | 14.307720000 | 6.656907000 | 13.476251000 |
| 1 | 12.670333000 | 7.263605000 | 13.561929000 |
| 1 | 13.508102000 | 4.584438000 | 16.242220000 |
| 1 | 13.833048000 | 4.623951000 | 14.563737000 |
| 7 | 13.669443000 | 9.076302000 | 10.874338000 |
| 6 | 12.586558000 | 9.012813000 | 9.914759000 |
| 6 | 11.467749000 | 8.100697000 | 10.448690000 |
| 8 | 11.750553000 | 7.023531000 | 10.985428000 |
| 1 | 14.555611000 | 8.595590000 | 10.696547000 |
| 1 | 12.206260000 | 10.018105000 | 9.704719000 |
| 7 | 10.205231000 | 8.540668000 | 10.279831000 |
| 6 | 9.103357000 | 8.030459000 | 11.104188000 |
| 6 | 9.162770000 | 8.722788000 | 12.473726000 |
| 6 | 7.883656000 | 8.680583000 | 13.303940000 |
| 6 | 8.042947000 | 9.476855000 | 14.607453000 |
| 7 | 8.971668000 | 8.853289000 | 15.545464000 |
| 6 | 10.216128000 | 9.230030000 | 15.815370000 |
| 7 | 10.862551000 | 10.156161000 | 15.054507000 |
| 7 | 10.832137000 | 8.752288000 | 16.908227000 |
| 1 | 10.115272000 | 9.508631000 | 9.996961000 |
| 1 | 9.207912000 | 6.947494000 | 11.190560000 |
| 1 | 9.988309000 | 8.269167000 | 13.032748000 |
| 1 | 9.437129000 | 9.774016000 | 12.304525000 |
| 1 | 7.054015000 | 9.126100000 | 12.736604000 |
| 1 | 7.616718000 | 7.643674000 | 13.533399000 |
| 1 | 8.389678000 | 10.495399000 | 14.405449000 |
| 1 | 7.077592000 | 9.552024000 | 15.120398000 |
| 1 | 8.648476000 | 7.989708000 | 16.026604000 |
| 1 | 11.751656000 | 10.463669000 | 15.424417000 |
| 1 | 10.883101000 | 10.064837000 | 14.041769000 |
| 1 | 10.314139000 | 8.089007000 | 17.505864000 |
| 1 | 11.822822000 | 8.525208000 | 16.823407000 |
| 7 | 6.086400000 | 0.944578000 | 8.269143000 |
| 6 | 7.226354000 | 1.863908000 | 8.325145000 |
| 6 | 8.559463000 | 1.117621000 | 8.118347000 |
| 8 | 8.691422000 | -0.061497000 | 8.447993000 |
| 6 | 7.300218000 | 2.541185000 | 9.713662000 |
| 6 | 6.387693000 | 3.721058000 | 9.976311000 |
| 6 | 5.155243000 | 3.913993000 | 9.340857000 |
| 6 | 6.777313000 | 4.641762000 | 10.959395000 |
| 6 | 4.326806000 | 4.984472000 | 9.683786000 |
| 6 | 5.953701000 | 5.705855000 | 11.317677000 |
| 6 | 4.718617000 | 5.871734000 | 10.688229000 |
| 8 | 3.848385000 | 6.879523000 | 11.037254000 |
| 1 | 6.357223000 | 0.140811000 | 8.841431000 |
| 1 | 7.106976000 | 2.631095000 | 7.549596000 |
| 1 | 7.076197000 | 1.759232000 | 10.447921000 |
| 1 | 8.327927000 | 2.858894000 | 9.918841000 |
| 1 | 4.829736000 | 3.203807000 | 8.587548000 |
| 1 | 7.738072000 | 4.520168000 | 11.449939000 |
| 1 | 3.368657000 | 5.132016000 | 9.194599000 |
| 1 | 6.267746000 | 6.383306000 | 12.108280000 |
| 1 | 4.277539000 | 7.428245000 | 11.711938000 |
| 7 | 9.588682000 | 1.853444000 | 7.630671000 |
| 6 | 10.980540000 | 1.445220000 | 7.808323000 |
| 6 | 11.687709000 | 2.676003000 | 8.357617000 |
| 8 | 11.824178000 | 3.673437000 | 7.646582000 |
| 1 | 9.443945000 | 2.831102000 | 7.409815000 |
| 1 | 10.996628000 | 0.587706000 | 8.483708000 |

| | | | |
|---|---|---|---|
| 7 | 12.024664000 | 2.667759000 | 9.666555000 |
| 6 | 12.389984000 | 3.943612000 | 10.244708000 |
| 6 | 13.783295000 | 4.370909000 | 9.760918000 |
| 8 | 14.718811000 | 3.593133000 | 9.558015000 |
| 6 | 12.275816000 | 3.967105000 | 11.770372000 |
| 6 | 10.816824000 | 4.175379000 | 12.191717000 |
| 8 | 9.895688000 | 4.009628000 | 11.391064000 |
| 7 | 10.617850000 | 4.566691000 | 13.468252000 |
| 1 | 11.850512000 | 1.855060000 | 10.252315000 |
| 1 | 11.682516000 | 4.680649000 | 9.859686000 |
| 1 | 12.869296000 | 4.801025000 | 12.159257000 |
| 1 | 12.673066000 | 3.051850000 | 12.218592000 |
| 1 | 11.389953000 | 4.712609000 | 14.105164000 |
| 1 | 9.661575000 | 4.835861000 | 13.739517000 |
| 7 | 13.881714000 | 5.712333000 | 9.581263000 |
| 6 | 15.084797000 | 6.306218000 | 9.038376000 |
| 6 | 16.125680000 | 6.489182000 | 10.139290000 |
| 8 | 16.151689000 | 7.514258000 | 10.836284000 |
| 1 | 13.147758000 | 6.297684000 | 9.982102000 |
| 1 | 15.458649000 | 5.659463000 | 8.240039000 |
| 7 | 16.937659000 | 5.423423000 | 10.302855000 |
| 6 | 17.834049000 | 5.262639000 | 11.439445000 |
| 6 | 17.439076000 | 4.052613000 | 12.297708000 |
| 6 | 16.004872000 | 4.173586000 | 12.808657000 |
| 16 | 15.421193000 | 2.779540000 | 13.839563000 |
| 6 | 16.442621000 | 3.031783000 | 15.331393000 |
| 1 | 16.640910000 | 4.583515000 | 9.808328000 |
| 1 | 17.783658000 | 6.190691000 | 12.014935000 |
| 1 | 17.534740000 | 3.129357000 | 11.713592000 |
| 1 | 18.145977000 | 3.977393000 | 13.131789000 |
| 1 | 15.881172000 | 5.106716000 | 13.372587000 |
| 1 | 15.316028000 | 4.194809000 | 11.961974000 |
| 1 | 16.145552000 | 2.276189000 | 16.060938000 |
| 1 | 16.258123000 | 4.020700000 | 15.762141000 |
| 1 | 17.505821000 | 2.913898000 | 15.108436000 |
| 6 | 4.756524000 | -3.274263000 | 20.693917000 |
| 6 | 4.177863000 | -2.915081000 | 19.334885000 |
| 8 | 4.835255000 | -2.988497000 | 18.294516000 |
| 6 | 5.165605000 | -2.018062000 | 21.506047000 |
| 6 | 6.148333000 | -1.146697000 | 20.787831000 |
| 7 | 7.413163000 | -0.840055000 | 21.268095000 |
| 6 | 6.075170000 | -0.510687000 | 19.573453000 |
| 6 | 8.045283000 | -0.052573000 | 20.357268000 |
| 7 | 7.253934000 | 0.156554000 | 19.319010000 |
| 1 | 4.022181000 | -3.843498000 | 21.275872000 |
| 1 | 4.271394000 | -1.431514000 | 21.752334000 |
| 1 | 5.587221000 | -2.349161000 | 22.461385000 |
| 1 | 5.282625000 | -0.514847000 | 18.846607000 |
| 1 | 9.051988000 | 0.321135000 | 20.468431000 |
| 1 | 7.794408000 | -1.134365000 | 22.156777000 |
| 7 | 2.888263000 | -2.469544000 | 19.336425000 |
| 6 | 2.224997000 | -2.058350000 | 18.127367000 |
| 6 | 2.025047000 | -0.542135000 | 18.002909000 |
| 8 | 2.458954000 | 0.255137000 | 18.842805000 |
| 1 | 2.407507000 | -2.356748000 | 20.217385000 |
| 1 | 2.856315000 | -2.385185000 | 17.296392000 |
| 7 | 1.329518000 | -0.190145000 | 16.897581000 |
| 6 | 1.358845000 | 1.146147000 | 16.305885000 |
| 6 | 2.399822000 | 1.222325000 | 15.170429000 |
| 6 | 3.741536000 | 0.759927000 | 15.633032000 |
| 7 | 4.243084000 | -0.498750000 | 15.341095000 |
| 6 | 4.677028000 | 1.348351000 | 16.444830000 |
| 6 | 5.452439000 | -0.618590000 | 15.930073000 |
| 7 | 5.736367000 | 0.479337000 | 16.623115000 |
| 1 | 1.088663000 | -0.943280000 | 16.262798000 |
| 1 | 1.618434000 | 1.846539000 | 17.099990000 |
| 1 | 2.075527000 | 0.609688000 | 14.320998000 |
| 1 | 2.464998000 | 2.250243000 | 14.800394000 |
| 1 | 4.703753000 | 2.345886000 | 16.847443000 |
| 1 | 6.077791000 | -1.490285000 | 15.851700000 |

| | | | |
|---|---|---|---|
| 1 | 3.888561000 | -1.133249000 | 14.610706000 |
| 6 | 5.253062000 | -5.529976000 | 14.799162000 |
| 6 | 3.722369000 | -5.485128000 | 14.920321000 |
| 6 | 5.928415000 | -4.232016000 | 15.256605000 |
| 6 | 3.227372000 | -5.454490000 | 16.370084000 |
| 1 | 5.646272000 | -6.377021000 | 15.379302000 |
| 1 | 3.296745000 | -6.360046000 | 14.413049000 |
| 1 | 3.353722000 | -4.600503000 | 14.382751000 |
| 1 | 5.813870000 | -4.065521000 | 16.329026000 |
| 1 | 5.474528000 | -3.374066000 | 14.748816000 |
| 1 | 7.003692000 | -4.238157000 | 15.036704000 |
| 1 | 2.132944000 | -5.402284000 | 16.413060000 |
| 1 | 3.633069000 | -4.597380000 | 16.915901000 |
| 1 | 3.534581000 | -6.360699000 | 16.907574000 |
| 7 | 7.198860000 | -4.867110000 | 18.744791000 |
| 6 | 8.646564000 | -4.769525000 | 18.589571000 |
| 6 | 9.198297000 | -4.398692000 | 17.192384000 |
| 6 | 8.984734000 | -2.959175000 | 16.841424000 |
| 7 | 9.591063000 | -2.330093000 | 15.759308000 |
| 6 | 8.262005000 | -1.968638000 | 17.454360000 |
| 6 | 9.246320000 | -1.017765000 | 15.754336000 |
| 7 | 8.436033000 | -0.772817000 | 16.772265000 |
| 1 | 6.701317000 | -4.034552000 | 18.435750000 |
| 1 | 9.075229000 | -5.735592000 | 18.875392000 |
| 1 | 8.740331000 | -5.053659000 | 16.437474000 |
| 1 | 10.276673000 | -4.611934000 | 17.172895000 |
| 1 | 7.623998000 | -2.033182000 | 18.316317000 |
| 1 | 9.600866000 | -0.297152000 | 15.035287000 |
| 1 | 10.235581000 | -2.756005000 | 15.106656000 |
| 7 | 9.599376000 | -1.937373000 | 10.785604000 |
| 6 | 8.794969000 | -1.220775000 | 11.778181000 |
| 6 | 7.342663000 | -1.578121000 | 11.456116000 |
| 8 | 6.771793000 | -1.219178000 | 10.425568000 |
| 6 | 9.052802000 | 0.314242000 | 11.827057000 |
| 6 | 10.410965000 | 0.607998000 | 12.383946000 |
| 7 | 11.568813000 | 0.237276000 | 11.708522000 |
| 6 | 10.749004000 | 1.127586000 | 13.617947000 |
| 6 | 12.570861000 | 0.504309000 | 12.538319000 |
| 7 | 12.125989000 | 1.049223000 | 13.698567000 |
| 1 | 9.325213000 | -1.576522000 | 9.868208000 |
| 1 | 9.032837000 | -1.631702000 | 12.771745000 |
| 1 | 8.961519000 | 0.686748000 | 10.807534000 |
| 1 | 8.299266000 | 0.825010000 | 12.431378000 |
| 1 | 10.163044000 | 1.588890000 | 14.400523000 |
| 1 | 13.620348000 | 0.344161000 | 12.333817000 |
| 1 | 12.683144000 | 1.239241000 | 14.544346000 |
| 7 | 6.683952000 | -2.364621000 | 12.360297000 |
| 6 | 5.438855000 | -3.018638000 | 11.966675000 |
| 6 | 4.208814000 | -2.112131000 | 12.127731000 |
| 8 | 3.551484000 | -2.098578000 | 13.186378000 |
| 1 | 7.197633000 | -2.754238000 | 13.137125000 |
| 1 | 5.528826000 | -3.345306000 | 10.925815000 |
| 7 | 3.919870000 | -1.386490000 | 11.038720000 |
| 6 | 2.871589000 | -0.374062000 | 10.948193000 |
| 6 | 3.430922000 | 1.028622000 | 10.647066000 |
| 6 | 4.166522000 | 1.670819000 | 11.798316000 |
| 6 | 5.289668000 | 1.063304000 | 12.371500000 |
| 6 | 3.779928000 | 2.919927000 | 12.295738000 |
| 6 | 5.987395000 | 1.654681000 | 13.416676000 |
| 6 | 4.484647000 | 3.541114000 | 13.322987000 |
| 6 | 5.583428000 | 2.899189000 | 13.897177000 |
| 8 | 6.248438000 | 3.522563000 | 14.918731000 |
| 1 | 4.648401000 | -1.372242000 | 10.328146000 |
| 1 | 2.324825000 | -0.394580000 | 11.893608000 |
| 1 | 2.586955000 | 1.664437000 | 10.355396000 |
| 1 | 4.091381000 | 0.974777000 | 9.768890000 |
| 1 | 5.647366000 | 0.129604000 | 11.972330000 |
| 1 | 2.940552000 | 3.439926000 | 11.844125000 |
| 1 | 6.838073000 | 1.162633000 | 13.873979000 |
| 1 | 4.212772000 | 4.533408000 | 13.656605000 |

| | | | |
|---|---|---|---|
| 1 | 6.945650000 | 2.911085000 | 15.275705000 |
| 6 | 5.324183000 | 7.191371000 | 16.984734000 |
| 6 | 4.727383000 | 6.460230000 | 15.776361000 |
| 6 | 3.627713000 | 5.491120000 | 16.232293000 |
| 6 | 4.197011000 | 7.445955000 | 14.727673000 |
| 1 | 4.546523000 | 7.747194000 | 17.528313000 |
| 1 | 5.812409000 | 6.492386000 | 17.671585000 |
| 1 | 5.526976000 | 5.871069000 | 15.316315000 |
| 1 | 3.121854000 | 5.012167000 | 15.388110000 |
| 1 | 2.860549000 | 6.025905000 | 16.812196000 |
| 1 | 4.052628000 | 4.698618000 | 16.856751000 |
| 1 | 3.795585000 | 6.921092000 | 13.851193000 |
| 1 | 3.392592000 | 8.071777000 | 15.135789000 |
| 1 | 4.996841000 | 8.114897000 | 14.385936000 |
| 6 | 1.599537000 | 4.301334000 | 19.464033000 |
| 7 | 2.527669000 | 3.184906000 | 19.380088000 |
| 6 | 3.862348000 | 3.308900000 | 19.468903000 |
| 7 | 4.454019000 | 4.502974000 | 19.517684000 |
| 7 | 4.610489000 | 2.197882000 | 19.570970000 |
| 1 | 1.809009000 | 4.912818000 | 20.348309000 |
| 1 | 1.640984000 | 4.935046000 | 18.569982000 |
| 1 | 2.181844000 | 2.236897000 | 19.250752000 |
| 1 | 3.949434000 | 5.324630000 | 19.219949000 |
| 1 | 5.492697000 | 4.545202000 | 19.484462000 |
| 1 | 4.126901000 | 1.317260000 | 19.434248000 |
| 1 | 5.557055000 | 2.255207000 | 19.180326000 |
| 26 | 7.683749000 | 1.079984000 | 17.228163000 |
| 6 | 9.202880000 | 3.554766000 | 18.326103000 |
| 6 | 9.560012000 | 4.378961000 | 17.085905000 |
| 6 | 9.216126000 | 5.861818000 | 17.289163000 |
| 6 | 7.702361000 | 3.592310000 | 18.603407000 |
| 8 | 9.799106000 | 6.501505000 | 18.202116000 |
| 8 | 8.369570000 | 6.374439000 | 16.497090000 |
| 8 | 7.233803000 | 4.475436000 | 19.327708000 |
| 8 | 6.950633000 | 2.687726000 | 18.033721000 |
| 16 | 9.760369000 | 1.782947000 | 18.124426000 |
| 1 | 9.740326000 | 3.948432000 | 19.190357000 |
| 1 | 10.638868000 | 4.286064000 | 16.923642000 |
| 1 | 9.050204000 | 3.981791000 | 16.207954000 |
| 8 | 8.100473000 | 1.725592000 | 15.778840000 |
| 8 | 15.359820000 | 1.075457000 | 18.060157000 |
| 1 | 14.671241000 | 0.785184000 | 17.416818000 |
| 1 | 16.169213000 | 0.603416000 | 17.816499000 |
| 8 | 13.301681000 | 1.161723000 | 16.247168000 |
| 1 | 12.412206000 | 1.035702000 | 16.652304000 |
| 1 | 13.486405000 | 2.095130000 | 16.534925000 |
| 8 | 12.192748000 | 3.526451000 | 19.558330000 |
| 1 | 12.242269000 | 4.529774000 | 19.417256000 |
| 1 | 12.640494000 | 3.372106000 | 20.404503000 |
| 8 | 12.303820000 | 6.069218000 | 19.031415000 |
| 1 | 11.351585000 | 6.236695000 | 18.818001000 |
| 1 | 12.768622000 | 6.278841000 | 18.195586000 |
| 1 | 9.023324000 | -4.037428000 | 19.314913000 |
| 1 | 6.830765000 | -5.638463000 | 18.191544000 |
| 1 | 1.255756000 | -2.563089000 | 18.030136000 |
| 1 | 0.358865000 | 1.398973000 | 15.939838000 |
| 1 | 0.588642000 | 3.901487000 | 19.555024000 |
| 1 | 6.094278000 | 7.906232000 | 16.680253000 |
| 1 | 18.863750000 | 5.145398000 | 11.081942000 |
| 1 | 14.849786000 | 7.284022000 | 8.612396000 |
| 1 | 11.437211000 | 1.162883000 | 6.854070000 |
| 1 | 6.005920000 | 0.580359000 | 7.319281000 |
| 1 | 10.568972000 | -1.639750000 | 10.911559000 |
| 1 | 5.625179000 | -3.913721000 | 20.512963000 |
| 1 | 8.164266000 | 8.238346000 | 10.581888000 |
| 1 | 12.973107000 | 8.598387000 | 8.978922000 |
| 1 | 14.881283000 | 10.386540000 | 13.379528000 |
| 1 | 2.179444000 | -0.670985000 | 10.151013000 |
| 1 | 5.287509000 | -3.891042000 | 12.598202000 |
| 1 | 5.523894000 | -5.734994000 | 13.754594000 |

| | | | |
|---|---|---|---|
| 8 | 7.995333000 | 5.501061000 | 14.020223000 |
| 1 | 8.091840000 | 5.796564000 | 14.961636000 |
| 1 | 7.330533000 | 4.789673000 | 14.084189000 |
| 8 | 13.966865000 | 3.401550000 | 17.572653000 |
| 1 | 13.297571000 | 3.336680000 | 18.312498000 |
| 1 | 14.739556000 | 2.879740000 | 17.876173000 |
| 8 | 10.823602000 | 1.738577000 | 17.020864000 |

**$^5Re_R$:**

| | | | |
|---|---|---|---|
| 6 | 15.395149000 | 7.257553000 | 14.142615000 |
| 6 | 15.098823000 | 7.988951000 | 12.838840000 |
| 8 | 14.292995000 | 8.929108000 | 12.789002000 |
| 6 | 14.102346000 | 6.920076000 | 14.890020000 |
| 6 | 13.270301000 | 5.880295000 | 14.118916000 |
| 6 | 11.943532000 | 5.645119000 | 14.799108000 |
| 8 | 11.852527000 | 4.921733000 | 15.818840000 |
| 7 | 10.884461000 | 6.291159000 | 14.289422000 |
| 1 | 15.991623000 | 6.358652000 | 13.950566000 |
| 1 | 14.328900000 | 6.541133000 | 15.892437000 |
| 1 | 13.519171000 | 7.839240000 | 15.005219000 |
| 1 | 13.104463000 | 6.220498000 | 13.093287000 |
| 1 | 13.804788000 | 4.926249000 | 14.084867000 |
| 1 | 9.968487000 | 6.171877000 | 14.725916000 |
| 1 | 10.917627000 | 6.568201000 | 13.307729000 |
| 7 | 15.783069000 | 7.553022000 | 11.757453000 |
| 6 | 15.691310000 | 8.252222000 | 10.485503000 |
| 6 | 14.410446000 | 7.897119000 | 9.720431000 |
| 8 | 14.403563000 | 7.023581000 | 8.835686000 |
| 1 | 16.374532000 | 6.721456000 | 11.823995000 |
| 1 | 15.735922000 | 9.330261000 | 10.668732000 |
| 7 | 13.317500000 | 8.579192000 | 10.088815000 |
| 6 | 11.992975000 | 8.272060000 | 9.551924000 |
| 6 | 10.917223000 | 8.836532000 | 10.474805000 |
| 6 | 9.520045000 | 8.331058000 | 10.111296000 |
| 6 | 8.403114000 | 8.907218000 | 10.989127000 |
| 7 | 8.526731000 | 8.524631000 | 12.399758000 |
| 6 | 9.046418000 | 9.266070000 | 13.366393000 |
| 7 | 9.647443000 | 10.447407000 | 13.098174000 |
| 7 | 8.921289000 | 8.891710000 | 14.651549000 |
| 1 | 13.402773000 | 9.103757000 | 10.958281000 |
| 1 | 11.876922000 | 7.190975000 | 9.460644000 |
| 1 | 11.157448000 | 8.529953000 | 11.498862000 |
| 1 | 10.949024000 | 9.938418000 | 10.430805000 |
| 1 | 9.514723000 | 7.239472000 | 10.192146000 |
| 1 | 9.283992000 | 8.579531000 | 9.068717000 |
| 1 | 8.360931000 | 9.999663000 | 10.922479000 |
| 1 | 7.435588000 | 8.529164000 | 10.647273000 |
| 1 | 8.212259000 | 7.558753000 | 12.617576000 |
| 1 | 9.970528000 | 11.024673000 | 13.859774000 |
| 1 | 10.056022000 | 10.612230000 | 12.191297000 |
| 1 | 8.489717000 | 7.987616000 | 14.881099000 |
| 1 | 9.638008000 | 9.180291000 | 15.304162000 |
| 7 | 6.455219000 | 2.715863000 | 8.083509000 |
| 6 | 7.539304000 | 3.652725000 | 7.809135000 |
| 6 | 8.877364000 | 2.948472000 | 7.596419000 |
| 8 | 9.218893000 | 1.999673000 | 8.307482000 |
| 6 | 7.792156000 | 4.611390000 | 9.007216000 |
| 6 | 6.640287000 | 5.533283000 | 9.298955000 |
| 6 | 5.664065000 | 5.199363000 | 10.247606000 |
| 6 | 6.506598000 | 6.751404000 | 8.617456000 |
| 6 | 4.590485000 | 6.054638000 | 10.510569000 |
| 6 | 5.439479000 | 7.609878000 | 8.862500000 |
| 6 | 4.470590000 | 7.261066000 | 9.811805000 |
| 8 | 3.446371000 | 8.140205000 | 10.004485000 |
| 1 | 6.838752000 | 1.963501000 | 8.656803000 |
| 1 | 7.271968000 | 4.247097000 | 6.927492000 |
| 1 | 8.007892000 | 3.988513000 | 9.877307000 |
| 1 | 8.696889000 | 5.191892000 | 8.792676000 |
| 1 | 5.758991000 | 4.260859000 | 10.785148000 |
| 1 | 7.254646000 | 7.032441000 | 7.879440000 |
| 1 | 3.845146000 | 5.779338000 | 11.252909000 |
| 1 | 5.333355000 | 8.549213000 | 8.329441000 |
| 1 | 2.861744000 | 7.807335000 | 10.702965000 |
| 7 | 9.703549000 | 3.478202000 | 6.648477000 |
| 6 | 11.131947000 | 3.281553000 | 6.816500000 |
| 6 | 11.599517000 | 4.087991000 | 8.037474000 |
| 8 | 11.111422000 | 5.187016000 | 8.285014000 |
| 1 | 9.437077000 | 4.386887000 | 6.290491000 |
| 1 | 11.329445000 | 2.213905000 | 6.927920000 |
| 7 | 12.562375000 | 3.507126000 | 8.791342000 |
| 6 | 13.114024000 | 4.147110000 | 9.979422000 |
| 6 | 14.576675000 | 3.710624000 | 10.065922000 |
| 8 | 14.884265000 | 2.563330000 | 10.423229000 |
| 6 | 12.341210000 | 3.762546000 | 11.256920000 |
| 6 | 11.178544000 | 4.705286000 | 11.549757000 |
| 8 | 11.365487000 | 5.935614000 | 11.533966000 |
| 7 | 10.020285000 | 4.136346000 | 11.928385000 |
| 1 | 12.830545000 | 2.547708000 | 8.614339000 |
| 1 | 13.047398000 | 5.223478000 | 9.824334000 |
| 1 | 13.008611000 | 3.834015000 | 12.122920000 |
| 1 | 12.013382000 | 2.723169000 | 11.204241000 |
| 1 | 9.852209000 | 3.131151000 | 11.752013000 |
| 1 | 9.204319000 | 4.745324000 | 12.052466000 |
| 7 | 15.485367000 | 4.639414000 | 9.721825000 |
| 6 | 16.901967000 | 4.331794000 | 9.779843000 |
| 6 | 17.342549000 | 4.165697000 | 11.239026000 |
| 8 | 17.286804000 | 5.108747000 | 12.036654000 |
| 1 | 15.175156000 | 5.546661000 | 9.345413000 |
| 1 | 17.125370000 | 3.436006000 | 9.190672000 |
| 7 | 17.773270000 | 2.931315000 | 11.577999000 |
| 6 | 18.032727000 | 2.540117000 | 12.958710000 |
| 6 | 16.826021000 | 1.826528000 | 13.592698000 |
| 6 | 15.598975000 | 2.732359000 | 13.664538000 |
| 16 | 14.055908000 | 1.923244000 | 14.233792000 |
| 6 | 13.771843000 | 0.701471000 | 12.901552000 |
| 1 | 17.570829000 | 2.191094000 | 10.918219000 |
| 1 | 18.280783000 | 3.453142000 | 13.504084000 |
| 1 | 16.592049000 | 0.932418000 | 13.002275000 |
| 1 | 17.104820000 | 1.480427000 | 14.595771000 |
| 1 | 15.387287000 | 3.157021000 | 12.684310000 |
| 1 | 15.768392000 | 3.560103000 | 14.360777000 |
| 1 | 14.441822000 | -0.154916000 | 13.000622000 |
| 1 | 12.743432000 | 0.353691000 | 13.012595000 |
| 1 | 13.916766000 | 1.188631000 | 11.934560000 |
| 6 | 12.798135000 | 5.503038000 | 19.659434000 |
| 6 | 11.475792000 | 4.796436000 | 19.930787000 |
| 7 | 10.826725000 | 4.426906000 | 18.642208000 |
| 1 | 12.635001000 | 6.414559000 | 19.075162000 |
| 1 | 13.469797000 | 4.848600000 | 19.093910000 |
| 1 | 10.777868000 | 5.433968000 | 20.480171000 |
| 1 | 11.616488000 | 3.875885000 | 20.503857000 |
| 1 | 11.457985000 | 3.799528000 | 18.101863000 |
| 1 | 10.621802000 | 5.294821000 | 18.100692000 |
| 1 | 9.925092000 | 3.906836000 | 18.792464000 |
| 6 | 5.792585000 | -3.212548000 | 23.216558000 |
| 6 | 5.022465000 | -3.723737000 | 22.003237000 |
| 8 | 5.572191000 | -4.312541000 | 21.070621000 |
| 6 | 5.419489000 | -1.808880000 | 23.738685000 |
| 6 | 5.750194000 | -0.718194000 | 22.765314000 |
| 7 | 6.489971000 | 0.406085000 | 23.112380000 |
| 6 | 5.511619000 | -0.575824000 | 21.419577000 |
| 6 | 6.670641000 | 1.160629000 | 21.994959000 |
| 7 | 6.099044000 | 0.584099000 | 20.952649000 |
| 1 | 5.646212000 | -3.944475000 | 24.021675000 |
| 1 | 4.360410000 | -1.769756000 | 24.030212000 |
| 1 | 5.973647000 | -1.644543000 | 24.669925000 |
| 1 | 4.952315000 | -1.221154000 | 20.762781000 |
| 1 | 7.224540000 | 2.087571000 | 21.966290000 |
| 1 | 6.824935000 | 0.630537000 | 24.039174000 |
| 7 | 3.669546000 | -3.572885000 | 22.042132000 |

| | | | |
|---|---|---|---|
| 6 | 2.837196000 | -4.079160000 | 20.978032000 |
| 6 | 2.776122000 | -3.134753000 | 19.768901000 |
| 8 | 2.848962000 | -1.912226000 | 19.913975000 |
| 1 | 3.250495000 | -2.972058000 | 22.735338000 |
| 1 | 3.214236000 | -5.062178000 | 20.683846000 |
| 7 | 2.576249000 | -3.751626000 | 18.582716000 |
| 6 | 2.466415000 | -3.054148000 | 17.306150000 |
| 6 | 3.802276000 | -3.001551000 | 16.540511000 |
| 6 | 4.889759000 | -2.340225000 | 17.319647000 |
| 7 | 5.733036000 | -3.023145000 | 18.183991000 |
| 6 | 5.253911000 | -1.024576000 | 17.443243000 |
| 6 | 6.562611000 | -2.136222000 | 18.783193000 |
| 7 | 6.284846000 | -0.907554000 | 18.361463000 |
| 1 | 2.573978000 | -4.762981000 | 18.568710000 |
| 1 | 2.125606000 | -2.039237000 | 17.508778000 |
| 1 | 3.635640000 | -2.450116000 | 15.609853000 |
| 1 | 4.106680000 | -4.013642000 | 16.246991000 |
| 1 | 4.868606000 | -0.158515000 | 16.933658000 |
| 1 | 7.344985000 | -2.456180000 | 19.457557000 |
| 1 | 5.702217000 | -4.004610000 | 18.421071000 |
| 6 | 7.217944000 | -3.423173000 | 13.686930000 |
| 6 | 5.902642000 | -2.669457000 | 13.446442000 |
| 6 | 7.169390000 | -4.378437000 | 14.883620000 |
| 6 | 4.746319000 | -3.551773000 | 12.968839000 |
| 1 | 7.488591000 | -3.979660000 | 12.778862000 |
| 1 | 5.623749000 | -2.159217000 | 14.378299000 |
| 1 | 6.052186000 | -1.878897000 | 12.701254000 |
| 1 | 8.131343000 | -4.882418000 | 15.027529000 |
| 1 | 6.407915000 | -5.156322000 | 14.752752000 |
| 1 | 6.940816000 | -3.827260000 | 15.803907000 |
| 1 | 5.021228000 | -4.087960000 | 12.053234000 |
| 1 | 4.461378000 | -4.300764000 | 13.716783000 |
| 1 | 3.866913000 | -2.944155000 | 12.731941000 |
| 7 | 8.578125000 | -4.203562000 | 20.364259000 |
| 6 | 9.602072000 | -3.824186000 | 21.341355000 |
| 6 | 10.690497000 | -2.947585000 | 20.700311000 |
| 6 | 10.122820000 | -1.691022000 | 20.119161000 |
| 7 | 10.676671000 | -1.029627000 | 19.037180000 |
| 6 | 8.970020000 | -1.006008000 | 20.416120000 |
| 6 | 9.865425000 | -0.003625000 | 18.707804000 |
| 7 | 8.804622000 | 0.028237000 | 19.514892000 |
| 1 | 7.740478000 | -4.560530000 | 20.821666000 |
| 1 | 10.089694000 | -4.680585000 | 21.833343000 |
| 1 | 11.197731000 | -3.510655000 | 19.905884000 |
| 1 | 11.458627000 | -2.726818000 | 21.454073000 |
| 1 | 8.222596000 | -1.215585000 | 21.163963000 |
| 1 | 10.050879000 | 0.678603000 | 17.894376000 |
| 1 | 11.524584000 | -1.305664000 | 18.485181000 |
| 7 | 9.805254000 | 1.310387000 | 11.200199000 |
| 6 | 8.587859000 | 0.588079000 | 11.569040000 |
| 6 | 7.370151000 | 1.469344000 | 11.269031000 |
| 8 | 7.352390000 | 2.666201000 | 11.564539000 |
| 6 | 8.523280000 | 0.309027000 | 13.093228000 |
| 6 | 9.793800000 | -0.036023000 | 13.808575000 |
| 7 | 10.363836000 | 0.886611000 | 14.661914000 |
| 6 | 10.506701000 | -1.202614000 | 14.030472000 |
| 6 | 11.336690000 | 0.267738000 | 15.368233000 |
| 7 | 11.463492000 | -1.003733000 | 15.007002000 |
| 1 | 9.809456000 | 1.460654000 | 10.188298000 |
| 1 | 8.495010000 | -0.362811000 | 11.021606000 |
| 1 | 8.110522000 | 1.197035000 | 13.579782000 |
| 1 | 7.792491000 | -0.488873000 | 13.251066000 |
| 1 | 10.367322000 | -2.170793000 | 13.569999000 |
| 1 | 11.902299000 | 0.784731000 | 16.127796000 |
| 1 | 10.045782000 | 1.860119000 | 14.808816000 |
| 7 | 6.307966000 | 0.833769000 | 10.719329000 |
| 6 | 5.000951000 | 1.457750000 | 10.670496000 |
| 6 | 3.934320000 | 0.379466000 | 10.794509000 |
| 8 | 4.225808000 | -0.797343000 | 10.991421000 |
| 1 | 6.288002000 | -0.180969000 | 10.677254000 |
| 1 | 4.899721000 | 2.138722000 | 11.522768000 |
| 7 | 2.651800000 | 0.841436000 | 10.701039000 |
| 6 | 1.573900000 | 0.148985000 | 11.397115000 |
| 6 | 1.270826000 | 0.804662000 | 12.775614000 |
| 6 | 2.541627000 | 1.243735000 | 13.469665000 |
| 6 | 3.495617000 | 0.303950000 | 13.894645000 |
| 6 | 2.901355000 | 2.599975000 | 13.523092000 |
| 6 | 4.758472000 | 0.695975000 | 14.317875000 |
| 6 | 4.162401000 | 3.013641000 | 13.958719000 |
| 6 | 5.112464000 | 2.053804000 | 14.337808000 |
| 8 | 6.362958000 | 2.364116000 | 14.719726000 |
| 1 | 2.555915000 | 1.847157000 | 10.623796000 |
| 1 | 1.913767000 | -0.879805000 | 11.524440000 |
| 1 | 0.618894000 | 1.674421000 | 12.628094000 |
| 1 | 0.700017000 | 0.081753000 | 13.373778000 |
| 1 | 3.265290000 | -0.756549000 | 13.834601000 |
| 1 | 2.186640000 | 3.353447000 | 13.195762000 |
| 1 | 5.500678000 | -0.031928000 | 14.618978000 |
| 1 | 4.418917000 | 4.065522000 | 14.004419000 |
| 1 | 6.493020000 | 3.348924000 | 14.774158000 |
| 6 | 3.218958000 | 7.933865000 | 13.353895000 |
| 6 | 4.292236000 | 7.196318000 | 14.163132000 |
| 6 | 5.347078000 | 8.186787000 | 14.669645000 |
| 6 | 3.660564000 | 6.411246000 | 15.318296000 |
| 1 | 3.661238000 | 8.514034000 | 12.536867000 |
| 1 | 2.664073000 | 8.632494000 | 13.991879000 |
| 1 | 4.802018000 | 6.484535000 | 13.497404000 |
| 1 | 5.857618000 | 8.681693000 | 13.834303000 |
| 1 | 4.882791000 | 8.967580000 | 15.285491000 |
| 1 | 6.103297000 | 7.686400000 | 15.279835000 |
| 1 | 4.391965000 | 5.785583000 | 15.836357000 |
| 1 | 3.209469000 | 7.095558000 | 16.048606000 |
| 1 | 2.862486000 | 5.749060000 | 14.958013000 |
| 6 | 0.353770000 | -0.200096000 | 18.578103000 |
| 7 | 1.142527000 | 0.341928000 | 17.464989000 |
| 6 | 2.266744000 | 1.089014000 | 17.678057000 |
| 7 | 3.003358000 | 0.898719000 | 18.778989000 |
| 7 | 2.612206000 | 2.010032000 | 16.790084000 |
| 1 | 0.968232000 | -0.853913000 | 19.196965000 |
| 1 | -0.070458000 | 0.594749000 | 19.203237000 |
| 1 | 0.580815000 | 0.682987000 | 16.694937000 |
| 1 | 2.908833000 | 0.013514000 | 19.271027000 |
| 1 | 3.958303000 | 1.278789000 | 18.737259000 |
| 1 | 3.471279000 | 2.593730000 | 16.909989000 |
| 1 | 2.142656000 | 2.069016000 | 15.895216000 |
| 26 | 7.106052000 | 0.896384000 | 18.920514000 |
| 6 | 7.491515000 | 5.491485000 | 15.863403000 |
| 6 | 7.193394000 | 5.063580000 | 17.294464000 |
| 6 | 7.157407000 | 3.544410000 | 17.515942000 |
| 6 | 5.750532000 | 2.966142000 | 17.528847000 |
| 8 | 6.871500000 | 4.966793000 | 14.899380000 |
| 8 | 8.349111000 | 6.426536000 | 15.704986000 |
| 8 | 4.787370000 | 3.615275000 | 17.091674000 |
| 8 | 5.617555000 | 1.763448000 | 18.003818000 |
| 16 | 7.998802000 | 3.012308000 | 19.082398000 |
| 1 | 7.938579000 | 5.520104000 | 17.946384000 |
| 1 | 6.217628000 | 5.485253000 | 17.560865000 |
| 1 | 7.688243000 | 3.047779000 | 16.699856000 |
| 8 | 7.976747000 | 0.339805000 | 16.328234000 |
| 8 | 12.772255000 | -1.690769000 | 17.415908000 |
| 1 | 13.413047000 | -0.952474000 | 17.373624000 |
| 1 | 12.376157000 | -1.718847000 | 16.509535000 |
| 8 | 7.764143000 | 5.893529000 | 12.567171000 |
| 1 | 7.395116000 | 5.540537000 | 13.429744000 |
| 1 | 7.088054000 | 5.717447000 | 11.886777000 |
| 1 | 13.286712000 | 5.773680000 | 20.599949000 |
| 1 | 16.008662000 | 7.930346000 | 14.754813000 |
| 1 | 9.108348000 | -3.251789000 | 22.137040000 |
| 1 | 8.932726000 | -4.938131000 | 19.754650000 |
| 1 | 6.850289000 | -3.232476000 | 22.944981000 |

| | | | |
|---|---|---|---|
| 1 | 1.818541000 | -4.205049000 | 21.359957000 |
| 1 | 1.703976000 | -3.557373000 | 16.702675000 |
| 1 | -0.457383000 | -0.794977000 | 18.153824000 |
| 8 | 7.854420000 | -0.807066000 | 15.927303000 |
| 1 | 2.485186000 | 7.230507000 | 12.934169000 |
| 1 | 0.672781000 | 0.132014000 | 10.776347000 |
| 1 | 4.882651000 | 2.053564000 | 9.756877000 |
| 1 | 10.607176000 | 0.730247000 | 11.441658000 |
| 1 | 6.159469000 | 2.279260000 | 7.211459000 |
| 1 | 11.653257000 | 3.642341000 | 5.925116000 |
| 1 | 17.448990000 | 5.174959000 | 9.351703000 |
| 1 | 18.908414000 | 1.883669000 | 12.977645000 |
| 1 | 16.541754000 | 7.962572000 | 9.866102000 |
| 1 | 11.901237000 | 8.697275000 | 8.544888000 |
| 1 | 8.020990000 | -2.696519000 | 13.859891000 |
| 8 | 9.590213000 | 3.502157000 | 15.068763000 |
| 1 | 10.371848000 | 3.957912000 | 15.458579000 |
| 1 | 9.554781000 | 3.826762000 | 14.149637000 |
| 8 | 12.473832000 | 2.711474000 | 17.234611000 |
| 1 | 12.650560000 | 3.337072000 | 16.503025000 |
| 1 | 13.266922000 | 2.152391000 | 17.371466000 |
| 8 | 10.080007000 | 6.969066000 | 17.675322000 |
| 1 | 9.578407000 | 7.407789000 | 18.379692000 |
| 1 | 9.408309000 | 6.789822000 | 16.961318000 |
| 8 | 14.391667000 | 0.687163000 | 17.152211000 |
| 1 | 15.300076000 | 0.626173000 | 17.488265000 |
| 1 | 14.471187000 | 0.845376000 | 16.179866000 |

**$^{5}$TS1$_{R}$:**

| | | | |
|---|---|---|---|
| 6 | 15.181470000 | 7.202979000 | 14.276438000 |
| 6 | 14.969581000 | 7.948592000 | 12.964503000 |
| 8 | 14.183625000 | 8.903104000 | 12.876950000 |
| 6 | 13.841293000 | 6.832284000 | 14.919512000 |
| 6 | 13.072603000 | 5.813568000 | 14.058553000 |
| 6 | 11.690518000 | 5.583112000 | 14.617248000 |
| 8 | 11.501937000 | 4.855737000 | 15.622563000 |
| 7 | 10.678794000 | 6.227848000 | 14.025188000 |
| 1 | 15.803824000 | 6.315838000 | 14.116283000 |
| 1 | 14.000372000 | 6.419603000 | 15.921484000 |
| 1 | 13.242174000 | 7.742813000 | 15.023081000 |
| 1 | 12.997344000 | 6.170623000 | 13.027862000 |
| 1 | 13.601504000 | 4.856847000 | 14.053293000 |
| 1 | 9.739876000 | 6.102248000 | 14.410967000 |
| 1 | 10.780185000 | 6.499077000 | 13.047491000 |
| 7 | 15.699837000 | 7.501074000 | 11.917730000 |
| 6 | 15.668444000 | 8.186485000 | 10.636309000 |
| 6 | 14.408600000 | 7.835145000 | 9.835308000 |
| 8 | 14.406434000 | 6.925314000 | 8.989484000 |
| 1 | 16.269793000 | 6.656884000 | 12.011855000 |
| 1 | 15.723252000 | 9.266027000 | 10.807237000 |
| 7 | 13.327932000 | 8.567889000 | 10.138084000 |
| 6 | 11.998492000 | 8.271773000 | 9.607980000 |
| 6 | 10.945375000 | 8.802540000 | 10.577718000 |
| 6 | 9.531887000 | 8.322755000 | 10.247765000 |
| 6 | 8.452748000 | 8.901143000 | 11.170647000 |
| 7 | 8.594980000 | 8.482282000 | 12.571426000 |
| 6 | 9.156076000 | 9.188638000 | 13.543789000 |
| 7 | 9.828029000 | 10.331128000 | 13.282677000 |
| 7 | 8.998811000 | 8.822740000 | 14.830067000 |
| 1 | 13.409549000 | 9.120286000 | 10.990018000 |
| 1 | 11.891149000 | 7.192811000 | 9.484825000 |
| 1 | 11.219131000 | 8.465034000 | 11.583921000 |
| 1 | 10.982982000 | 9.905526000 | 10.567915000 |
| 1 | 9.512449000 | 7.230258000 | 10.307574000 |
| 1 | 9.265654000 | 8.595593000 | 9.218907000 |
| 1 | 8.441518000 | 9.995798000 | 11.135212000 |
| 1 | 7.467667000 | 8.557687000 | 10.842125000 |
| 1 | 8.241159000 | 7.528557000 | 12.774079000 |
| 1 | 10.175405000 | 10.883136000 | 14.052735000 |
| 1 | 10.269769000 | 10.464379000 | 12.385865000 |
| 1 | 8.511338000 | 7.941078000 | 15.016852000 |
| 1 | 9.773192000 | 8.979954000 | 15.464677000 |
| 7 | 6.411459000 | 2.862130000 | 7.911173000 |
| 6 | 7.563555000 | 3.727511000 | 7.681759000 |
| 6 | 8.865247000 | 2.942947000 | 7.514276000 |
| 8 | 9.135913000 | 1.997944000 | 8.260814000 |
| 6 | 7.832674000 | 4.656009000 | 8.898029000 |
| 6 | 6.727345000 | 5.631478000 | 9.194785000 |
| 6 | 5.739681000 | 5.335181000 | 10.143682000 |
| 6 | 6.660371000 | 6.869226000 | 8.540743000 |
| 6 | 4.723441000 | 6.247574000 | 10.439830000 |
| 6 | 5.649134000 | 7.784842000 | 8.816468000 |
| 6 | 4.671840000 | 7.475837000 | 9.770785000 |
| 8 | 3.709882000 | 8.414231000 | 10.002453000 |
| 1 | 6.715791000 | 2.090674000 | 8.505976000 |
| 1 | 7.366311000 | 4.343138000 | 6.796281000 |
| 1 | 7.995836000 | 4.007305000 | 9.759783000 |
| 1 | 8.772133000 | 5.190727000 | 8.715478000 |
| 1 | 5.785991000 | 4.382539000 | 10.662346000 |
| 1 | 7.416904000 | 7.121791000 | 7.801389000 |
| 1 | 3.971491000 | 6.002569000 | 11.185998000 |
| 1 | 5.595097000 | 8.740345000 | 8.304760000 |
| 1 | 3.088040000 | 8.083538000 | 10.669084000 |
| 7 | 9.737332000 | 3.389809000 | 6.565674000 |
| 6 | 11.150280000 | 3.117547000 | 6.759050000 |
| 6 | 11.651559000 | 3.944342000 | 7.954901000 |
| 8 | 11.175450000 | 5.050717000 | 8.190513000 |
| 1 | 9.527859000 | 4.295307000 | 6.164893000 |
| 1 | 11.286724000 | 2.045529000 | 6.912794000 |
| 7 | 12.629992000 | 3.376296000 | 8.701771000 |
| 6 | 13.137520000 | 4.022218000 | 9.906666000 |
| 6 | 14.591145000 | 3.586040000 | 10.087505000 |
| 8 | 14.880723000 | 2.435771000 | 10.448183000 |
| 6 | 12.293688000 | 3.659232000 | 11.145253000 |
| 6 | 11.134364000 | 4.624688000 | 11.354139000 |
| 8 | 11.347060000 | 5.849736000 | 11.344206000 |
| 7 | 9.939388000 | 4.087533000 | 11.654496000 |
| 1 | 12.877960000 | 2.406265000 | 8.554014000 |
| 1 | 13.085596000 | 5.097699000 | 9.737858000 |
| 1 | 12.910630000 | 3.735000000 | 12.047802000 |
| 1 | 11.955262000 | 2.622807000 | 11.086806000 |
| 1 | 9.750297000 | 3.080113000 | 11.525108000 |
| 1 | 9.141505000 | 4.715933000 | 11.782847000 |
| 7 | 15.514027000 | 4.530966000 | 9.834020000 |
| 6 | 16.923954000 | 4.237167000 | 10.005152000 |
| 6 | 17.255761000 | 4.094229000 | 11.495326000 |
| 8 | 17.141271000 | 5.049591000 | 12.272204000 |
| 1 | 15.218235000 | 5.451356000 | 9.480134000 |
| 1 | 17.194872000 | 3.334542000 | 9.447929000 |
| 7 | 17.661085000 | 2.867450000 | 11.886593000 |
| 6 | 17.842572000 | 2.507408000 | 13.288561000 |
| 6 | 16.605086000 | 1.800833000 | 13.867514000 |
| 6 | 15.374215000 | 2.705288000 | 13.869270000 |
| 16 | 13.793248000 | 1.871022000 | 14.275378000 |
| 6 | 13.678769000 | 0.646296000 | 12.920646000 |
| 1 | 17.514013000 | 2.113503000 | 11.227652000 |
| 1 | 18.059246000 | 3.431836000 | 13.827815000 |
| 1 | 16.405977000 | 0.901823000 | 13.272430000 |
| 1 | 16.829461000 | 1.463751000 | 14.886516000 |
| 1 | 15.243220000 | 3.169657000 | 12.891925000 |
| 1 | 15.477830000 | 3.503065000 | 14.611201000 |
| 1 | 14.348693000 | -0.199528000 | 13.088669000 |
| 1 | 12.650044000 | 0.281846000 | 12.919485000 |
| 1 | 13.912197000 | 1.134366000 | 11.971280000 |
| 6 | 12.639658000 | 5.927606000 | 20.036184000 |
| 6 | 11.449974000 | 4.980785000 | 20.123469000 |
| 7 | 10.925020000 | 4.684085000 | 18.758067000 |
| 1 | 12.348709000 | 6.880420000 | 19.580411000 |
| 1 | 13.436701000 | 5.484729000 | 19.430641000 |
| 1 | 10.626790000 | 5.402206000 | 20.706774000 |

| | | | |
|---|---|---|---|
| 1 | 11.727405000 | 4.024951000 | 20.575778000 |
| 1 | 11.684705000 | 4.243606000 | 18.187937000 |
| 1 | 10.604959000 | 5.564995000 | 18.294940000 |
| 1 | 10.102621000 | 4.036223000 | 18.812515000 |
| 6 | 5.756089000 | -2.847925000 | 23.556433000 |
| 6 | 5.276951000 | -3.560013000 | 22.296858000 |
| 8 | 6.043935000 | -4.009119000 | 21.442743000 |
| 6 | 5.187232000 | -1.430648000 | 23.762385000 |
| 6 | 5.604701000 | -0.489433000 | 22.677318000 |
| 7 | 5.237989000 | 0.850876000 | 22.640333000 |
| 6 | 6.380145000 | -0.670210000 | 21.560624000 |
| 6 | 5.795464000 | 1.415922000 | 21.534275000 |
| 7 | 6.490896000 | 0.516843000 | 20.863758000 |
| 1 | 5.491408000 | -3.465195000 | 24.422795000 |
| 1 | 4.090325000 | -1.462660000 | 23.838248000 |
| 1 | 5.536182000 | -1.066827000 | 24.737933000 |
| 1 | 6.854806000 | -1.572772000 | 21.215644000 |
| 1 | 5.697947000 | 2.455340000 | 21.256805000 |
| 1 | 4.669872000 | 1.331311000 | 23.324189000 |
| 7 | 3.929121000 | -3.737914000 | 22.195247000 |
| 6 | 3.371222000 | -4.323581000 | 21.004751000 |
| 6 | 3.309520000 | -3.333057000 | 19.830539000 |
| 8 | 3.352580000 | -2.116528000 | 20.016672000 |
| 1 | 3.317175000 | -3.197503000 | 22.788956000 |
| 1 | 3.981497000 | -5.188553000 | 20.731003000 |
| 7 | 3.150195000 | -3.921407000 | 18.621853000 |
| 6 | 3.014904000 | -3.198571000 | 17.362645000 |
| 6 | 4.332425000 | -3.119429000 | 16.571240000 |
| 6 | 5.424499000 | -2.404572000 | 17.295238000 |
| 7 | 6.304832000 | -3.020991000 | 18.175035000 |
| 6 | 5.774166000 | -1.078551000 | 17.323666000 |
| 6 | 7.136496000 | -2.087477000 | 18.695273000 |
| 7 | 6.829254000 | -0.892808000 | 18.197560000 |
| 1 | 3.143455000 | -4.932470000 | 18.585015000 |
| 1 | 2.661387000 | -2.193345000 | 17.586153000 |
| 1 | 4.134898000 | -2.586993000 | 15.636930000 |
| 1 | 4.652857000 | -4.128544000 | 16.286826000 |
| 1 | 5.342920000 | -0.245800000 | 16.794478000 |
| 1 | 7.924162000 | -2.339277000 | 19.397481000 |
| 1 | 6.294024000 | -3.989884000 | 18.459671000 |
| 6 | 6.621227000 | -3.586824000 | 12.976601000 |
| 6 | 5.319636000 | -2.778056000 | 12.899436000 |
| 6 | 6.690493000 | -4.596667000 | 14.126109000 |
| 6 | 4.066095000 | -3.614376000 | 12.629494000 |
| 1 | 6.775192000 | -4.106956000 | 12.021492000 |
| 1 | 5.185107000 | -2.217591000 | 13.835357000 |
| 1 | 5.397036000 | -2.017252000 | 12.113237000 |
| 1 | 7.636223000 | -5.149202000 | 14.106878000 |
| 1 | 5.879885000 | -5.332216000 | 14.074348000 |
| 1 | 6.629392000 | -4.083750000 | 15.093648000 |
| 1 | 4.190244000 | -4.224709000 | 11.727520000 |
| 1 | 3.833428000 | -4.293667000 | 13.458490000 |
| 1 | 3.199012000 | -2.965703000 | 12.464572000 |
| 7 | 9.005905000 | -3.796396000 | 20.886207000 |
| 6 | 9.865435000 | -3.257099000 | 21.944727000 |
| 6 | 11.044491000 | -2.444841000 | 21.362578000 |
| 6 | 10.531082000 | -1.412655000 | 20.410877000 |
| 7 | 10.736185000 | -1.484381000 | 19.046960000 |
| 6 | 9.576299000 | -0.434331000 | 20.583073000 |
| 6 | 9.916810000 | -0.603511000 | 18.445839000 |
| 7 | 9.181013000 | 0.041981000 | 19.346315000 |
| 1 | 8.111941000 | -4.100480000 | 21.270925000 |
| 1 | 10.263482000 | -4.021992000 | 22.630589000 |
| 1 | 11.740452000 | -3.112033000 | 20.840119000 |
| 1 | 11.602709000 | -1.993601000 | 22.191581000 |
| 1 | 9.096518000 | -0.094924000 | 21.487131000 |
| 1 | 9.855125000 | -0.498332000 | 17.376762000 |
| 1 | 11.445605000 | -1.996641000 | 18.488292000 |
| 7 | 9.661744000 | 1.215107000 | 11.112056000 |
| 6 | 8.384697000 | 0.536859000 | 11.336284000 |

| | | | |
|---|---|---|---|
| 6 | 7.238382000 | 1.528001000 | 11.113032000 |
| 8 | 7.298392000 | 2.697404000 | 11.497709000 |
| 6 | 8.215979000 | 0.023227000 | 12.800591000 |
| 6 | 9.448542000 | -0.368834000 | 13.551146000 |
| 7 | 10.179986000 | 0.581819000 | 14.236372000 |
| 6 | 10.004373000 | -1.579298000 | 13.922713000 |
| 6 | 11.092330000 | -0.067567000 | 14.998326000 |
| 7 | 11.026780000 | -1.381879000 | 14.829227000 |
| 1 | 9.748115000 | 1.396174000 | 10.108919000 |
| 1 | 8.260368000 | -0.315247000 | 10.651216000 |
| 1 | 7.716814000 | 0.798810000 | 13.390226000 |
| 1 | 7.543968000 | -0.836892000 | 12.757862000 |
| 1 | 9.716318000 | -2.572789000 | 13.608787000 |
| 1 | 11.755018000 | 0.442359000 | 15.679108000 |
| 1 | 9.967192000 | 1.596183000 | 14.266162000 |
| 7 | 6.136389000 | 1.003754000 | 10.521394000 |
| 6 | 4.863274000 | 1.694968000 | 10.537475000 |
| 6 | 3.749422000 | 0.663179000 | 10.674424000 |
| 8 | 3.992799000 | -0.537022000 | 10.772968000 |
| 1 | 6.058869000 | -0.002295000 | 10.404542000 |
| 1 | 4.833626000 | 2.360929000 | 11.406614000 |
| 7 | 2.489508000 | 1.186019000 | 10.699904000 |
| 6 | 1.413636000 | 0.477684000 | 11.383743000 |
| 6 | 1.199681000 | 1.018087000 | 12.826103000 |
| 6 | 2.515961000 | 1.324672000 | 13.506402000 |
| 6 | 3.420612000 | 0.301488000 | 13.833153000 |
| 6 | 2.959897000 | 2.649788000 | 13.648139000 |
| 6 | 4.713073000 | 0.586886000 | 14.255728000 |
| 6 | 4.250041000 | 2.955557000 | 14.083467000 |
| 6 | 5.145794000 | 1.916259000 | 14.377326000 |
| 8 | 6.417972000 | 2.129346000 | 14.768984000 |
| 1 | 2.433474000 | 2.197505000 | 10.704290000 |
| 1 | 1.713102000 | -0.571100000 | 11.404379000 |
| 1 | 0.592080000 | 1.930531000 | 12.783005000 |
| 1 | 0.610406000 | 0.274866000 | 13.379548000 |
| 1 | 3.127324000 | -0.736193000 | 13.697195000 |
| 1 | 2.285253000 | 3.466391000 | 13.396498000 |
| 1 | 5.417676000 | -0.207449000 | 14.470850000 |
| 1 | 4.571701000 | 3.982788000 | 14.201255000 |
| 1 | 6.611730000 | 3.102047000 | 14.800076000 |
| 6 | 3.359131000 | 7.991584000 | 13.364589000 |
| 6 | 4.372139000 | 7.216153000 | 14.215576000 |
| 6 | 5.388787000 | 8.182596000 | 14.834568000 |
| 6 | 3.658935000 | 6.390053000 | 15.292508000 |
| 1 | 3.858126000 | 8.595768000 | 12.598786000 |
| 1 | 2.770191000 | 8.673875000 | 13.989682000 |
| 1 | 4.925479000 | 6.527955000 | 13.560478000 |
| 1 | 5.951762000 | 8.714478000 | 14.057710000 |
| 1 | 4.883253000 | 8.935210000 | 15.452709000 |
| 1 | 6.105323000 | 7.655859000 | 15.470357000 |
| 1 | 4.352779000 | 5.749785000 | 15.844359000 |
| 1 | 3.151705000 | 7.048395000 | 16.009674000 |
| 1 | 2.893065000 | 5.739998000 | 14.850012000 |
| 6 | 0.704594000 | -0.641852000 | 18.657776000 |
| 7 | 1.430962000 | -0.013660000 | 17.548976000 |
| 6 | 2.514945000 | 0.795030000 | 17.751071000 |
| 7 | 3.318022000 | 0.608424000 | 18.804627000 |
| 7 | 2.758576000 | 1.774281000 | 16.891427000 |
| 1 | 1.399405000 | -1.138697000 | 19.333982000 |
| 1 | 0.112301000 | 0.088714000 | 19.222132000 |
| 1 | 0.826581000 | 0.317601000 | 16.806933000 |
| 1 | 3.301925000 | -0.293545000 | 19.278502000 |
| 1 | 4.247050000 | 1.048644000 | 18.727221000 |
| 1 | 3.582623000 | 2.407099000 | 17.006628000 |
| 1 | 2.249437000 | 1.838927000 | 16.018728000 |
| 26 | 7.451452000 | 0.935139000 | 18.881529000 |
| 6 | 7.459415000 | 5.394358000 | 15.824251000 |
| 6 | 7.152161000 | 5.098775000 | 17.286534000 |
| 6 | 7.207464000 | 3.599588000 | 17.608622000 |
| 6 | 5.845896000 | 2.926191000 | 17.637057000 |

| | | | |
|---|---|---|---|
| 8 | 6.835230000 | 4.801863000 | 14.905338000 |
| 8 | 8.357380000 | 6.271949000 | 15.590981000 |
| 8 | 4.825173000 | 3.502783000 | 17.230064000 |
| 8 | 5.831285000 | 1.710701000 | 18.097261000 |
| 16 | 8.064509000 | 3.182967000 | 19.197783000 |
| 1 | 7.863503000 | 5.643487000 | 17.908690000 |
| 1 | 6.148280000 | 5.477604000 | 17.506468000 |
| 1 | 7.760757000 | 3.089649000 | 16.814318000 |
| 8 | 9.426806000 | 1.675139000 | 16.692725000 |
| 8 | 12.579582000 | -2.094938000 | 17.135873000 |
| 1 | 13.002034000 | -1.213307000 | 17.164616000 |
| 1 | 12.145520000 | -2.111399000 | 16.249780000 |
| 8 | 7.754914000 | 5.843330000 | 12.574052000 |
| 1 | 7.395447000 | 5.415540000 | 13.403828000 |
| 1 | 7.059724000 | 5.751961000 | 11.896730000 |
| 1 | 13.033157000 | 6.132119000 | 21.036271000 |
| 1 | 15.734607000 | 7.874634000 | 14.944441000 |
| 1 | 9.251664000 | -2.576551000 | 22.547190000 |
| 1 | 9.447812000 | -4.608197000 | 20.457639000 |
| 1 | 6.846165000 | -2.806454000 | 23.504396000 |
| 1 | 2.357905000 | -4.679261000 | 21.219721000 |
| 1 | 2.251011000 | -3.699207000 | 16.758313000 |
| 1 | 0.037344000 | -1.399511000 | 18.240897000 |
| 8 | 10.440295000 | 2.187565000 | 17.092170000 |
| 1 | 2.649016000 | 7.310488000 | 12.874198000 |
| 1 | 0.486913000 | 0.553035000 | 10.806589000 |
| 1 | 4.740675000 | 2.312857000 | 9.638597000 |
| 1 | 10.415780000 | 0.587234000 | 11.386214000 |
| 1 | 6.122565000 | 2.441745000 | 7.029094000 |
| 1 | 11.699534000 | 3.412983000 | 5.860247000 |
| 1 | 17.497987000 | 5.077171000 | 9.607080000 |
| 1 | 18.716521000 | 1.853337000 | 13.369267000 |
| 1 | 16.536957000 | 7.877185000 | 10.052269000 |
| 1 | 11.882434000 | 8.734649000 | 8.620119000 |
| 1 | 7.456738000 | -2.888637000 | 13.094877000 |
| 8 | 9.503624000 | 3.250159000 | 14.498430000 |
| 1 | 10.203379000 | 3.749489000 | 14.973582000 |
| 1 | 9.489022000 | 3.639638000 | 13.604850000 |
| 8 | 13.111300000 | 3.587414000 | 17.362821000 |
| 1 | 12.678998000 | 3.900317000 | 16.534079000 |
| 1 | 13.088158000 | 2.609819000 | 17.346336000 |
| 8 | 10.053951000 | 7.089773000 | 17.493990000 |
| 1 | 9.472440000 | 7.582812000 | 18.094446000 |
| 1 | 9.436917000 | 6.724383000 | 16.799412000 |
| 8 | 13.581700000 | 0.687217000 | 17.218622000 |
| 1 | 14.366258000 | 0.687786000 | 17.791778000 |
| 1 | 13.920349000 | 0.881305000 | 16.312182000 |

**$^{5}$IM1$_R$:**

| | | | |
|---|---|---|---|
| 6 | 16.007929000 | 5.280389000 | 13.866808000 |
| 6 | 15.925044000 | 5.756023000 | 12.422254000 |
| 8 | 15.349288000 | 6.809268000 | 12.112061000 |
| 6 | 14.639915000 | 5.351229000 | 14.550902000 |
| 6 | 13.626334000 | 4.414508000 | 13.869131000 |
| 6 | 12.268375000 | 4.554474000 | 14.508276000 |
| 8 | 12.039870000 | 4.097836000 | 15.657446000 |
| 7 | 11.344627000 | 5.238045000 | 13.828725000 |
| 1 | 16.417456000 | 4.265723000 | 13.911063000 |
| 1 | 14.738409000 | 5.078139000 | 15.606703000 |
| 1 | 14.270545000 | 6.380849000 | 14.500281000 |
| 1 | 13.558487000 | 4.634880000 | 12.801620000 |
| 1 | 13.952158000 | 3.377741000 | 13.980237000 |
| 1 | 10.427305000 | 5.377685000 | 14.255753000 |
| 1 | 11.423588000 | 5.273954000 | 12.811610000 |
| 7 | 16.521173000 | 4.949378000 | 11.515066000 |
| 6 | 16.592500000 | 5.320458000 | 10.110802000 |
| 6 | 15.262445000 | 5.069378000 | 9.388461000 |
| 8 | 15.051626000 | 4.018708000 | 8.759666000 |
| 1 | 16.907818000 | 4.050254000 | 11.811909000 |
| 1 | 16.875249000 | 6.374857000 | 10.033259000 |
| 7 | 14.357738000 | 6.050830000 | 9.514269000 |
| 6 | 12.985316000 | 5.925893000 | 9.026399000 |
| 6 | 12.087891000 | 6.891026000 | 9.796561000 |
| 6 | 10.600283000 | 6.672209000 | 9.518245000 |
| 6 | 9.682748000 | 7.660108000 | 10.249015000 |
| 7 | 9.752329000 | 7.546720000 | 11.710937000 |
| 6 | 10.475066000 | 8.309358000 | 12.520264000 |
| 7 | 11.366684000 | 9.200759000 | 12.035080000 |
| 7 | 10.274845000 | 8.265398000 | 13.849891000 |
| 1 | 14.574461000 | 6.748540000 | 10.224376000 |
| 1 | 12.641368000 | 4.899742000 | 9.164151000 |
| 1 | 12.282435000 | 6.747436000 | 10.865682000 |
| 1 | 12.374376000 | 7.924317000 | 9.535023000 |
| 1 | 10.338289000 | 5.651153000 | 9.811751000 |
| 1 | 10.396022000 | 6.764677000 | 8.444011000 |
| 1 | 9.902551000 | 8.696038000 | 9.969159000 |
| 1 | 8.642671000 | 7.466921000 | 9.971727000 |
| 1 | 9.206029000 | 6.765892000 | 12.120168000 |
| 1 | 11.851875000 | 9.809836000 | 12.676896000 |
| 1 | 11.796292000 | 9.046943000 | 11.135679000 |
| 1 | 9.613918000 | 7.574819000 | 14.220137000 |
| 1 | 11.072157000 | 8.404806000 | 14.459583000 |
| 7 | 6.361896000 | 1.575324000 | 8.535522000 |
| 6 | 7.634607000 | 2.166883000 | 8.135467000 |
| 6 | 8.768356000 | 1.142565000 | 8.115175000 |
| 8 | 8.909574000 | 0.329354000 | 9.032743000 |
| 6 | 8.104347000 | 3.253691000 | 9.140988000 |
| 6 | 7.203544000 | 4.457102000 | 9.217965000 |
| 6 | 6.224054000 | 4.564215000 | 10.214676000 |
| 6 | 7.322968000 | 5.508865000 | 8.298964000 |
| 6 | 5.396363000 | 5.687365000 | 10.299604000 |
| 6 | 6.500136000 | 6.629732000 | 8.363729000 |
| 6 | 5.528722000 | 6.723482000 | 9.368300000 |
| 8 | 4.754377000 | 7.845893000 | 9.382782000 |
| 1 | 6.557356000 | 0.900075000 | 9.275525000 |
| 1 | 7.508518000 | 2.623124000 | 7.146661000 |
| 1 | 8.173145000 | 2.776269000 | 10.119975000 |
| 1 | 9.119759000 | 3.557341000 | 8.860373000 |
| 1 | 6.127617000 | 3.759920000 | 10.936873000 |
| 1 | 8.077570000 | 5.450484000 | 7.517802000 |
| 1 | 4.649062000 | 5.753867000 | 11.087196000 |
| 1 | 6.590063000 | 7.440431000 | 7.647873000 |
| 1 | 4.130986000 | 7.798557000 | 10.124126000 |
| 7 | 9.651127000 | 1.227989000 | 7.078521000 |
| 6 | 11.002257000 | 0.743990000 | 7.303288000 |
| 6 | 11.677385000 | 1.659449000 | 8.336485000 |
| 8 | 11.445898000 | 2.865055000 | 8.343905000 |
| 1 | 9.589331000 | 2.069629000 | 6.519488000 |
| 1 | 10.955786000 | -0.293119000 | 7.640504000 |
| 7 | 12.511393000 | 1.049784000 | 9.212140000 |
| 6 | 13.135243000 | 1.782529000 | 10.307755000 |
| 6 | 14.455137000 | 1.078131000 | 10.625880000 |
| 8 | 14.479127000 | -0.006268000 | 11.226992000 |
| 6 | 12.229120000 | 1.832077000 | 11.551617000 |
| 6 | 11.313770000 | 3.048481000 | 11.592599000 |
| 8 | 11.754098000 | 4.170978000 | 11.287892000 |
| 7 | 10.081914000 | 2.850589000 | 12.097142000 |
| 1 | 12.534932000 | 0.039219000 | 9.260113000 |
| 1 | 13.322318000 | 2.794680000 | 9.950386000 |
| 1 | 12.845035000 | 1.904149000 | 12.454446000 |
| 1 | 11.663074000 | 0.903728000 | 11.646258000 |
| 1 | 9.688482000 | 1.897062000 | 12.167436000 |
| 1 | 9.431596000 | 3.641015000 | 12.101892000 |
| 7 | 15.564673000 | 1.716638000 | 10.213686000 |
| 6 | 16.877143000 | 1.159624000 | 10.486358000 |
| 6 | 17.193269000 | 1.285520000 | 11.981922000 |
| 8 | 17.387358000 | 2.389394000 | 12.504487000 |
| 1 | 15.479095000 | 2.567374000 | 9.640008000 |
| 1 | 16.927335000 | 0.118317000 | 10.152033000 |
| 7 | 17.215900000 | 0.121989000 | 12.667019000 |

| | | | |
|---|---|---|---|
| 6 | 17.285528000 | 0.059216000 | 14.122482000 |
| 6 | 15.902895000 | -0.197726000 | 14.745516000 |
| 6 | 14.940806000 | 0.959231000 | 14.485945000 |
| 16 | 13.188486000 | 0.643686000 | 14.927632000 |
| 6 | 12.801504000 | -0.851479000 | 13.950563000 |
| 1 | 16.828209000 | -0.681305000 | 12.187830000 |
| 1 | 17.701162000 | 1.010616000 | 14.460559000 |
| 1 | 15.497267000 | -1.126683000 | 14.328009000 |
| 1 | 16.020258000 | -0.360447000 | 15.824118000 |
| 1 | 14.967303000 | 1.241056000 | 13.433121000 |
| 1 | 15.217028000 | 1.832121000 | 15.085873000 |
| 1 | 13.199595000 | -1.751147000 | 14.423403000 |
| 1 | 11.713171000 | -0.927618000 | 13.932540000 |
| 1 | 13.195278000 | -0.733473000 | 12.938332000 |
| 6 | 13.497938000 | 6.349763000 | 19.692093000 |
| 6 | 12.267181000 | 5.453675000 | 19.704475000 |
| 7 | 12.038250000 | 4.865307000 | 18.353112000 |
| 1 | 13.381390000 | 7.166404000 | 18.971475000 |
| 1 | 14.394219000 | 5.780386000 | 19.423803000 |
| 1 | 11.362100000 | 6.004663000 | 19.974240000 |
| 1 | 12.375996000 | 4.623050000 | 20.406375000 |
| 1 | 12.848996000 | 4.279181000 | 18.043917000 |
| 1 | 11.848070000 | 5.614534000 | 17.652443000 |
| 1 | 11.223550000 | 4.222157000 | 18.364437000 |
| 6 | 4.285005000 | -1.560487000 | 23.523079000 |
| 6 | 4.431126000 | -2.427551000 | 22.288611000 |
| 8 | 5.502176000 | -2.601349000 | 21.712797000 |
| 6 | 3.935872000 | -0.101973000 | 23.114063000 |
| 6 | 5.041329000 | 0.582142000 | 22.376765000 |
| 7 | 5.870068000 | 1.530135000 | 22.959848000 |
| 6 | 5.535866000 | 0.419258000 | 21.106301000 |
| 6 | 6.808256000 | 1.902771000 | 22.052838000 |
| 7 | 6.626014000 | 1.242164000 | 20.923515000 |
| 1 | 3.495870000 | -1.948367000 | 24.177136000 |
| 1 | 3.038406000 | -0.112726000 | 22.485779000 |
| 1 | 3.681931000 | 0.462542000 | 24.017994000 |
| 1 | 5.202439000 | -0.256984000 | 20.337601000 |
| 1 | 7.579975000 | 2.633260000 | 22.239830000 |
| 1 | 5.780558000 | 1.895612000 | 23.897859000 |
| 7 | 3.266347000 | -2.970805000 | 21.816560000 |
| 6 | 3.222466000 | -3.513401000 | 20.486201000 |
| 6 | 3.166595000 | -2.415967000 | 19.404708000 |
| 8 | 3.132809000 | -1.218402000 | 19.701737000 |
| 1 | 2.395844000 | -2.662751000 | 22.227237000 |
| 1 | 4.123258000 | -4.114719000 | 20.331780000 |
| 7 | 3.137815000 | -2.890092000 | 18.141413000 |
| 6 | 3.135588000 | -2.071457000 | 16.934928000 |
| 6 | 4.452914000 | -2.194345000 | 16.153180000 |
| 6 | 5.633609000 | -1.659231000 | 16.901572000 |
| 7 | 6.695756000 | -2.448189000 | 17.328189000 |
| 6 | 5.943353000 | -0.393943000 | 17.339303000 |
| 6 | 7.580846000 | -1.669878000 | 17.986658000 |
| 7 | 7.148112000 | -0.417510000 | 18.018931000 |
| 1 | 3.169597000 | -3.894359000 | 18.016404000 |
| 1 | 2.971108000 | -1.037662000 | 17.238545000 |
| 1 | 4.326728000 | -1.668247000 | 15.200377000 |
| 1 | 4.624221000 | -3.244178000 | 15.886649000 |
| 1 | 5.413168000 | 0.534167000 | 17.220436000 |
| 1 | 8.495374000 | -2.045475000 | 18.405542000 |
| 1 | 6.789210000 | -3.454235000 | 17.273197000 |
| 6 | 6.554173000 | -3.711784000 | 13.451738000 |
| 6 | 5.300191000 | -3.079453000 | 12.831014000 |
| 6 | 6.587735000 | -5.240365000 | 13.362608000 |
| 6 | 5.219391000 | -3.206319000 | 11.306804000 |
| 1 | 7.446462000 | -3.300639000 | 12.959011000 |
| 1 | 4.404166000 | -3.534050000 | 13.276264000 |
| 1 | 5.252125000 | -2.017185000 | 13.103650000 |
| 1 | 7.484650000 | -5.648054000 | 13.841864000 |
| 1 | 6.585932000 | -5.583351000 | 12.323143000 |
| 1 | 5.712762000 | -5.682485000 | 13.855774000 |

| | | | |
|---|---|---|---|
| 1 | 6.110401000 | -2.779747000 | 10.825723000 |
| 1 | 5.150465000 | -4.251679000 | 10.989823000 |
| 1 | 4.339653000 | -2.678452000 | 10.926552000 |
| 7 | 7.170331000 | -4.334679000 | 19.791929000 |
| 6 | 8.045308000 | -4.609621000 | 20.955584000 |
| 6 | 9.450495000 | -4.019636000 | 20.789927000 |
| 6 | 9.491880000 | -2.531734000 | 20.614383000 |
| 7 | 10.547612000 | -1.923244000 | 19.954850000 |
| 6 | 8.618537000 | -1.508423000 | 20.916069000 |
| 6 | 10.283150000 | -0.601001000 | 19.862733000 |
| 7 | 9.113879000 | -0.307524000 | 20.422606000 |
| 1 | 6.628330000 | -3.501251000 | 20.016604000 |
| 1 | 8.157637000 | -5.693843000 | 21.056479000 |
| 1 | 9.921110000 | -4.483011000 | 19.913639000 |
| 1 | 10.053018000 | -4.317498000 | 21.659939000 |
| 1 | 7.656089000 | -1.565106000 | 21.402964000 |
| 1 | 10.925598000 | 0.110603000 | 19.370612000 |
| 1 | 11.308084000 | -2.375431000 | 19.409446000 |
| 7 | 9.192504000 | 0.066916000 | 12.083339000 |
| 6 | 7.826515000 | -0.229927000 | 12.515613000 |
| 6 | 6.895923000 | 0.869013000 | 11.990781000 |
| 8 | 7.197767000 | 2.061195000 | 12.075429000 |
| 6 | 7.680642000 | -0.208347000 | 14.064586000 |
| 6 | 8.839000000 | -0.679974000 | 14.889021000 |
| 7 | 9.515451000 | 0.218656000 | 15.690209000 |
| 6 | 9.398346000 | -1.909814000 | 15.193002000 |
| 6 | 10.412534000 | -0.465240000 | 16.430966000 |
| 7 | 10.377814000 | -1.765151000 | 16.158116000 |
| 1 | 9.241689000 | -0.008377000 | 11.064094000 |
| 1 | 7.486438000 | -1.203435000 | 12.135051000 |
| 1 | 7.465971000 | 0.817383000 | 14.373339000 |
| 1 | 6.793036000 | -0.796119000 | 14.318238000 |
| 1 | 9.140867000 | -2.881040000 | 14.794568000 |
| 1 | 11.045578000 | 0.019488000 | 17.154282000 |
| 1 | 9.401473000 | 1.245717000 | 15.683309000 |
| 7 | 5.715974000 | 0.449247000 | 11.473931000 |
| 6 | 4.643152000 | 1.385011000 | 11.203692000 |
| 6 | 3.305482000 | 0.708260000 | 11.453810000 |
| 8 | 3.226747000 | -0.447768000 | 11.863796000 |
| 1 | 5.426936000 | -0.517300000 | 11.578884000 |
| 1 | 4.732423000 | 2.226989000 | 11.898229000 |
| 7 | 2.222745000 | 1.511119000 | 11.227822000 |
| 6 | 1.002329000 | 1.319104000 | 12.003004000 |
| 6 | 0.965425000 | 2.248679000 | 13.251440000 |
| 6 | 2.331679000 | 2.391355000 | 13.888792000 |
| 6 | 3.012010000 | 1.271770000 | 14.394392000 |
| 6 | 3.043858000 | 3.599835000 | 13.819559000 |
| 6 | 4.341584000 | 1.348591000 | 14.784488000 |
| 6 | 4.374480000 | 3.701506000 | 14.233832000 |
| 6 | 5.039652000 | 2.565082000 | 14.719562000 |
| 8 | 6.319895000 | 2.574039000 | 15.128523000 |
| 1 | 2.438732000 | 2.468654000 | 10.975428000 |
| 1 | 0.998694000 | 0.268913000 | 12.298221000 |
| 1 | 0.600867000 | 3.239846000 | 12.956279000 |
| 1 | 0.221693000 | 1.840965000 | 13.949937000 |
| 1 | 2.509517000 | 0.309455000 | 14.421545000 |
| 1 | 2.551938000 | 4.484116000 | 13.418111000 |
| 1 | 4.874127000 | 0.473204000 | 15.128514000 |
| 1 | 4.900954000 | 4.646703000 | 14.188405000 |
| 1 | 6.701247000 | 3.490677000 | 15.083489000 |
| 6 | 4.461211000 | 8.315215000 | 12.754646000 |
| 6 | 5.381562000 | 7.589798000 | 13.743682000 |
| 6 | 6.618680000 | 8.444639000 | 14.042405000 |
| 6 | 4.626362000 | 7.230169000 | 15.029631000 |
| 1 | 5.003069000 | 8.604929000 | 11.847463000 |
| 1 | 4.054366000 | 9.231044000 | 13.200385000 |
| 1 | 5.730614000 | 6.658976000 | 13.273910000 |
| 1 | 7.204752000 | 8.622536000 | 13.132435000 |
| 1 | 6.327097000 | 9.422240000 | 14.446535000 |
| 1 | 7.270962000 | 7.961298000 | 14.774513000 |

| | | | |
|---|---|---|---|
| 1 | 5.225797000 | 6.592250000 | 15.686169000 |
| 1 | 4.351000000 | 8.137474000 | 15.582024000 |
| 1 | 3.697085000 | 6.690444000 | 14.807146000 |
| 6 | 0.749772000 | 0.653401000 | 18.421675000 |
| 7 | 1.511274000 | 1.469175000 | 17.474224000 |
| 6 | 2.667286000 | 2.110116000 | 17.804713000 |
| 7 | 3.462556000 | 1.630984000 | 18.767888000 |
| 7 | 2.993870000 | 3.230041000 | 17.172873000 |
| 1 | 1.380488000 | -0.122426000 | 18.855760000 |
| 1 | 0.319859000 | 1.258194000 | 19.229976000 |
| 1 | 0.945738000 | 1.981523000 | 16.810025000 |
| 1 | 3.307707000 | 0.693192000 | 19.127201000 |
| 1 | 4.434781000 | 1.978398000 | 18.788146000 |
| 1 | 3.937087000 | 3.660976000 | 17.321423000 |
| 1 | 2.515421000 | 3.470091000 | 16.313643000 |
| 26 | 7.899937000 | 1.181743000 | 19.205966000 |
| 6 | 8.126684000 | 5.603778000 | 15.722084000 |
| 6 | 7.888379000 | 5.581911000 | 17.227474000 |
| 6 | 7.779829000 | 4.168010000 | 17.813603000 |
| 6 | 6.341298000 | 3.622571000 | 17.912244000 |
| 8 | 7.326198000 | 5.037172000 | 14.933332000 |
| 8 | 9.149765000 | 6.251503000 | 15.319788000 |
| 8 | 5.411845000 | 4.368725000 | 17.543042000 |
| 8 | 6.211680000 | 2.434452000 | 18.361096000 |
| 16 | 8.604354000 | 3.991832000 | 19.450420000 |
| 1 | 8.711361000 | 6.120660000 | 17.699692000 |
| 1 | 6.955339000 | 6.117008000 | 17.431351000 |
| 1 | 8.333680000 | 3.437734000 | 17.209332000 |
| 8 | 9.551110000 | 1.851772000 | 18.332933000 |
| 8 | 12.359799000 | -2.599508000 | 18.018602000 |
| 1 | 12.786919000 | -1.721014000 | 18.024785000 |
| 1 | 11.699366000 | -2.537261000 | 17.284286000 |
| 8 | 8.362435000 | 5.262424000 | 12.444157000 |
| 1 | 7.941640000 | 5.170201000 | 13.345804000 |
| 1 | 7.648199000 | 5.128470000 | 11.793916000 |
| 1 | 13.655995000 | 6.786708000 | 20.682213000 |
| 1 | 16.712268000 | 5.940433000 | 14.387902000 |
| 1 | 7.595397000 | -4.244009000 | 21.888826000 |
| 1 | 6.483503000 | -5.086865000 | 19.741869000 |
| 1 | 5.230250000 | -1.585139000 | 24.069858000 |
| 1 | 2.354119000 | -4.172941000 | 20.387418000 |
| 1 | 2.295699000 | -2.374637000 | 16.299690000 |
| 1 | -0.058908000 | 0.167628000 | 17.872042000 |
| 8 | 9.968711000 | 3.103681000 | 18.967918000 |
| 1 | 3.605195000 | 7.685587000 | 12.472620000 |
| 1 | 0.124518000 | 1.493847000 | 11.373845000 |
| 1 | 4.720677000 | 1.785246000 | 10.184619000 |
| 1 | 9.817491000 | -0.628690000 | 12.487087000 |
| 1 | 5.980781000 | 1.037097000 | 7.758431000 |
| 1 | 11.558774000 | 0.781081000 | 6.362566000 |
| 1 | 17.616087000 | 1.741437000 | 9.931306000 |
| 1 | 17.979177000 | -0.737938000 | 14.408603000 |
| 1 | 17.358361000 | 4.712174000 | 9.626937000 |
| 1 | 12.956075000 | 6.144228000 | 7.951709000 |
| 1 | 6.631733000 | -3.407465000 | 14.504622000 |
| 8 | 9.537407000 | 2.951995000 | 15.237421000 |
| 1 | 10.409161000 | 3.280253000 | 15.551301000 |
| 1 | 9.620091000 | 2.950300000 | 14.265484000 |
| 8 | 13.851504000 | 2.998635000 | 17.327828000 |
| 1 | 13.263375000 | 3.164729000 | 16.553703000 |
| 1 | 13.582191000 | 2.129590000 | 17.689624000 |
| 8 | 11.211375000 | 7.080060000 | 16.793151000 |
| 1 | 10.866409000 | 7.776162000 | 17.373544000 |
| 1 | 10.411013000 | 6.733583000 | 16.306781000 |
| 8 | 13.423488000 | 0.156913000 | 18.011558000 |
| 1 | 14.341775000 | -0.005198000 | 18.285868000 |
| 1 | 13.438926000 | 0.143907000 | 17.019612000 |

**$^{5}$TS2$_{R}$:**

| | | | |
|---|---|---|---|
| 6 | 15.988951000 | 5.304662000 | 13.905347000 |
| 6 | 15.909811000 | 5.795312000 | 12.465257000 |
| 8 | 15.320057000 | 6.843047000 | 12.162306000 |
| 6 | 14.618407000 | 5.360741000 | 14.585732000 |
| 6 | 13.614838000 | 4.419295000 | 13.895947000 |
| 6 | 12.249305000 | 4.556545000 | 14.519848000 |
| 8 | 12.010778000 | 4.104581000 | 15.668806000 |
| 7 | 11.330429000 | 5.234136000 | 13.828032000 |
| 1 | 16.404313000 | 4.291941000 | 13.940021000 |
| 1 | 14.715871000 | 5.082401000 | 15.640308000 |
| 1 | 14.241203000 | 6.387745000 | 14.539540000 |
| 1 | 13.557622000 | 4.637989000 | 12.827484000 |
| 1 | 13.942388000 | 3.383666000 | 14.013254000 |
| 1 | 10.408576000 | 5.374811000 | 14.244853000 |
| 1 | 11.421749000 | 5.269764000 | 12.811691000 |
| 7 | 16.527605000 | 5.009491000 | 11.554468000 |
| 6 | 16.608638000 | 5.397980000 | 10.155402000 |
| 6 | 15.290451000 | 5.135847000 | 9.415314000 |
| 8 | 15.101486000 | 4.090542000 | 8.770935000 |
| 1 | 16.928471000 | 4.114188000 | 11.844295000 |
| 1 | 16.876399000 | 6.457309000 | 10.092695000 |
| 7 | 14.370269000 | 6.102304000 | 9.543827000 |
| 6 | 13.004135000 | 5.961464000 | 9.042935000 |
| 6 | 12.090549000 | 6.915606000 | 9.807125000 |
| 6 | 10.607280000 | 6.683445000 | 9.518053000 |
| 6 | 9.677567000 | 7.662288000 | 10.245609000 |
| 7 | 9.739151000 | 7.544672000 | 11.707434000 |
| 6 | 10.453042000 | 8.307588000 | 12.524667000 |
| 7 | 11.342304000 | 9.206763000 | 12.049358000 |
| 7 | 10.244247000 | 8.256494000 | 13.852531000 |
| 1 | 14.568099000 | 6.791260000 | 10.268020000 |
| 1 | 12.669837000 | 4.931572000 | 9.176916000 |
| 1 | 12.278287000 | 6.771301000 | 10.877338000 |
| 1 | 12.368861000 | 7.952146000 | 9.549670000 |
| 1 | 10.352574000 | 5.659485000 | 9.808087000 |
| 1 | 10.409155000 | 6.776282000 | 8.442728000 |
| 1 | 9.891956000 | 8.700598000 | 9.970389000 |
| 1 | 8.640684000 | 7.462403000 | 9.961141000 |
| 1 | 9.199623000 | 6.756004000 | 12.110273000 |
| 1 | 11.820921000 | 9.812805000 | 12.699012000 |
| 1 | 11.782114000 | 9.056802000 | 11.154205000 |
| 1 | 9.580826000 | 7.563237000 | 14.212146000 |
| 1 | 11.037142000 | 8.386004000 | 14.470887000 |
| 7 | 6.410588000 | 1.547720000 | 8.527929000 |
| 6 | 7.679801000 | 2.145599000 | 8.126250000 |
| 6 | 8.819943000 | 1.128505000 | 8.104785000 |
| 8 | 8.961166000 | 0.309402000 | 9.016937000 |
| 6 | 8.142364000 | 3.236353000 | 9.131077000 |
| 6 | 7.236884000 | 4.436885000 | 9.194761000 |
| 6 | 6.247968000 | 4.545983000 | 10.181678000 |
| 6 | 7.357879000 | 5.481771000 | 8.268010000 |
| 6 | 5.410885000 | 5.663367000 | 10.248441000 |
| 6 | 6.526236000 | 6.596808000 | 8.314835000 |
| 6 | 5.543836000 | 6.691567000 | 9.308661000 |
| 8 | 4.760016000 | 7.807553000 | 9.304642000 |
| 1 | 6.609907000 | 0.874197000 | 9.268502000 |
| 1 | 7.550515000 | 2.600862000 | 7.137420000 |
| 1 | 8.206199000 | 2.763081000 | 10.112545000 |
| 1 | 9.158738000 | 3.541686000 | 8.856014000 |
| 1 | 6.150539000 | 3.747627000 | 10.910348000 |
| 1 | 8.119997000 | 5.421838000 | 7.494327000 |
| 1 | 4.655913000 | 5.731590000 | 11.028431000 |
| 1 | 6.617022000 | 7.401801000 | 7.592685000 |
| 1 | 4.127875000 | 7.760588000 | 10.038531000 |
| 7 | 9.707694000 | 1.227305000 | 7.073584000 |
| 6 | 11.061752000 | 0.752899000 | 7.301617000 |
| 6 | 11.730134000 | 1.675851000 | 8.332797000 |
| 8 | 11.485582000 | 2.878989000 | 8.340480000 |
| 1 | 9.642649000 | 2.072571000 | 6.520338000 |
| 1 | 11.020851000 | -0.283263000 | 7.642313000 |
| 7 | 12.573538000 | 1.075499000 | 9.205815000 |

| | | | |
|---|---|---|---|
| 6 | 13.189950000 | 1.813760000 | 10.302146000 |
| 6 | 14.519261000 | 1.125430000 | 10.616632000 |
| 8 | 14.557500000 | 0.034853000 | 11.205484000 |
| 6 | 12.283070000 | 1.849169000 | 11.546527000 |
| 6 | 11.354387000 | 3.055839000 | 11.588580000 |
| 8 | 11.785673000 | 4.184261000 | 11.289873000 |
| 7 | 10.123127000 | 2.843802000 | 12.084585000 |
| 1 | 12.609679000 | 0.065217000 | 9.252825000 |
| 1 | 13.363417000 | 2.829252000 | 9.947314000 |
| 1 | 12.897753000 | 1.925279000 | 12.449985000 |
| 1 | 11.726881000 | 0.914283000 | 11.637317000 |
| 1 | 9.739314000 | 1.886769000 | 12.158853000 |
| 1 | 9.466534000 | 3.628400000 | 12.096842000 |
| 7 | 15.620785000 | 1.784005000 | 10.214763000 |
| 6 | 16.939936000 | 1.240319000 | 10.481791000 |
| 6 | 17.256693000 | 1.356011000 | 11.977928000 |
| 8 | 17.438345000 | 2.457138000 | 12.510880000 |
| 1 | 15.526294000 | 2.637650000 | 9.646985000 |
| 1 | 17.002322000 | 0.202795000 | 10.137896000 |
| 7 | 17.296197000 | 0.186700000 | 12.651899000 |
| 6 | 17.373569000 | 0.111952000 | 14.106531000 |
| 6 | 15.997157000 | -0.164736000 | 14.734209000 |
| 6 | 15.020394000 | 0.983869000 | 14.491859000 |
| 16 | 13.275589000 | 0.639103000 | 14.937340000 |
| 6 | 12.900243000 | -0.833058000 | 13.920917000 |
| 1 | 16.917443000 | -0.617278000 | 12.166745000 |
| 1 | 17.781224000 | 1.064645000 | 14.450641000 |
| 1 | 15.599005000 | -1.093842000 | 14.309538000 |
| 1 | 16.121471000 | -0.336781000 | 15.810486000 |
| 1 | 15.038009000 | 1.277828000 | 13.442018000 |
| 1 | 15.287485000 | 1.853861000 | 15.100048000 |
| 1 | 13.297327000 | -1.742654000 | 14.375310000 |
| 1 | 11.812414000 | -0.912529000 | 13.891168000 |
| 1 | 13.302088000 | -0.691495000 | 12.914922000 |
| 6 | 13.670764000 | 6.449271000 | 19.550716000 |
| 6 | 12.409890000 | 5.602181000 | 19.649828000 |
| 7 | 12.107233000 | 4.955929000 | 18.341972000 |
| 1 | 13.557317000 | 7.234856000 | 18.795666000 |
| 1 | 14.535479000 | 5.834228000 | 19.279466000 |
| 1 | 11.536739000 | 6.201400000 | 19.923000000 |
| 1 | 12.512077000 | 4.804287000 | 20.389567000 |
| 1 | 12.876176000 | 4.317929000 | 18.035053000 |
| 1 | 11.916741000 | 5.671229000 | 17.610217000 |
| 1 | 11.259143000 | 4.352858000 | 18.421796000 |
| 6 | 4.468859000 | -1.600451000 | 23.595184000 |
| 6 | 4.419046000 | -2.441062000 | 22.335406000 |
| 8 | 5.395557000 | -2.650931000 | 21.618067000 |
| 6 | 4.129316000 | -0.118237000 | 23.273259000 |
| 6 | 5.196699000 | 0.580026000 | 22.492436000 |
| 7 | 6.021990000 | 1.546421000 | 23.049128000 |
| 6 | 5.648215000 | 0.435122000 | 21.203176000 |
| 6 | 6.917364000 | 1.946278000 | 22.110292000 |
| 7 | 6.712691000 | 1.285496000 | 20.986159000 |
| 1 | 3.752073000 | -1.978906000 | 24.332662000 |
| 1 | 3.186051000 | -0.079858000 | 22.716734000 |
| 1 | 3.958119000 | 0.410540000 | 24.217277000 |
| 1 | 5.300647000 | -0.243052000 | 20.441189000 |
| 1 | 7.676040000 | 2.696066000 | 22.273747000 |
| 1 | 5.956901000 | 1.905755000 | 23.991538000 |
| 7 | 3.185314000 | -2.925106000 | 22.003596000 |
| 6 | 2.975079000 | -3.453690000 | 20.683061000 |
| 6 | 3.059483000 | -2.362976000 | 19.596977000 |
| 8 | 3.070397000 | -1.163245000 | 19.886335000 |
| 1 | 2.380861000 | -2.599427000 | 22.520685000 |
| 1 | 3.727463000 | -4.222722000 | 20.481589000 |
| 7 | 3.072551000 | -2.847852000 | 18.338309000 |
| 6 | 3.077210000 | -2.045575000 | 17.120985000 |
| 6 | 4.371305000 | -2.222525000 | 16.315566000 |
| 6 | 5.575352000 | -1.627967000 | 16.977228000 |
| 7 | 6.739194000 | -2.343424000 | 17.238120000 |
| 6 | 5.853057000 | -0.342939000 | 17.375663000 |
| 6 | 7.646297000 | -1.503783000 | 17.780096000 |
| 7 | 7.140666000 | -0.282988000 | 17.879530000 |
| 1 | 3.064315000 | -3.853777000 | 18.224074000 |
| 1 | 2.945261000 | -1.003501000 | 17.411279000 |
| 1 | 4.216046000 | -1.767933000 | 15.329602000 |
| 1 | 4.541246000 | -3.288596000 | 16.121888000 |
| 1 | 5.251824000 | 0.546218000 | 17.322949000 |
| 1 | 8.642101000 | -1.802452000 | 18.049842000 |
| 1 | 6.836178000 | -3.353034000 | 17.322284000 |
| 6 | 6.520811000 | -3.657184000 | 13.590461000 |
| 6 | 5.273539000 | -3.034144000 | 12.947367000 |
| 6 | 6.525239000 | -5.188613000 | 13.579024000 |
| 6 | 5.191387000 | -3.219761000 | 11.429304000 |
| 1 | 7.415099000 | -3.288323000 | 13.068501000 |
| 1 | 4.373051000 | -3.462198000 | 13.409655000 |
| 1 | 5.237338000 | -1.961237000 | 13.178797000 |
| 1 | 7.421066000 | -5.588354000 | 14.067060000 |
| 1 | 6.501382000 | -5.585516000 | 12.559073000 |
| 1 | 5.649541000 | -5.587398000 | 14.106956000 |
| 1 | 6.086509000 | -2.819528000 | 10.933113000 |
| 1 | 5.115902000 | -4.276345000 | 11.153256000 |
| 1 | 4.316874000 | -2.700050000 | 11.027544000 |
| 7 | 6.799860000 | -4.360988000 | 19.445474000 |
| 6 | 7.712150000 | -4.762401000 | 20.543934000 |
| 6 | 9.131416000 | -4.209366000 | 20.376745000 |
| 6 | 9.234900000 | -2.714298000 | 20.348114000 |
| 7 | 10.348233000 | -2.091020000 | 19.804429000 |
| 6 | 8.379159000 | -1.689378000 | 20.689534000 |
| 6 | 10.132284000 | -0.758424000 | 19.810492000 |
| 7 | 8.939708000 | -0.470672000 | 20.326919000 |
| 1 | 6.329698000 | -3.506776000 | 19.744361000 |
| 1 | 7.786954000 | -5.854296000 | 20.543825000 |
| 1 | 9.549608000 | -4.601801000 | 19.440644000 |
| 1 | 9.752526000 | -4.617446000 | 21.186751000 |
| 1 | 7.399162000 | -1.752195000 | 21.137637000 |
| 1 | 10.822598000 | -0.028162000 | 19.421593000 |
| 1 | 11.150932000 | -2.516897000 | 19.295246000 |
| 7 | 9.227808000 | 0.065468000 | 12.057408000 |
| 6 | 7.868277000 | -0.228880000 | 12.511984000 |
| 6 | 6.930878000 | 0.870080000 | 11.997299000 |
| 8 | 7.228922000 | 2.063131000 | 12.087477000 |
| 6 | 7.754703000 | -0.204422000 | 14.062594000 |
| 6 | 8.931976000 | -0.681472000 | 14.856495000 |
| 7 | 9.599968000 | 0.202786000 | 15.679221000 |
| 6 | 9.508975000 | -1.911776000 | 15.122874000 |
| 6 | 10.506131000 | -0.487985000 | 16.400921000 |
| 7 | 10.488941000 | -1.780401000 | 16.089367000 |
| 1 | 9.262697000 | -0.006384000 | 11.037322000 |
| 1 | 7.520428000 | -1.203528000 | 12.140381000 |
| 1 | 7.548697000 | 0.821568000 | 14.376072000 |
| 1 | 6.871733000 | -0.789728000 | 14.338783000 |
| 1 | 9.259820000 | -2.875799000 | 14.702380000 |
| 1 | 11.127443000 | -0.017175000 | 17.143953000 |
| 1 | 9.457438000 | 1.225139000 | 15.712058000 |
| 7 | 5.752552000 | 0.449314000 | 11.478399000 |
| 6 | 4.680630000 | 1.383666000 | 11.199233000 |
| 6 | 3.342135000 | 0.700614000 | 11.428322000 |
| 8 | 3.262908000 | -0.463967000 | 11.812516000 |
| 1 | 5.466292000 | -0.519232000 | 11.572559000 |
| 1 | 4.759160000 | 2.223230000 | 11.897994000 |
| 7 | 2.257939000 | 1.504335000 | 11.213319000 |
| 6 | 1.029703000 | 1.279887000 | 11.966579000 |
| 6 | 0.968825000 | 2.167441000 | 13.245367000 |
| 6 | 2.329574000 | 2.316843000 | 13.892902000 |
| 6 | 3.025718000 | 1.198267000 | 14.380127000 |
| 6 | 3.024589000 | 3.536413000 | 13.844517000 |
| 6 | 4.354905000 | 1.286050000 | 14.771682000 |
| 6 | 4.353146000 | 3.649509000 | 14.261289000 |
| 6 | 5.034706000 | 2.514405000 | 14.727485000 |

| | | | |
|---|---|---|---|
| 8 | 6.314065000 | 2.540319000 | 15.139408000 |
| 1 | 2.469596000 | 2.469326000 | 10.987807000 |
| 1 | 1.030415000 | 0.220823000 | 12.227917000 |
| 1 | 0.585830000 | 3.160352000 | 12.981201000 |
| 1 | 0.230911000 | 1.722006000 | 13.926529000 |
| 1 | 2.537631000 | 0.227992000 | 14.386853000 |
| 1 | 2.520539000 | 4.420350000 | 13.457701000 |
| 1 | 4.898909000 | 0.411772000 | 15.102838000 |
| 1 | 4.866853000 | 4.602444000 | 14.232404000 |
| 1 | 6.683958000 | 3.461141000 | 15.081672000 |
| 6 | 4.441941000 | 8.286614000 | 12.673303000 |
| 6 | 5.353461000 | 7.567083000 | 13.674643000 |
| 6 | 6.591372000 | 8.420494000 | 13.973829000 |
| 6 | 4.588232000 | 7.222830000 | 14.958667000 |
| 1 | 4.990539000 | 8.566160000 | 11.767077000 |
| 1 | 4.035844000 | 9.207963000 | 13.108250000 |
| 1 | 5.701986000 | 6.630138000 | 13.216945000 |
| 1 | 7.183155000 | 8.589099000 | 13.065891000 |
| 1 | 6.300852000 | 9.402222000 | 14.368567000 |
| 1 | 7.237807000 | 7.940881000 | 14.713541000 |
| 1 | 5.185456000 | 6.596676000 | 15.628377000 |
| 1 | 4.305693000 | 8.136268000 | 15.497102000 |
| 1 | 3.662559000 | 6.676875000 | 14.736192000 |
| 6 | 0.707132000 | 0.653359000 | 18.554471000 |
| 7 | 1.499045000 | 1.387205000 | 17.564563000 |
| 6 | 2.645231000 | 2.055197000 | 17.876324000 |
| 7 | 3.426590000 | 1.639093000 | 18.882324000 |
| 7 | 2.979927000 | 3.131922000 | 17.181298000 |
| 1 | 1.323842000 | -0.083488000 | 19.068327000 |
| 1 | 0.255714000 | 1.324144000 | 19.295993000 |
| 1 | 0.952209000 | 1.849280000 | 16.849345000 |
| 1 | 3.281183000 | 0.713312000 | 19.277861000 |
| 1 | 4.392059000 | 1.995782000 | 18.900033000 |
| 1 | 3.917035000 | 3.577894000 | 17.322078000 |
| 1 | 2.499055000 | 3.337835000 | 16.314298000 |
| 26 | 7.893903000 | 1.226424000 | 19.183464000 |
| 6 | 8.078560000 | 5.597627000 | 15.692790000 |
| 6 | 7.801353000 | 5.599594000 | 17.190384000 |
| 6 | 7.742313000 | 4.188469000 | 17.792481000 |
| 6 | 6.327984000 | 3.606013000 | 17.914872000 |
| 8 | 7.295504000 | 5.014549000 | 14.896238000 |
| 8 | 9.114456000 | 6.230630000 | 15.302939000 |
| 8 | 5.367324000 | 4.306630000 | 17.544122000 |
| 8 | 6.232548000 | 2.422906000 | 18.402066000 |
| 16 | 8.594026000 | 4.051671000 | 19.423711000 |
| 1 | 8.591067000 | 6.178352000 | 17.672465000 |
| 1 | 6.843009000 | 6.098313000 | 17.365077000 |
| 1 | 8.317613000 | 3.480489000 | 17.185591000 |
| 8 | 9.511740000 | 1.677911000 | 18.585338000 |
| 8 | 12.343276000 | -2.665604000 | 18.031732000 |
| 1 | 12.785058000 | -1.794863000 | 18.083117000 |
| 1 | 11.749470000 | -2.581966000 | 17.244345000 |
| 8 | 8.374429000 | 5.238840000 | 12.425974000 |
| 1 | 7.936860000 | 5.144320000 | 13.319419000 |
| 1 | 7.674353000 | 5.093402000 | 11.763068000 |
| 1 | 13.881510000 | 6.926872000 | 20.511923000 |
| 1 | 16.688326000 | 5.962670000 | 14.435798000 |
| 1 | 7.309968000 | -4.463692000 | 21.521277000 |
| 1 | 6.059424000 | -5.060693000 | 19.394414000 |
| 1 | 5.472375000 | -1.676433000 | 24.019482000 |
| 1 | 1.989338000 | -3.927559000 | 20.640075000 |
| 1 | 2.216785000 | -2.332677000 | 16.505628000 |
| 1 | -0.086467000 | 0.124379000 | 18.022959000 |
| 8 | 9.976656000 | 3.357252000 | 18.970806000 |
| 1 | 3.585456000 | 7.657647000 | 12.391414000 |
| 1 | 0.158595000 | 1.468740000 | 11.332251000 |
| 1 | 4.769666000 | 1.786013000 | 10.181776000 |
| 1 | 9.857918000 | -0.631572000 | 12.450578000 |
| 1 | 6.031012000 | 1.006957000 | 7.751876000 |
| 1 | 11.619221000 | 0.790090000 | 6.361415000 |
| 1 | 17.671290000 | 1.836117000 | 9.931504000 |
| 1 | 18.077097000 | -0.680365000 | 14.381855000 |
| 1 | 17.388783000 | 4.806554000 | 9.673478000 |
| 1 | 12.982391000 | 6.180826000 | 7.968349000 |
| 1 | 6.614205000 | -3.302374000 | 14.625203000 |
| 8 | 9.518044000 | 2.948171000 | 15.308385000 |
| 1 | 10.385503000 | 3.311346000 | 15.599368000 |
| 1 | 9.566982000 | 2.962305000 | 14.335283000 |
| 8 | 13.811514000 | 2.959061000 | 17.328545000 |
| 1 | 13.210778000 | 3.132487000 | 16.566241000 |
| 1 | 13.525212000 | 2.102477000 | 17.704965000 |
| 8 | 11.197678000 | 7.119790000 | 16.726946000 |
| 1 | 10.849217000 | 7.809890000 | 17.312490000 |
| 1 | 10.392632000 | 6.737639000 | 16.279288000 |
| 8 | 13.427953000 | 0.083911000 | 18.031712000 |
| 1 | 14.340709000 | -0.032616000 | 18.344265000 |
| 1 | 13.488670000 | 0.085996000 | 17.042441000 |

**$^{5}$IM2$_{R}$:**

| | | | |
|---|---|---|---|
| 6 | 16.253627000 | 5.551978000 | 13.886990000 |
| 6 | 16.057177000 | 6.122830000 | 12.486617000 |
| 8 | 15.389055000 | 7.148234000 | 12.288569000 |
| 6 | 14.932846000 | 5.510471000 | 14.660064000 |
| 6 | 13.963581000 | 4.477540000 | 14.055636000 |
| 6 | 12.602890000 | 4.581776000 | 14.700764000 |
| 8 | 12.349205000 | 4.032604000 | 15.798282000 |
| 7 | 11.694992000 | 5.336227000 | 14.062738000 |
| 1 | 16.717083000 | 4.560600000 | 13.834101000 |
| 1 | 15.113972000 | 5.272337000 | 15.713707000 |
| 1 | 14.477035000 | 6.504783000 | 14.616615000 |
| 1 | 13.870466000 | 4.643058000 | 12.978396000 |
| 1 | 14.341081000 | 3.463977000 | 14.221006000 |
| 1 | 10.761834000 | 5.430828000 | 14.467038000 |
| 1 | 11.790108000 | 5.453288000 | 13.054081000 |
| 7 | 16.677090000 | 5.447046000 | 11.494253000 |
| 6 | 16.693000000 | 5.951070000 | 10.130459000 |
| 6 | 15.372241000 | 5.677407000 | 9.400852000 |
| 8 | 15.225061000 | 4.688671000 | 8.662467000 |
| 1 | 17.142630000 | 4.557704000 | 11.689882000 |
| 1 | 16.899433000 | 7.025799000 | 10.148186000 |
| 7 | 14.402529000 | 6.571033000 | 9.640368000 |
| 6 | 13.040614000 | 6.397239000 | 9.141549000 |
| 6 | 12.090255000 | 7.281690000 | 9.941266000 |
| 6 | 10.623079000 | 7.007896000 | 9.611180000 |
| 6 | 9.644170000 | 7.906323000 | 10.375255000 |
| 7 | 9.695539000 | 7.692983000 | 11.824535000 |
| 6 | 10.349848000 | 8.433660000 | 12.705606000 |
| 7 | 11.150094000 | 9.451186000 | 12.311078000 |
| 7 | 10.170353000 | 8.235895000 | 14.023361000 |
| 1 | 14.572117000 | 7.192435000 | 10.430633000 |
| 1 | 12.744353000 | 5.350865000 | 9.232307000 |
| 1 | 12.265513000 | 7.088270000 | 11.005777000 |
| 1 | 12.333557000 | 8.339396000 | 9.743716000 |
| 1 | 10.402858000 | 5.959802000 | 9.839000000 |
| 1 | 10.440971000 | 7.151181000 | 8.538514000 |
| 1 | 9.820353000 | 8.966484000 | 10.164912000 |
| 1 | 8.619617000 | 7.684948000 | 10.062747000 |
| 1 | 9.227534000 | 6.819478000 | 12.138702000 |
| 1 | 11.593283000 | 10.029453000 | 13.009075000 |
| 1 | 11.583478000 | 9.425547000 | 11.400839000 |
| 1 | 9.589834000 | 7.450893000 | 14.346261000 |
| 1 | 10.931185000 | 8.467090000 | 14.648350000 |
| 7 | 6.661362000 | 1.770934000 | 8.242499000 |
| 6 | 7.897211000 | 2.458405000 | 7.884199000 |
| 6 | 9.093223000 | 1.507348000 | 7.817471000 |
| 8 | 9.227317000 | 0.591033000 | 8.631852000 |
| 6 | 8.293857000 | 3.518688000 | 8.950360000 |
| 6 | 7.313558000 | 4.653242000 | 9.083017000 |
| 6 | 6.271832000 | 4.604521000 | 10.020483000 |
| 6 | 7.418588000 | 5.794989000 | 8.275763000 |

| | | | |
|---|---|---|---|
| 6 | 5.370346000 | 5.663477000 | 10.154006000 |
| 6 | 6.522677000 | 6.853987000 | 8.391208000 |
| 6 | 5.489583000 | 6.790816000 | 9.334341000 |
| 8 | 4.643489000 | 7.858932000 | 9.403699000 |
| 1 | 6.896278000 | 1.070380000 | 8.946197000 |
| 1 | 7.750658000 | 2.959147000 | 6.919827000 |
| 1 | 8.390116000 | 2.993399000 | 9.903088000 |
| 1 | 9.286997000 | 3.904140000 | 8.694062000 |
| 1 | 6.187388000 | 3.730189000 | 10.657203000 |
| 1 | 8.219705000 | 5.856734000 | 7.542656000 |
| 1 | 4.573199000 | 5.609163000 | 10.891966000 |
| 1 | 6.601857000 | 7.734293000 | 7.761440000 |
| 1 | 3.983669000 | 7.702491000 | 10.097033000 |
| 7 | 10.036328000 | 1.774205000 | 6.867625000 |
| 6 | 11.396019000 | 1.335679000 | 7.122355000 |
| 6 | 11.988989000 | 2.186644000 | 8.256445000 |
| 8 | 11.656343000 | 3.360276000 | 8.394046000 |
| 1 | 9.956991000 | 2.673822000 | 6.410154000 |
| 1 | 11.380770000 | 0.273660000 | 7.373835000 |
| 7 | 12.886445000 | 1.564241000 | 9.058249000 |
| 6 | 13.497408000 | 2.240742000 | 10.198847000 |
| 6 | 14.875878000 | 1.608647000 | 10.404952000 |
| 8 | 15.010311000 | 0.514685000 | 10.969729000 |
| 6 | 12.621392000 | 2.136916000 | 11.464538000 |
| 6 | 11.647999000 | 3.306515000 | 11.581671000 |
| 8 | 12.060572000 | 4.465928000 | 11.411812000 |
| 7 | 10.391804000 | 3.018972000 | 11.987131000 |
| 1 | 13.001920000 | 0.560824000 | 8.995706000 |
| 1 | 13.605980000 | 3.290139000 | 9.925509000 |
| 1 | 13.257882000 | 2.178030000 | 12.356648000 |
| 1 | 12.105447000 | 1.174685000 | 11.497310000 |
| 1 | 10.022188000 | 2.052486000 | 11.903962000 |
| 1 | 9.707667000 | 3.785201000 | 11.963357000 |
| 7 | 15.915201000 | 2.309370000 | 9.914956000 |
| 6 | 17.263920000 | 1.789302000 | 10.033353000 |
| 6 | 17.734483000 | 1.839882000 | 11.492295000 |
| 8 | 17.817271000 | 2.911863000 | 12.102513000 |
| 1 | 15.749861000 | 3.182600000 | 9.396270000 |
| 1 | 17.315625000 | 0.770792000 | 9.634427000 |
| 7 | 18.043989000 | 0.651623000 | 12.052773000 |
| 6 | 18.325572000 | 0.510907000 | 13.478018000 |
| 6 | 17.086216000 | 0.049730000 | 14.263214000 |
| 6 | 15.937365000 | 1.052210000 | 14.159951000 |
| 16 | 14.314471000 | 0.455128000 | 14.770083000 |
| 6 | 14.010110000 | -0.960664000 | 13.652186000 |
| 1 | 17.756132000 | -0.177618000 | 11.549514000 |
| 1 | 18.674550000 | 1.485042000 | 13.826008000 |
| 1 | 16.769989000 | -0.923602000 | 13.870625000 |
| 1 | 17.363646000 | -0.108024000 | 15.312110000 |
| 1 | 15.780577000 | 1.346240000 | 13.122487000 |
| 1 | 16.154843000 | 1.955066000 | 14.739003000 |
| 1 | 14.730458000 | -1.761855000 | 13.831419000 |
| 1 | 13.012074000 | -1.333989000 | 13.888660000 |
| 1 | 14.068321000 | -0.620198000 | 12.616868000 |
| 6 | 13.624666000 | 5.367928000 | 19.359768000 |
| 6 | 12.319399000 | 4.848791000 | 19.950272000 |
| 7 | 11.468391000 | 4.254598000 | 18.878096000 |
| 1 | 13.424809000 | 6.131059000 | 18.600253000 |
| 1 | 14.181005000 | 4.548950000 | 18.894173000 |
| 1 | 11.741592000 | 5.648617000 | 20.422569000 |
| 1 | 12.495129000 | 4.070721000 | 20.697809000 |
| 1 | 11.984394000 | 3.457355000 | 18.437933000 |
| 1 | 11.279911000 | 4.985206000 | 18.158175000 |
| 1 | 10.538754000 | 3.906610000 | 19.239962000 |
| 6 | 4.760844000 | -1.388331000 | 23.718136000 |
| 6 | 4.035690000 | -2.162278000 | 22.625637000 |
| 8 | 4.607392000 | -2.871534000 | 21.798191000 |
| 6 | 4.754513000 | 0.146259000 | 23.504922000 |
| 6 | 5.565681000 | 0.582449000 | 22.324332000 |
| 7 | 6.566643000 | 1.542201000 | 22.399865000 |
| 6 | 5.576157000 | 0.166641000 | 21.017417000 |
| 6 | 7.145345000 | 1.677595000 | 21.180263000 |
| 7 | 6.565365000 | 0.844061000 | 20.328001000 |
| 1 | 4.290225000 | -1.611251000 | 24.683215000 |
| 1 | 3.721148000 | 0.504279000 | 23.406700000 |
| 1 | 5.151230000 | 0.615272000 | 24.412158000 |
| 1 | 4.954248000 | -0.566658000 | 20.537426000 |
| 1 | 7.940717000 | 2.374635000 | 20.923336000 |
| 1 | 6.817302000 | 2.070345000 | 23.224153000 |
| 7 | 2.677570000 | -2.033008000 | 22.626319000 |
| 6 | 1.884337000 | -2.629291000 | 21.584408000 |
| 6 | 1.934187000 | -1.841163000 | 20.262880000 |
| 8 | 2.325365000 | -0.674114000 | 20.223447000 |
| 1 | 2.247031000 | -1.348499000 | 23.230993000 |
| 1 | 2.245175000 | -3.648236000 | 21.414213000 |
| 7 | 1.462088000 | -2.531706000 | 19.198148000 |
| 6 | 1.402556000 | -2.024282000 | 17.832512000 |
| 6 | 2.555401000 | -2.551562000 | 16.958003000 |
| 6 | 3.888673000 | -2.166908000 | 17.507964000 |
| 7 | 4.687194000 | -3.014608000 | 18.260388000 |
| 6 | 4.551102000 | -0.966226000 | 17.514211000 |
| 6 | 5.789581000 | -2.342110000 | 18.670411000 |
| 7 | 5.719042000 | -1.088912000 | 18.237796000 |
| 1 | 1.200191000 | -3.498050000 | 19.345561000 |
| 1 | 1.456616000 | -0.937325000 | 17.880159000 |
| 1 | 2.435505000 | -2.142382000 | 15.948597000 |
| 1 | 2.483666000 | -3.641843000 | 16.855918000 |
| 1 | 4.281127000 | -0.031544000 | 17.052175000 |
| 1 | 6.563871000 | -2.797013000 | 19.277596000 |
| 1 | 4.516480000 | -3.993306000 | 18.440022000 |
| 6 | 6.260923000 | -3.072528000 | 14.906440000 |
| 6 | 5.134722000 | -2.702239000 | 13.929420000 |
| 6 | 6.125592000 | -4.480763000 | 15.493757000 |
| 6 | 5.196718000 | -3.452841000 | 12.596726000 |
| 1 | 7.220793000 | -3.003348000 | 14.379069000 |
| 1 | 4.163015000 | -2.888381000 | 14.410004000 |
| 1 | 5.160482000 | -1.626376000 | 13.719901000 |
| 1 | 6.907086000 | -4.677845000 | 16.236483000 |
| 1 | 6.199523000 | -5.253216000 | 14.721282000 |
| 1 | 5.152649000 | -4.605449000 | 15.986621000 |
| 1 | 6.132036000 | -3.217894000 | 12.070323000 |
| 1 | 5.162594000 | -4.539207000 | 12.732306000 |
| 1 | 4.369040000 | -3.158095000 | 11.946776000 |
| 7 | 7.140039000 | -4.434119000 | 20.976542000 |
| 6 | 8.280252000 | -4.172830000 | 21.867734000 |
| 6 | 9.566238000 | -3.868387000 | 21.079984000 |
| 6 | 9.401971000 | -2.649263000 | 20.229403000 |
| 7 | 10.110825000 | -2.407156000 | 19.062001000 |
| 6 | 8.487713000 | -1.628313000 | 20.316719000 |
| 6 | 9.604740000 | -1.306037000 | 18.478924000 |
| 7 | 8.613910000 | -0.806960000 | 19.211278000 |
| 1 | 6.271233000 | -4.165638000 | 21.434505000 |
| 1 | 8.484375000 | -4.999618000 | 22.563716000 |
| 1 | 9.814592000 | -4.724848000 | 20.439834000 |
| 1 | 10.401344000 | -3.757830000 | 21.785511000 |
| 1 | 7.710825000 | -1.466607000 | 21.045911000 |
| 1 | 9.934985000 | -0.908222000 | 17.535890000 |
| 1 | 10.935189000 | -2.892526000 | 18.646178000 |
| 7 | 9.498905000 | 0.248866000 | 11.627212000 |
| 6 | 8.177382000 | -0.147666000 | 12.125412000 |
| 6 | 7.140677000 | 0.864372000 | 11.627417000 |
| 8 | 7.351461000 | 2.077454000 | 11.678109000 |
| 6 | 8.131109000 | -0.113911000 | 13.674338000 |
| 6 | 9.344494000 | -0.651815000 | 14.361201000 |
| 7 | 10.198051000 | 0.206226000 | 15.017626000 |
| 6 | 9.880149000 | -1.909413000 | 14.569105000 |
| 6 | 11.168869000 | -0.525911000 | 15.602809000 |
| 7 | 11.015517000 | -1.823083000 | 15.350473000 |
| 1 | 9.499432000 | 0.221325000 | 10.604793000 |
| 1 | 7.898712000 | -1.155185000 | 11.777739000 |

| | | | |
|---|---|---|---|
| 1 | 7.988213000 | 0.920483000 | 13.991735000 |
| 1 | 7.241960000 | -0.655279000 | 14.007626000 |
| 1 | 9.502052000 | -2.862816000 | 14.231290000 |
| 1 | 11.933390000 | -0.089776000 | 16.224949000 |
| 1 | 10.073538000 | 1.231744000 | 15.084802000 |
| 7 | 5.982535000 | 0.338681000 | 11.157003000 |
| 6 | 4.826171000 | 1.169784000 | 10.886266000 |
| 6 | 3.562677000 | 0.361988000 | 11.151661000 |
| 8 | 3.613077000 | -0.793305000 | 11.565117000 |
| 1 | 5.769073000 | -0.643097000 | 11.306138000 |
| 1 | 4.836699000 | 2.026347000 | 11.568208000 |
| 7 | 2.394053000 | 1.033689000 | 10.927223000 |
| 6 | 1.185764000 | 0.660055000 | 11.653623000 |
| 6 | 0.967787000 | 1.539805000 | 12.923153000 |
| 6 | 2.283925000 | 1.862273000 | 13.593015000 |
| 6 | 3.055411000 | 0.853497000 | 14.195984000 |
| 6 | 2.886566000 | 3.116389000 | 13.408680000 |
| 6 | 4.378431000 | 1.071984000 | 14.552670000 |
| 6 | 4.215959000 | 3.355962000 | 13.765939000 |
| 6 | 4.981680000 | 2.317625000 | 14.314428000 |
| 8 | 6.281370000 | 2.438506000 | 14.608863000 |
| 1 | 2.486160000 | 2.010381000 | 10.675155000 |
| 1 | 1.320500000 | -0.386275000 | 11.930679000 |
| 1 | 0.466263000 | 2.472256000 | 12.636882000 |
| 1 | 0.280085000 | 1.004076000 | 13.590533000 |
| 1 | 2.632035000 | -0.139916000 | 14.320747000 |
| 1 | 2.318564000 | 3.914655000 | 12.934010000 |
| 1 | 4.986569000 | 0.287851000 | 14.987079000 |
| 1 | 4.678371000 | 4.318602000 | 13.577181000 |
| 1 | 6.613611000 | 3.374900000 | 14.534748000 |
| 6 | 4.278493000 | 7.946361000 | 12.837172000 |
| 6 | 5.280278000 | 7.202381000 | 13.728279000 |
| 6 | 6.501984000 | 8.084544000 | 14.009934000 |
| 6 | 4.618818000 | 6.735378000 | 15.030306000 |
| 1 | 4.757522000 | 8.313023000 | 11.922641000 |
| 1 | 3.862119000 | 8.815729000 | 13.360693000 |
| 1 | 5.637635000 | 6.315580000 | 13.185605000 |
| 1 | 7.029534000 | 8.337150000 | 13.081969000 |
| 1 | 6.199726000 | 9.025166000 | 14.487878000 |
| 1 | 7.208645000 | 7.581793000 | 14.674884000 |
| 1 | 5.279275000 | 6.081950000 | 15.607090000 |
| 1 | 4.346866000 | 7.595727000 | 15.655279000 |
| 1 | 3.697654000 | 6.173520000 | 14.830035000 |
| 6 | 0.169899000 | 1.372698000 | 18.650458000 |
| 7 | 1.031615000 | 1.721191000 | 17.514852000 |
| 6 | 2.304482000 | 2.188233000 | 17.687892000 |
| 7 | 3.036755000 | 1.802104000 | 18.738043000 |
| 7 | 2.807500000 | 3.041844000 | 16.805738000 |
| 1 | 0.677446000 | 0.673971000 | 19.314554000 |
| 1 | -0.126941000 | 2.259277000 | 19.224064000 |
| 1 | 0.539728000 | 2.197316000 | 16.768395000 |
| 1 | 2.740838000 | 0.993329000 | 19.280346000 |
| 1 | 4.054554000 | 1.927130000 | 18.639201000 |
| 1 | 3.789367000 | 3.382102000 | 16.904366000 |
| 1 | 2.357485000 | 3.176957000 | 15.908685000 |
| 26 | 7.194381000 | 0.477298000 | 18.357562000 |
| 6 | 8.242659000 | 5.186300000 | 15.507441000 |
| 6 | 7.995197000 | 4.755644000 | 16.943750000 |
| 6 | 7.626466000 | 3.279450000 | 17.106188000 |
| 6 | 6.139383000 | 3.067838000 | 17.411976000 |
| 8 | 7.490815000 | 4.775716000 | 14.582953000 |
| 8 | 9.190410000 | 6.020578000 | 15.311415000 |
| 8 | 5.341466000 | 3.977567000 | 17.122127000 |
| 8 | 5.767888000 | 1.937113000 | 17.904096000 |
| 16 | 8.788281000 | 2.400086000 | 18.232795000 |
| 1 | 8.864272000 | 5.015072000 | 17.548043000 |
| 1 | 7.161610000 | 5.364166000 | 17.310080000 |
| 1 | 7.811765000 | 2.729008000 | 16.183498000 |
| 8 | 7.754624000 | 0.273728000 | 16.837216000 |
| 8 | 12.290702000 | -3.045248000 | 17.563613000 |
| 1 | 12.838202000 | -2.251313000 | 17.724351000 |
| 1 | 11.888170000 | -2.858646000 | 16.679419000 |
| 8 | 8.524379000 | 5.258745000 | 12.196385000 |
| 1 | 8.093042000 | 5.086930000 | 13.084659000 |
| 1 | 7.829805000 | 5.123602000 | 11.525101000 |
| 1 | 14.242816000 | 5.812223000 | 20.145758000 |
| 1 | 16.960279000 | 6.212963000 | 14.404399000 |
| 1 | 8.030574000 | -3.304127000 | 22.488111000 |
| 1 | 7.086564000 | -5.427801000 | 20.763607000 |
| 1 | 5.788539000 | -1.756161000 | 23.747627000 |
| 1 | 0.843989000 | -2.690477000 | 21.919817000 |
| 1 | 0.436470000 | -2.299727000 | 17.396171000 |
| 1 | -0.724406000 | 0.883457000 | 18.258815000 |
| 8 | 8.978977000 | 3.305871000 | 19.513562000 |
| 1 | 3.433756000 | 7.299400000 | 12.560562000 |
| 1 | 0.315808000 | 0.725725000 | 10.993050000 |
| 1 | 4.855669000 | 1.568674000 | 9.864542000 |
| 1 | 10.180061000 | -0.430379000 | 11.964119000 |
| 1 | 6.322832000 | 1.248817000 | 7.435262000 |
| 1 | 11.995849000 | 1.470963000 | 6.217363000 |
| 1 | 17.924908000 | 2.428247000 | 9.442584000 |
| 1 | 19.140151000 | -0.208979000 | 13.605748000 |
| 1 | 17.490741000 | 5.445836000 | 9.583533000 |
| 1 | 13.007503000 | 6.657479000 | 8.076302000 |
| 1 | 6.307183000 | -2.338007000 | 15.719275000 |
| 8 | 9.939377000 | 2.930227000 | 15.004695000 |
| 1 | 10.732237000 | 3.307252000 | 15.450966000 |
| 1 | 10.104300000 | 3.060334000 | 14.047531000 |
| 8 | 13.207765000 | 2.382470000 | 17.787526000 |
| 1 | 13.111956000 | 2.737332000 | 16.878934000 |
| 1 | 13.187510000 | 1.405206000 | 17.745198000 |
| 8 | 10.897252000 | 6.558926000 | 17.324252000 |
| 1 | 10.385889000 | 7.141437000 | 17.907355000 |
| 1 | 10.269679000 | 6.343264000 | 16.581940000 |
| 8 | 13.688801000 | -0.451499000 | 17.759008000 |
| 1 | 14.386289000 | -0.439700000 | 18.435436000 |
| 1 | 14.161259000 | -0.370804000 | 16.895695000 |

**$^{5}$TS3$_{R}$:**

| | | | |
|---|---|---|---|
| 6 | 16.240318000 | 5.481865000 | 13.963625000 |
| 6 | 16.062994000 | 6.067636000 | 12.566690000 |
| 8 | 15.399582000 | 7.095916000 | 12.369622000 |
| 6 | 14.914221000 | 5.456843000 | 14.728106000 |
| 6 | 13.931209000 | 4.448621000 | 14.105246000 |
| 6 | 12.565237000 | 4.572507000 | 14.734980000 |
| 8 | 12.290672000 | 4.032633000 | 15.832932000 |
| 7 | 11.674696000 | 5.333147000 | 14.080982000 |
| 1 | 16.687248000 | 4.483219000 | 13.905120000 |
| 1 | 15.084536000 | 5.204379000 | 15.780331000 |
| 1 | 14.476873000 | 6.459698000 | 14.692031000 |
| 1 | 13.852516000 | 4.625410000 | 13.028858000 |
| 1 | 14.287737000 | 3.426752000 | 14.263915000 |
| 1 | 10.737722000 | 5.440683000 | 14.472535000 |
| 1 | 11.788053000 | 5.448454000 | 13.073467000 |
| 7 | 16.697402000 | 5.402901000 | 11.575448000 |
| 6 | 16.732471000 | 5.927383000 | 10.220004000 |
| 6 | 15.425185000 | 5.663259000 | 9.463512000 |
| 8 | 15.293231000 | 4.683446000 | 8.709833000 |
| 1 | 17.155997000 | 4.509251000 | 11.766979000 |
| 1 | 16.936122000 | 7.002053000 | 10.258291000 |
| 7 | 14.448613000 | 6.552279000 | 9.691538000 |
| 6 | 13.103001000 | 6.379976000 | 9.148966000 |
| 6 | 12.126216000 | 7.272450000 | 9.906202000 |
| 6 | 10.671219000 | 6.985880000 | 9.533368000 |
| 6 | 9.662774000 | 7.880678000 | 10.263090000 |
| 7 | 9.687740000 | 7.688878000 | 11.715790000 |
| 6 | 10.319456000 | 8.448299000 | 12.596996000 |
| 7 | 11.102153000 | 9.479863000 | 12.202341000 |
| 7 | 10.134370000 | 8.254986000 | 13.914101000 |
| 1 | 14.596154000 | 7.164685000 | 10.492788000 |

| | | | |
|---|---|---|---|
| 1 | 12.799203000 | 5.335810000 | 9.238911000 |
| 1 | 12.266974000 | 7.092496000 | 10.977968000 |
| 1 | 12.371585000 | 8.328417000 | 9.702759000 |
| 1 | 10.452738000 | 5.937032000 | 9.760226000 |
| 1 | 10.520009000 | 7.121271000 | 8.454854000 |
| 1 | 9.827212000 | 8.940281000 | 10.040789000 |
| 1 | 8.648422000 | 7.639982000 | 9.932782000 |
| 1 | 9.220687000 | 6.818487000 | 12.039627000 |
| 1 | 11.531175000 | 10.069659000 | 12.899416000 |
| 1 | 11.538889000 | 9.460167000 | 11.293722000 |
| 1 | 9.570096000 | 7.462333000 | 14.249544000 |
| 1 | 10.875258000 | 8.529592000 | 14.544712000 |
| 7 | 6.650952000 | 1.805234000 | 8.209072000 |
| 6 | 7.890057000 | 2.485696000 | 7.849361000 |
| 6 | 9.084532000 | 1.532128000 | 7.799924000 |
| 8 | 9.208946000 | 0.620838000 | 8.621517000 |
| 6 | 8.283024000 | 3.558249000 | 8.904874000 |
| 6 | 7.295181000 | 4.687365000 | 9.023914000 |
| 6 | 6.255359000 | 4.643971000 | 9.963417000 |
| 6 | 7.387118000 | 5.815836000 | 8.196491000 |
| 6 | 5.341798000 | 5.694665000 | 10.078655000 |
| 6 | 6.479511000 | 6.866592000 | 8.293909000 |
| 6 | 5.447865000 | 6.808345000 | 9.239022000 |
| 8 | 4.589846000 | 7.867911000 | 9.290036000 |
| 1 | 6.880308000 | 1.108812000 | 8.918701000 |
| 1 | 7.749892000 | 2.975780000 | 6.878552000 |
| 1 | 8.382138000 | 3.043533000 | 9.862885000 |
| 1 | 9.274055000 | 3.946072000 | 8.644090000 |
| 1 | 6.178410000 | 3.778174000 | 10.613016000 |
| 1 | 8.186510000 | 5.873087000 | 7.460946000 |
| 1 | 4.545850000 | 5.644453000 | 10.818145000 |
| 1 | 6.548087000 | 7.736382000 | 7.648522000 |
| 1 | 3.929711000 | 7.714557000 | 9.983769000 |
| 7 | 10.037390000 | 1.791402000 | 6.857860000 |
| 6 | 11.392356000 | 1.346492000 | 7.124761000 |
| 6 | 11.984503000 | 2.195097000 | 8.260853000 |
| 8 | 11.658863000 | 3.371041000 | 8.397497000 |
| 1 | 9.965053000 | 2.687868000 | 6.393252000 |
| 1 | 11.369369000 | 0.285098000 | 7.378448000 |
| 7 | 12.876003000 | 1.567510000 | 9.064765000 |
| 6 | 13.497709000 | 2.241717000 | 10.200457000 |
| 6 | 14.865238000 | 1.587604000 | 10.408253000 |
| 8 | 14.975176000 | 0.475451000 | 10.942900000 |
| 6 | 12.624372000 | 2.154287000 | 11.470018000 |
| 6 | 11.659260000 | 3.330860000 | 11.584096000 |
| 8 | 12.086405000 | 4.489300000 | 11.437510000 |
| 7 | 10.394047000 | 3.048608000 | 11.957400000 |
| 1 | 12.988570000 | 0.563695000 | 9.002597000 |
| 1 | 13.621910000 | 3.288199000 | 9.923013000 |
| 1 | 13.263211000 | 2.196271000 | 12.360171000 |
| 1 | 12.102229000 | 1.195813000 | 11.509168000 |
| 1 | 10.023492000 | 2.082868000 | 11.874705000 |
| 1 | 9.713024000 | 3.817533000 | 11.939121000 |
| 7 | 15.920835000 | 2.290910000 | 9.959914000 |
| 6 | 17.263248000 | 1.758081000 | 10.094778000 |
| 6 | 17.699243000 | 1.776255000 | 11.564670000 |
| 8 | 17.794298000 | 2.837592000 | 12.191642000 |
| 1 | 15.774599000 | 3.173511000 | 9.451077000 |
| 1 | 17.316893000 | 0.747981000 | 9.675337000 |
| 7 | 17.960420000 | 0.571060000 | 12.114127000 |
| 6 | 18.199981000 | 0.398901000 | 13.543368000 |
| 6 | 16.928583000 | -0.041896000 | 14.288060000 |
| 6 | 15.808906000 | 0.991096000 | 14.171544000 |
| 16 | 14.160549000 | 0.432337000 | 14.749101000 |
| 6 | 13.842101000 | -0.966512000 | 13.613786000 |
| 1 | 17.656932000 | -0.240339000 | 11.591095000 |
| 1 | 18.566103000 | 1.356939000 | 13.917650000 |
| 1 | 16.596599000 | -1.000186000 | 13.872104000 |
| 1 | 17.174525000 | -0.222974000 | 15.341110000 |
| 1 | 15.680687000 | 1.296186000 | 13.133509000 |
| 1 | 16.040532000 | 1.884061000 | 14.760368000 |
| 1 | 14.527780000 | -1.793492000 | 13.811087000 |
| 1 | 12.825915000 | -1.306684000 | 13.818793000 |
| 1 | 13.946960000 | -0.624847000 | 12.582458000 |
| 6 | 13.713534000 | 5.408880000 | 19.177837000 |
| 6 | 12.463396000 | 4.925360000 | 19.902327000 |
| 7 | 11.520753000 | 4.299748000 | 18.926579000 |
| 1 | 13.450234000 | 6.143481000 | 18.409900000 |
| 1 | 14.221181000 | 4.566500000 | 18.698559000 |
| 1 | 11.933766000 | 5.748159000 | 20.390737000 |
| 1 | 12.700031000 | 4.172990000 | 20.659034000 |
| 1 | 11.991965000 | 3.477068000 | 18.477822000 |
| 1 | 11.277412000 | 5.006292000 | 18.195864000 |
| 1 | 10.626866000 | 3.978113000 | 19.356081000 |
| 6 | 4.685930000 | -1.176646000 | 23.953433000 |
| 6 | 4.321744000 | -2.132376000 | 22.827466000 |
| 8 | 5.159072000 | -2.731237000 | 22.156545000 |
| 6 | 4.500063000 | 0.310764000 | 23.554160000 |
| 6 | 5.403642000 | 0.736626000 | 22.440347000 |
| 7 | 6.424749000 | 1.666151000 | 22.588035000 |
| 6 | 5.498995000 | 0.316733000 | 21.137407000 |
| 6 | 7.086407000 | 1.778987000 | 21.406164000 |
| 7 | 6.547285000 | 0.963184000 | 20.517198000 |
| 1 | 4.064961000 | -1.389780000 | 24.831836000 |
| 1 | 3.455693000 | 0.485390000 | 23.267278000 |
| 1 | 4.680860000 | 0.929123000 | 24.440252000 |
| 1 | 4.894645000 | -0.405478000 | 20.616361000 |
| 1 | 7.907402000 | 2.455348000 | 21.206693000 |
| 1 | 6.629414000 | 2.189283000 | 23.428007000 |
| 7 | 2.977613000 | -2.275436000 | 22.607202000 |
| 6 | 2.480116000 | -2.973449000 | 21.453549000 |
| 6 | 2.221516000 | -2.042743000 | 20.256139000 |
| 8 | 2.489309000 | -0.841034000 | 20.302926000 |
| 1 | 2.342613000 | -1.659589000 | 23.095466000 |
| 1 | 3.240780000 | -3.705583000 | 21.167846000 |
| 7 | 1.679784000 | -2.664564000 | 19.180941000 |
| 6 | 1.498215000 | -2.041050000 | 17.874805000 |
| 6 | 2.634126000 | -2.391557000 | 16.890365000 |
| 6 | 3.972879000 | -2.036692000 | 17.445616000 |
| 7 | 4.753119000 | -2.929345000 | 18.168855000 |
| 6 | 4.649632000 | -0.846480000 | 17.517780000 |
| 6 | 5.858818000 | -2.288246000 | 18.622401000 |
| 7 | 5.810016000 | -1.015237000 | 18.250008000 |
| 1 | 1.496321000 | -3.657295000 | 19.255163000 |
| 1 | 1.478424000 | -0.962812000 | 18.026202000 |
| 1 | 2.461558000 | -1.841979000 | 15.958893000 |
| 1 | 2.596520000 | -3.457646000 | 16.632870000 |
| 1 | 4.397800000 | 0.111475000 | 17.094390000 |
| 1 | 6.621553000 | -2.784230000 | 19.211378000 |
| 1 | 4.600395000 | -3.924127000 | 18.250384000 |
| 6 | 6.192598000 | -3.054705000 | 14.919942000 |
| 6 | 5.182554000 | -2.704723000 | 13.817757000 |
| 6 | 6.059891000 | -4.486688000 | 15.446184000 |
| 6 | 5.432364000 | -3.423747000 | 12.489592000 |
| 1 | 7.211190000 | -2.909210000 | 14.540998000 |
| 1 | 4.165754000 | -2.933786000 | 14.170630000 |
| 1 | 5.191827000 | -1.624400000 | 13.632385000 |
| 1 | 6.756295000 | -4.670815000 | 16.272198000 |
| 1 | 6.266403000 | -5.228189000 | 14.667354000 |
| 1 | 5.040725000 | -4.677829000 | 15.810274000 |
| 1 | 6.442883000 | -3.204094000 | 12.117157000 |
| 1 | 5.353765000 | -4.511805000 | 12.589062000 |
| 1 | 4.712407000 | -3.096381000 | 11.734464000 |
| 7 | 6.996415000 | -4.644969000 | 20.705216000 |
| 6 | 8.064278000 | -4.449625000 | 21.701260000 |
| 6 | 9.399560000 | -4.068004000 | 21.044117000 |
| 6 | 9.299688000 | -2.766160000 | 20.314305000 |
| 7 | 10.066479000 | -2.441898000 | 19.205857000 |
| 6 | 8.428750000 | -1.713997000 | 20.473978000 |
| 6 | 9.639866000 | -1.259293000 | 18.726833000 |

| | | | |
|---|---|---|---|
| 7 | 8.641541000 | -0.786211000 | 19.468526000 |
| 1 | 6.103823000 | -4.458824000 | 21.156598000 |
| 1 | 8.226547000 | -5.334380000 | 22.334434000 |
| 1 | 9.702805000 | -4.855134000 | 20.341428000 |
| 1 | 10.178468000 | -4.030761000 | 21.818450000 |
| 1 | 7.626193000 | -1.594285000 | 21.185739000 |
| 1 | 10.018783000 | -0.786078000 | 17.838186000 |
| 1 | 10.864775000 | -2.934154000 | 18.753787000 |
| 7 | 9.509790000 | 0.282711000 | 11.600145000 |
| 6 | 8.180364000 | -0.116265000 | 12.068986000 |
| 6 | 7.150179000 | 0.907428000 | 11.581105000 |
| 8 | 7.362370000 | 2.119470000 | 11.647467000 |
| 6 | 8.109888000 | -0.113506000 | 13.617700000 |
| 6 | 9.306256000 | -0.666100000 | 14.323705000 |
| 7 | 10.127189000 | 0.180476000 | 15.031859000 |
| 6 | 9.829148000 | -1.929186000 | 14.541129000 |
| 6 | 11.056076000 | -0.561451000 | 15.666328000 |
| 7 | 10.918475000 | -1.854665000 | 15.386201000 |
| 1 | 9.528992000 | 0.261420000 | 10.577865000 |
| 1 | 7.901185000 | -1.115341000 | 11.697811000 |
| 1 | 7.961980000 | 0.914775000 | 13.954632000 |
| 1 | 7.213817000 | -0.663414000 | 13.917646000 |
| 1 | 9.485109000 | -2.878620000 | 14.155845000 |
| 1 | 11.786183000 | -0.135711000 | 16.335792000 |
| 1 | 9.975378000 | 1.196703000 | 15.133507000 |
| 7 | 5.989653000 | 0.389134000 | 11.109739000 |
| 6 | 4.829285000 | 1.220040000 | 10.858088000 |
| 6 | 3.571876000 | 0.393296000 | 11.098204000 |
| 8 | 3.632234000 | -0.778856000 | 11.460962000 |
| 1 | 5.781104000 | -0.597227000 | 11.232994000 |
| 1 | 4.832145000 | 2.060450000 | 11.560424000 |
| 7 | 2.398344000 | 1.066686000 | 10.910618000 |
| 6 | 1.196743000 | 0.652879000 | 11.626983000 |
| 6 | 0.972623000 | 1.486499000 | 12.925798000 |
| 6 | 2.287300000 | 1.818331000 | 13.594782000 |
| 6 | 3.102428000 | 0.807173000 | 14.130743000 |
| 6 | 2.838729000 | 3.104133000 | 13.484090000 |
| 6 | 4.422854000 | 1.053175000 | 14.481970000 |
| 6 | 4.163158000 | 3.371891000 | 13.838010000 |
| 6 | 4.978406000 | 2.330454000 | 14.303445000 |
| 8 | 6.282538000 | 2.479903000 | 14.570949000 |
| 1 | 2.484588000 | 2.055610000 | 10.708942000 |
| 1 | 1.346070000 | -0.400378000 | 11.867894000 |
| 1 | 0.447440000 | 2.416101000 | 12.675532000 |
| 1 | 0.303333000 | 0.913647000 | 13.580968000 |
| 1 | 2.720745000 | -0.208427000 | 14.196540000 |
| 1 | 2.231947000 | 3.909373000 | 13.073577000 |
| 1 | 5.059998000 | 0.264620000 | 14.864346000 |
| 1 | 4.576675000 | 4.368181000 | 13.730523000 |
| 1 | 6.588652000 | 3.423916000 | 14.508487000 |
| 6 | 4.278980000 | 8.042392000 | 12.714684000 |
| 6 | 5.262728000 | 7.277704000 | 13.608585000 |
| 6 | 6.510556000 | 8.126488000 | 13.876908000 |
| 6 | 4.591128000 | 6.844760000 | 14.917976000 |
| 1 | 4.764629000 | 8.390315000 | 11.796461000 |
| 1 | 3.887640000 | 8.925853000 | 13.233914000 |
| 1 | 5.590005000 | 6.375487000 | 13.071729000 |
| 1 | 7.043911000 | 8.349874000 | 12.944949000 |
| 1 | 6.237028000 | 9.082465000 | 14.341469000 |
| 1 | 7.202070000 | 7.612466000 | 14.548907000 |
| 1 | 5.229030000 | 6.169882000 | 15.496172000 |
| 1 | 4.356669000 | 7.718985000 | 15.538663000 |
| 1 | 3.646675000 | 6.319583000 | 14.726791000 |
| 6 | 0.297271000 | 1.305653000 | 19.008806000 |
| 7 | 1.079474000 | 1.606321000 | 17.803302000 |
| 6 | 2.356152000 | 2.086843000 | 17.879064000 |
| 7 | 3.130076000 | 1.787686000 | 18.929201000 |
| 7 | 2.823946000 | 2.858265000 | 16.908330000 |
| 1 | 0.797956000 | 0.544950000 | 19.608132000 |
| 1 | 0.133436000 | 2.199529000 | 19.622578000 |
| 1 | 0.538953000 | 2.033848000 | 17.061677000 |
| 1 | 2.873869000 | 0.994256000 | 19.510405000 |
| 1 | 4.142189000 | 1.933898000 | 18.793328000 |
| 1 | 3.789505000 | 3.257759000 | 16.962383000 |
| 1 | 2.311399000 | 2.980804000 | 16.044103000 |
| 26 | 7.281422000 | 0.537662000 | 18.515158000 |
| 6 | 8.201764000 | 5.280322000 | 15.487802000 |
| 6 | 7.902619000 | 4.872727000 | 16.921619000 |
| 6 | 7.599117000 | 3.380026000 | 17.077792000 |
| 6 | 6.136665000 | 3.101282000 | 17.453348000 |
| 8 | 7.460537000 | 4.866454000 | 14.556291000 |
| 8 | 9.172943000 | 6.089656000 | 15.302589000 |
| 8 | 5.294023000 | 3.952936000 | 17.111907000 |
| 8 | 5.832585000 | 1.995385000 | 18.028891000 |
| 16 | 8.866261000 | 2.551952000 | 18.127624000 |
| 1 | 8.726242000 | 5.180583000 | 17.565706000 |
| 1 | 7.019188000 | 5.440817000 | 17.230656000 |
| 1 | 7.754613000 | 2.851846000 | 16.141202000 |
| 8 | 8.286436000 | 0.825241000 | 17.124712000 |
| 8 | 12.187985000 | -3.060848000 | 17.609103000 |
| 1 | 12.742517000 | -2.270077000 | 17.757045000 |
| 1 | 11.770061000 | -2.876183000 | 16.731150000 |
| 8 | 8.510695000 | 5.267505000 | 12.158041000 |
| 1 | 8.077396000 | 5.119457000 | 13.049490000 |
| 1 | 7.815467000 | 5.126148000 | 11.488908000 |
| 1 | 14.400963000 | 5.877257000 | 19.888413000 |
| 1 | 16.955533000 | 6.126086000 | 14.490509000 |
| 1 | 7.743281000 | -3.643050000 | 22.367011000 |
| 1 | 6.992382000 | -5.613654000 | 20.391405000 |
| 1 | 5.728375000 | -1.367375000 | 24.216417000 |
| 1 | 1.562829000 | -3.517874000 | 21.705932000 |
| 1 | 0.529583000 | -2.348685000 | 17.467678000 |
| 1 | -0.667927000 | 0.907663000 | 18.690093000 |
| 8 | 8.884616000 | 3.404325000 | 19.441520000 |
| 1 | 3.416354000 | 7.416873000 | 12.444582000 |
| 1 | 0.323636000 | 0.728976000 | 10.971802000 |
| 1 | 4.857886000 | 1.641920000 | 9.845423000 |
| 1 | 10.184892000 | -0.396943000 | 11.947894000 |
| 1 | 6.312257000 | 1.279096000 | 7.404551000 |
| 1 | 11.999843000 | 1.475747000 | 6.223997000 |
| 1 | 17.940723000 | 2.405102000 | 9.532386000 |
| 1 | 18.991050000 | -0.345256000 | 13.679594000 |
| 1 | 17.539375000 | 5.432490000 | 9.677020000 |
| 1 | 13.108947000 | 6.629464000 | 8.080819000 |
| 1 | 6.085122000 | -2.349876000 | 15.751444000 |
| 8 | 9.886684000 | 2.933346000 | 14.982620000 |
| 1 | 10.680121000 | 3.293881000 | 15.437851000 |
| 1 | 10.068780000 | 3.050906000 | 14.028147000 |
| 8 | 13.148475000 | 2.375279000 | 17.806700000 |
| 1 | 13.045169000 | 2.723377000 | 16.895863000 |
| 1 | 13.150668000 | 1.397384000 | 17.772516000 |
| 8 | 10.847442000 | 6.542550000 | 17.369759000 |
| 1 | 10.345039000 | 7.152679000 | 17.931660000 |
| 1 | 10.241574000 | 6.360514000 | 16.600781000 |
| 8 | 13.623705000 | -0.468852000 | 17.759265000 |
| 1 | 14.363988000 | -0.496471000 | 18.387658000 |
| 1 | 14.039631000 | -0.393860000 | 16.866589000 |

**$^5$Prod$_R$:**

| | | | |
|---|---|---|---|
| 6 | 15.968445000 | 5.211670000 | 14.295560000 |
| 6 | 15.972565000 | 5.775651000 | 12.880236000 |
| 8 | 15.430527000 | 6.857899000 | 12.611034000 |
| 6 | 14.571899000 | 5.305759000 | 14.917030000 |
| 6 | 13.562745000 | 4.434598000 | 14.153140000 |
| 6 | 12.178095000 | 4.554699000 | 14.739318000 |
| 8 | 11.909304000 | 4.123616000 | 15.887663000 |
| 7 | 11.257772000 | 5.171951000 | 13.988789000 |
| 1 | 16.333997000 | 4.178897000 | 14.294607000 |
| 1 | 14.603455000 | 4.993284000 | 15.965954000 |
| 1 | 14.246119000 | 6.351143000 | 14.889033000 |

| | | | |
|---|---|---|---|
| 1 | 13.546405000 | 4.704825000 | 13.094502000 |
| 1 | 13.856389000 | 3.384908000 | 14.224710000 |
| 1 | 10.306300000 | 5.254442000 | 14.351402000 |
| 1 | 11.401424000 | 5.219618000 | 12.977899000 |
| 7 | 16.604301000 | 5.018943000 | 11.955893000 |
| 6 | 16.763170000 | 5.483311000 | 10.586631000 |
| 6 | 15.474934000 | 5.310815000 | 9.771098000 |
| 8 | 15.296168000 | 4.325809000 | 9.035103000 |
| 1 | 16.969732000 | 4.097023000 | 12.208625000 |
| 1 | 17.066746000 | 6.534783000 | 10.597978000 |
| 7 | 14.567452000 | 6.283410000 | 9.935559000 |
| 6 | 13.232665000 | 6.219254000 | 9.346392000 |
| 6 | 12.284418000 | 7.118536000 | 10.133876000 |
| 6 | 10.827625000 | 6.952896000 | 9.701764000 |
| 6 | 9.858312000 | 7.882107000 | 10.440095000 |
| 7 | 9.775093000 | 7.604130000 | 11.878125000 |
| 6 | 10.390009000 | 8.269244000 | 12.846698000 |
| 7 | 11.330273000 | 9.199662000 | 12.571039000 |
| 7 | 10.028377000 | 8.082321000 | 14.128350000 |
| 1 | 14.745940000 | 6.917482000 | 10.713323000 |
| 1 | 12.874496000 | 5.188184000 | 9.365912000 |
| 1 | 12.381553000 | 6.864586000 | 11.195620000 |
| 1 | 12.599802000 | 8.167916000 | 10.004347000 |
| 1 | 10.528409000 | 5.912929000 | 9.867982000 |
| 1 | 10.725566000 | 7.152969000 | 8.627538000 |
| 1 | 10.124956000 | 8.935312000 | 10.301644000 |
| 1 | 8.848531000 | 7.749701000 | 10.040238000 |
| 1 | 9.212631000 | 6.768053000 | 12.125054000 |
| 1 | 11.726134000 | 9.738941000 | 13.326437000 |
| 1 | 11.868264000 | 9.135891000 | 11.720168000 |
| 1 | 9.334292000 | 7.359530000 | 14.334792000 |
| 1 | 10.732565000 | 8.146647000 | 14.856170000 |
| 7 | 6.514812000 | 1.661396000 | 8.292741000 |
| 6 | 7.788746000 | 2.284925000 | 7.947744000 |
| 6 | 8.941622000 | 1.284489000 | 7.901846000 |
| 8 | 9.066138000 | 0.410704000 | 8.764306000 |
| 6 | 8.221608000 | 3.328741000 | 9.014815000 |
| 6 | 7.359952000 | 4.561177000 | 9.038521000 |
| 6 | 6.230523000 | 4.647831000 | 9.863849000 |
| 6 | 7.661565000 | 5.658255000 | 8.217335000 |
| 6 | 5.424683000 | 5.789626000 | 9.865425000 |
| 6 | 6.865612000 | 6.799010000 | 8.203474000 |
| 6 | 5.734693000 | 6.865917000 | 9.027045000 |
| 8 | 4.989265000 | 8.006338000 | 8.967498000 |
| 1 | 6.702757000 | 0.955763000 | 9.005705000 |
| 1 | 7.677996000 | 2.787754000 | 6.980034000 |
| 1 | 8.195491000 | 2.825928000 | 9.983176000 |
| 1 | 9.264072000 | 3.605016000 | 8.820705000 |
| 1 | 5.997724000 | 3.814315000 | 10.518133000 |
| 1 | 8.537973000 | 5.614199000 | 7.574877000 |
| 1 | 4.552491000 | 5.841074000 | 10.512753000 |
| 1 | 7.093845000 | 7.642232000 | 7.559535000 |
| 1 | 4.237550000 | 7.932652000 | 9.575622000 |
| 7 | 9.863615000 | 1.466774000 | 6.913142000 |
| 6 | 11.219908000 | 1.008942000 | 7.156891000 |
| 6 | 11.843261000 | 1.884342000 | 8.255735000 |
| 8 | 11.561417000 | 3.077234000 | 8.342644000 |
| 1 | 9.795434000 | 2.341155000 | 6.407565000 |
| 1 | 11.191338000 | -0.044534000 | 7.441106000 |
| 7 | 12.699362000 | 1.256378000 | 9.094163000 |
| 6 | 13.287023000 | 1.942983000 | 10.238081000 |
| 6 | 14.593732000 | 1.216013000 | 10.565410000 |
| 8 | 14.586635000 | 0.073747000 | 11.047662000 |
| 6 | 12.345816000 | 1.924760000 | 11.458665000 |
| 6 | 11.418483000 | 3.125127000 | 11.558556000 |
| 8 | 11.862635000 | 4.274329000 | 11.394570000 |
| 7 | 10.157644000 | 2.876137000 | 11.974068000 |
| 1 | 12.770394000 | 0.246730000 | 9.080305000 |
| 1 | 13.485185000 | 2.971759000 | 9.937927000 |
| 1 | 12.937555000 | 1.954173000 | 12.379059000 |

| | | | |
|---|---|---|---|
| 1 | 11.787145000 | 0.987486000 | 11.484967000 |
| 1 | 9.766830000 | 1.919646000 | 11.933940000 |
| 1 | 9.496872000 | 3.661373000 | 11.965167000 |
| 7 | 15.720374000 | 1.901672000 | 10.303470000 |
| 6 | 17.021709000 | 1.327312000 | 10.598364000 |
| 6 | 17.258068000 | 1.326685000 | 12.113725000 |
| 8 | 17.433111000 | 2.381845000 | 12.736022000 |
| 1 | 15.662917000 | 2.802182000 | 9.807954000 |
| 1 | 17.095057000 | 0.318193000 | 10.180750000 |
| 7 | 17.228044000 | 0.111861000 | 12.701849000 |
| 6 | 17.198482000 | -0.061383000 | 14.151338000 |
| 6 | 15.764846000 | -0.266314000 | 14.669594000 |
| 6 | 14.882099000 | 0.940186000 | 14.360957000 |
| 16 | 13.100306000 | 0.719804000 | 14.732845000 |
| 6 | 12.696585000 | -0.735380000 | 13.700127000 |
| 1 | 16.859103000 | -0.644477000 | 12.138249000 |
| 1 | 17.651895000 | 0.832011000 | 14.585340000 |
| 1 | 15.345966000 | -1.163939000 | 14.199896000 |
| 1 | 15.788825000 | -0.452174000 | 15.750112000 |
| 1 | 14.963521000 | 1.212563000 | 13.308895000 |
| 1 | 15.197705000 | 1.803373000 | 14.954962000 |
| 1 | 12.981510000 | -1.664812000 | 14.195789000 |
| 1 | 11.612489000 | -0.732724000 | 13.575313000 |
| 1 | 13.189173000 | -0.641092000 | 12.729028000 |
| 6 | 13.977518000 | 5.897716000 | 18.916356000 |
| 6 | 12.674281000 | 5.726199000 | 19.684925000 |
| 7 | 11.653668000 | 5.054305000 | 18.831621000 |
| 1 | 13.820378000 | 6.491904000 | 18.009939000 |
| 1 | 14.390879000 | 4.925934000 | 18.625425000 |
| 1 | 12.259211000 | 6.689975000 | 19.991838000 |
| 1 | 12.810046000 | 5.114881000 | 20.580698000 |
| 1 | 11.964473000 | 4.106312000 | 18.532695000 |
| 1 | 11.443589000 | 5.623206000 | 17.982774000 |
| 1 | 10.721324000 | 4.909280000 | 19.296544000 |
| 6 | 4.742542000 | -1.120075000 | 24.236824000 |
| 6 | 4.807952000 | -1.894358000 | 22.929313000 |
| 8 | 5.865394000 | -2.252778000 | 22.417900000 |
| 6 | 4.783102000 | 0.412405000 | 23.986003000 |
| 6 | 5.962371000 | 0.798689000 | 23.160798000 |
| 7 | 7.268186000 | 0.753361000 | 23.633122000 |
| 6 | 6.079920000 | 1.033238000 | 21.817126000 |
| 6 | 8.110708000 | 0.939136000 | 22.590426000 |
| 7 | 7.416554000 | 1.110309000 | 21.476874000 |
| 1 | 3.839018000 | -1.372945000 | 24.802982000 |
| 1 | 3.877516000 | 0.724635000 | 23.456061000 |
| 1 | 4.777477000 | 0.928337000 | 24.953456000 |
| 1 | 5.311537000 | 1.121192000 | 21.067386000 |
| 1 | 9.186712000 | 0.922287000 | 22.663166000 |
| 1 | 7.547404000 | 0.600930000 | 24.592250000 |
| 7 | 3.595825000 | -2.146416000 | 22.347540000 |
| 6 | 3.505637000 | -2.790070000 | 21.069003000 |
| 6 | 3.285623000 | -1.822023000 | 19.895931000 |
| 8 | 3.247815000 | -0.599688000 | 20.043555000 |
| 1 | 2.760543000 | -1.750544000 | 22.753878000 |
| 1 | 4.455275000 | -3.312007000 | 20.913443000 |
| 7 | 3.123634000 | -2.462197000 | 18.713053000 |
| 6 | 3.081191000 | -1.828719000 | 17.403371000 |
| 6 | 4.318976000 | -2.180919000 | 16.567528000 |
| 6 | 5.577627000 | -1.633630000 | 17.163886000 |
| 7 | 6.782092000 | -2.323997000 | 17.145478000 |
| 6 | 5.861763000 | -0.451124000 | 17.803888000 |
| 6 | 7.722593000 | -1.578200000 | 17.762919000 |
| 7 | 7.195440000 | -0.434611000 | 18.175750000 |
| 1 | 3.190858000 | -3.472505000 | 18.723244000 |
| 1 | 3.015636000 | -0.752284000 | 17.558034000 |
| 1 | 4.165864000 | -1.810454000 | 15.546359000 |
| 1 | 4.397112000 | -3.271984000 | 16.473193000 |
| 1 | 5.220025000 | 0.379294000 | 18.045914000 |
| 1 | 8.746678000 | -1.890163000 | 17.873039000 |
| 1 | 6.922603000 | -3.278525000 | 16.844155000 |

| | | | |
|---|---|---|---|
| 6 | 6.253713000 | -3.441850000 | 13.736940000 |
| 6 | 4.951871000 | -3.009180000 | 13.048631000 |
| 6 | 6.267451000 | -4.903690000 | 14.192298000 |
| 6 | 4.794469000 | -3.551041000 | 11.626021000 |
| 1 | 7.093778000 | -3.268129000 | 13.049853000 |
| 1 | 4.093728000 | -3.331988000 | 13.655709000 |
| 1 | 4.894535000 | -1.914514000 | 13.016433000 |
| 1 | 7.216132000 | -5.167131000 | 14.675788000 |
| 1 | 6.129173000 | -5.591230000 | 13.351732000 |
| 1 | 5.457558000 | -5.100696000 | 14.907803000 |
| 1 | 5.638191000 | -3.239072000 | 10.995199000 |
| 1 | 4.762864000 | -4.645709000 | 11.607537000 |
| 1 | 3.876905000 | -3.171797000 | 11.168751000 |
| 7 | 6.962779000 | -4.169986000 | 19.735700000 |
| 6 | 7.896971000 | -4.609118000 | 20.792189000 |
| 6 | 9.347759000 | -4.258939000 | 20.439210000 |
| 6 | 9.585132000 | -2.786411000 | 20.299568000 |
| 7 | 10.641814000 | -2.269615000 | 19.570820000 |
| 6 | 8.876185000 | -1.691122000 | 20.737852000 |
| 6 | 10.539596000 | -0.922804000 | 19.575617000 |
| 7 | 9.473635000 | -0.527433000 | 20.264904000 |
| 1 | 6.758023000 | -3.182452000 | 19.879980000 |
| 1 | 7.826901000 | -5.698508000 | 20.871225000 |
| 1 | 9.608957000 | -4.751045000 | 19.494393000 |
| 1 | 10.010208000 | -4.679634000 | 21.208185000 |
| 1 | 7.970535000 | -1.661743000 | 21.327021000 |
| 1 | 11.247051000 | -0.281385000 | 19.077139000 |
| 1 | 11.338467000 | -2.762968000 | 18.965857000 |
| 7 | 9.196805000 | 0.096196000 | 11.783271000 |
| 6 | 7.846764000 | -0.221229000 | 12.254953000 |
| 6 | 6.877914000 | 0.849303000 | 11.731003000 |
| 8 | 7.128743000 | 2.051678000 | 11.832415000 |
| 6 | 7.751097000 | -0.165435000 | 13.800750000 |
| 6 | 8.874297000 | -0.763397000 | 14.593496000 |
| 7 | 9.603071000 | 0.054623000 | 15.434874000 |
| 6 | 9.375133000 | -2.034595000 | 14.823583000 |
| 6 | 10.476696000 | -0.711909000 | 16.122686000 |
| 7 | 10.372783000 | -1.990750000 | 15.780263000 |
| 1 | 9.217431000 | 0.059692000 | 10.760326000 |
| 1 | 7.514762000 | -1.211317000 | 11.905225000 |
| 1 | 7.669462000 | 0.883994000 | 14.088037000 |
| 1 | 6.801501000 | -0.629399000 | 14.088483000 |
| 1 | 9.074389000 | -2.968908000 | 14.371533000 |
| 1 | 11.183601000 | -0.309334000 | 16.828673000 |
| 1 | 9.535604000 | 1.085947000 | 15.455376000 |
| 7 | 5.728144000 | 0.384060000 | 11.183121000 |
| 6 | 4.618769000 | 1.268945000 | 10.886747000 |
| 6 | 3.312174000 | 0.534283000 | 11.159780000 |
| 8 | 3.300426000 | -0.623051000 | 11.571692000 |
| 1 | 5.476476000 | -0.594163000 | 11.281942000 |
| 1 | 4.680341000 | 2.136638000 | 11.551343000 |
| 7 | 2.180871000 | 1.266179000 | 10.945386000 |
| 6 | 0.948627000 | 0.922595000 | 11.645071000 |
| 6 | 0.772741000 | 1.737525000 | 12.962877000 |
| 6 | 2.100095000 | 1.967135000 | 13.650833000 |
| 6 | 2.802186000 | 0.917527000 | 14.266860000 |
| 6 | 2.774322000 | 3.189550000 | 13.489058000 |
| 6 | 4.120640000 | 1.070918000 | 14.681691000 |
| 6 | 4.101830000 | 3.358205000 | 13.883062000 |
| 6 | 4.792175000 | 2.284275000 | 14.470423000 |
| 8 | 6.077401000 | 2.369120000 | 14.847193000 |
| 1 | 2.312797000 | 2.238245000 | 10.694536000 |
| 1 | 1.019317000 | -0.143922000 | 11.861263000 |
| 1 | 0.312834000 | 2.705766000 | 12.733136000 |
| 1 | 0.061387000 | 1.196714000 | 13.599959000 |
| 1 | 2.323840000 | -0.053117000 | 14.371685000 |
| 1 | 2.257937000 | 4.020467000 | 13.012294000 |
| 1 | 4.665655000 | 0.263836000 | 15.157324000 |
| 1 | 4.614632000 | 4.301924000 | 13.733176000 |
| 1 | 6.464236000 | 3.257431000 | 14.596479000 |

| | | | |
|---|---|---|---|
| 6 | 4.391686000 | 8.300770000 | 12.296865000 |
| 6 | 5.217947000 | 7.469463000 | 13.284804000 |
| 6 | 6.453225000 | 8.253554000 | 13.741894000 |
| 6 | 4.361217000 | 7.041012000 | 14.481591000 |
| 1 | 4.998841000 | 8.624688000 | 11.445163000 |
| 1 | 3.997101000 | 9.201604000 | 12.782372000 |
| 1 | 5.572488000 | 6.564516000 | 12.771153000 |
| 1 | 7.111397000 | 8.481193000 | 12.895055000 |
| 1 | 6.162802000 | 9.205752000 | 14.203244000 |
| 1 | 7.031277000 | 7.689511000 | 14.479120000 |
| 1 | 4.929132000 | 6.396878000 | 15.159981000 |
| 1 | 4.020060000 | 7.915154000 | 15.049886000 |
| 1 | 3.469007000 | 6.489424000 | 14.159083000 |
| 6 | 0.667080000 | 0.734856000 | 18.185697000 |
| 7 | 1.587538000 | 1.435788000 | 17.290514000 |
| 6 | 2.554279000 | 2.274451000 | 17.731063000 |
| 7 | 3.085223000 | 2.125733000 | 18.947938000 |
| 7 | 2.970095000 | 3.273735000 | 16.952177000 |
| 1 | 1.218078000 | 0.129153000 | 18.906955000 |
| 1 | 0.016342000 | 1.432353000 | 18.725404000 |
| 1 | 1.273429000 | 1.607272000 | 16.343907000 |
| 1 | 3.046624000 | 1.213225000 | 19.398535000 |
| 1 | 3.895371000 | 2.732144000 | 19.143843000 |
| 1 | 3.845734000 | 3.735874000 | 17.220687000 |
| 1 | 2.628138000 | 3.373251000 | 16.005373000 |
| 26 | 8.114292000 | 1.008737000 | 19.474435000 |
| 6 | 7.714133000 | 5.239788000 | 15.406862000 |
| 6 | 7.086119000 | 5.049915000 | 16.777991000 |
| 6 | 7.310085000 | 3.650130000 | 17.366547000 |
| 6 | 6.241500000 | 3.257095000 | 18.398221000 |
| 8 | 7.118043000 | 4.742198000 | 14.405793000 |
| 8 | 8.785937000 | 5.912273000 | 15.312166000 |
| 8 | 5.178444000 | 3.926421000 | 18.435164000 |
| 8 | 6.443877000 | 2.221296000 | 19.092589000 |
| 16 | 9.052748000 | 3.600281000 | 17.967907000 |
| 1 | 7.464259000 | 5.805688000 | 17.472629000 |
| 1 | 6.007788000 | 5.194601000 | 16.696480000 |
| 1 | 7.284421000 | 2.883250000 | 16.589472000 |
| 8 | 9.315301000 | 2.047548000 | 18.229036000 |
| 8 | 12.377284000 | -2.951857000 | 17.622558000 |
| 1 | 12.846797000 | -2.095813000 | 17.704001000 |
| 1 | 11.753131000 | -2.829065000 | 16.869233000 |
| 8 | 8.414168000 | 5.224235000 | 12.138468000 |
| 1 | 7.901110000 | 5.024009000 | 12.974883000 |
| 1 | 7.779311000 | 5.134135000 | 11.403311000 |
| 1 | 14.717322000 | 6.408862000 | 19.538973000 |
| 1 | 16.675767000 | 5.807444000 | 14.885673000 |
| 1 | 7.640988000 | -4.193346000 | 21.777370000 |
| 1 | 6.077730000 | -4.656499000 | 19.872826000 |
| 1 | 5.607166000 | -1.426043000 | 24.831628000 |
| 1 | 2.703083000 | -3.537836000 | 21.074647000 |
| 1 | 2.174576000 | -2.144033000 | 16.874550000 |
| 1 | 0.051146000 | 0.068333000 | 17.579375000 |
| 8 | 9.042709000 | 4.346949000 | 19.302959000 |
| 1 | 3.529550000 | 7.731866000 | 11.921356000 |
| 1 | 0.089488000 | 1.078541000 | 10.986276000 |
| 1 | 4.678169000 | 1.642021000 | 9.856399000 |
| 1 | 9.834474000 | -0.615750000 | 12.135589000 |
| 1 | 6.159841000 | 1.152712000 | 7.483884000 |
| 1 | 11.803529000 | 1.106016000 | 6.236916000 |
| 1 | 17.783936000 | 1.956014000 | 10.133413000 |
| 1 | 17.820517000 | -0.922286000 | 14.414927000 |
| 1 | 17.546349000 | 4.894028000 | 10.107219000 |
| 1 | 13.279174000 | 6.533975000 | 8.296370000 |
| 1 | 6.442400000 | -2.793379000 | 14.599804000 |
| 8 | 9.583943000 | 2.771677000 | 15.011164000 |
| 1 | 10.314087000 | 3.240277000 | 15.469913000 |
| 1 | 9.828267000 | 2.793589000 | 14.063459000 |
| 8 | 12.085783000 | 2.334524000 | 17.971388000 |
| 1 | 12.170807000 | 2.708780000 | 17.065948000 |

| | | | |
|---|---|---|---|
| 1 | 11.132311000 | 2.083793000 | 18.056154000 |
| 8 | 10.658407000 | 6.934046000 | 16.953701000 |
| 1 | 10.170119000 | 7.536346000 | 17.537948000 |
| 1 | 9.944363000 | 6.446965000 | 16.463097000 |
| 8 | 13.351749000 | -0.233313000 | 17.783150000 |
| 1 | 13.063869000 | 0.650227000 | 18.096246000 |
| 1 | 13.402410000 | -0.089590000 | 16.814899000 |

**$^3Re_R$:**

| | | | |
|---|---|---|---|
| 6 | 15.632834000 | 7.556514000 | 14.179300000 |
| 6 | 15.293431000 | 8.255092000 | 12.869734000 |
| 8 | 14.496550000 | 9.201908000 | 12.820288000 |
| 6 | 14.382327000 | 7.366876000 | 15.037219000 |
| 6 | 13.391800000 | 6.381742000 | 14.402471000 |
| 6 | 12.127260000 | 6.278052000 | 15.231595000 |
| 8 | 12.181463000 | 5.968297000 | 16.438654000 |
| 7 | 10.978069000 | 6.588001000 | 14.604014000 |
| 1 | 16.137495000 | 6.604097000 | 13.980860000 |
| 1 | 14.652361000 | 7.014759000 | 16.037699000 |
| 1 | 13.895644000 | 8.340701000 | 15.152703000 |
| 1 | 13.151997000 | 6.663797000 | 13.373517000 |
| 1 | 13.840698000 | 5.379123000 | 14.363968000 |
| 1 | 10.116283000 | 6.290288000 | 15.057121000 |
| 1 | 10.970976000 | 6.516507000 | 13.582788000 |
| 7 | 15.935830000 | 7.784261000 | 11.776363000 |
| 6 | 15.769502000 | 8.431272000 | 10.485701000 |
| 6 | 14.459141000 | 8.018983000 | 9.805459000 |
| 8 | 14.433535000 | 7.118431000 | 8.947022000 |
| 1 | 16.503738000 | 6.937660000 | 11.843415000 |
| 1 | 15.801222000 | 9.515866000 | 10.626603000 |
| 7 | 13.363653000 | 8.673207000 | 10.215403000 |
| 6 | 12.032755000 | 8.329516000 | 9.720826000 |
| 6 | 10.940735000 | 8.796487000 | 10.678294000 |
| 6 | 9.575302000 | 8.264598000 | 10.232850000 |
| 6 | 8.392548000 | 8.762173000 | 11.066871000 |
| 7 | 8.405735000 | 8.278475000 | 12.450250000 |
| 6 | 8.800910000 | 8.968043000 | 13.507576000 |
| 7 | 9.432568000 | 10.157112000 | 13.370866000 |
| 7 | 8.498634000 | 8.537429000 | 14.747326000 |
| 1 | 13.475879000 | 9.254554000 | 11.044480000 |
| 1 | 11.965457000 | 7.249537000 | 9.586202000 |
| 1 | 11.169769000 | 8.421065000 | 11.681117000 |
| 1 | 10.930524000 | 9.899376000 | 10.710452000 |
| 1 | 9.604839000 | 7.171211000 | 10.262119000 |
| 1 | 9.385124000 | 8.554747000 | 9.191145000 |
| 1 | 8.343036000 | 9.856194000 | 11.081125000 |
| 1 | 7.458186000 | 8.405612000 | 10.625376000 |
| 1 | 8.062072000 | 7.310073000 | 12.586048000 |
| 1 | 9.686581000 | 10.671875000 | 14.200815000 |
| 1 | 9.979756000 | 10.335670000 | 12.542098000 |
| 1 | 8.101504000 | 7.595345000 | 14.832193000 |
| 1 | 9.162989000 | 8.742366000 | 15.484375000 |
| 7 | 6.499737000 | 3.112819000 | 7.682728000 |
| 6 | 7.611892000 | 4.031967000 | 7.495721000 |
| 6 | 8.954167000 | 3.310263000 | 7.349851000 |
| 8 | 9.211392000 | 2.281357000 | 7.976164000 |
| 6 | 7.783503000 | 4.949248000 | 8.739146000 |
| 6 | 6.553396000 | 5.740026000 | 9.094363000 |
| 6 | 5.556231000 | 5.186393000 | 9.911060000 |
| 6 | 6.367545000 | 7.047197000 | 8.627335000 |
| 6 | 4.424310000 | 5.920843000 | 10.266703000 |
| 6 | 5.240523000 | 7.792320000 | 8.969725000 |
| 6 | 4.264391000 | 7.230003000 | 9.798457000 |
| 8 | 3.186653000 | 8.005690000 | 10.117263000 |
| 1 | 6.772168000 | 2.494948000 | 8.443810000 |
| 1 | 7.414280000 | 4.651165000 | 6.611620000 |
| 1 | 8.061026000 | 4.300453000 | 9.572826000 |
| 1 | 8.633042000 | 5.616424000 | 8.561826000 |
| 1 | 5.676932000 | 4.165819000 | 10.259482000 |
| 1 | 7.122918000 | 7.494502000 | 7.985036000 |
| 1 | 3.665874000 | 5.480036000 | 10.907782000 |
| 1 | 5.099378000 | 8.805063000 | 8.606310000 |
| 1 | 2.574172000 | 7.490587000 | 10.664768000 |
| 7 | 9.869910000 | 3.906021000 | 6.525686000 |
| 6 | 11.271226000 | 3.645817000 | 6.788964000 |
| 6 | 11.678156000 | 4.348453000 | 8.091571000 |
| 8 | 11.190985000 | 5.431424000 | 8.400590000 |
| 1 | 9.659404000 | 4.857998000 | 6.252328000 |
| 1 | 11.420850000 | 2.566174000 | 6.844446000 |
| 7 | 12.603475000 | 3.701519000 | 8.840881000 |
| 6 | 13.136762000 | 4.260576000 | 10.075613000 |
| 6 | 14.595140000 | 3.808098000 | 10.167526000 |
| 8 | 14.892900000 | 2.660067000 | 10.532747000 |
| 6 | 12.343622000 | 3.814303000 | 11.319073000 |
| 6 | 11.125076000 | 4.693311000 | 11.586761000 |
| 8 | 11.298170000 | 5.904420000 | 11.833732000 |
| 7 | 9.936308000 | 4.089721000 | 11.587801000 |
| 1 | 12.864101000 | 2.752501000 | 8.607002000 |
| 1 | 13.087055000 | 5.345029000 | 9.982725000 |
| 1 | 12.987301000 | 3.920558000 | 12.199046000 |
| 1 | 12.076358000 | 2.758307000 | 11.232568000 |
| 1 | 9.824027000 | 3.092752000 | 11.352033000 |
| 1 | 9.101382000 | 4.638201000 | 11.812142000 |
| 7 | 15.511391000 | 4.730420000 | 9.826267000 |
| 6 | 16.926611000 | 4.421044000 | 9.905384000 |
| 6 | 17.347067000 | 4.295486000 | 11.374505000 |
| 8 | 17.321571000 | 5.270076000 | 12.133518000 |
| 1 | 15.206762000 | 5.642893000 | 9.456834000 |
| 1 | 17.151269000 | 3.507579000 | 9.344878000 |
| 7 | 17.718782000 | 3.056445000 | 11.767511000 |
| 6 | 17.938187000 | 2.710526000 | 13.166763000 |
| 6 | 16.719804000 | 2.002684000 | 13.785665000 |
| 6 | 15.472536000 | 2.883705000 | 13.762389000 |
| 16 | 13.926903000 | 2.055960000 | 14.300846000 |
| 6 | 13.740065000 | 0.754225000 | 13.029203000 |
| 1 | 17.490386000 | 2.301391000 | 11.133158000 |
| 1 | 18.160277000 | 3.643074000 | 13.689540000 |
| 1 | 16.528967000 | 1.077269000 | 13.229287000 |
| 1 | 16.963515000 | 1.709721000 | 14.814013000 |
| 1 | 15.288455000 | 3.249473000 | 12.753542000 |
| 1 | 15.589964000 | 3.748289000 | 14.423007000 |
| 1 | 14.482204000 | -0.034664000 | 13.166766000 |
| 1 | 12.746238000 | 0.323637000 | 13.165880000 |
| 1 | 13.845624000 | 1.207418000 | 12.040768000 |
| 6 | 12.238883000 | 6.529273000 | 19.683417000 |
| 6 | 11.338896000 | 5.388213000 | 20.139738000 |
| 7 | 10.574404000 | 4.838308000 | 18.979105000 |
| 1 | 11.644666000 | 7.338014000 | 19.248382000 |
| 1 | 12.944487000 | 6.195153000 | 18.918462000 |
| 1 | 10.610935000 | 5.725828000 | 20.882721000 |
| 1 | 11.910920000 | 4.562640000 | 20.572308000 |
| 1 | 11.221934000 | 4.363924000 | 18.309313000 |
| 1 | 10.061020000 | 5.611075000 | 18.484347000 |
| 1 | 9.902934000 | 4.099351000 | 19.244463000 |
| 6 | 5.537323000 | -2.359948000 | 23.960931000 |
| 6 | 4.915806000 | -3.118622000 | 22.793897000 |
| 8 | 5.558757000 | -3.844786000 | 22.037383000 |
| 6 | 5.408416000 | -0.820800000 | 23.857975000 |
| 6 | 6.182378000 | -0.249654000 | 22.711804000 |
| 7 | 7.330256000 | 0.513495000 | 22.882339000 |
| 6 | 6.047078000 | -0.385100000 | 21.351428000 |
| 6 | 7.833718000 | 0.812269000 | 21.655935000 |
| 7 | 7.083402000 | 0.268639000 | 20.716575000 |
| 1 | 5.059852000 | -2.697450000 | 24.889350000 |
| 1 | 4.353600000 | -0.531956000 | 23.774923000 |
| 1 | 5.762342000 | -0.383297000 | 24.798242000 |
| 1 | 5.283380000 | -0.909392000 | 20.799171000 |
| 1 | 8.726506000 | 1.392742000 | 21.481173000 |
| 1 | 7.707379000 | 0.820393000 | 23.768256000 |
| 7 | 3.565104000 | -2.969632000 | 22.664728000 |

| | | | |
|---|---|---|---|
| 6 | 2.826829000 | -3.649898000 | 21.633940000 |
| 6 | 2.801415000 | -2.894890000 | 20.293554000 |
| 8 | 3.048251000 | -1.689256000 | 20.222358000 |
| 1 | 3.080991000 | -2.297831000 | 23.241233000 |
| 1 | 3.273800000 | -4.638021000 | 21.491568000 |
| 7 | 2.440194000 | -3.668315000 | 19.243504000 |
| 6 | 2.396988000 | -3.215262000 | 17.857621000 |
| 6 | 3.671271000 | -3.585189000 | 17.074721000 |
| 6 | 4.894022000 | -3.002354000 | 17.697943000 |
| 7 | 5.636375000 | -3.658233000 | 18.671004000 |
| 6 | 5.466492000 | -1.761043000 | 17.596955000 |
| 6 | 6.613988000 | -2.835734000 | 19.113258000 |
| 7 | 6.523534000 | -1.669213000 | 18.483627000 |
| 1 | 2.281093000 | -4.652596000 | 19.416640000 |
| 1 | 2.290689000 | -2.131433000 | 17.878006000 |
| 1 | 3.566877000 | -3.211440000 | 16.052704000 |
| 1 | 3.758733000 | -4.676837000 | 17.000417000 |
| 1 | 5.219359000 | -0.937993000 | 16.946528000 |
| 1 | 7.353165000 | -3.150598000 | 19.837035000 |
| 1 | 5.482016000 | -4.592477000 | 19.020766000 |
| 6 | 6.444676000 | -2.532357000 | 13.175121000 |
| 6 | 5.204866000 | -1.746547000 | 13.626578000 |
| 6 | 6.869047000 | -3.624551000 | 14.161454000 |
| 6 | 3.901676000 | -2.537496000 | 13.491108000 |
| 1 | 6.253328000 | -2.971185000 | 12.184940000 |
| 1 | 5.355542000 | -1.423388000 | 14.663132000 |
| 1 | 5.107215000 | -0.836021000 | 13.029173000 |
| 1 | 7.753986000 | -4.165517000 | 13.807087000 |
| 1 | 6.073850000 | -4.364011000 | 14.314480000 |
| 1 | 7.111954000 | -3.180491000 | 15.133615000 |
| 1 | 3.676040000 | -2.717110000 | 12.436903000 |
| 1 | 3.953066000 | -3.510301000 | 13.995811000 |
| 1 | 3.057129000 | -1.976495000 | 13.911035000 |
| 7 | 8.472572000 | -4.722223000 | 21.290750000 |
| 6 | 9.671923000 | -4.329419000 | 22.040313000 |
| 6 | 10.817286000 | -3.896906000 | 21.105194000 |
| 6 | 10.419675000 | -2.730943000 | 20.256506000 |
| 7 | 11.023581000 | -2.361807000 | 19.063472000 |
| 6 | 9.336422000 | -1.896643000 | 20.375372000 |
| 6 | 10.305575000 | -1.369360000 | 18.501748000 |
| 7 | 9.263950000 | -1.072700000 | 19.269154000 |
| 1 | 7.638504000 | -4.668476000 | 21.872902000 |
| 1 | 10.051946000 | -5.117450000 | 22.708148000 |
| 1 | 11.095242000 | -4.739855000 | 20.457699000 |
| 1 | 11.704579000 | -3.671788000 | 21.712901000 |
| 1 | 8.574698000 | -1.880237000 | 21.135498000 |
| 1 | 10.535950000 | -0.926547000 | 17.550108000 |
| 1 | 11.877245000 | -2.675860000 | 18.601248000 |
| 7 | 9.676358000 | 1.204566000 | 10.876951000 |
| 6 | 8.465049000 | 0.550612000 | 11.400907000 |
| 6 | 7.253124000 | 1.188080000 | 10.721981000 |
| 8 | 7.134344000 | 2.413937000 | 10.661748000 |
| 6 | 8.354385000 | 0.781998000 | 12.916080000 |
| 6 | 9.502988000 | 0.239835000 | 13.711178000 |
| 7 | 10.143325000 | 1.040508000 | 14.628284000 |
| 6 | 10.083727000 | -1.006804000 | 13.867599000 |
| 6 | 11.055692000 | 0.288277000 | 15.288095000 |
| 7 | 11.053686000 | -0.965391000 | 14.854926000 |
| 1 | 9.632521000 | 1.300173000 | 9.860511000 |
| 1 | 8.456860000 | -0.531901000 | 11.199141000 |
| 1 | 8.287020000 | 1.861910000 | 13.080285000 |
| 1 | 7.417565000 | 0.354755000 | 13.288355000 |
| 1 | 9.844499000 | -1.928856000 | 13.356638000 |
| 1 | 11.666419000 | 0.688489000 | 16.082472000 |
| 1 | 10.105983000 | 2.065376000 | 14.699781000 |
| 7 | 6.360076000 | 0.324648000 | 10.178845000 |
| 6 | 5.160820000 | 0.765842000 | 9.499785000 |
| 6 | 3.915376000 | 0.133814000 | 10.118484000 |
| 8 | 3.993158000 | -0.854982000 | 10.836819000 |
| 1 | 6.393973000 | -0.655244000 | 10.432098000 |

| | | | |
|---|---|---|---|
| 1 | 5.118609000 | 1.853554000 | 9.558665000 |
| 7 | 2.747578000 | 0.736423000 | 9.759376000 |
| 6 | 1.515796000 | 0.530162000 | 10.508720000 |
| 6 | 1.272213000 | 1.676355000 | 11.519917000 |
| 6 | 2.428469000 | 1.839311000 | 12.475499000 |
| 6 | 2.481795000 | 1.125206000 | 13.679917000 |
| 6 | 3.547154000 | 2.606258000 | 12.124101000 |
| 6 | 3.607414000 | 1.164401000 | 14.496843000 |
| 6 | 4.694001000 | 2.636477000 | 12.914114000 |
| 6 | 4.730696000 | 1.908812000 | 14.110297000 |
| 8 | 5.818705000 | 1.880727000 | 14.910011000 |
| 1 | 2.823005000 | 1.602831000 | 9.240823000 |
| 1 | 1.627549000 | -0.422748000 | 11.029023000 |
| 1 | 1.104021000 | 2.607510000 | 10.962031000 |
| 1 | 0.341973000 | 1.462294000 | 12.060205000 |
| 1 | 1.629277000 | 0.516219000 | 13.973885000 |
| 1 | 3.534715000 | 3.175090000 | 11.197705000 |
| 1 | 3.649471000 | 0.611096000 | 15.429791000 |
| 1 | 5.565700000 | 3.200690000 | 12.602799000 |
| 1 | 6.412694000 | 2.650791000 | 14.725855000 |
| 6 | 2.854677000 | 6.522037000 | 13.439534000 |
| 6 | 4.151332000 | 6.033892000 | 14.092810000 |
| 6 | 5.035536000 | 7.209901000 | 14.521418000 |
| 6 | 3.842068000 | 5.108764000 | 15.275748000 |
| 1 | 3.059365000 | 7.229353000 | 12.628380000 |
| 1 | 2.223918000 | 7.042591000 | 14.171302000 |
| 1 | 4.715365000 | 5.448014000 | 13.355732000 |
| 1 | 5.344439000 | 7.809621000 | 13.656432000 |
| 1 | 4.498938000 | 7.871239000 | 15.213918000 |
| 1 | 5.936000000 | 6.848357000 | 15.025630000 |
| 1 | 4.755236000 | 4.690579000 | 15.708525000 |
| 1 | 3.313739000 | 5.664189000 | 16.064438000 |
| 1 | 3.210947000 | 4.272328000 | 14.952170000 |
| 6 | 0.701360000 | 0.188202000 | 18.523661000 |
| 7 | 1.344025000 | 1.462333000 | 18.246548000 |
| 6 | 2.678435000 | 1.656937000 | 18.148181000 |
| 7 | 3.539667000 | 0.710866000 | 18.520146000 |
| 7 | 3.127035000 | 2.826583000 | 17.687648000 |
| 1 | 0.806769000 | -0.506660000 | 17.682215000 |
| 1 | 1.112344000 | -0.277285000 | 19.423761000 |
| 1 | 0.757086000 | 2.253769000 | 18.030570000 |
| 1 | 3.262986000 | -0.100453000 | 19.068152000 |
| 1 | 4.552305000 | 0.880033000 | 18.379508000 |
| 1 | 4.153961000 | 3.014012000 | 17.647439000 |
| 1 | 2.542172000 | 3.383471000 | 17.081157000 |
| 26 | 7.732946000 | 0.005496000 | 18.615463000 |
| 6 | 7.905881000 | 4.695420000 | 15.547253000 |
| 6 | 8.376801000 | 3.999765000 | 16.815896000 |
| 6 | 8.000502000 | 2.556090000 | 17.072786000 |
| 6 | 6.566927000 | 2.373903000 | 17.552620000 |
| 8 | 7.356557000 | 4.077027000 | 14.599783000 |
| 8 | 8.189227000 | 5.941296000 | 15.512549000 |
| 8 | 5.784059000 | 3.337060000 | 17.547662000 |
| 8 | 6.250784000 | 1.187828000 | 17.978464000 |
| 16 | 9.118196000 | 1.832318000 | 18.362182000 |
| 1 | 9.466138000 | 4.073315000 | 16.756465000 |
| 1 | 8.052552000 | 4.614525000 | 17.657728000 |
| 1 | 8.124611000 | 1.954904000 | 16.169191000 |
| 8 | 8.322623000 | -0.547068000 | 16.639622000 |
| 8 | 12.811961000 | -1.880788000 | 16.964041000 |
| 1 | 13.100760000 | -0.956194000 | 17.067224000 |
| 1 | 12.274653000 | -1.861897000 | 16.137809000 |
| 8 | 7.558306000 | 5.634549000 | 12.401233000 |
| 1 | 7.369960000 | 5.096273000 | 13.218945000 |
| 1 | 6.818319000 | 5.504089000 | 11.782112000 |
| 1 | 12.798731000 | 6.921557000 | 20.537376000 |
| 1 | 16.350443000 | 8.196342000 | 14.708993000 |
| 1 | 9.403500000 | -3.482889000 | 22.683861000 |
| 1 | 8.559861000 | -5.684228000 | 20.969858000 |
| 1 | 6.591283000 | -2.643629000 | 24.001508000 |

| | | | |
|---|---|---|---|
| 1 | 1.793784000 | -3.791974000 | 21.969986000 |
| 1 | 1.511192000 | -3.642313000 | 17.376299000 |
| 1 | -0.361202000 | 0.372408000 | 18.688799000 |
| 8 | 7.411819000 | -0.521598000 | 15.767599000 |
| 1 | 2.274755000 | 5.681226000 | 13.037584000 |
| 1 | 0.672856000 | 0.449499000 | 9.814052000 |
| 1 | 5.204388000 | 0.495122000 | 8.435677000 |
| 1 | 10.483086000 | 0.632174000 | 11.116427000 |
| 1 | 6.406648000 | 2.528011000 | 6.852624000 |
| 1 | 11.876566000 | 4.040925000 | 5.967399000 |
| 1 | 17.481264000 | 5.249883000 | 9.459920000 |
| 1 | 18.818315000 | 2.063299000 | 13.237153000 |
| 1 | 16.591771000 | 8.132976000 | 9.833199000 |
| 1 | 11.887321000 | 8.782701000 | 8.731574000 |
| 1 | 7.283742000 | -1.836353000 | 13.054851000 |
| 8 | 10.958580000 | 3.601888000 | 14.852266000 |
| 1 | 11.386001000 | 3.542120000 | 15.736311000 |
| 1 | 11.686182000 | 3.408672000 | 14.236794000 |
| 8 | 12.327471000 | 3.518116000 | 17.277900000 |
| 1 | 12.808088000 | 4.318029000 | 16.969727000 |
| 1 | 12.923508000 | 2.747527000 | 17.414482000 |
| 8 | 9.176559000 | 6.965180000 | 17.779032000 |
| 1 | 8.393613000 | 7.241501000 | 18.281270000 |
| 1 | 8.805704000 | 6.577583000 | 16.933597000 |
| 8 | 13.728523000 | 1.052767000 | 17.291277000 |
| 1 | 14.539000000 | 0.890239000 | 17.801111000 |
| 1 | 14.020585000 | 1.179155000 | 16.356025000 |

**$^7$Re$_R$:**

| | | | |
|---|---|---|---|
| 6 | 15.496215000 | 7.608693000 | 13.768221000 |
| 6 | 15.113711000 | 8.205413000 | 12.420282000 |
| 8 | 14.288081000 | 9.124105000 | 12.323269000 |
| 6 | 14.260228000 | 7.391387000 | 14.642423000 |
| 6 | 13.327061000 | 6.322397000 | 14.055586000 |
| 6 | 12.076103000 | 6.184844000 | 14.898873000 |
| 8 | 12.150364000 | 5.893772000 | 16.109754000 |
| 7 | 10.906604000 | 6.427855000 | 14.286179000 |
| 1 | 16.058699000 | 6.679004000 | 13.626262000 |
| 1 | 14.553278000 | 7.100790000 | 15.655993000 |
| 1 | 13.720245000 | 8.341154000 | 14.716060000 |
| 1 | 13.069071000 | 6.553957000 | 13.019000000 |
| 1 | 13.832674000 | 5.348403000 | 14.058667000 |
| 1 | 10.036331000 | 6.273038000 | 14.797050000 |
| 1 | 10.875554000 | 6.472553000 | 13.265710000 |
| 7 | 15.747817000 | 7.675637000 | 11.349481000 |
| 6 | 15.552772000 | 8.230743000 | 10.020096000 |
| 6 | 14.232761000 | 7.763160000 | 9.394608000 |
| 8 | 14.190881000 | 6.789846000 | 8.621473000 |
| 1 | 16.355990000 | 6.862496000 | 11.467187000 |
| 1 | 15.577873000 | 9.322977000 | 10.083074000 |
| 7 | 13.148838000 | 8.459940000 | 9.762793000 |
| 6 | 11.798111000 | 8.070487000 | 9.364536000 |
| 6 | 10.773659000 | 8.637269000 | 10.343723000 |
| 6 | 9.372197000 | 8.083135000 | 10.079887000 |
| 6 | 8.281255000 | 8.682819000 | 10.972431000 |
| 7 | 8.439027000 | 8.349677000 | 12.391744000 |
| 6 | 8.952449000 | 9.137156000 | 13.325873000 |
| 7 | 9.575639000 | 10.292755000 | 13.002368000 |
| 7 | 8.786455000 | 8.834926000 | 14.625082000 |
| 1 | 13.282765000 | 9.092784000 | 10.550054000 |
| 1 | 11.727719000 | 6.982114000 | 9.338721000 |
| 1 | 11.088075000 | 8.371647000 | 11.359179000 |
| 1 | 10.768904000 | 9.737643000 | 10.260744000 |
| 1 | 9.395801000 | 6.997531000 | 10.216680000 |
| 1 | 9.083275000 | 8.271009000 | 9.037802000 |
| 1 | 8.236266000 | 9.772650000 | 10.872300000 |
| 1 | 7.305746000 | 8.292739000 | 10.668692000 |
| 1 | 8.133200000 | 7.393774000 | 12.655502000 |
| 1 | 9.898170000 | 10.901937000 | 13.739202000 |
| 1 | 10.019175000 | 10.388036000 | 12.101432000 |
| 1 | 8.348409000 | 7.935452000 | 14.855131000 |
| 1 | 9.509356000 | 9.106970000 | 15.280043000 |
| 7 | 6.293151000 | 2.512959000 | 8.305272000 |
| 6 | 7.385709000 | 3.418073000 | 7.967220000 |
| 6 | 8.710512000 | 2.682941000 | 7.774267000 |
| 8 | 9.038357000 | 1.757768000 | 8.522140000 |
| 6 | 7.667376000 | 4.435093000 | 9.110245000 |
| 6 | 6.511014000 | 5.351631000 | 9.402071000 |
| 6 | 5.577175000 | 5.037972000 | 10.397644000 |
| 6 | 6.328498000 | 6.540239000 | 8.681310000 |
| 6 | 4.499575000 | 5.882372000 | 10.672695000 |
| 6 | 5.257152000 | 7.390472000 | 8.939208000 |
| 6 | 4.333465000 | 7.061668000 | 9.939241000 |
| 8 | 3.305316000 | 7.934078000 | 10.149211000 |
| 1 | 6.675012000 | 1.778596000 | 8.902482000 |
| 1 | 7.113803000 | 3.970217000 | 7.059908000 |
| 1 | 7.923658000 | 3.856863000 | 10.001240000 |
| 1 | 8.555386000 | 5.015995000 | 8.837539000 |
| 1 | 5.710296000 | 4.122839000 | 10.965360000 |
| 1 | 7.040710000 | 6.806040000 | 7.903378000 |
| 1 | 3.791797000 | 5.622965000 | 11.455965000 |
| 1 | 5.114472000 | 8.308214000 | 8.377900000 |
| 1 | 2.732395000 | 7.592265000 | 10.852953000 |
| 7 | 9.539917000 | 3.156235000 | 6.799880000 |
| 6 | 10.965213000 | 2.941291000 | 6.966920000 |
| 6 | 11.462929000 | 3.805145000 | 8.136140000 |
| 8 | 10.958171000 | 4.900769000 | 8.363247000 |
| 1 | 9.287342000 | 4.053812000 | 6.405632000 |
| 1 | 11.142620000 | 1.878164000 | 7.139249000 |
| 7 | 12.474767000 | 3.281322000 | 8.867786000 |
| 6 | 13.061502000 | 3.993066000 | 9.996983000 |
| 6 | 14.536464000 | 3.593028000 | 10.057288000 |
| 8 | 14.883902000 | 2.477873000 | 10.475358000 |
| 6 | 12.354523000 | 3.661293000 | 11.324278000 |
| 6 | 11.182399000 | 4.582231000 | 11.633968000 |
| 8 | 11.326352000 | 5.814698000 | 11.550495000 |
| 7 | 10.073268000 | 3.998964000 | 12.127613000 |
| 1 | 12.768163000 | 2.325201000 | 8.714422000 |
| 1 | 12.965952000 | 5.058439000 | 9.788967000 |
| 1 | 13.055281000 | 3.805260000 | 12.153958000 |
| 1 | 12.058738000 | 2.610708000 | 11.345006000 |
| 1 | 9.903562000 | 2.989427000 | 11.978589000 |
| 1 | 9.242700000 | 4.587957000 | 12.265535000 |
| 7 | 15.409129000 | 4.523345000 | 9.633706000 |
| 6 | 16.836590000 | 4.267169000 | 9.689950000 |
| 6 | 17.301743000 | 4.250588000 | 11.151635000 |
| 8 | 17.277129000 | 5.275155000 | 11.842913000 |
| 1 | 15.059371000 | 5.396581000 | 9.213456000 |
| 1 | 17.078116000 | 3.327978000 | 9.181590000 |
| 7 | 17.708007000 | 3.048937000 | 11.616491000 |
| 6 | 17.980752000 | 2.799673000 | 13.027224000 |
| 6 | 16.787193000 | 2.134962000 | 13.735441000 |
| 6 | 15.556171000 | 3.038035000 | 13.746850000 |
| 16 | 14.011913000 | 2.259935000 | 14.360237000 |
| 6 | 13.797671000 | 0.894765000 | 13.157641000 |
| 1 | 17.475701000 | 2.248652000 | 11.041756000 |
| 1 | 18.219877000 | 3.765208000 | 13.477803000 |
| 1 | 16.552387000 | 1.195136000 | 13.222161000 |
| 1 | 17.078882000 | 1.873494000 | 14.759770000 |
| 1 | 15.345516000 | 3.398617000 | 12.740391000 |
| 1 | 15.718910000 | 3.910774000 | 14.387141000 |
| 1 | 14.428632000 | 0.040977000 | 13.411368000 |
| 1 | 12.753175000 | 0.585779000 | 13.216747000 |
| 1 | 14.030320000 | 1.262938000 | 12.155544000 |
| 6 | 12.951387000 | 6.586184000 | 19.288414000 |
| 6 | 11.864622000 | 5.748091000 | 19.949074000 |
| 7 | 10.893985000 | 5.261892000 | 18.924446000 |
| 1 | 12.516562000 | 7.444627000 | 18.767995000 |
| 1 | 13.504221000 | 5.999316000 | 18.549177000 |
| 1 | 11.301240000 | 6.327999000 | 20.684974000 |

| | | | |
|---|---|---|---|
| 1 | 12.274560000 | 4.868643000 | 20.453239000 |
| 1 | 11.370844000 | 4.637009000 | 18.238671000 |
| 1 | 10.473672000 | 6.072717000 | 18.408142000 |
| 1 | 10.132930000 | 4.672002000 | 19.321222000 |
| 6 | 4.985498000 | -3.527376000 | 22.870132000 |
| 6 | 4.253339000 | -3.790354000 | 21.557997000 |
| 8 | 4.737089000 | -4.513267000 | 20.685611000 |
| 6 | 4.770307000 | -2.161874000 | 23.558107000 |
| 6 | 5.434533000 | -1.034511000 | 22.826734000 |
| 7 | 6.468159000 | -0.292509000 | 23.387263000 |
| 6 | 5.332628000 | -0.575759000 | 21.536912000 |
| 6 | 6.945798000 | 0.560991000 | 22.447444000 |
| 7 | 6.288259000 | 0.400125000 | 21.310672000 |
| 1 | 4.680015000 | -4.328235000 | 23.556908000 |
| 1 | 3.702641000 | -1.958155000 | 23.713434000 |
| 1 | 5.189470000 | -2.228372000 | 24.568395000 |
| 1 | 4.649928000 | -0.883266000 | 20.759515000 |
| 1 | 7.765166000 | 1.245557000 | 22.606925000 |
| 1 | 6.794489000 | -0.362193000 | 24.341359000 |
| 7 | 3.001211000 | -3.269272000 | 21.434078000 |
| 6 | 2.176918000 | -3.575343000 | 20.286846000 |
| 6 | 2.497606000 | -2.687757000 | 19.076698000 |
| 8 | 2.700691000 | -1.479252000 | 19.220845000 |
| 1 | 2.654374000 | -2.609443000 | 22.112434000 |
| 1 | 2.301694000 | -4.634003000 | 20.045064000 |
| 7 | 2.473485000 | -3.323259000 | 17.883819000 |
| 6 | 2.831118000 | -2.698099000 | 16.614920000 |
| 6 | 4.308378000 | -2.936033000 | 16.252832000 |
| 6 | 5.233967000 | -2.377442000 | 17.281189000 |
| 7 | 5.702803000 | -3.117254000 | 18.356874000 |
| 6 | 5.718407000 | -1.109893000 | 17.489345000 |
| 6 | 6.427885000 | -2.302125000 | 19.158754000 |
| 7 | 6.457009000 | -1.075212000 | 18.657256000 |
| 1 | 2.365616000 | -4.329036000 | 17.891068000 |
| 1 | 2.650984000 | -1.627892000 | 16.708264000 |
| 1 | 4.504098000 | -2.470596000 | 15.283681000 |
| 1 | 4.488057000 | -4.010090000 | 16.120390000 |
| 1 | 5.593204000 | -0.224311000 | 16.890850000 |
| 1 | 6.922801000 | -2.664639000 | 20.046283000 |
| 1 | 5.411806000 | -4.044472000 | 18.642005000 |
| 6 | 7.452708000 | -3.515657000 | 12.871601000 |
| 6 | 6.205446000 | -2.866101000 | 12.255100000 |
| 6 | 7.403295000 | -3.634071000 | 14.398408000 |
| 6 | 4.924019000 | -3.693345000 | 12.386256000 |
| 1 | 7.597139000 | -4.511184000 | 12.430846000 |
| 1 | 6.036905000 | -1.880700000 | 12.710188000 |
| 1 | 6.387486000 | -2.679625000 | 11.188271000 |
| 1 | 8.340980000 | -4.035883000 | 14.797466000 |
| 1 | 6.596278000 | -4.300714000 | 14.721567000 |
| 1 | 7.239878000 | -2.658588000 | 14.872397000 |
| 1 | 5.033360000 | -4.669059000 | 11.897988000 |
| 1 | 4.667235000 | -3.880619000 | 13.435682000 |
| 1 | 4.088760000 | -3.164665000 | 11.918973000 |
| 7 | 8.256504000 | -4.441663000 | 21.015857000 |
| 6 | 9.647484000 | -4.388076000 | 21.501925000 |
| 6 | 10.573953000 | -3.660989000 | 20.519569000 |
| 6 | 10.164299000 | -2.251177000 | 20.229545000 |
| 7 | 10.837844000 | -1.430828000 | 19.335214000 |
| 6 | 9.105566000 | -1.501566000 | 20.677945000 |
| 6 | 10.178698000 | -0.257520000 | 19.264351000 |
| 7 | 9.109480000 | -0.265261000 | 20.058169000 |
| 1 | 7.712489000 | -5.085022000 | 21.588424000 |
| 1 | 10.077254000 | -5.383830000 | 21.689250000 |
| 1 | 10.616987000 | -4.228698000 | 19.577766000 |
| 1 | 11.593271000 | -3.687550000 | 20.926872000 |
| 1 | 8.333376000 | -1.783463000 | 21.373589000 |
| 1 | 10.464965000 | 0.568805000 | 18.632581000 |
| 1 | 11.673956000 | -1.631888000 | 18.744168000 |
| 7 | 9.747561000 | 1.164463000 | 11.390613000 |
| 6 | 8.514294000 | 0.456006000 | 11.739501000 |
| 6 | 7.315617000 | 1.384811000 | 11.510749000 |
| 8 | 7.359201000 | 2.583921000 | 11.794565000 |
| 6 | 8.473607000 | 0.071902000 | 13.248403000 |
| 6 | 9.771237000 | -0.172250000 | 13.951657000 |
| 7 | 10.328279000 | 0.829456000 | 14.717689000 |
| 6 | 10.523591000 | -1.296404000 | 14.247032000 |
| 6 | 11.333613000 | 0.295643000 | 15.446471000 |
| 7 | 11.494286000 | -0.994303000 | 15.182829000 |
| 1 | 9.739739000 | 1.346375000 | 10.383813000 |
| 1 | 8.385374000 | -0.454019000 | 11.133828000 |
| 1 | 7.971639000 | 0.875444000 | 13.793862000 |
| 1 | 7.842753000 | -0.812482000 | 13.353511000 |
| 1 | 10.398202000 | -2.302787000 | 13.872635000 |
| 1 | 11.898126000 | 0.873549000 | 16.160291000 |
| 1 | 10.022186000 | 1.813957000 | 14.798751000 |
| 7 | 6.194783000 | 0.788500000 | 11.038139000 |
| 6 | 4.920772000 | 1.480252000 | 11.032934000 |
| 6 | 3.798846000 | 0.455405000 | 11.100690000 |
| 8 | 4.027802000 | -0.750380000 | 11.139644000 |
| 1 | 6.128722000 | -0.223146000 | 10.974442000 |
| 1 | 4.857162000 | 2.123847000 | 11.917726000 |
| 7 | 2.542290000 | 0.991361000 | 11.142835000 |
| 6 | 1.470090000 | 0.271120000 | 11.820592000 |
| 6 | 1.287971000 | 0.759595000 | 13.287605000 |
| 6 | 2.616347000 | 1.074132000 | 13.940611000 |
| 6 | 3.580928000 | 0.074250000 | 14.135432000 |
| 6 | 3.004206000 | 2.399762000 | 14.179575000 |
| 6 | 4.882719000 | 0.385979000 | 14.510858000 |
| 6 | 4.300029000 | 2.732140000 | 14.574688000 |
| 6 | 5.260229000 | 1.721055000 | 14.724758000 |
| 8 | 6.540432000 | 1.973599000 | 15.060170000 |
| 1 | 2.503381000 | 2.001909000 | 11.204290000 |
| 1 | 1.757255000 | -0.780942000 | 11.795889000 |
| 1 | 0.660997000 | 1.659929000 | 13.295133000 |
| 1 | 0.728564000 | -0.012798000 | 13.832083000 |
| 1 | 3.331603000 | -0.958348000 | 13.906705000 |
| 1 | 2.280952000 | 3.199848000 | 14.027372000 |
| 1 | 5.634781000 | -0.388446000 | 14.611407000 |
| 1 | 4.585826000 | 3.761292000 | 14.753222000 |
| 1 | 6.727028000 | 2.956423000 | 15.065515000 |
| 6 | 3.160931000 | 7.555273000 | 13.581915000 |
| 6 | 4.281114000 | 6.900562000 | 14.398277000 |
| 6 | 5.330818000 | 7.940868000 | 14.802102000 |
| 6 | 3.714151000 | 6.186881000 | 15.631706000 |
| 1 | 3.565156000 | 8.095216000 | 12.718906000 |
| 1 | 2.608250000 | 8.279931000 | 14.192534000 |
| 1 | 4.783989000 | 6.152357000 | 13.768468000 |
| 1 | 5.794289000 | 8.394415000 | 13.917696000 |
| 1 | 4.874649000 | 8.747923000 | 15.389828000 |
| 1 | 6.121274000 | 7.484880000 | 15.403424000 |
| 1 | 4.496742000 | 5.661530000 | 16.187595000 |
| 1 | 3.235551000 | 6.907855000 | 16.307423000 |
| 1 | 2.950335000 | 5.453126000 | 15.336116000 |
| 6 | 0.632553000 | 0.400732000 | 17.209601000 |
| 7 | 1.118370000 | 1.750322000 | 17.449438000 |
| 6 | 2.415562000 | 2.057934000 | 17.670964000 |
| 7 | 3.296353000 | 1.108045000 | 17.967876000 |
| 7 | 2.800502000 | 3.339100000 | 17.620627000 |
| 1 | 1.087952000 | -0.022516000 | 16.306741000 |
| 1 | 0.843565000 | -0.251101000 | 18.061864000 |
| 1 | 0.493081000 | 2.520561000 | 17.268717000 |
| 1 | 3.009877000 | 0.185260000 | 18.289802000 |
| 1 | 4.283750000 | 1.375942000 | 18.121696000 |
| 1 | 3.820943000 | 3.553434000 | 17.674645000 |
| 1 | 2.249128000 | 3.986806000 | 17.075912000 |
| 26 | 7.338363000 | 0.796326000 | 19.450996000 |
| 6 | 7.783542000 | 5.200093000 | 15.859054000 |
| 6 | 8.113190000 | 4.679912000 | 17.247473000 |
| 6 | 7.783928000 | 3.256153000 | 17.644878000 |
| 6 | 6.299511000 | 2.991944000 | 17.840882000 |

| | | | |
|---|---|---|---|
| 8 | 7.100872000 | 4.568897000 | 15.013496000 |
| 8 | 8.298123000 | 6.349501000 | 15.618078000 |
| 8 | 5.467180000 | 3.869713000 | 17.557736000 |
| 8 | 5.982654000 | 1.820333000 | 18.301852000 |
| 16 | 8.674394000 | 2.835908000 | 19.228309000 |
| 1 | 9.194618000 | 4.784711000 | 17.290213000 |
| 1 | 7.681304000 | 5.376640000 | 17.974124000 |
| 1 | 8.141154000 | 2.569794000 | 16.874380000 |
| 8 | 8.493768000 | 0.028705000 | 16.790579000 |
| 8 | 12.921973000 | -1.546866000 | 17.552979000 |
| 1 | 13.318316000 | -0.654727000 | 17.563681000 |
| 1 | 12.494878000 | -1.603376000 | 16.661385000 |
| 8 | 7.746339000 | 5.696344000 | 12.654995000 |
| 1 | 7.426402000 | 5.288541000 | 13.508931000 |
| 1 | 7.098190000 | 5.444313000 | 11.973619000 |
| 1 | 13.652446000 | 6.949835000 | 20.045353000 |
| 1 | 16.171194000 | 8.323311000 | 14.256236000 |
| 1 | 9.646627000 | -3.855222000 | 22.460641000 |
| 1 | 8.244165000 | -4.841681000 | 20.077388000 |
| 1 | 6.047766000 | -3.668787000 | 22.655739000 |
| 1 | 1.129913000 | -3.405061000 | 20.556751000 |
| 1 | 2.173891000 | -3.093357000 | 15.833468000 |
| 1 | -0.448749000 | 0.447114000 | 17.072369000 |
| 8 | 8.628275000 | -1.166392000 | 16.602190000 |
| 1 | 2.435884000 | 6.809732000 | 13.225624000 |
| 1 | 0.533534000 | 0.376965000 | 11.264580000 |
| 1 | 4.828493000 | 2.116905000 | 10.143666000 |
| 1 | 10.533510000 | 0.548217000 | 11.592823000 |
| 1 | 5.965622000 | 2.046467000 | 7.460809000 |
| 1 | 11.485895000 | 3.237916000 | 6.051618000 |
| 1 | 17.349313000 | 5.083885000 | 9.177144000 |
| 1 | 18.864710000 | 2.158809000 | 13.106408000 |
| 1 | 16.367009000 | 7.894228000 | 9.376087000 |
| 1 | 11.597934000 | 8.434560000 | 8.348743000 |
| 1 | 8.337900000 | -2.930578000 | 12.586840000 |
| 8 | 10.057547000 | 3.503431000 | 15.148447000 |
| 1 | 10.836254000 | 3.551607000 | 15.742653000 |
| 1 | 10.318428000 | 3.878343000 | 14.285453000 |
| 8 | 12.158703000 | 3.510182000 | 17.116305000 |
| 1 | 12.595759000 | 4.262457000 | 16.656099000 |
| 1 | 12.816000000 | 2.806720000 | 17.310634000 |
| 8 | 9.709887000 | 7.486903000 | 17.615237000 |
| 1 | 9.024204000 | 7.878539000 | 18.179449000 |
| 1 | 9.197217000 | 7.038241000 | 16.884954000 |
| 8 | 13.901803000 | 1.282259000 | 17.386208000 |
| 1 | 14.712740000 | 1.348390000 | 17.916430000 |
| 1 | 14.188074000 | 1.383701000 | 16.445416000 |

**$^{1}$Re$_{R}$:**

| | | | |
|---|---|---|---|
| 6 | 15.579887000 | 7.542125000 | 13.927873000 |
| 6 | 15.215696000 | 8.201455000 | 12.604373000 |
| 8 | 14.397175000 | 9.129527000 | 12.539146000 |
| 6 | 14.336925000 | 7.285614000 | 14.780922000 |
| 6 | 13.411251000 | 6.239516000 | 14.141250000 |
| 6 | 12.178186000 | 6.032987000 | 14.997193000 |
| 8 | 12.277296000 | 5.614113000 | 16.167055000 |
| 7 | 10.997929000 | 6.370102000 | 14.448661000 |
| 1 | 16.142274000 | 6.618547000 | 13.750274000 |
| 1 | 14.623132000 | 6.950067000 | 15.782515000 |
| 1 | 13.793104000 | 8.229639000 | 14.891609000 |
| 1 | 13.134104000 | 6.532370000 | 13.125591000 |
| 1 | 13.935003000 | 5.276942000 | 14.077943000 |
| 1 | 10.139078000 | 6.185451000 | 14.966339000 |
| 1 | 10.939526000 | 6.500114000 | 13.437596000 |
| 7 | 15.861939000 | 7.717596000 | 11.519241000 |
| 6 | 15.695711000 | 8.336865000 | 10.214629000 |
| 6 | 14.382897000 | 7.912831000 | 9.545262000 |
| 8 | 14.343915000 | 6.977256000 | 8.726679000 |
| 1 | 16.460196000 | 6.893326000 | 11.607250000 |
| 1 | 15.729689000 | 9.424636000 | 10.329745000 |

| | | | |
|---|---|---|---|
| 7 | 13.299851000 | 8.602215000 | 9.928659000 |
| 6 | 11.952885000 | 8.242235000 | 9.491874000 |
| 6 | 10.919196000 | 8.802186000 | 10.463706000 |
| 6 | 9.510640000 | 8.294485000 | 10.150907000 |
| 6 | 8.421588000 | 8.883992000 | 11.052916000 |
| 7 | 8.562903000 | 8.487930000 | 12.457196000 |
| 6 | 9.098809000 | 9.214722000 | 13.425488000 |
| 7 | 9.722168000 | 10.385943000 | 13.159563000 |
| 7 | 8.962189000 | 8.838519000 | 14.708880000 |
| 1 | 13.425526000 | 9.191295000 | 10.750610000 |
| 1 | 11.864681000 | 7.156120000 | 9.437390000 |
| 1 | 11.202864000 | 8.493804000 | 11.476074000 |
| 1 | 10.944636000 | 9.904442000 | 10.418515000 |
| 1 | 9.504130000 | 7.203764000 | 10.243881000 |
| 1 | 9.244619000 | 8.532126000 | 9.112950000 |
| 1 | 8.399687000 | 9.977510000 | 10.995317000 |
| 1 | 7.441175000 | 8.526206000 | 10.725763000 |
| 1 | 8.252209000 | 7.519227000 | 12.667257000 |
| 1 | 10.066154000 | 10.946437000 | 13.924756000 |
| 1 | 10.151621000 | 10.531118000 | 12.258553000 |
| 1 | 8.518834000 | 7.937141000 | 14.924851000 |
| 1 | 9.691339000 | 9.095647000 | 15.361593000 |
| 7 | 6.437492000 | 2.788435000 | 8.079494000 |
| 6 | 7.540669000 | 3.708915000 | 7.827347000 |
| 6 | 8.869896000 | 2.981669000 | 7.639666000 |
| 8 | 9.180887000 | 2.028297000 | 8.358675000 |
| 6 | 7.789964000 | 4.664904000 | 9.029189000 |
| 6 | 6.629520000 | 5.571786000 | 9.333865000 |
| 6 | 5.661510000 | 5.216962000 | 10.283178000 |
| 6 | 6.478940000 | 6.795420000 | 8.665919000 |
| 6 | 4.579613000 | 6.056607000 | 10.559754000 |
| 6 | 5.403882000 | 7.639636000 | 8.925850000 |
| 6 | 4.443479000 | 7.269295000 | 9.875425000 |
| 8 | 3.409609000 | 8.134571000 | 10.083137000 |
| 1 | 6.799903000 | 2.026249000 | 8.653348000 |
| 1 | 7.299130000 | 4.308134000 | 6.941523000 |
| 1 | 8.019868000 | 4.041775000 | 9.895944000 |
| 1 | 8.686345000 | 5.255658000 | 8.809097000 |
| 1 | 5.768460000 | 4.272956000 | 10.808578000 |
| 1 | 7.219527000 | 7.092413000 | 7.926644000 |
| 1 | 3.840564000 | 5.764558000 | 11.301989000 |
| 1 | 5.285026000 | 8.583704000 | 8.403979000 |
| 1 | 2.832770000 | 7.785776000 | 10.780281000 |
| 7 | 9.724107000 | 3.492609000 | 6.705776000 |
| 6 | 11.143716000 | 3.270467000 | 6.908773000 |
| 6 | 11.598309000 | 4.073379000 | 8.136864000 |
| 8 | 11.105603000 | 5.170062000 | 8.383930000 |
| 1 | 9.483305000 | 4.406958000 | 6.343903000 |
| 1 | 11.318813000 | 2.199940000 | 7.029948000 |
| 7 | 12.557709000 | 3.494186000 | 8.896362000 |
| 6 | 13.136542000 | 4.152659000 | 10.061406000 |
| 6 | 14.606857000 | 3.733362000 | 10.106647000 |
| 8 | 14.939752000 | 2.600341000 | 10.485771000 |
| 6 | 12.411754000 | 3.775140000 | 11.367895000 |
| 6 | 11.235182000 | 4.691740000 | 11.685906000 |
| 8 | 11.387293000 | 5.925476000 | 11.654287000 |
| 7 | 10.104242000 | 4.099499000 | 12.120936000 |
| 1 | 12.844439000 | 2.542350000 | 8.708253000 |
| 1 | 13.052651000 | 5.226563000 | 9.896932000 |
| 1 | 13.104060000 | 3.884397000 | 12.210523000 |
| 1 | 12.112330000 | 2.725459000 | 11.348349000 |
| 1 | 9.929454000 | 3.100187000 | 11.909728000 |
| 1 | 9.276810000 | 4.699328000 | 12.231114000 |
| 7 | 15.491749000 | 4.660560000 | 9.701455000 |
| 6 | 16.913631000 | 4.372474000 | 9.725304000 |
| 6 | 17.404593000 | 4.299627000 | 11.176380000 |
| 8 | 17.369251000 | 5.290912000 | 11.913814000 |
| 1 | 15.156528000 | 5.548516000 | 9.300800000 |
| 1 | 17.126634000 | 3.445274000 | 9.182990000 |
| 7 | 17.850040000 | 3.089814000 | 11.580353000 |

| | | | | | | | |
|---|---|---|---|---|---|---|---|
| 6 | 18.147893000 | 2.788950000 | 12.976270000 | 1 | 5.028256000 | -4.092273000 | 11.992455000 |
| 6 | 16.985979000 | 2.050987000 | 13.664112000 | 1 | 4.399126000 | -4.385198000 | 13.619153000 |
| 6 | 15.708479000 | 2.887831000 | 13.677902000 | 1 | 3.795087000 | -3.021197000 | 12.651282000 |
| 16 | 14.194859000 | 2.020073000 | 14.242562000 | 7 | 8.493217000 | -4.319768000 | 20.615754000 |
| 6 | 14.039645000 | 0.694422000 | 12.992057000 | 6 | 9.498956000 | -3.866960000 | 21.581930000 |
| 1 | 17.634019000 | 2.308846000 | 10.973831000 | 6 | 10.654091000 | -3.123902000 | 20.887318000 |
| 1 | 18.355514000 | 3.742993000 | 13.465159000 | 6 | 10.160634000 | -1.923961000 | 20.142059000 |
| 1 | 16.806347000 | 1.108456000 | 13.133647000 | 7 | 10.750851000 | -1.416463000 | 18.997978000 |
| 1 | 17.284914000 | 1.787463000 | 14.685772000 | 6 | 9.017829000 | -1.182795000 | 20.320059000 |
| 1 | 15.493024000 | 3.258114000 | 12.676766000 | 6 | 9.967348000 | -0.431136000 | 18.516991000 |
| 1 | 15.815314000 | 3.751396000 | 14.342275000 | 7 | 8.894257000 | -0.271756000 | 19.288011000 |
| 1 | 14.845213000 | -0.036483000 | 13.089987000 | 1 | 7.603849000 | -4.510973000 | 21.074482000 |
| 1 | 13.091448000 | 0.191413000 | 13.191572000 | 1 | 9.924102000 | -4.677990000 | 22.193303000 |
| 1 | 14.053072000 | 1.139021000 | 11.994757000 | 1 | 11.155491000 | -3.802983000 | 20.185376000 |
| 6 | 12.746845000 | 5.699430000 | 19.572831000 | 1 | 11.406490000 | -2.851112000 | 21.639948000 |
| 6 | 11.503291000 | 4.876500000 | 19.885391000 | 1 | 8.247987000 | -1.286575000 | 21.066092000 |
| 7 | 10.728537000 | 4.621264000 | 18.638285000 | 1 | 10.177606000 | 0.116921000 | 17.614856000 |
| 1 | 12.474348000 | 6.649285000 | 19.103523000 | 1 | 11.612125000 | -1.710371000 | 18.488529000 |
| 1 | 13.403339000 | 5.162868000 | 18.881107000 | 7 | 9.802441000 | 1.325960000 | 11.238560000 |
| 1 | 10.840198000 | 5.397011000 | 20.581831000 | 6 | 8.566465000 | 0.589745000 | 11.508785000 |
| 1 | 11.755004000 | 3.905013000 | 20.320465000 | 6 | 7.361413000 | 1.489332000 | 11.220619000 |
| 1 | 11.317176000 | 4.105110000 | 17.953333000 | 8 | 7.349898000 | 2.680910000 | 11.533202000 |
| 1 | 10.421157000 | 5.533898000 | 18.228432000 | 6 | 8.435182000 | 0.204056000 | 13.008899000 |
| 1 | 9.894185000 | 4.014253000 | 18.806945000 | 6 | 9.693214000 | -0.123041000 | 13.747507000 |
| 6 | 5.526341000 | -2.923512000 | 23.415077000 | 7 | 10.276589000 | 0.827507000 | 14.556553000 |
| 6 | 4.863534000 | -3.555914000 | 22.197303000 | 6 | 10.437469000 | -1.270338000 | 13.961464000 |
| 8 | 5.495387000 | -4.091874000 | 21.285568000 | 6 | 11.292727000 | 0.243917000 | 15.230567000 |
| 6 | 5.173915000 | -1.438320000 | 23.662192000 | 7 | 11.430771000 | -1.032760000 | 14.893067000 |
| 6 | 5.706774000 | -0.525440000 | 22.600671000 | 1 | 9.842518000 | 1.533363000 | 10.237725000 |
| 7 | 6.469911000 | 0.599424000 | 22.885717000 | 1 | 8.495755000 | -0.320448000 | 10.893658000 |
| 6 | 5.649116000 | -0.569547000 | 21.228903000 | 1 | 7.961317000 | 1.037167000 | 13.534758000 |
| 6 | 6.841459000 | 1.177063000 | 21.713163000 | 1 | 7.742251000 | -0.637055000 | 13.079614000 |
| 7 | 6.369247000 | 0.481541000 | 20.694132000 | 1 | 10.301239000 | -2.248829000 | 13.522663000 |
| 1 | 5.238680000 | -3.507600000 | 24.298305000 | 1 | 11.891290000 | 0.778542000 | 15.950083000 |
| 1 | 4.087614000 | -1.313888000 | 23.768437000 | 1 | 9.953481000 | 1.794393000 | 14.715201000 |
| 1 | 5.595464000 | -1.151875000 | 24.632350000 | 7 | 6.292261000 | 0.864403000 | 10.667424000 |
| 1 | 5.138032000 | -1.279332000 | 20.601502000 | 6 | 4.985049000 | 1.488707000 | 10.641252000 |
| 1 | 7.440174000 | 2.071207000 | 21.628513000 | 6 | 3.917705000 | 0.410126000 | 10.755712000 |
| 1 | 6.693603000 | 0.943063000 | 23.809535000 | 8 | 4.206658000 | -0.768184000 | 10.947664000 |
| 7 | 3.501491000 | -3.546705000 | 22.196081000 | 1 | 6.270414000 | -0.149772000 | 10.615100000 |
| 6 | 2.764525000 | -4.113530000 | 21.095413000 | 1 | 4.890354000 | 2.155084000 | 11.505672000 |
| 6 | 2.748893000 | -3.203619000 | 19.856797000 | 7 | 2.636022000 | 0.874527000 | 10.658199000 |
| 8 | 2.881677000 | -1.982511000 | 19.958876000 | 6 | 1.550832000 | 0.179554000 | 11.339865000 |
| 1 | 3.003256000 | -2.996356000 | 22.879318000 | 6 | 1.236841000 | 0.823962000 | 12.721562000 |
| 1 | 3.198923000 | -5.085914000 | 20.845524000 | 6 | 2.504936000 | 1.247311000 | 13.429818000 |
| 7 | 2.517763000 | -3.853154000 | 18.693439000 | 6 | 3.449802000 | 0.295031000 | 13.848214000 |
| 6 | 2.406843000 | -3.199855000 | 17.393646000 | 6 | 2.877885000 | 2.599631000 | 13.493307000 |
| 6 | 3.718755000 | -3.243321000 | 16.588533000 | 6 | 4.717779000 | 0.670767000 | 14.269016000 |
| 6 | 4.852406000 | -2.580966000 | 17.297905000 | 6 | 4.145320000 | 2.996945000 | 13.926416000 |
| 7 | 5.704522000 | -3.249382000 | 18.167428000 | 6 | 5.087807000 | 2.024585000 | 14.291913000 |
| 6 | 5.262359000 | -1.274481000 | 17.342323000 | 8 | 6.345702000 | 2.315644000 | 14.660019000 |
| 6 | 6.584928000 | -2.366250000 | 18.692332000 | 1 | 2.541969000 | 1.880602000 | 10.583768000 |
| 7 | 6.329199000 | -1.152456000 | 18.215236000 | 1 | 1.887161000 | -0.851119000 | 11.461789000 |
| 1 | 2.449932000 | -4.862183000 | 18.719597000 | 1 | 0.591590000 | 1.699032000 | 12.575990000 |
| 1 | 2.124291000 | -2.161573000 | 17.564537000 | 1 | 0.655895000 | 0.098522000 | 13.306784000 |
| 1 | 3.543860000 | -2.739953000 | 15.632263000 | 1 | 3.208477000 | -0.762911000 | 13.783367000 |
| 1 | 3.976493000 | -4.281512000 | 16.344795000 | 1 | 2.170420000 | 3.362387000 | 13.172033000 |
| 1 | 4.882205000 | -0.419449000 | 16.809965000 | 1 | 5.455249000 | -0.064350000 | 14.561845000 |
| 1 | 7.371427000 | -2.670108000 | 19.371823000 | 1 | 4.414253000 | 4.045734000 | 13.973606000 |
| 1 | 5.660278000 | -4.223150000 | 18.430781000 | 1 | 6.497198000 | 3.296559000 | 14.732286000 |
| 6 | 7.127116000 | -3.379463000 | 13.722675000 | 6 | 3.244942000 | 7.859091000 | 13.440562000 |
| 6 | 5.780085000 | -2.686708000 | 13.473579000 | 6 | 4.336407000 | 7.124138000 | 14.227539000 |
| 6 | 7.079117000 | -4.468332000 | 14.798982000 | 6 | 5.395754000 | 8.117946000 | 14.717751000 |
| 6 | 4.687877000 | -3.599857000 | 12.910766000 | 6 | 3.729688000 | 6.332995000 | 15.392114000 |
| 1 | 7.490698000 | -3.811655000 | 12.779571000 | 1 | 3.670599000 | 8.445147000 | 12.619014000 |
| 1 | 5.443311000 | -2.239555000 | 14.418370000 | 1 | 2.697777000 | 8.551826000 | 14.091566000 |
| 1 | 5.912885000 | -1.852286000 | 12.774719000 | 1 | 4.837322000 | 6.417103000 | 13.549933000 |
| 1 | 8.067278000 | -4.916197000 | 14.951188000 | 1 | 5.892552000 | 8.613373000 | 13.874646000 |
| 1 | 6.389281000 | -5.276894000 | 14.531400000 | 1 | 4.937495000 | 8.898292000 | 15.338854000 |
| 1 | 6.755573000 | -4.045159000 | 15.757503000 | 1 | 6.162498000 | 7.620840000 | 15.317240000 |

| | | | |
|---|---|---|---|
| 1 | 4.470773000 | 5.699209000 | 15.886273000 |
| 1 | 3.301277000 | 7.013726000 | 16.139216000 |
| 1 | 2.919756000 | 5.676956000 | 15.048009000 |
| 6 | 0.384181000 | -0.273226000 | 18.581925000 |
| 7 | 1.160638000 | 0.251528000 | 17.452351000 |
| 6 | 2.304078000 | 0.976908000 | 17.640792000 |
| 7 | 3.072477000 | 0.760571000 | 18.715127000 |
| 7 | 2.639625000 | 1.903207000 | 16.755382000 |
| 1 | 1.006327000 | -0.912345000 | 19.208135000 |
| 1 | -0.037360000 | 0.531129000 | 19.196937000 |
| 1 | 0.587213000 | 0.609597000 | 16.698580000 |
| 1 | 2.964778000 | -0.117057000 | 19.218090000 |
| 1 | 4.036985000 | 1.112774000 | 18.640604000 |
| 1 | 3.510590000 | 2.470334000 | 16.867914000 |
| 1 | 2.157815000 | 1.973767000 | 15.867464000 |
| 26 | 7.238165000 | 0.638083000 | 18.645983000 |
| 6 | 7.542772000 | 5.405864000 | 15.907025000 |
| 6 | 7.233248000 | 4.911768000 | 17.315283000 |
| 6 | 7.216114000 | 3.383318000 | 17.460349000 |
| 6 | 5.810438000 | 2.793808000 | 17.475158000 |
| 8 | 6.974879000 | 4.871281000 | 14.916672000 |
| 8 | 8.339705000 | 6.396431000 | 15.794973000 |
| 8 | 4.837283000 | 3.462482000 | 17.092281000 |
| 8 | 5.685350000 | 1.569386000 | 17.894640000 |
| 16 | 8.110403000 | 2.758488000 | 18.955265000 |
| 1 | 7.952411000 | 5.357952000 | 18.002341000 |
| 1 | 6.244460000 | 5.304093000 | 17.579334000 |
| 1 | 7.727366000 | 2.933818000 | 16.606304000 |
| 8 | 8.072397000 | 0.606603000 | 16.658406000 |
| 8 | 12.840766000 | -1.742975000 | 17.266726000 |
| 1 | 13.315013000 | -0.888975000 | 17.283252000 |
| 1 | 12.411729000 | -1.757655000 | 16.375789000 |
| 8 | 7.828275000 | 5.853790000 | 12.603169000 |
| 1 | 7.474862000 | 5.483113000 | 13.464562000 |
| 1 | 7.160578000 | 5.655504000 | 11.921484000 |
| 1 | 13.302172000 | 5.905352000 | 20.492631000 |
| 1 | 16.250920000 | 8.232302000 | 14.455025000 |
| 1 | 9.005819000 | -3.180022000 | 22.280891000 |
| 1 | 8.802235000 | -5.179726000 | 20.166991000 |
| 1 | 6.605078000 | -3.029298000 | 23.280989000 |
| 1 | 1.729664000 | -4.277052000 | 21.413622000 |
| 1 | 1.602308000 | -3.685164000 | 16.831531000 |
| 1 | -0.428908000 | -0.878975000 | 18.176977000 |
| 8 | 7.589691000 | -0.221896000 | 15.860419000 |
| 1 | 2.508140000 | 7.153879000 | 13.029522000 |
| 1 | 0.655013000 | 0.169840000 | 10.711192000 |
| 1 | 4.857098000 | 2.100956000 | 9.739976000 |
| 1 | 10.588940000 | 0.720634000 | 11.469769000 |
| 1 | 6.148317000 | 2.362707000 | 7.199939000 |
| 1 | 11.693813000 | 3.617137000 | 6.029008000 |
| 1 | 17.434452000 | 5.194462000 | 9.228858000 |
| 1 | 19.055969000 | 2.179141000 | 13.020391000 |
| 1 | 16.518208000 | 8.022665000 | 9.569880000 |
| 1 | 11.781093000 | 8.633156000 | 8.480993000 |
| 1 | 7.863605000 | -2.626422000 | 14.028645000 |
| 8 | 9.938109000 | 3.494438000 | 15.176353000 |
| 1 | 10.791838000 | 3.454133000 | 15.652745000 |
| 1 | 10.133949000 | 3.878959000 | 14.298584000 |
| 8 | 12.278078000 | 3.123048000 | 16.886940000 |
| 1 | 12.724689000 | 3.925749000 | 16.534502000 |
| 1 | 12.947475000 | 2.439223000 | 17.110980000 |
| 8 | 9.809274000 | 7.152095000 | 17.867607000 |
| 1 | 9.217663000 | 7.534389000 | 18.534096000 |
| 1 | 9.212697000 | 6.905748000 | 17.104778000 |
| 8 | 14.030653000 | 0.950177000 | 17.203364000 |
| 1 | 14.830100000 | 0.979438000 | 17.753744000 |
| 1 | 14.339421000 | 1.078398000 | 16.271809000 |

(11) Frisch, M. J. T., G. W.; Schlegel, H. B.; Scuseria, G. E.; Robb, M. A.; Cheeseman, J. R.; Scalmani, G.; Barone, V.; Mennucci, B.; Petersson, G. A.; Nakatsuji, H.; Caricato, M.; Li, X.; Hratchian, H. P.; Izmaylov, A. F.; Bloino, J.; Zheng, G.; Sonnenberg, J. L.; Hada, M.; Ehara, M.; Toyota, K.; Fukuda, R.; Hasegawa, J.; Ishida, M.; Nakajima, T.; Honda, Y.; Kitao, O.; Nakai, H.; Vreven, T.; Montgomery Jr, J. A.; Peralta, J. E.; Ogliaro, F.; Bearpark, M.; Heyd, J. J.; Brothers, E.; Kudin, K. N.; Staroverov, V. N.; Keith, T.; Kobayashi, R.; Normand, J.; Raghavachari, K.; Rendell, A.; Burant, J. C.; Iyengar, S. S.; Tomasi, J.; Cossi, M.; Rega, N.;. Millam, J. M.; Klene, M.; Knox, J. E.; Cross, J. B.; Bakken, V.; Adamo, C.; Jaramillo, J.; Gomperts, R.; Stratmann, R. E.; Yazyev, O.; Austin, A. J.; Cammi, R.; Pomelli, C.; Ochterski, J. W.; Martin, R. L.; Morokuma, K.; Zakrzewski, V. G.; Voth, G. A.; Salvador, P.; Dannenberg, J. J.; Dapprich, S.; Daniels, A. D.; Farkas, O.; Foresman, J.

B.; Ortiz, J. V.; Cioslowski, J.; Fox, D. J. . Gaussian-09, Revision D.01. Gaussian, I., Ed.; Wallingford CT, 2013.
(12) Lee, C. T.; Yang, W. T.; Parr, R. G. Development of the Colle-Salvetti correlation-energy formula into a functional of the electron-density. *Phys Rev B* **1988**, *37* (2), 785-789.
(13) Kendall, R. A.; Dunning, T. H.; Harrison, R. J. Electron-affinities of the 1st-row atoms revisited - systematic basis-sets and wave-functions. *J Chem Phys* **1992**, *96* (9), 6796-6806.
(14) Trag, J.; Zahn, D. Improved GAFF2 parameters for fluorinated alkanes and mixed hydro- and fluorocarbons. *J Mol Model* **2019**, *25* (2), 39.
(15) Maier, J. A.; Martinez, C.; Kasavajhala, K.; Wickstrom, L.; Hauser, K. E.; Simmerling, C. ff14SB: Improving the accuracy of protein side chain and backbone parameters from ff99SB. *J Chem Theory Comput* **2015**, *11* (8), 3696-3713.
(16) Case, D. A.; Cheatham, T. E., 3rd; Darden, T.; Gohlke, H.; Luo, R.; Merz, K. M., Jr.; Onufriev, A.; Simmerling, C.; Wang, B.; Woods, R. J. The Amber biomolecular simulation programs. *J Comput Chem* **2005**, *26* (16), 1668-1688.
(17) de Visser, S. P.; Wong, H. P. H.; Zhang, Y.; Yadav, R.; Sastri, C. V. Tutorial review on the set-up and running of quantum mechanical cluster models for enzymatic reaction mechanisms. *Chem Eur J* **2024**, *30* (60), e202402468.
(18) Grimme, S.; Antony, J.; Ehrlich, S.; Krieg, H. A consistent and accurate ab initio parametrization of density functional dispersion correction (DFT-D) for the 94 elements H-Pu. *J Chem Phys* **2010**, *132* (15), 154104.
(19) Hay, P. J.; Wadt, W. R. Abinitio effective core potentials for molecular calculations - potentials for the transition-metal atoms Sc to Hg. *J Chem Phys* **1985**, *82* (1), 270-283.
(20) Francl, M. M.; Pietro, W. J.; Hehre, W. J.; Binkley, J. S.; Gordon, M. S.; Defrees, D. J.; Pople, J. A. Self-consistent molecular-orbital methods. 23. A polarization-type basis set for 2nd-row elements. *J Chem Phys* **1982**, *77* (7), 3654-3665.
